## Supplementary Files 20212102 for "A candidate sex determination locus in amphibians, evolved by structural variation between X- and Y-chromosomes": SupplFile_new2_female-male-assembly-comparison.pptx

### Slide 1
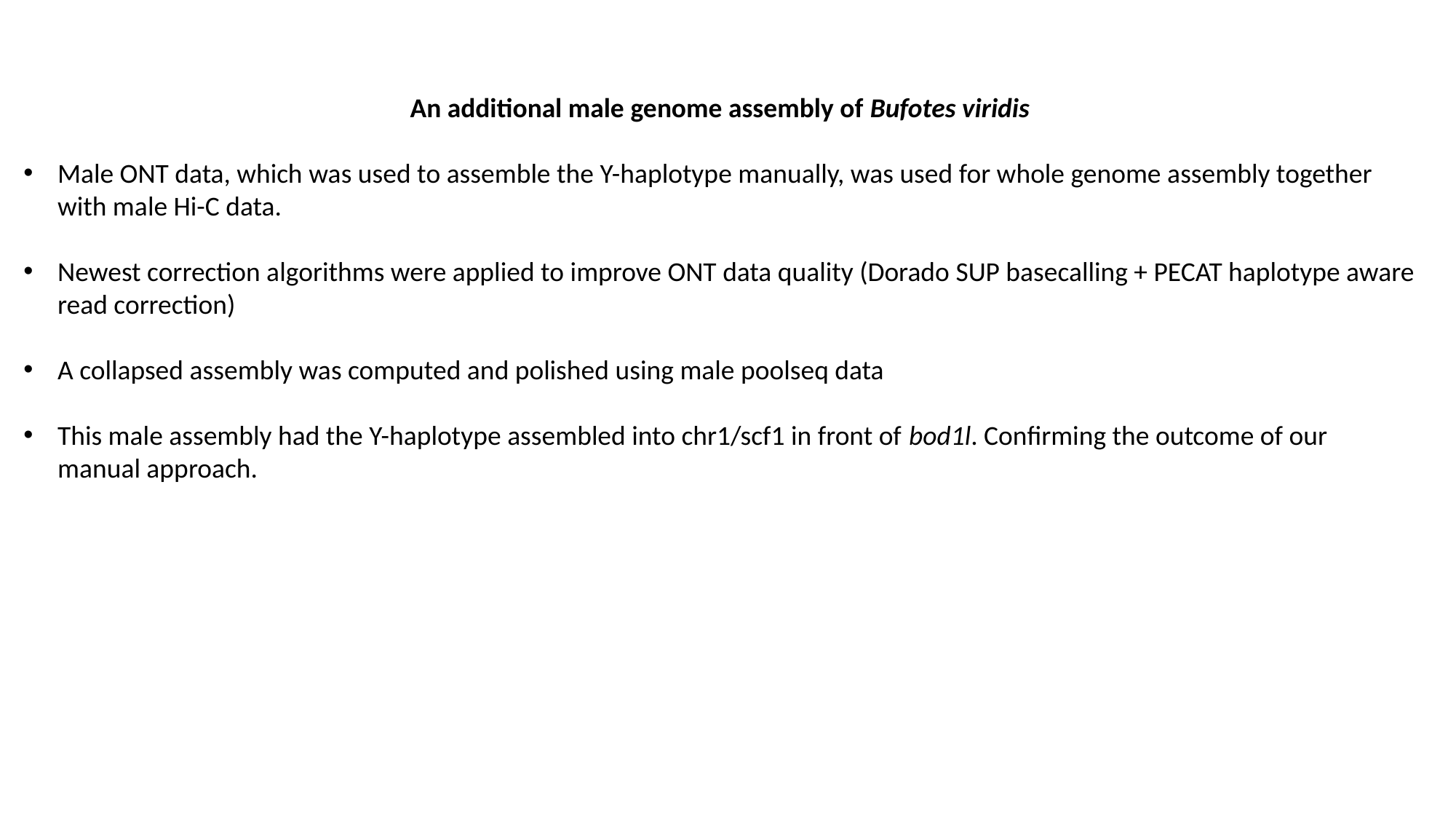

An additional male genome assembly of Bufotes viridis
Male ONT data, which was used to assemble the Y-haplotype manually, was used for whole genome assembly together with male Hi-C data.
Newest correction algorithms were applied to improve ONT data quality (Dorado SUP basecalling + PECAT haplotype aware read correction)
A collapsed assembly was computed and polished using male poolseq data
This male assembly had the Y-haplotype assembled into chr1/scf1 in front of bod1l. Confirming the outcome of our manual approach.

### Slide 2
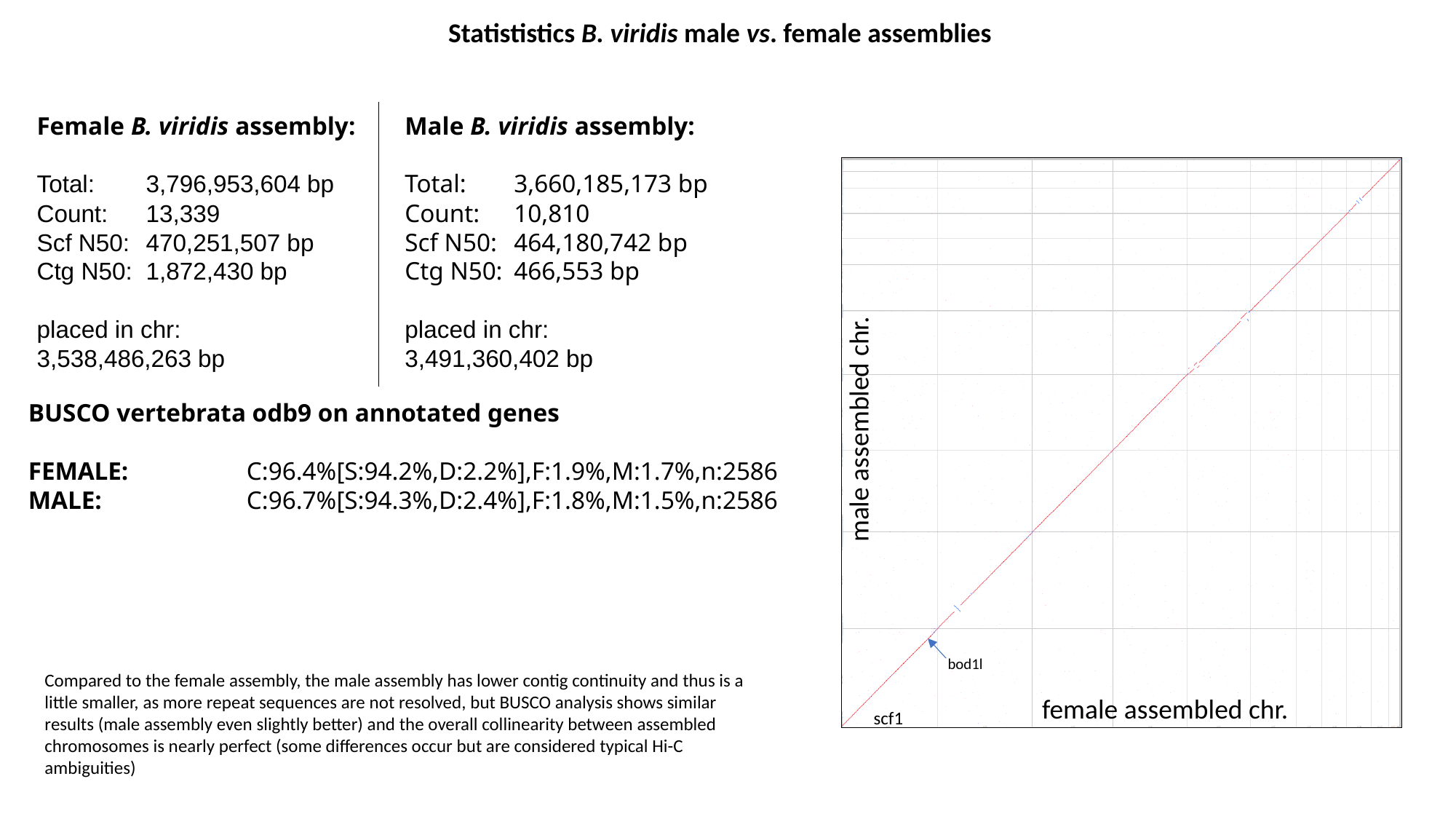

Statististics B. viridis male vs. female assemblies
Female B. viridis assembly:
Total: 	3,796,953,604 bp
Count: 	13,339
Scf N50:	470,251,507 bp
Ctg N50:	1,872,430 bp
placed in chr:
3,538,486,263 bp
Male B. viridis assembly:
Total: 	3,660,185,173 bp
Count: 	10,810
Scf N50:	464,180,742 bp
Ctg N50:	466,553 bp
placed in chr:
3,491,360,402 bp
BUSCO vertebrata odb9 on annotated genes
FEMALE:		C:96.4%[S:94.2%,D:2.2%],F:1.9%,M:1.7%,n:2586
MALE:		C:96.7%[S:94.3%,D:2.4%],F:1.8%,M:1.5%,n:2586
male assembled chr.
bod1l
Compared to the female assembly, the male assembly has lower contig continuity and thus is a little smaller, as more repeat sequences are not resolved, but BUSCO analysis shows similar results (male assembly even slightly better) and the overall collinearity between assembled chromosomes is nearly perfect (some differences occur but are considered typical Hi-C ambiguities)
female assembled chr.
scf1

### Slide 3
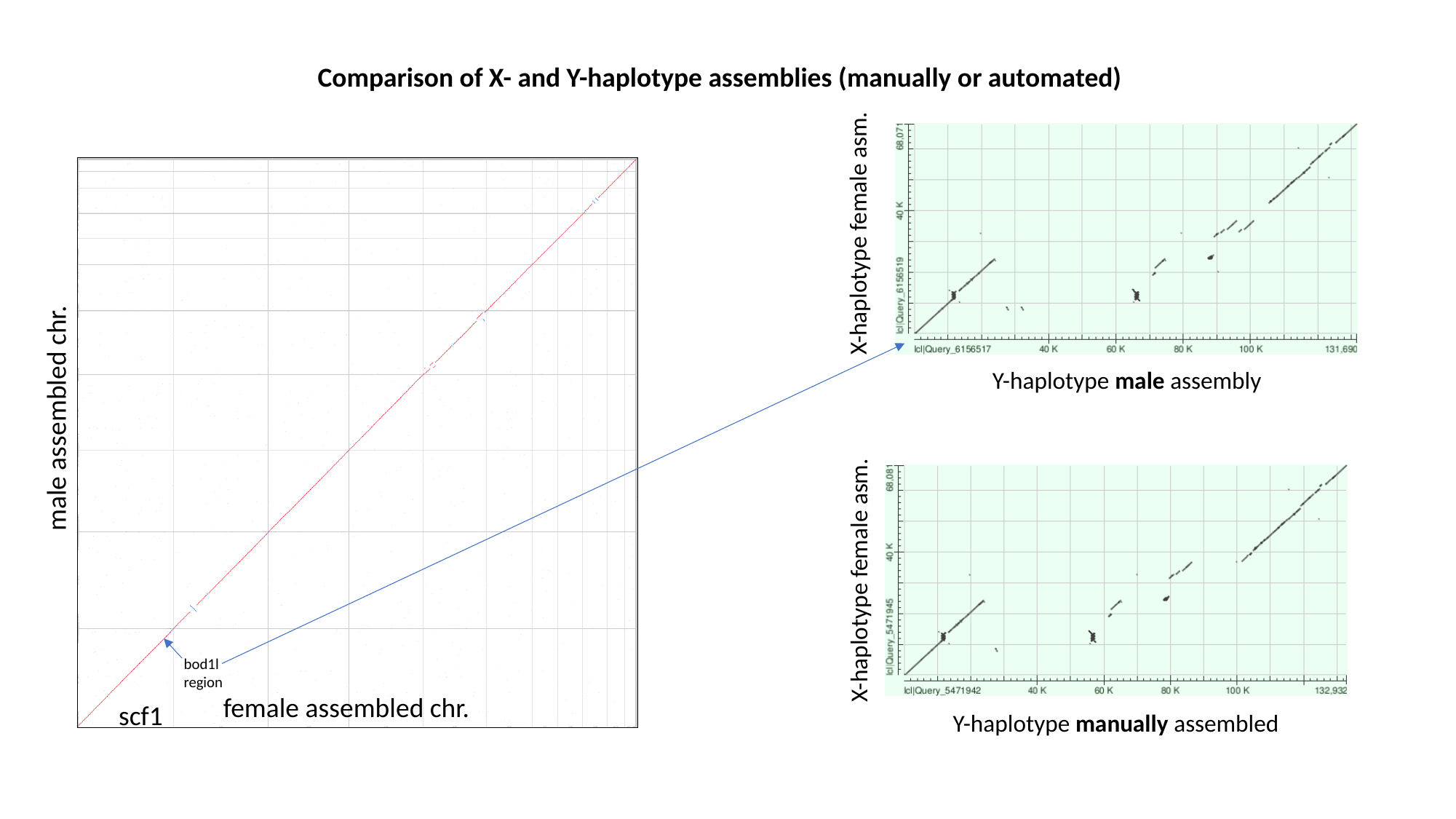

Comparison of X- and Y-haplotype assemblies (manually or automated)
X-haplotype female asm.
Y-haplotype male assembly
male assembled chr.
X-haplotype female asm.
bod1l region
female assembled chr.
scf1
Y-haplotype manually assembled

### Slide 4
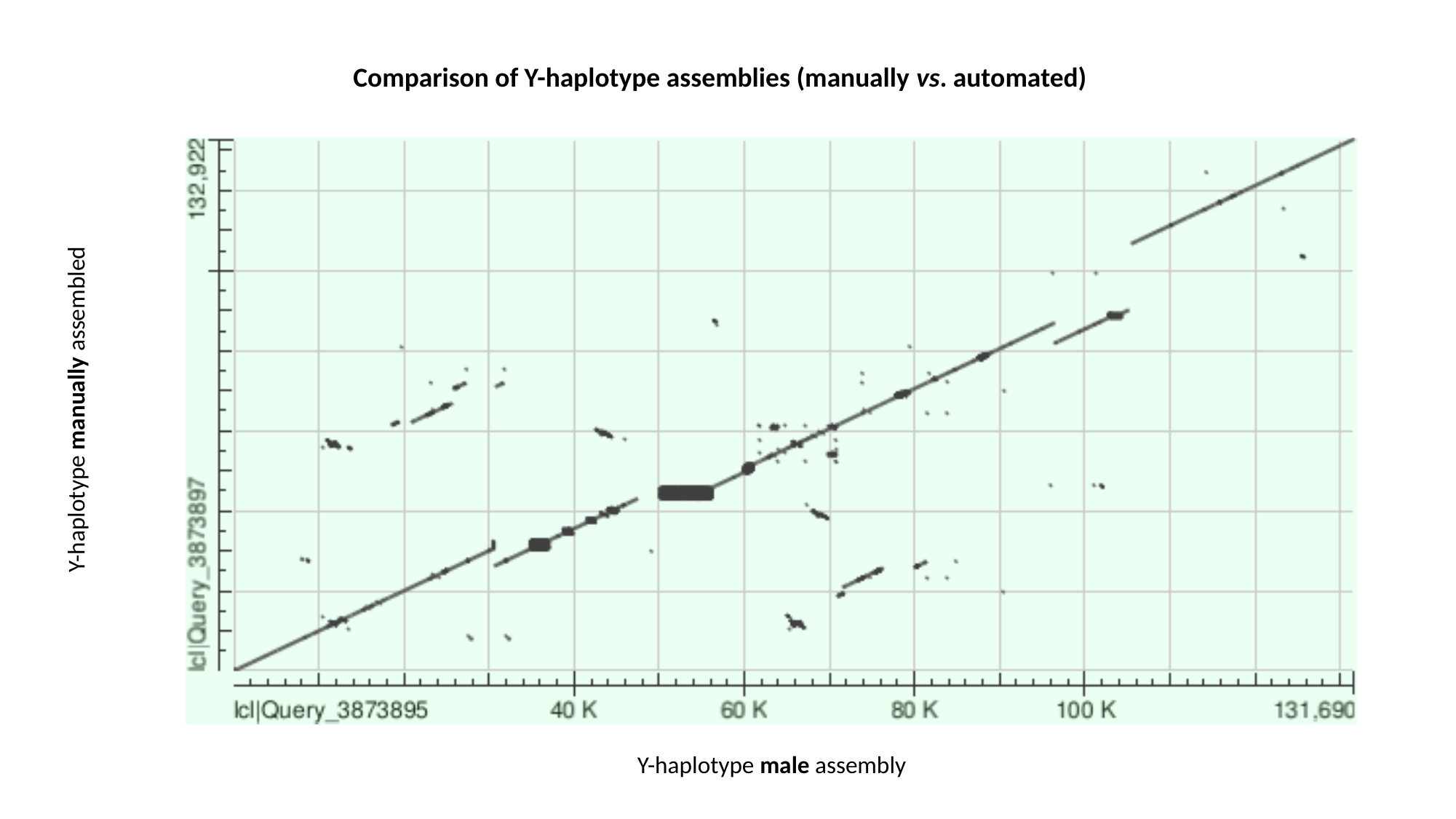

Comparison of Y-haplotype assemblies (manually vs. automated)
Y-haplotype manually assembled
Y-haplotype male assembly

### Slide 5
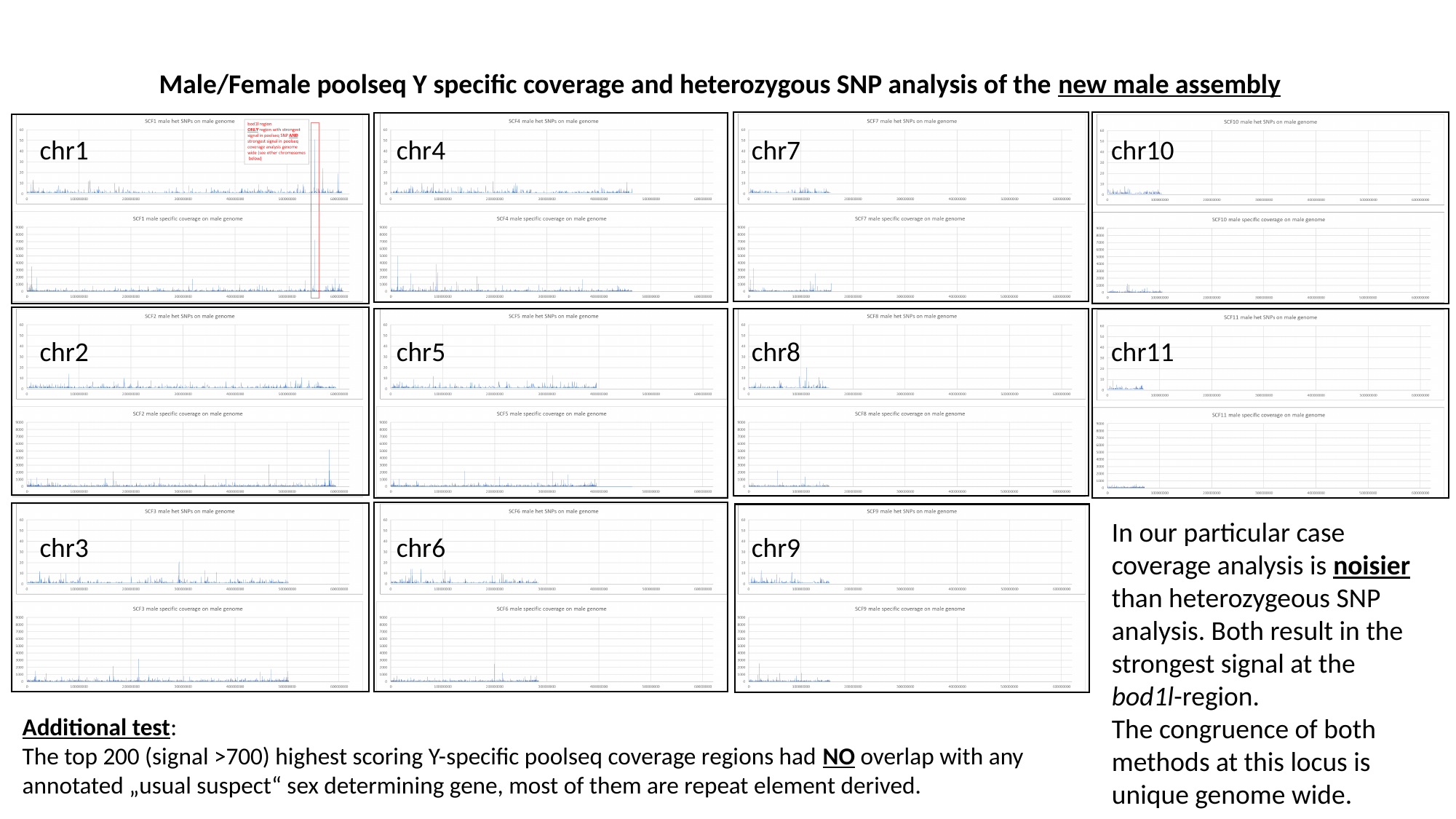

Male/Female poolseq Y specific coverage and heterozygous SNP analysis of the new male assembly
chr1
chr4
chr7
chr10
chr2
chr5
chr8
chr11
In our particular case coverage analysis is noisier than heterozygeous SNP analysis. Both result in the strongest signal at the bod1l-region.
The congruence of both methods at this locus is unique genome wide.
chr3
chr6
chr9
Additional test:
The top 200 (signal >700) highest scoring Y-specific poolseq coverage regions had NO overlap with any annotated „usual suspect“ sex determining gene, most of them are repeat element derived.

### Slide 6
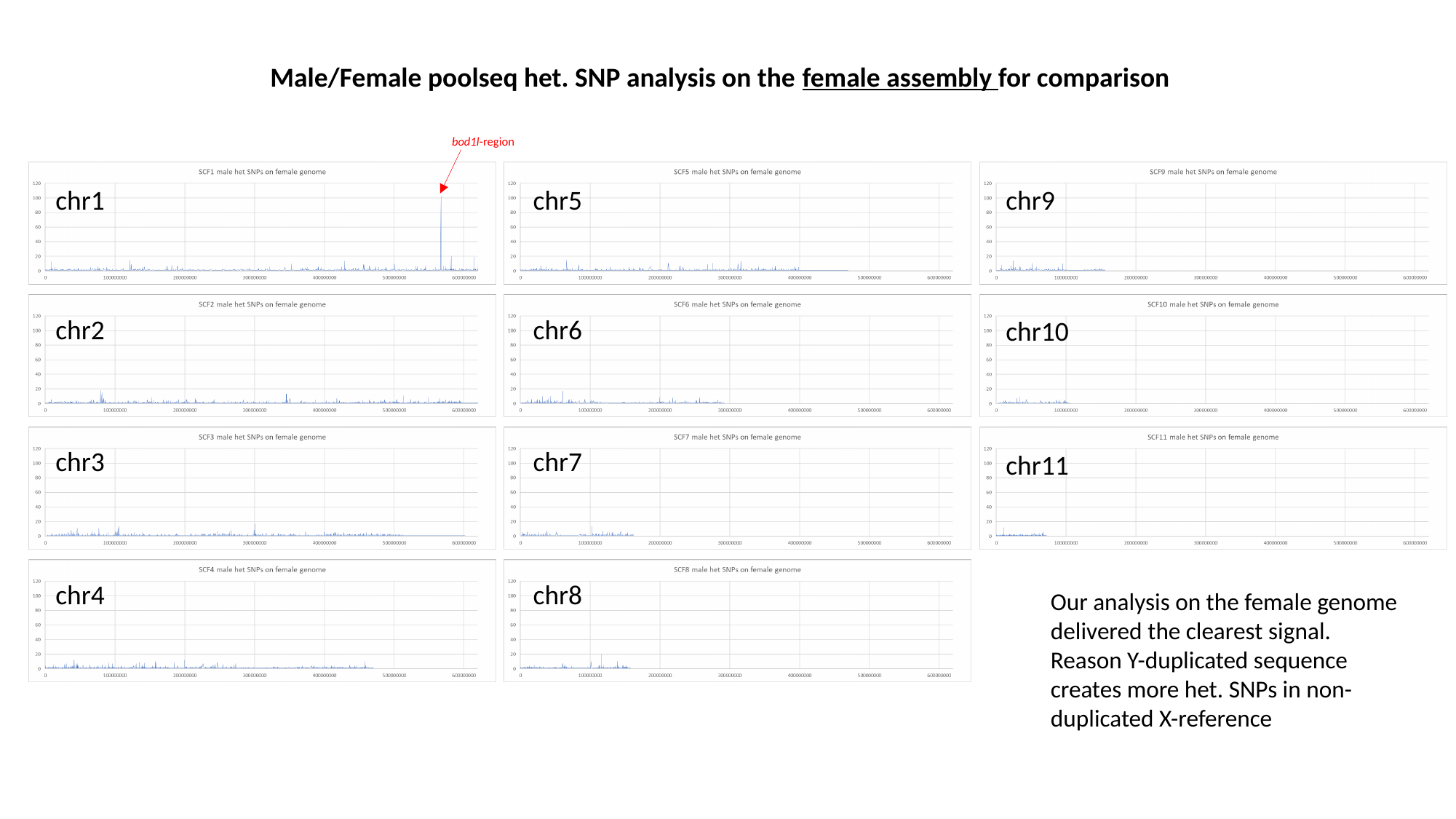

Male/Female poolseq het. SNP analysis on the female assembly for comparison
bod1l-region
chr1
chr5
chr9
chr2
chr6
chr10
chr3
chr7
chr11
chr4
chr8
Our analysis on the female genome delivered the clearest signal.
Reason Y-duplicated sequence creates more het. SNPs in non-duplicated X-reference
