## Supplementary Files 20212102 for "A candidate sex determination locus in amphibians, evolved by structural variation between X- and Y-chromosomes": SupplFile_new3.pdf

### Supplementary File S3

| sample no. | 1 | 2 | 3 | 4 | 5 | 6 | 7 | 8 | 9 | 10 | 11 | 12 | 13 | 14 | 15 | 16 | 17 | 18 | 19 | 20 | 21 | 22 | 23 | 24 | 25 | 26 | 27 | 28 | 29 | 30 | 31 | 32 | 33 | 34 | 35 | 36 | 37 | 38 | 39 | 40 | 41 | 42 | 43 | 44 | 45 | 46 | 47 | 48 | 49 |
|---|---|---|---|---|---|---|---|---|---|---|---|---|---|---|---|---|---|---|---|---|---|---|---|---|---|---|---|---|---|---|---|---|---|---|---|---|---|---|---|---|---|---|---|---|---|---|---|---|---|
| sex | F | F | F | F | F | F | F | F | F | F | F | F | F | F | F | F | F | F | F | F | F | F | F | F | F | F | F | F | M | M | M | M | M | M | M | M | M | M | M | M | M | M | M | M | M | M | M | M | M |
| scf1_566,831,998_G_A [AA609: Arginine to Lysine] | | | | | | | | | | | | | | | | | | | | | | | | | | | | | | | | | | | | | | | | | | | | | | | | | |
| scf1_566,832,081_T_G [AA637: Serine to Alanine] | | | | | | | | | | | | | | | | | | | | | | | | | | | | | | | | | | | | | | | | | | | | | | | | | |
| scf1_566,835,326_A_G [AA1718: Lysine to Glutamic acid] | | | | | | | | | | | | | | | | | | | | | | | | | | | | | | | | | | | | | | | | | | | | | | | | | |
| scf1_566,835,600_G_T [AA1810: Alanine to Serine] | | | | | | | | | | | | | | | | | | | | | | | | | | | | | | | | | | | | | | | | | | | | | | | | | |
| scf1_566,836,519_C_T [AA2116: Serine to Phenylalanine] | | | | | | | | | | | | | | | | | | | | | | | | | | | | | | | | | | | | | | | | | | | | | | | | | |
| scf1_566,836,636_A_G [AA2155: Histidine to Arginine] | | | | | | | | | | | | | | | | | | | | | | | | | | | | | | | | | | | | | | | | | | | | | | | | | |

homozygous reference | heterozygous | homozygous alternative

| sample no. | sample name short | sample name full |
|---|---|---|
| 1 | F_10_GR14_5x6-1:ref | F_10_GR14_5x6.1,/S32_1/S32_2 |
| 2 | F_10_GR14_5x6-2:ref | F_10_GR14_5x6.2,/S33_1/S33_2 |
| 3 | F_10_GR14_5x6-3:ref | F_10_GR14_5x6.3,/S36_1/S36_2 |
| 4 | F_10_GR14_5x6-4:ref | F_10_GR14_5x6.4,/S38_1/S38_2.fq.gz |
| 5 | F_15_GR14_5x6-1:ref | F_15_GR14_5x6.1,/S40_1/S40_2. |
| 6 | F_15_GR14_5x6-2:ref | F_15_GR14_5x6.2,/S41_1/S41_2 |
| 7 | F_15_GR14_5x6-3:ref | F_15_GR14_5x6.3,/S42_1/S42_2 |
| 8 | F_15_GR14_5x6-4:ref | F_15_GR14_5x6.4,/S45_1/S45_2. |
| 9 | F_adult_ADULT-1:ref | F_adult_ADULT.1,/S30_1/S30_2. |
| 10 | F_ca18_GR14_1x2-1:ref | F_ca18_GR14_1x2.1,/S2_1/S2_2. |
| 11 | F_ca18_GR14_1x2-2:ref | F_ca18_GR14_1x2.2,/S4_1/S4_2 |
| 12 | F_ca18_GR14_1x2-3:ref | F_ca18_GR14_1x2.3,/S6_1/S6_2 |
| 13 | F_ca18_GR14_1x2-4:ref | F_ca18_GR14_1x2.4,/S7_1/S7_2 |
| 14 | F_ca34_GR14_1x2-1:ref | F_ca34_GR14_1x2.1,/S13_1/S13_2 |
| 15 | F_ca34_GR14_1x2-2:ref | F_ca34_GR14_1x2.2,/S14_1/S14_2 |
| 16 | F_ca34_GR14_1x2-3:ref | F_ca34_GR14_1x2.3,/S12_1/S12_2 |
| 17 | F_ca34_GR14_1x2-4:ref | F_ca34_GR14_1x2.4,/S9_1/S9_2. |
| 18 | F_Gosner30_Z_2022-1:ref | F_Gosner30_Z_2022.1,/S22_1/S22_2 |
| 19 | F_Gosner30_Z_2022-2:ref | F_Gosner30_Z_2022.2,/S23_1/S23_2 |
| 20 | F_Gosner36-37_Z_2022-1:ref | F_Gosner36-37_Z_2022.1,/S25_1/S25_2 |
| 21 | F_Gosner38_Potamies-1:ref | F_Gosner38_Potamies.1,/S15_1/S15_2 |
| 22 | F_Gosner38_Potamies-2:ref | F_Gosner38_Potamies.2,/S16_1/S16_2 |
| 23 | F_Gosner38_Potamies-3:ref | F_Gosner38_Potamies.3,/S19_1/S19_2 |
| 24 | F_Gosner38_Potamies-4:ref | F_Gosner38_Potamies.4,/S20_1/S20_2 |
| 25 | F_Gosner43-44_Z_2022-1:ref | F_Gosner43-44_Z_2022.1,/S47_1/S47_2 |
| 26 | F_Gosner43-44_Z_2022-2:ref | F_Gosner43-44_Z_2022.2,/S50_1/S50 |
| 27 | F_Gosner43-44_Z_2022-3:ref | F_Gosner43-44_Z_2022.3,/S51_1/S51_2 |
| 28 | F_Gosner43-44_Z_2022-4:ref | F_Gosner43-44_Z_2022.4,/S52_1/S52_2 |

| sample no. | sample name short | sample name full |
|---|---|---|
| 29 | M_10_GR14_5x6-1:ref | M_10_GR14_5x6.1,/S31_1/S31_ |
| 30 | M_10_GR14_5x6-2:ref | M_10_GR14_5x6.2,/S35_1/S35_2 |
| 31 | M_10_GR14_5x6-3:ref | M_10_GR14_5x6.3,/S37_1/S37_2 |
| 32 | M_15_GR14_5x6-1:ref | M_15_GR14_5x6.1,/S39_1/S39_2 |
| 33 | M_15_GR14_5x6-2:ref | M_15_GR14_5x6.2,/S46_1/S46_2 |
| 34 | M_ca18_GR14_1x2-1:ref | M_ca18_GR14_1x2.1,/S1_1/S1_2 |
| 35 | M_ca18_GR14_1x2-2:ref | M_ca18_GR14_1x2.2,/S3_1/S3_2 |
| 36 | M_ca18_GR14_1x2-3:ref | M_ca18_GR14_1x2.3,/S5_1/S5_2 |
| 37 | M_ca18_GR14_1x2-4:ref | M_ca18_GR14_1x2.4,/S8_1/S8_2 |
| 38 | M_ca34_GR14_1x2-1:ref | M_ca34_GR14_1x2.1,/S10_1/S10_2 |
| 39 | M_ca34_GR14_1x2-2:ref | M_ca34_GR14_1x2.2,/S11_1/S11_2 |
| 40 | M_Gosner30_Z_2022-1:ref | M_Gosner30_Z_2022.1,/S21_1/S21_2 |
| 41 | M_Gosner30_Z_2022-2:ref | M_Gosner30_Z_2022.2,/S24_1/S24_2 |
| 42 | M_Gosner36-37_Z_2022-1:ref | M_Gosner36-37_Z_2022.1,/S26_1/S26_2 |
| 43 | M_Gosner36-37_Z_2022-2:ref | M_Gosner36-37_Z_2022.2,/S27_1/S27_2 |
| 44 | M_Gosner36-37_Z_2022-3:ref | M_Gosner36-37_Z_2022.3,/S28_1/S28_2 |
| 45 | M_Gosner38_Potamies-1:ref | M_Gosner38_Potamies.1,/S17_1/S17_2 |
| 46 | M_Gosner38_Potamies-2:ref | M_Gosner38_Potamies.2,/S18_1/S18_2 |
| 47 | M_Gosner43-44_Z_2022-1:ref | M_Gosner43-44_Z_2022.1,/S48_1/S48_2 |
| 48 | M_Gosner43-44_Z_2022-2:ref | M_Gosner43-44_Z_2022.2,/S49_1/S49_2 |
| 49 | M_subadult_ADULT-1:ref | M_subadult_ADULT.1,/S29_1/S29_2 |
