## Supplementary Files 20212102 for "A candidate sex determination locus in amphibians, evolved by structural variation between X- and Y-chromosomes": SupplFile_new6_ExpressionUsualSuspects_27Sept2023.pdf

### Toad Transcriptome

09/27/2022

#### Toad RNA-Seq analysis

| | |
|---|---|
| Gosner 23-24 | Stage_1_females |
| Gosner 23-24 | Stage_1_males |
| Gosner 27-29 | Stage_2_females |
| Gosner 27-29 | Stage_2_males |
| Gosner 30-33 | Stage_3_females |
| Gosner 30-33 | Stage_3_males |
| Gosner 34-38 | Stage_4_females |
| Gosner 34-38 | Stage_4_males |
| Gosner 43-44 | Stage_5a_female |
| Gosner 43-44 | Stage_5a_males |
| Gosner 45-46 | Stage_5b_female |
| Gosner 45-46 | Stage_5b_males |
| Gosner 46 (6 weeks after metamorphosis) | Stage_5c_males |
| adult | Stage_6_females |
| subadult | Stage_6_males |

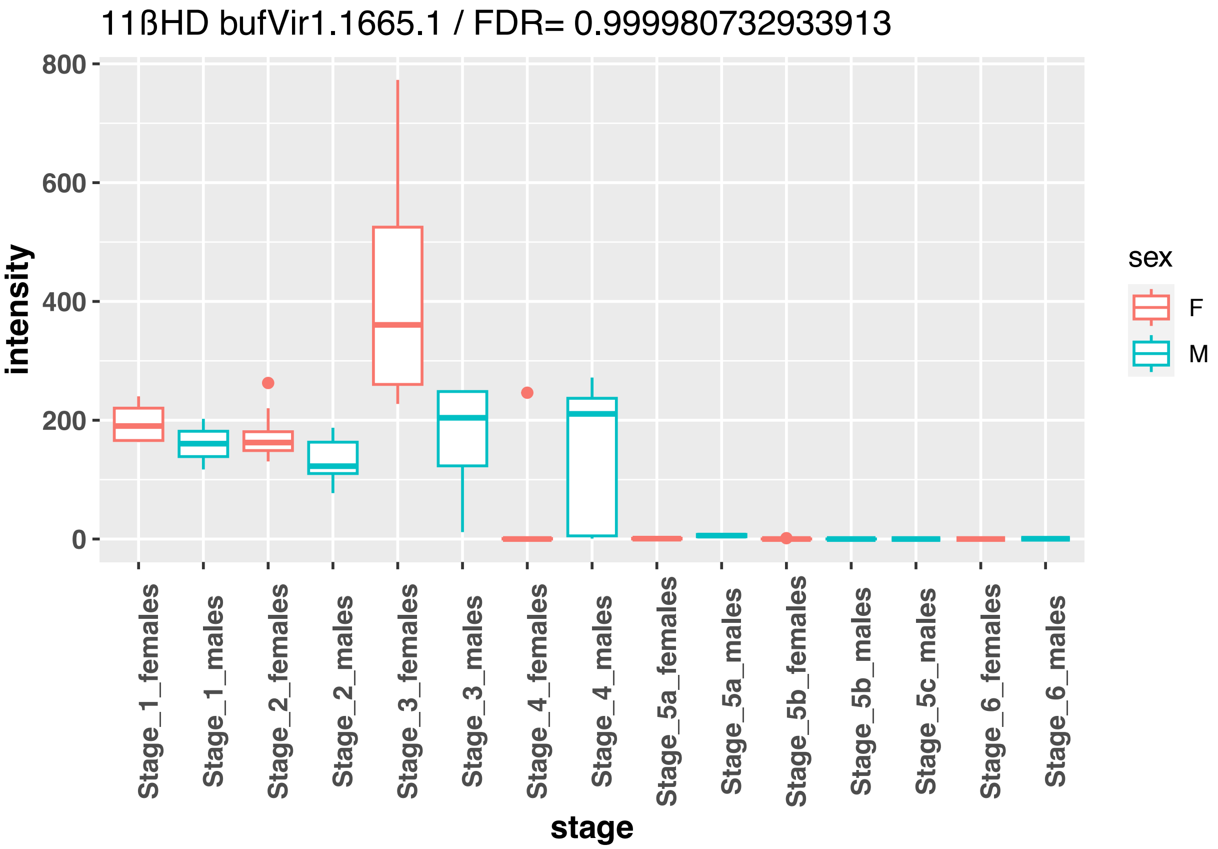
11ßHD bufVir1.1665.1 / FDR= 0.999980732933913
intensity
800
600
400
200
0
stage
Stage_1_females
Stage_1_males
Stage_2_females
Stage_2_males
Stage_3_females
Stage_3_males
Stage_4_females
Stage_4_males
Stage_5a_females
Stage_5a_males
Stage_5b_females
Stage_5b_males
Stage_5c_males
Stage_6_females
Stage_6_males
sex
F
M

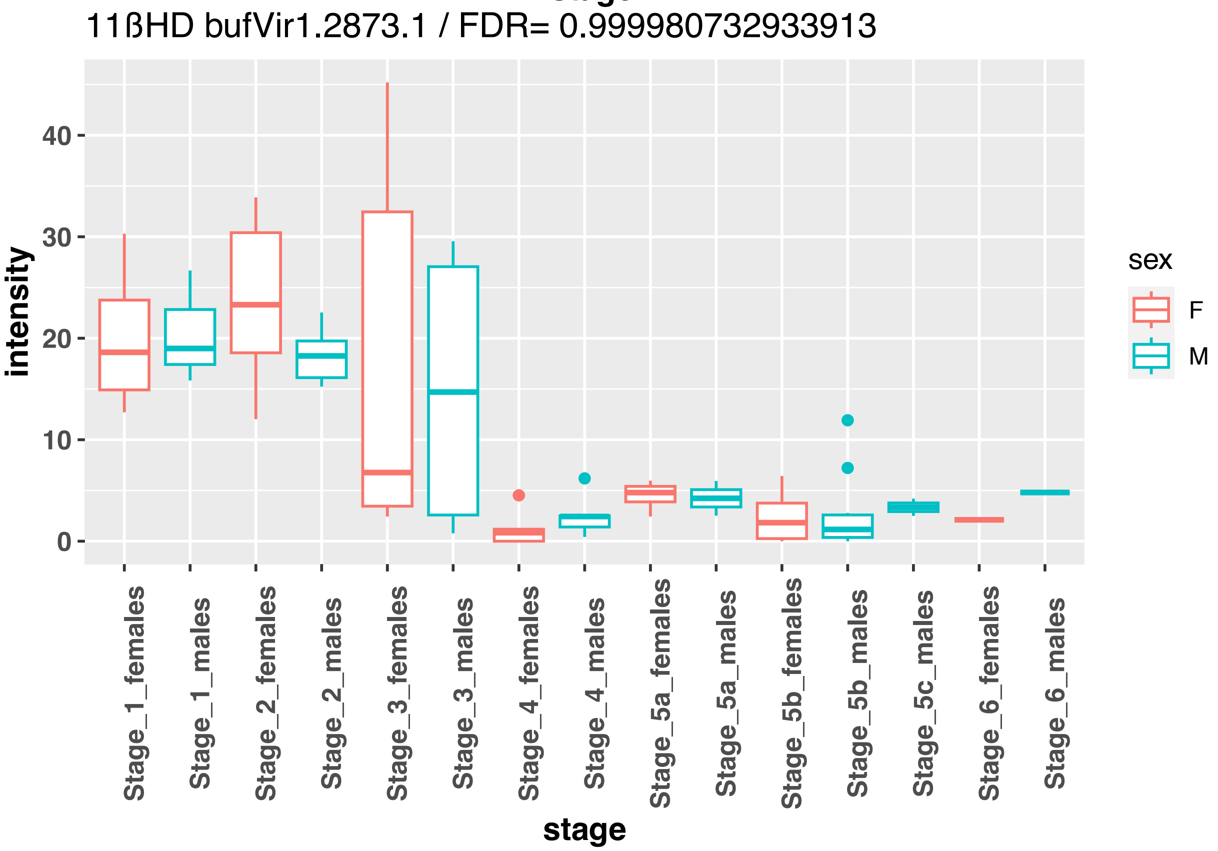
11ßHD bufVir1.2873.1 / FDR= 0.999980732933913
intensity
40
30
20
10
0
stage
Stage_1_females
Stage_1_males
Stage_2_females
Stage_2_males
Stage_3_females
Stage_3_males
Stage_4_females
Stage_4_males
Stage_5a_females
Stage_5a_males
Stage_5b_females
Stage_5b_males
Stage_5c_males
Stage_6_females
Stage_6_males
sex
F
M

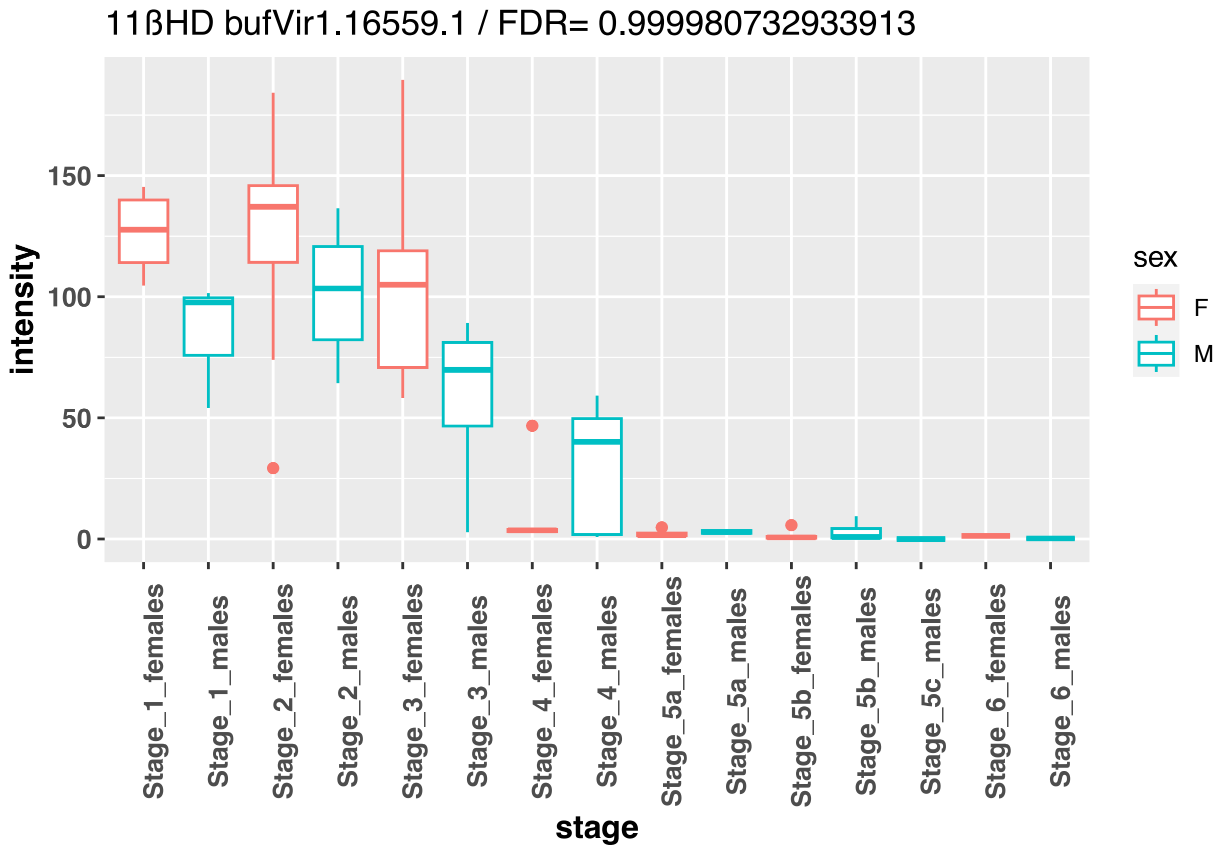
11ßHD bufVir1.16559.1 / FDR= 0.999980732933913
intensity
150
100
50
0
Stage_1_females
Stage_1_males
Stage_2_females
Stage_2_males
Stage_3_females
Stage_3_males
Stage_4_females
Stage_4_males
Stage_5a_females
Stage_5a_males
Stage_5b_females
Stage_5b_males
Stage_5c_males
Stage_6_females
Stage_6_males
stage
sex
F
M

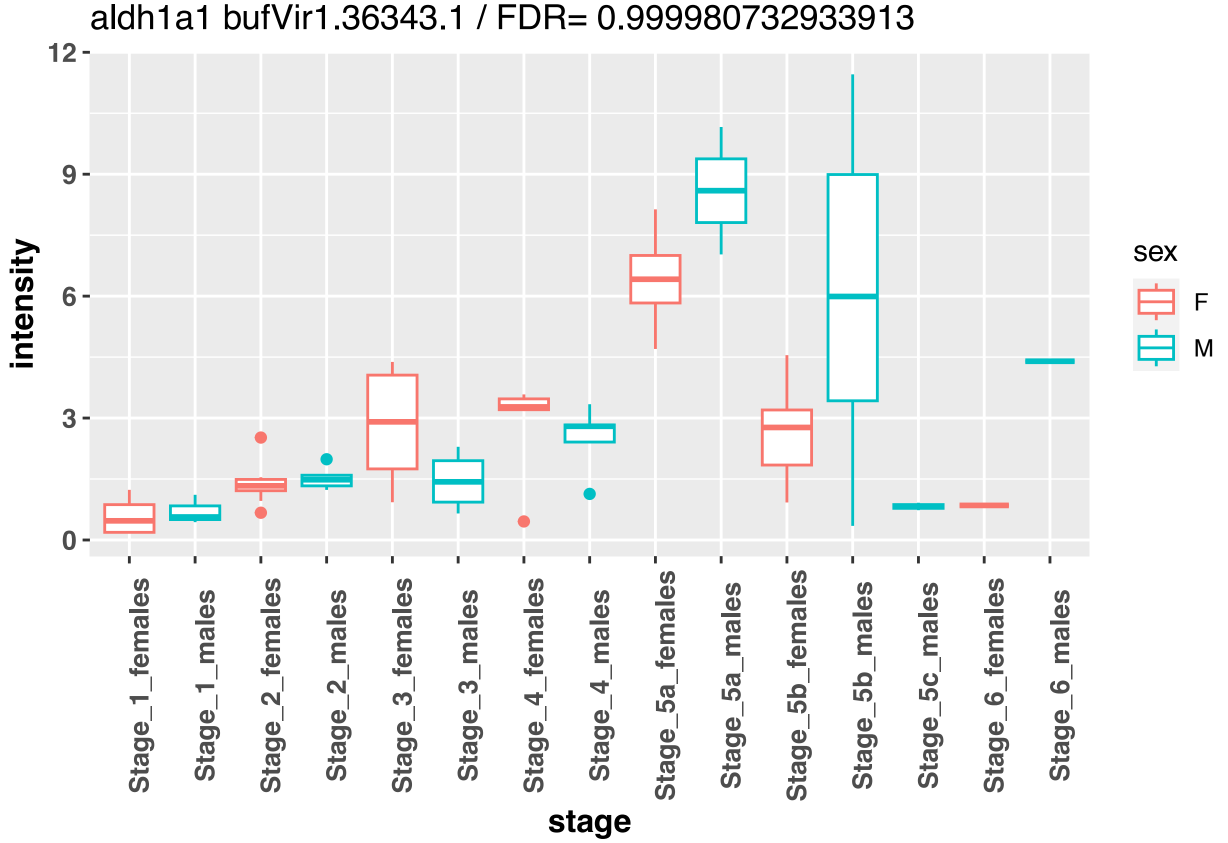
aldh1a1 bufVir1.36343.1 / FDR= 0.999980732933913
intensity
12
9
6
3
0
Stage_1_females
Stage_1_males
Stage_2_females
Stage_2_males
Stage_3_females
Stage_3_males
Stage_4_females
Stage_4_males
Stage_5a_females
Stage_5a_males
Stage_5b_females
Stage_5b_males
Stage_5c_males
Stage_6_females
Stage_6_males
stage
sex
F
M

aldh1a2 bufVir1.36343.1 / FDR= 0.999980732933913

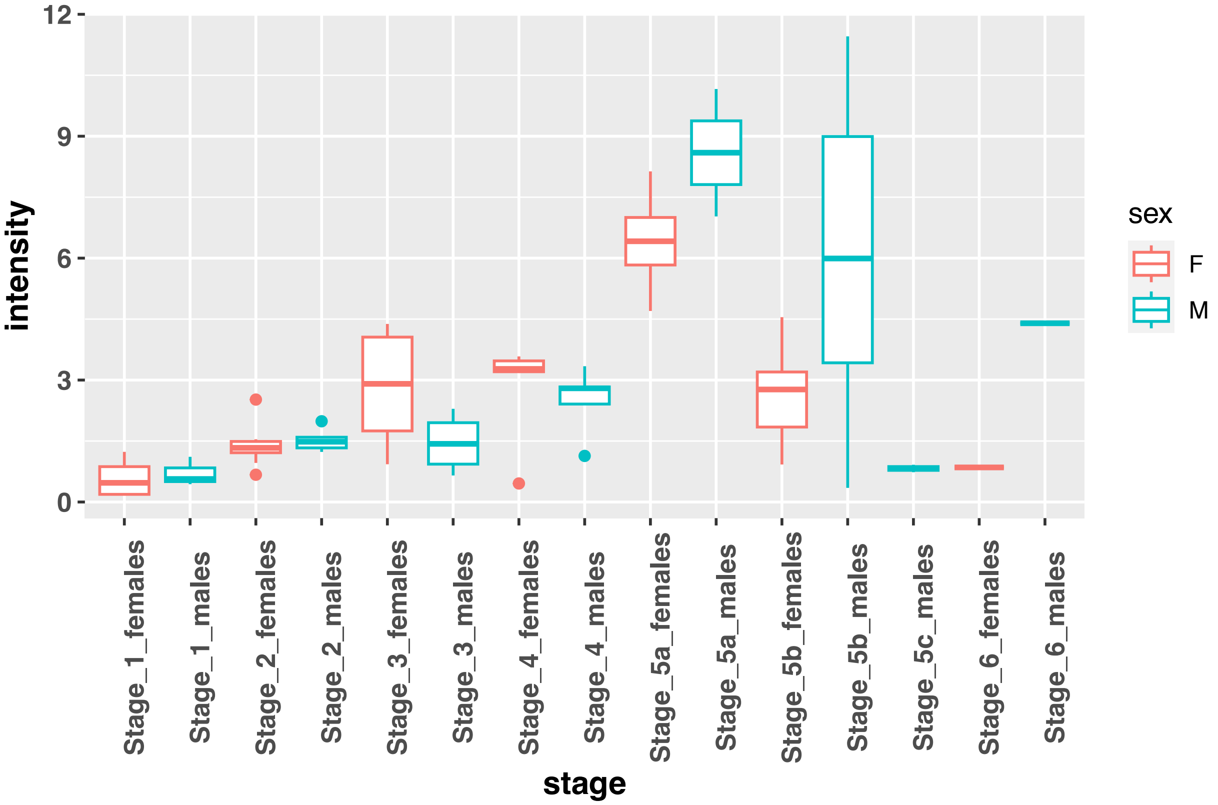


ck18 bufVir1.18618.1 / FDR= 0.92738135669211

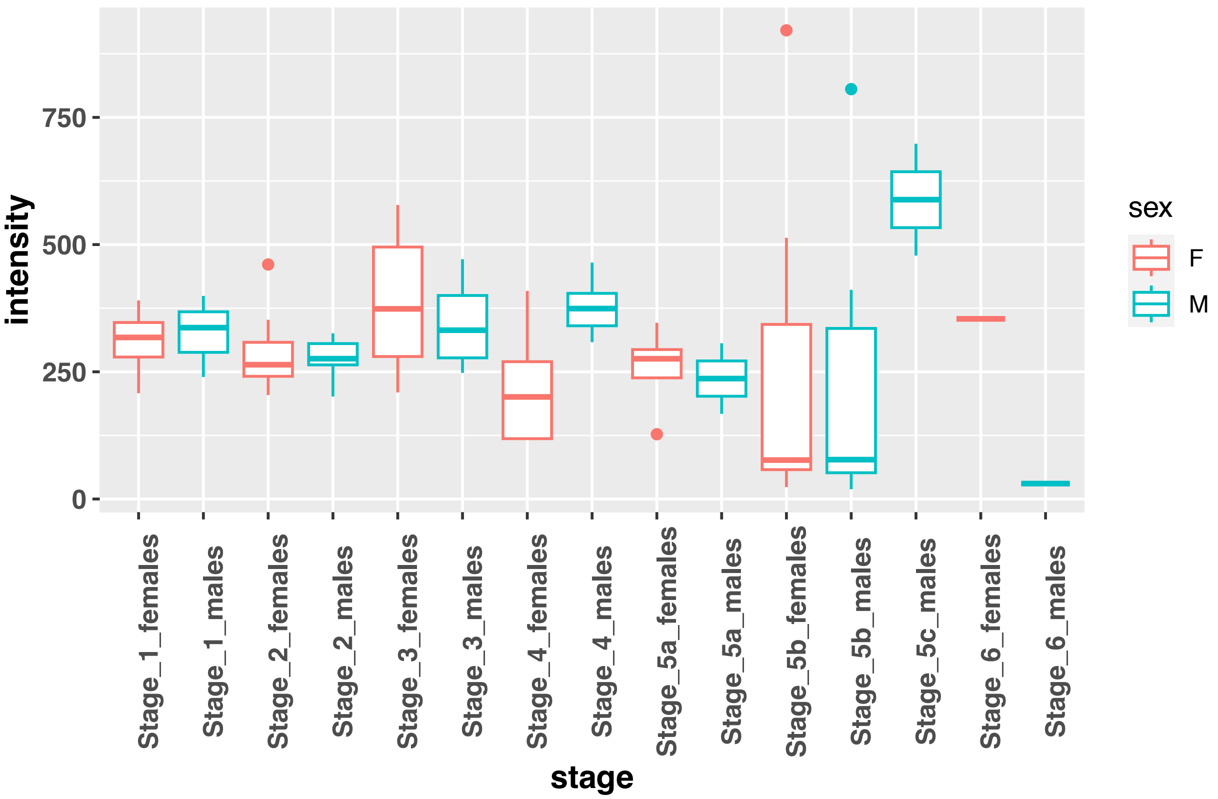

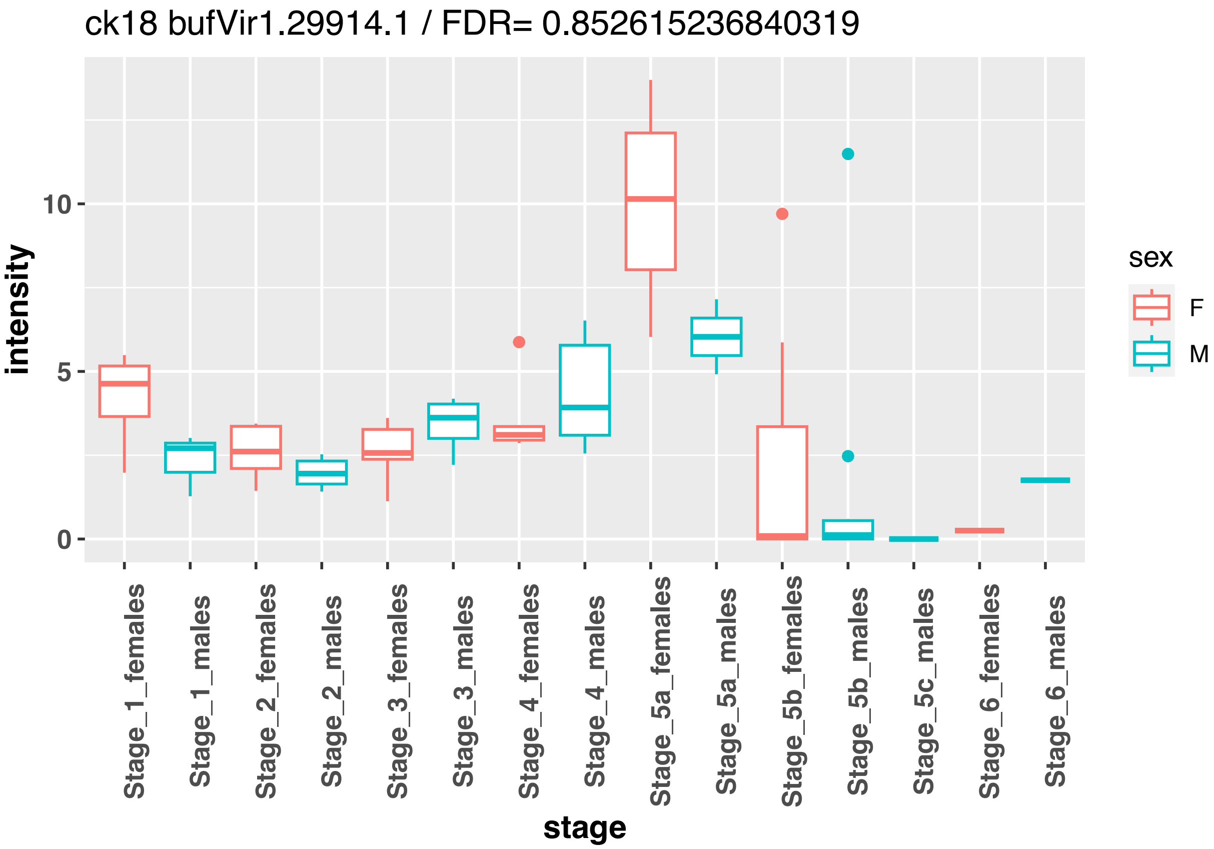
ck18 bufVir1.29914.1 / FDR= 0.852615236840319
intensity
stage
sex
F
M
Stage_1_females
Stage_1_males
Stage_2_females
Stage_2_males
Stage_3_females
Stage_3_males
Stage_4_females
Stage_4_males
Stage_5a_females
Stage_5a_males
Stage_5b_females
Stage_5b_males
Stage_5c_males
Stage_6_females
Stage_6_males

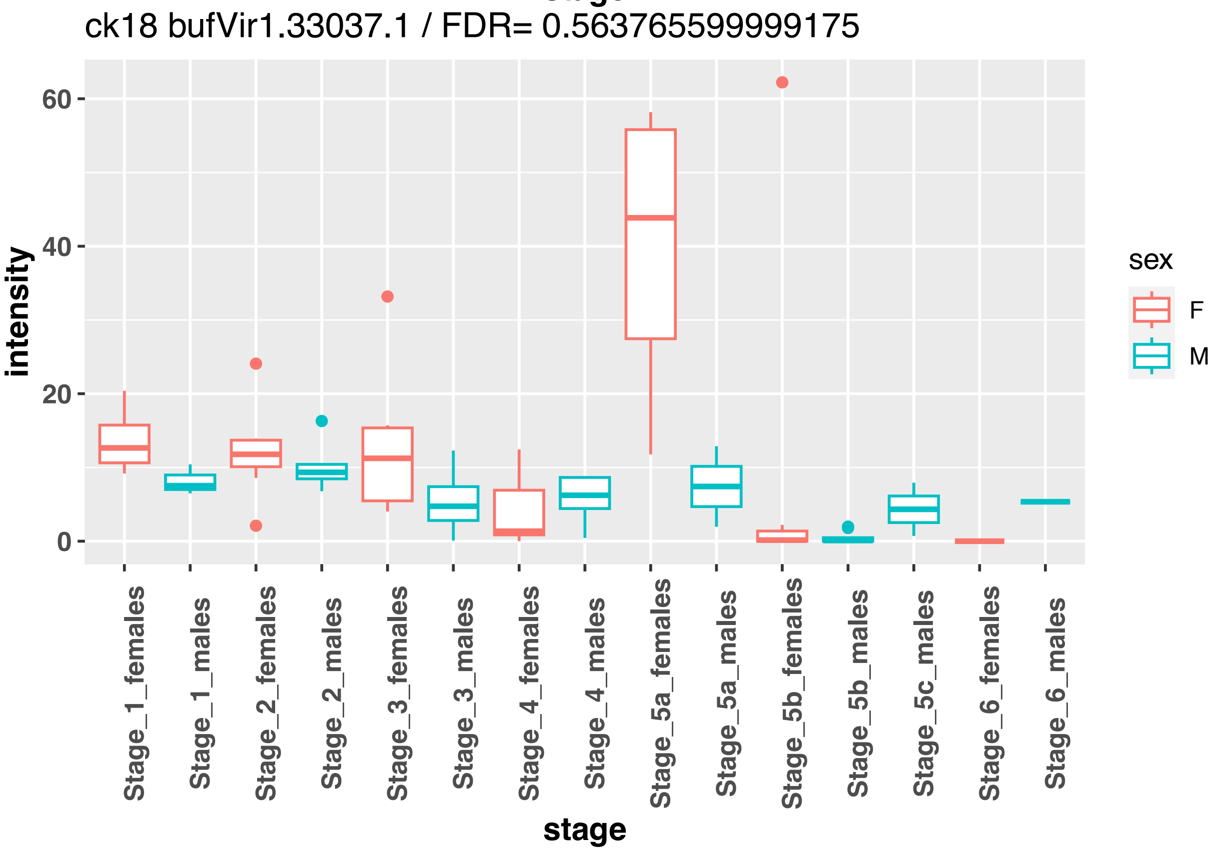
ck18 bufVir1.33037.1 / FDR= 0.563765599999175
intensity
stage
sex
F
M
Stage_1_females
Stage_1_males
Stage_2_females
Stage_2_males
Stage_3_females
Stage_3_males
Stage_4_females
Stage_4_males
Stage_5a_females
Stage_5a_males
Stage_5b_females
Stage_5b_males
Stage_5c_males
Stage_6_females
Stage_6_males

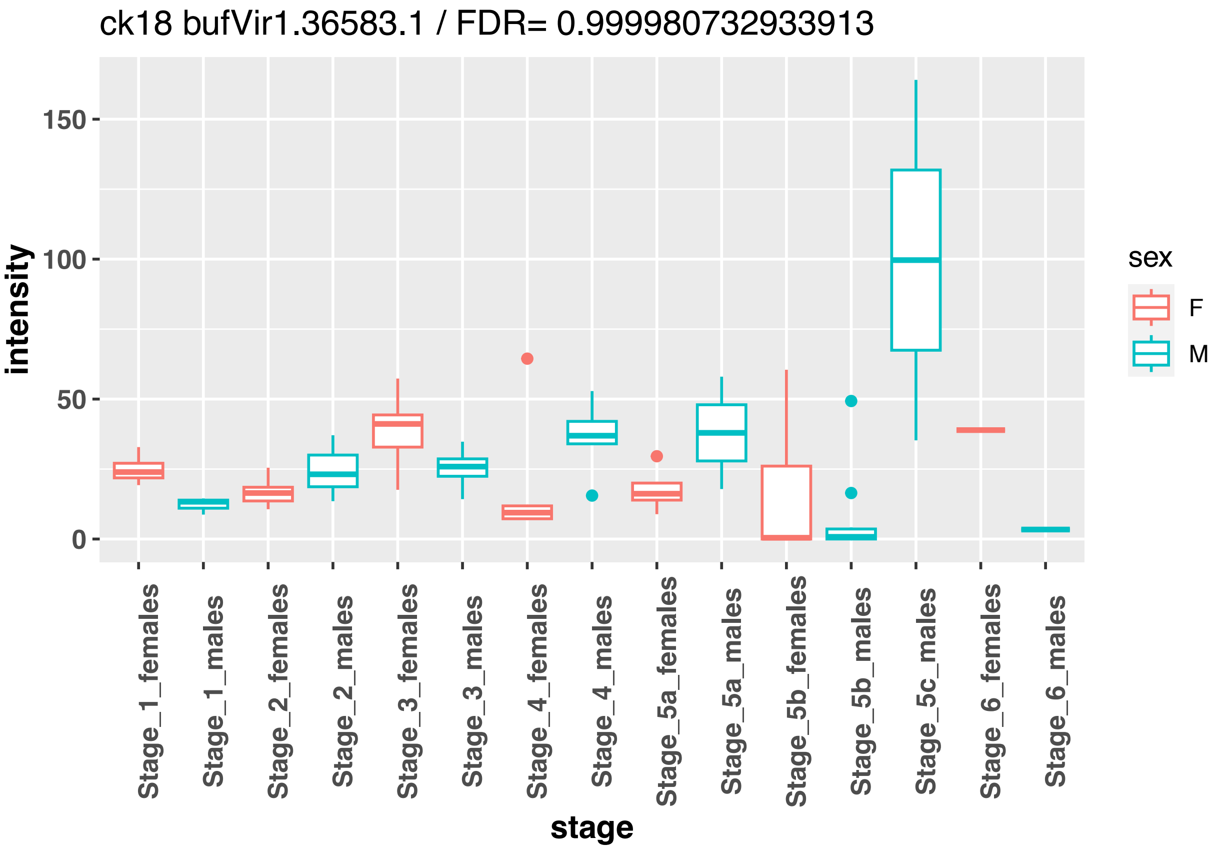
ck18 bufVir1.36583.1 / FDR= 0.999980732933913
intensity
150
100
50
0
stage
Stage_1_females
Stage_1_males
Stage_2_females
Stage_2_males
Stage_3_females
Stage_3_males
Stage_4_females
Stage_4_males
Stage_5a_females
Stage_5a_males
Stage_5b_females
Stage_5b_males
Stage_5c_males
Stage_6_females
Stage_6_males
sex
F
M

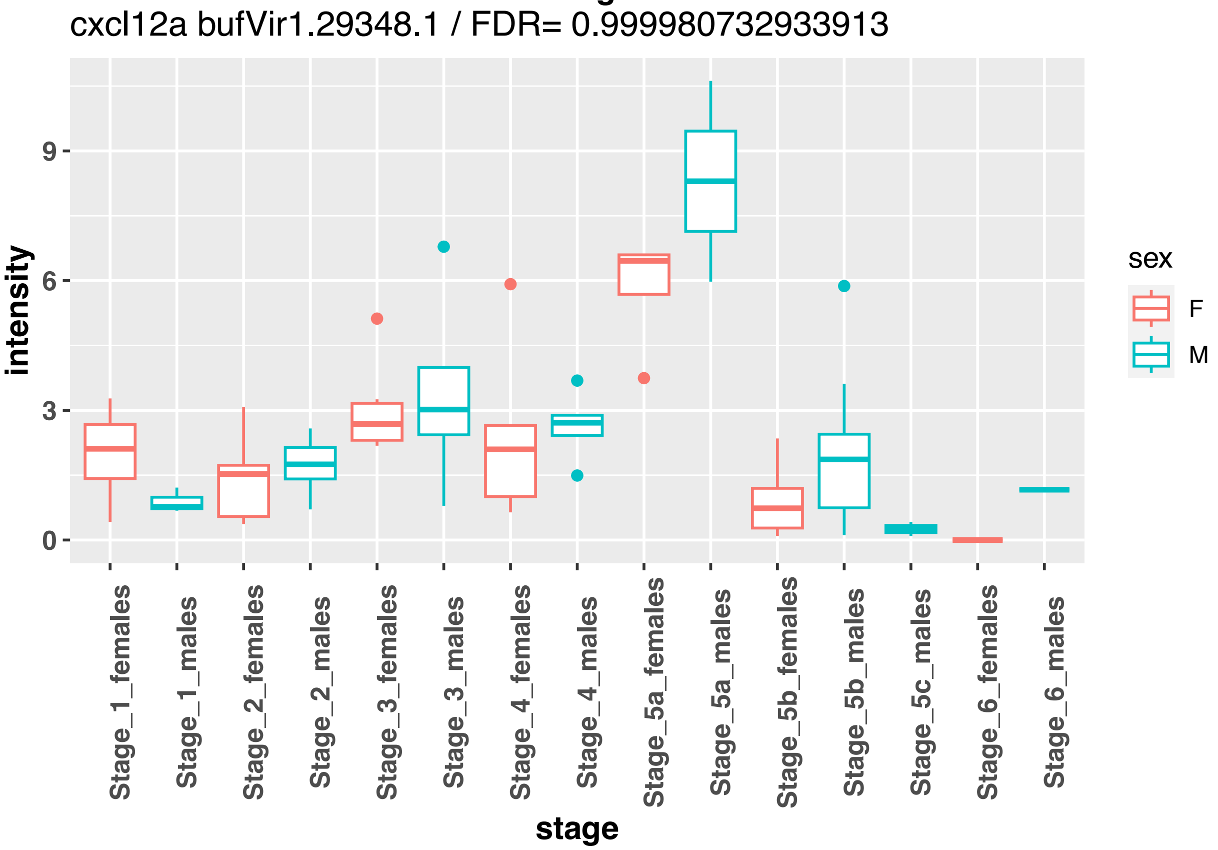
cxcl12a bufVir1.29348.1 / FDR= 0.999980732933913
intensity
9
6
3
0
stage
Stage_1_females
Stage_1_males
Stage_2_females
Stage_2_males
Stage_3_females
Stage_3_males
Stage_4_females
Stage_4_males
Stage_5a_females
Stage_5a_males
Stage_5b_females
Stage_5b_males
Stage_5c_males
Stage_6_females
Stage_6_males
sex
F
M

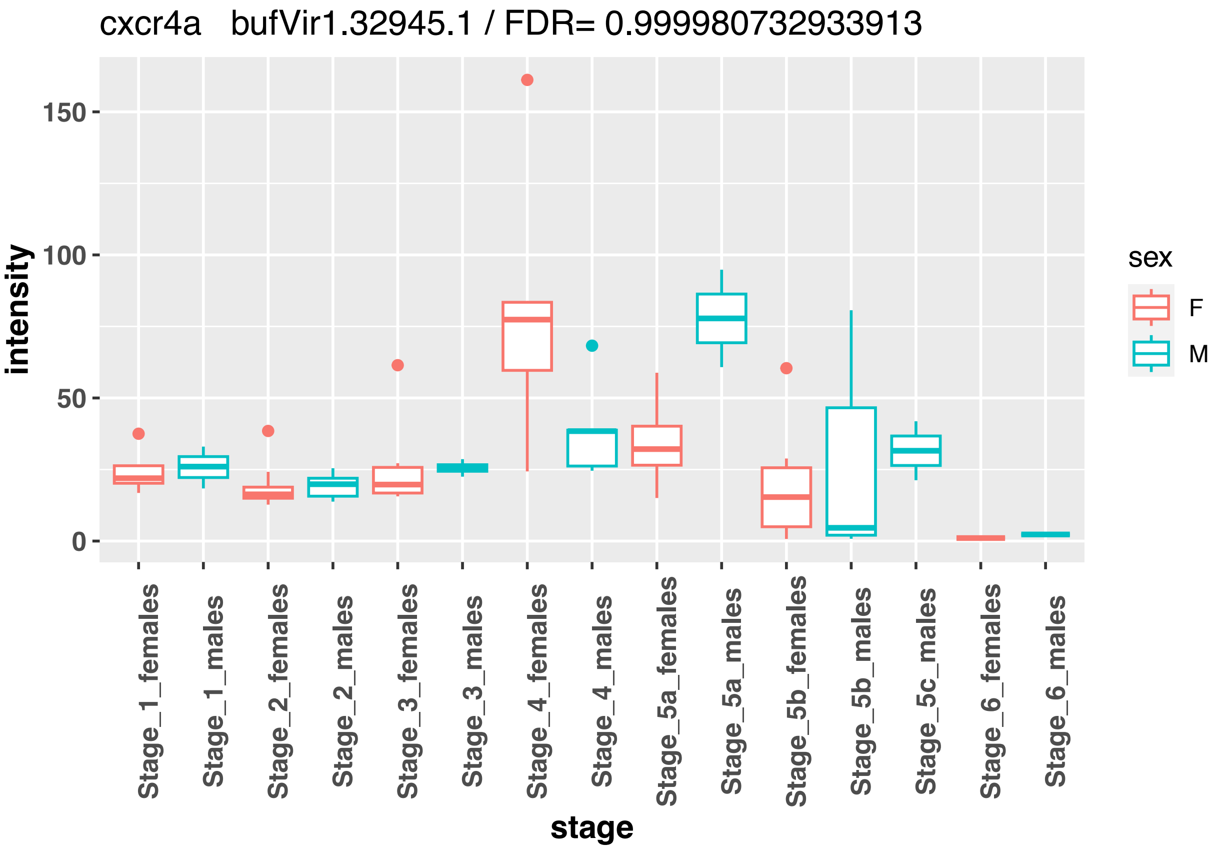
cxcr4a bufVir1.32945.1 / FDR= 0.999980732933913
intensity
150
100
50
0
Stage_1_females
Stage_1_males
Stage_2_females
Stage_2_males
Stage_3_females
Stage_3_males
Stage_4_females
Stage_4_males
Stage_5a_females
Stage_5a_males
Stage_5b_females
Stage_5b_males
Stage_5c_males
Stage_6_females
Stage_6_males
stage
sex
F
M

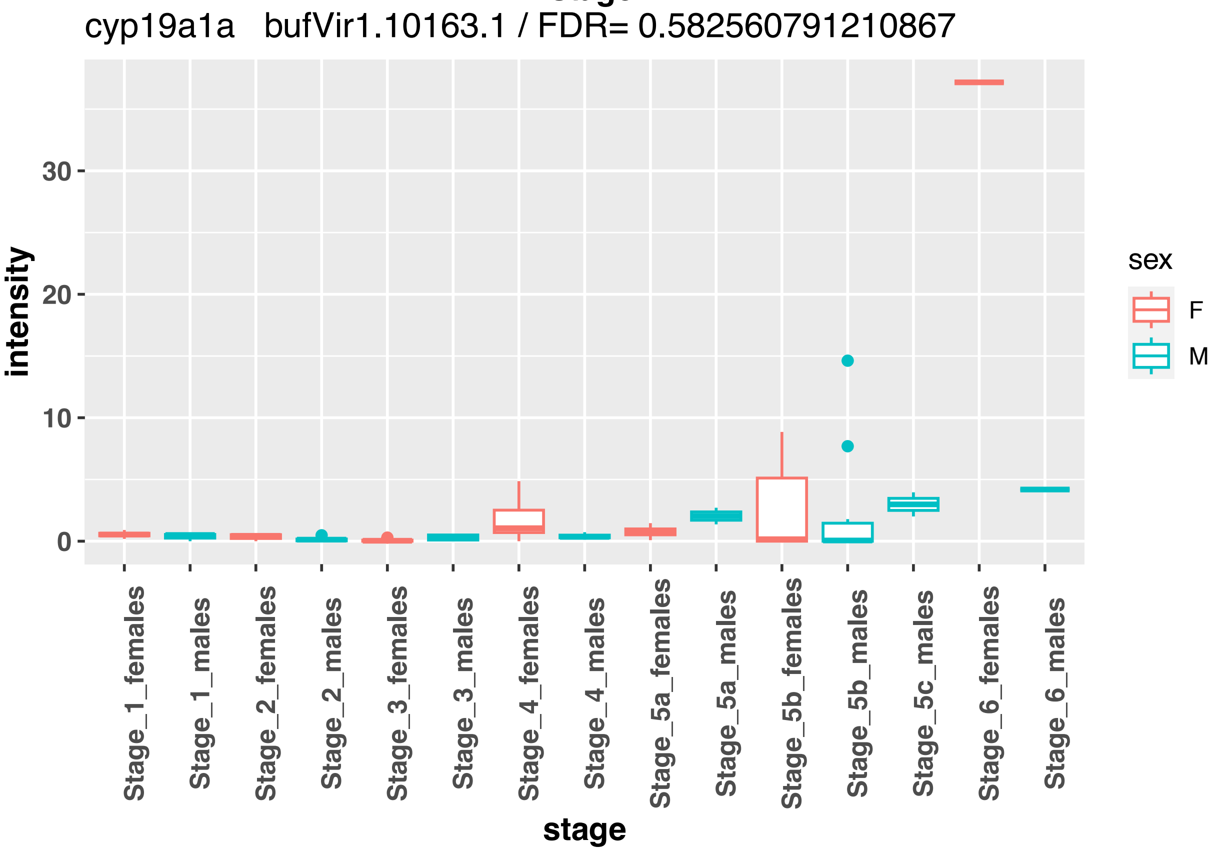
cyp19a1a bufVir1.10163.1 / FDR= 0.582560791210867
intensity
30
20
10
0
Stage_1_females
Stage_1_males
Stage_2_females
Stage_2_males
Stage_3_females
Stage_3_males
Stage_4_females
Stage_4_males
Stage_5a_females
Stage_5a_males
Stage_5b_females
Stage_5b_males
Stage_5c_males
Stage_6_females
Stage_6_males
stage
sex
F
M

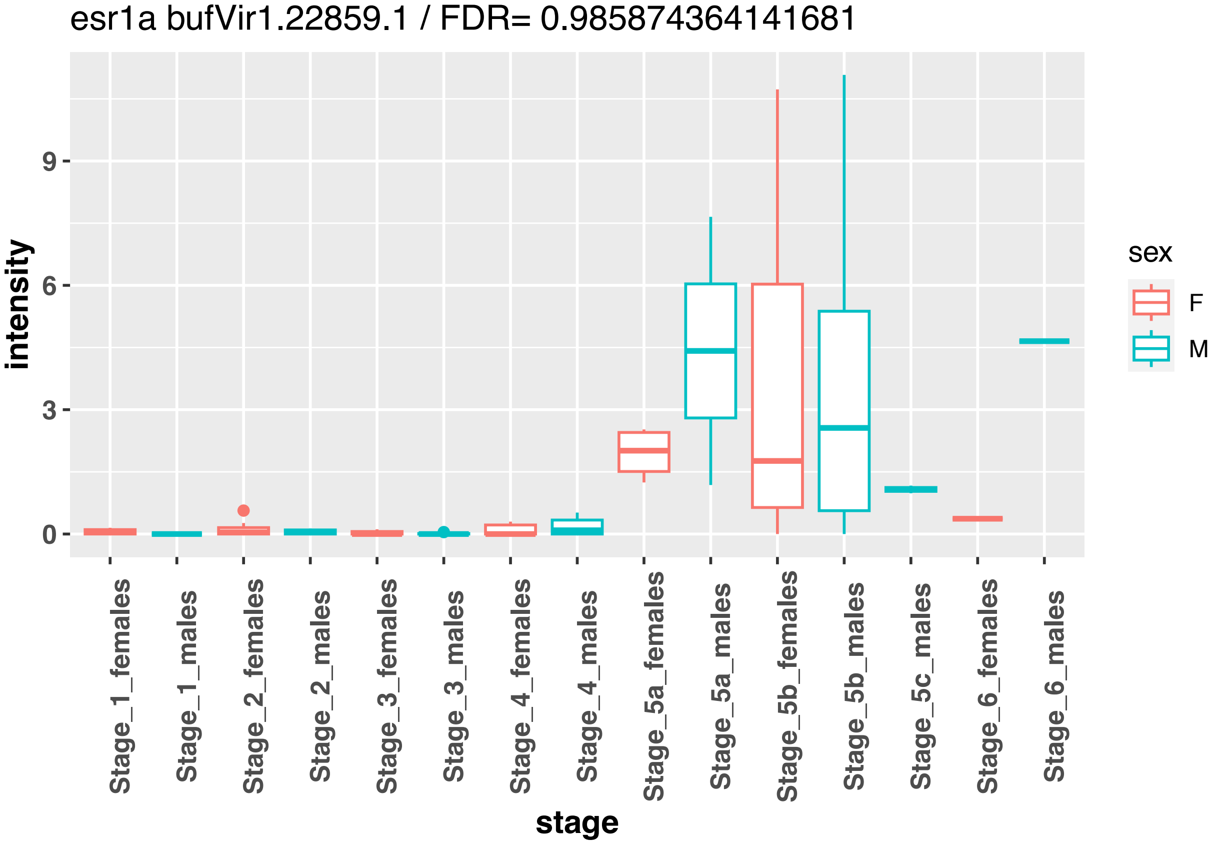
esr1a bufVir1.22859.1 / FDR= 0.985874364141681
intensity
stage
sex
F
M
0
3
6
9
Stage_1_females
Stage_1_males
Stage_2_females
Stage_2_males
Stage_3_females
Stage_3_males
Stage_4_females
Stage_4_males
Stage_5a_females
Stage_5a_males
Stage_5b_females
Stage_5b_males
Stage_5c_males
Stage_6_females
Stage_6_males

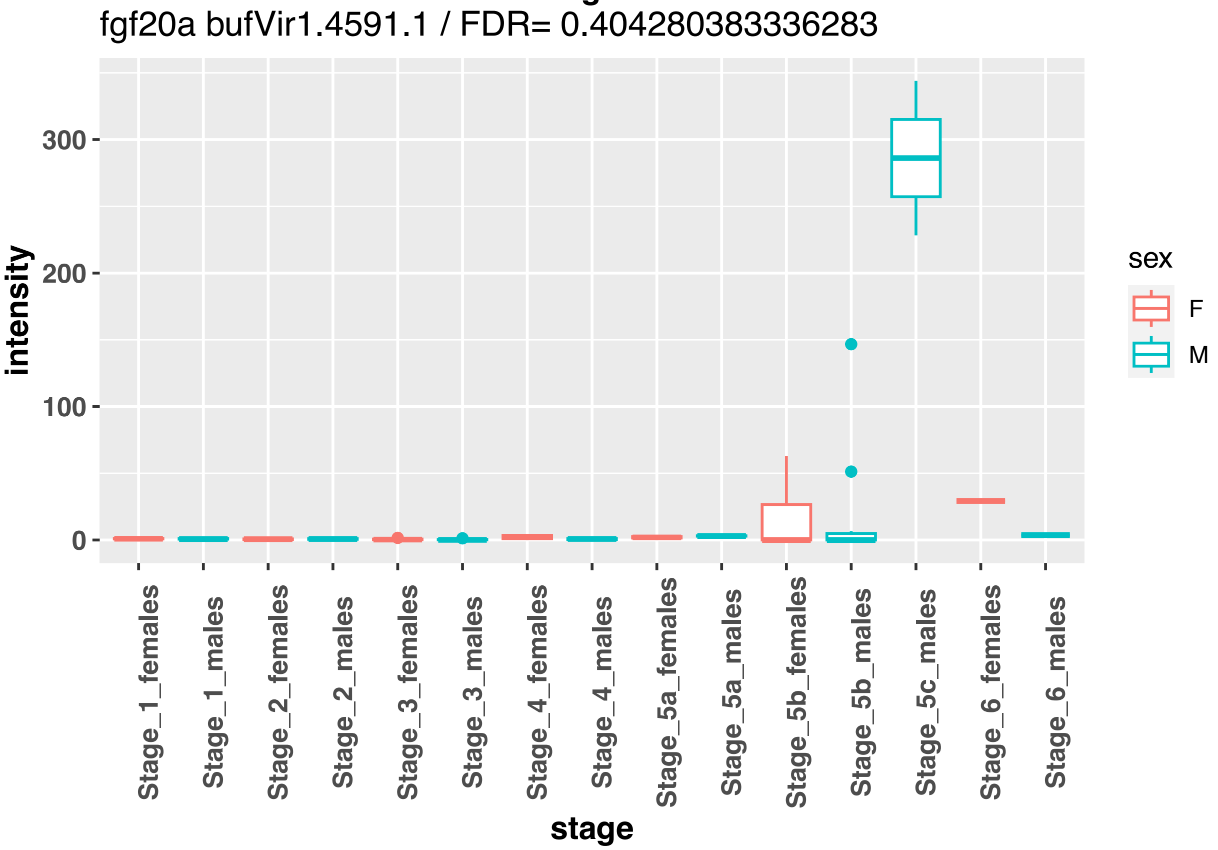
fgf20a bufVir1.4591.1 / FDR= 0.404280383336283
intensity
stage
sex
F
M
0
100
200
300
Stage_1_females
Stage_1_males
Stage_2_females
Stage_2_males
Stage_3_females
Stage_3_males
Stage_4_females
Stage_4_males
Stage_5a_females
Stage_5a_males
Stage_5b_females
Stage_5b_males
Stage_5c_males
Stage_6_females
Stage_6_males

nanos2 bufVir1.9233.1 / FDR= 0.496755733517493

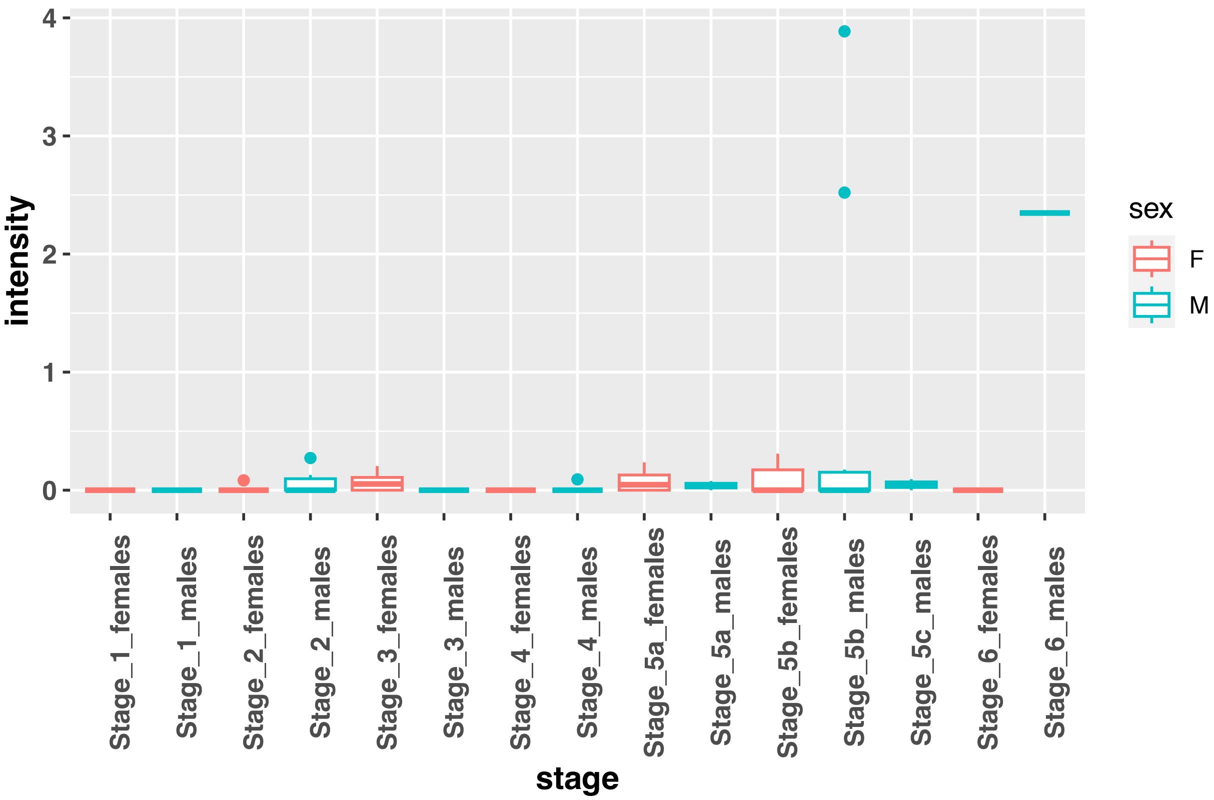


pdgfab bufVir1.31826.1 / FDR= 0.999980732933913

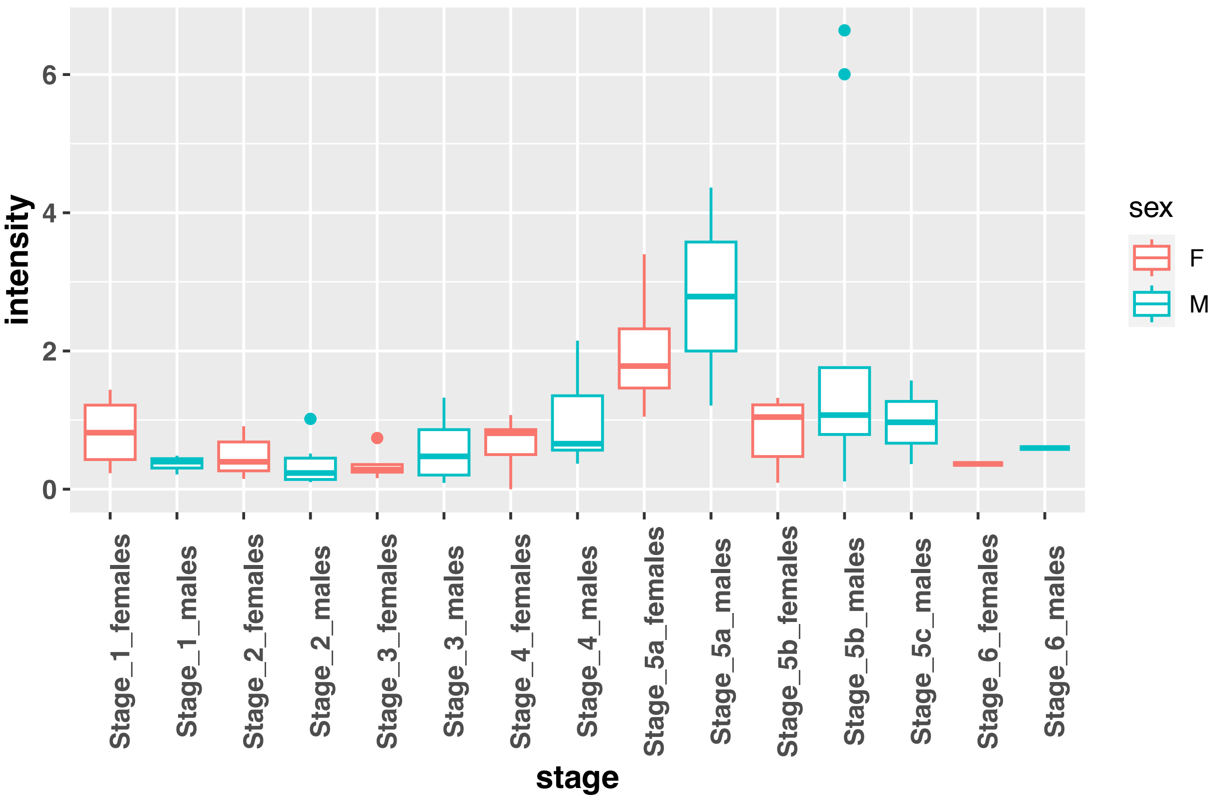

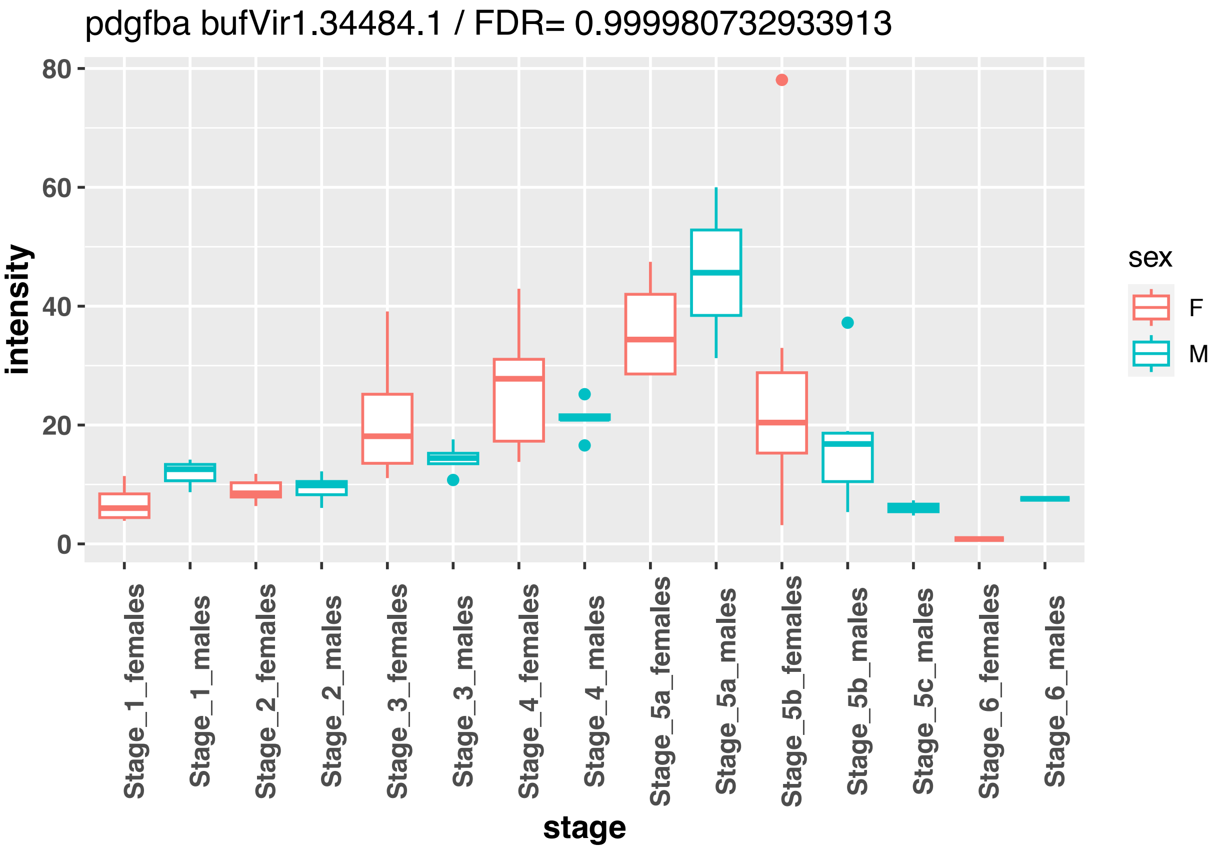
pdgfba bufVir1.34484.1 / FDR= 0.999980732933913
intensity
80
60
40
20
0
Stage_1_females
Stage_1_males
Stage_2_females
Stage_2_males
Stage_3_females
Stage_3_males
Stage_4_females
Stage_4_males
Stage_5a_females
Stage_5a_males
Stage_5b_females
Stage_5b_males
Stage_5c_males
Stage_6_females
Stage_6_males
stage
sex
F
M

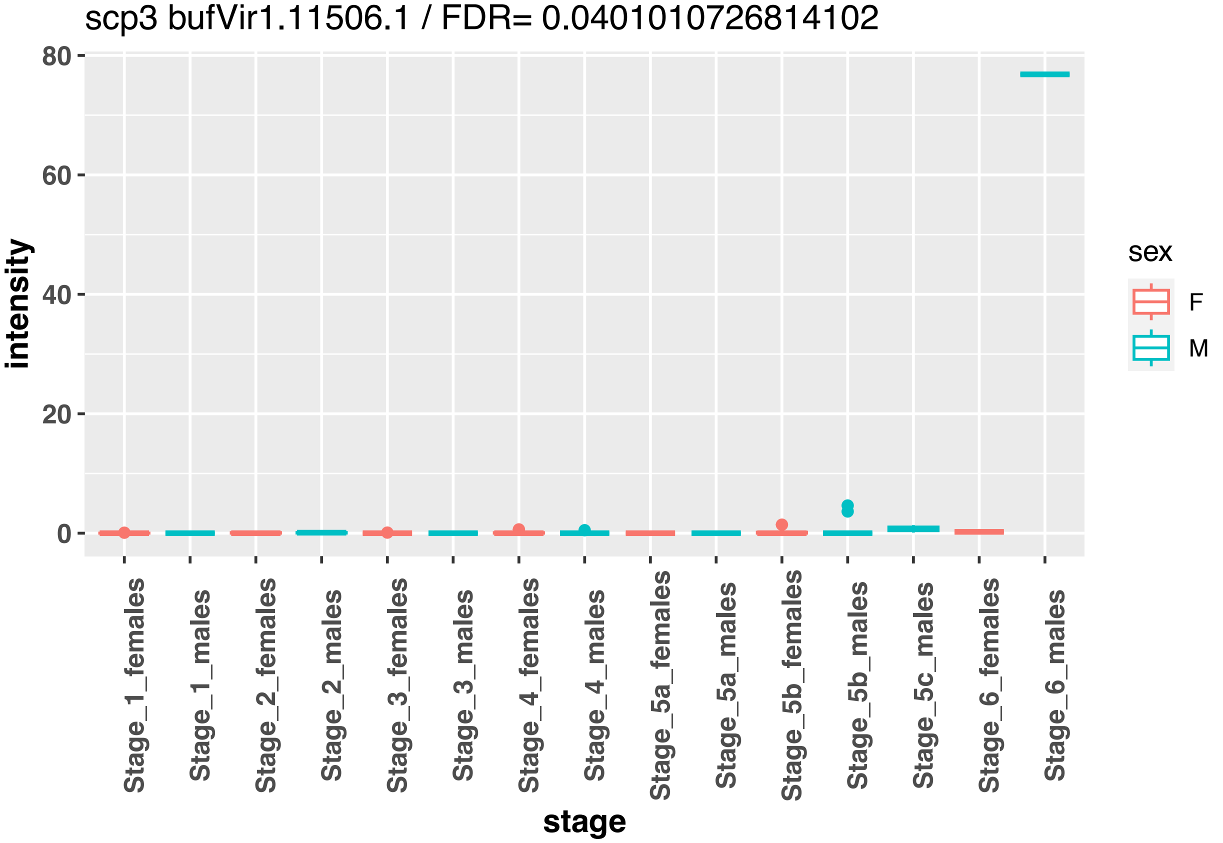
scp3 bufVir1.11506.1 / FDR= 0.0401010726814102
intensity
80
60
40
20
0
Stage_1_females
Stage_1_males
Stage_2_females
Stage_2_males
Stage_3_females
Stage_3_males
Stage_4_females
Stage_4_males
Stage_5a_females
Stage_5a_males
Stage_5b_females
Stage_5b_males
Stage_5c_males
Stage_6_females
Stage_6_males
stage
sex
F
M

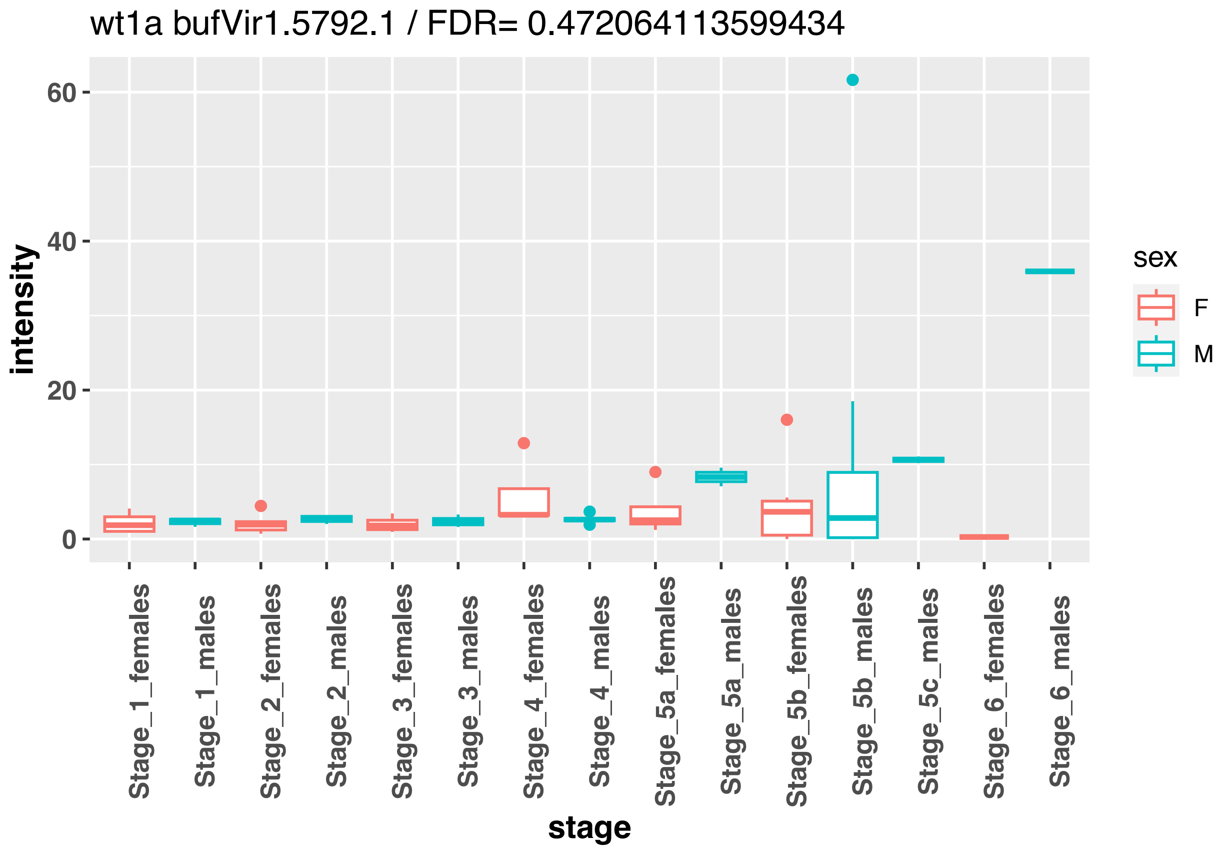
wt1a bufVir1.5792.1 / FDR= 0.472064113599434
intensity
60
40
20
0
Stage_1_females
Stage_1_males
Stage_2_females
Stage_2_males
Stage_3_females
Stage_3_males
Stage_4_females
Stage_4_males
Stage_5a_females
Stage_5a_males
Stage_5b_females
Stage_5b_males
Stage_5c_males
Stage_6_females
Stage_6_males
stage
sex
F
M

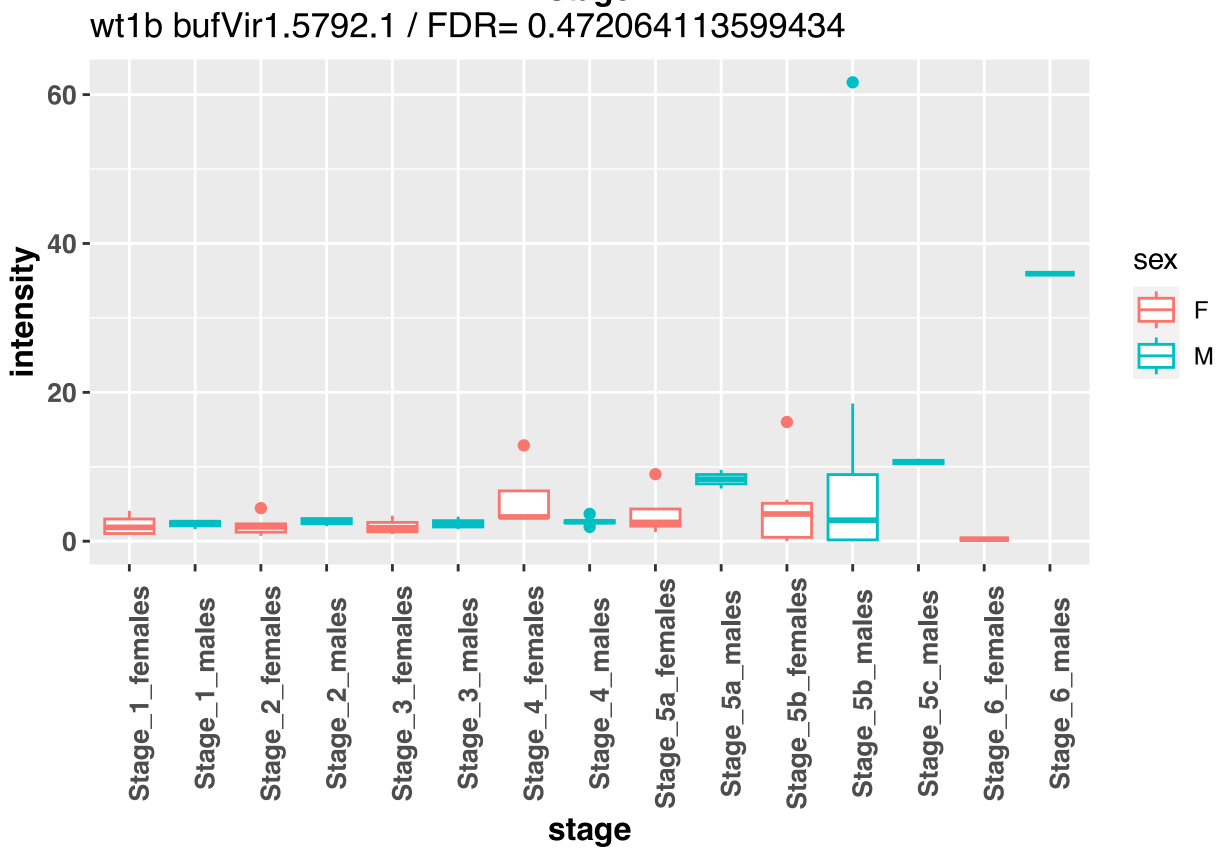
wt1b bufVir1.5792.1 / FDR= 0.472064113599434
intensity
60
40
20
0
Stage_1_females
Stage_1_males
Stage_2_females
Stage_2_males
Stage_3_females
Stage_3_males
Stage_4_females
Stage_4_males
Stage_5a_females
Stage_5a_males
Stage_5b_females
Stage_5b_males
Stage_5c_males
Stage_6_females
Stage_6_males
stage
sex
F
M

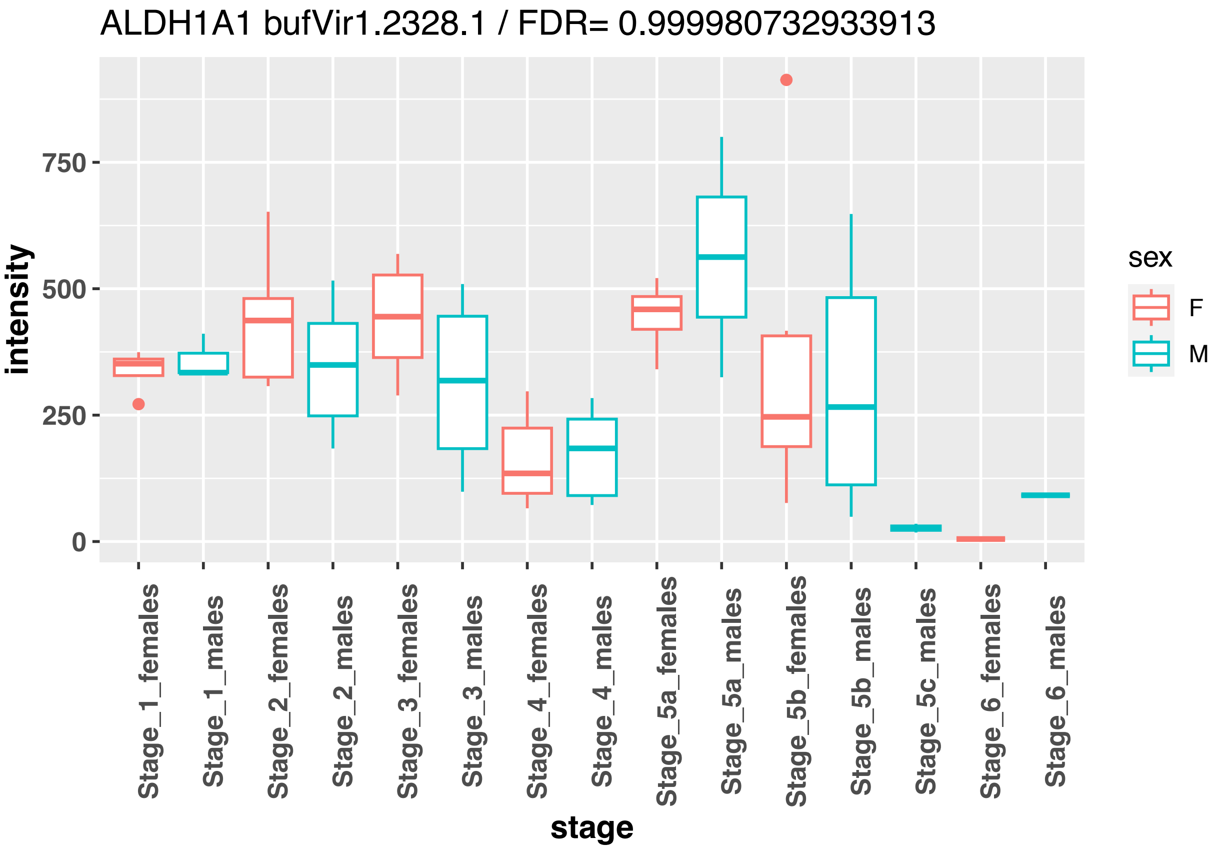
ALDH1A1 bufVir1.2328.1 / FDR= 0.999980732933913
intensity
750
500
250
0
Stage_1_females
Stage_1_males
Stage_2_females
Stage_2_males
Stage_3_females
Stage_3_males
Stage_4_females
Stage_4_males
Stage_5a_females
Stage_5a_males
Stage_5b_females
Stage_5b_males
Stage_5c_males
Stage_6_females
Stage_6_males
stage
sex
F
M

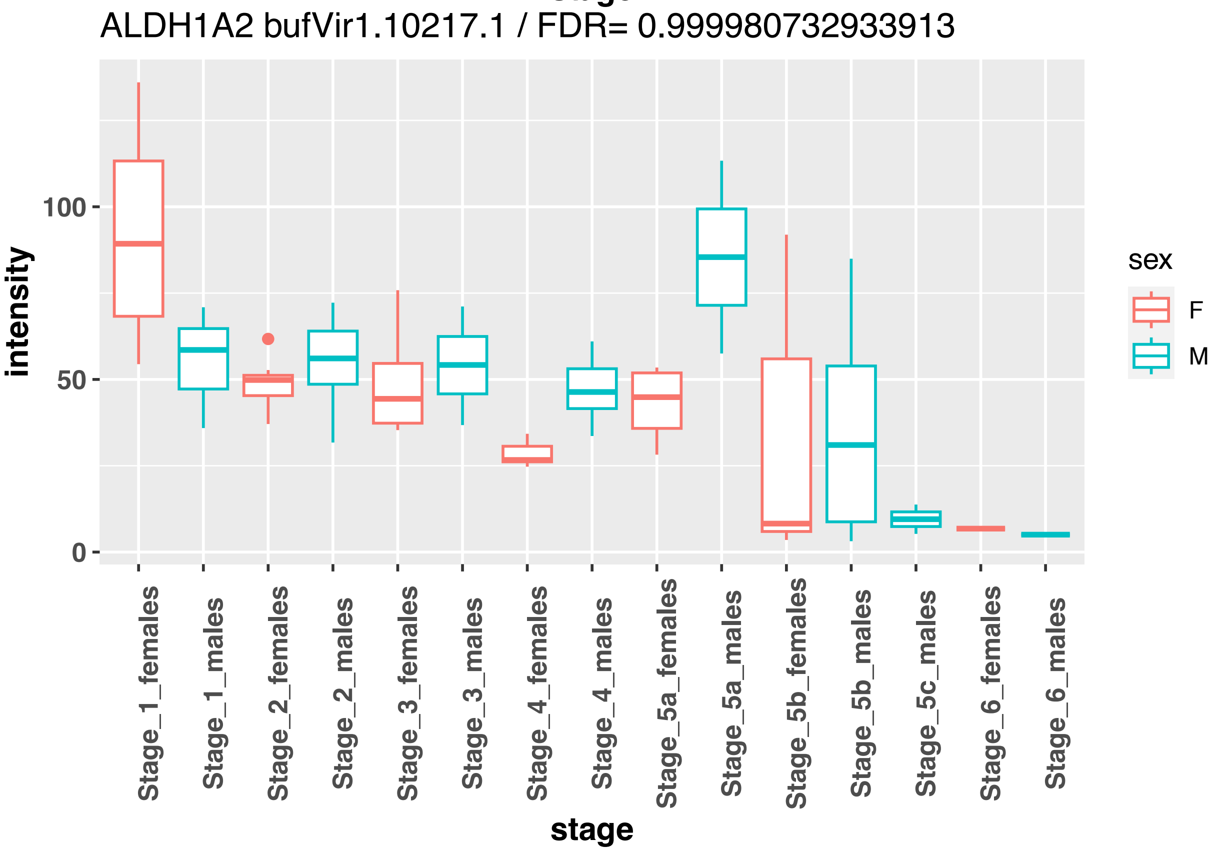
ALDH1A2 bufVir1.10217.1 / FDR= 0.999980732933913
intensity
100
50
0
Stage_1_females
Stage_1_males
Stage_2_females
Stage_2_males
Stage_3_females
Stage_3_males
Stage_4_females
Stage_4_males
Stage_5a_females
Stage_5a_males
Stage_5b_females
Stage_5b_males
Stage_5c_males
Stage_6_females
Stage_6_males
stage
sex
F
M

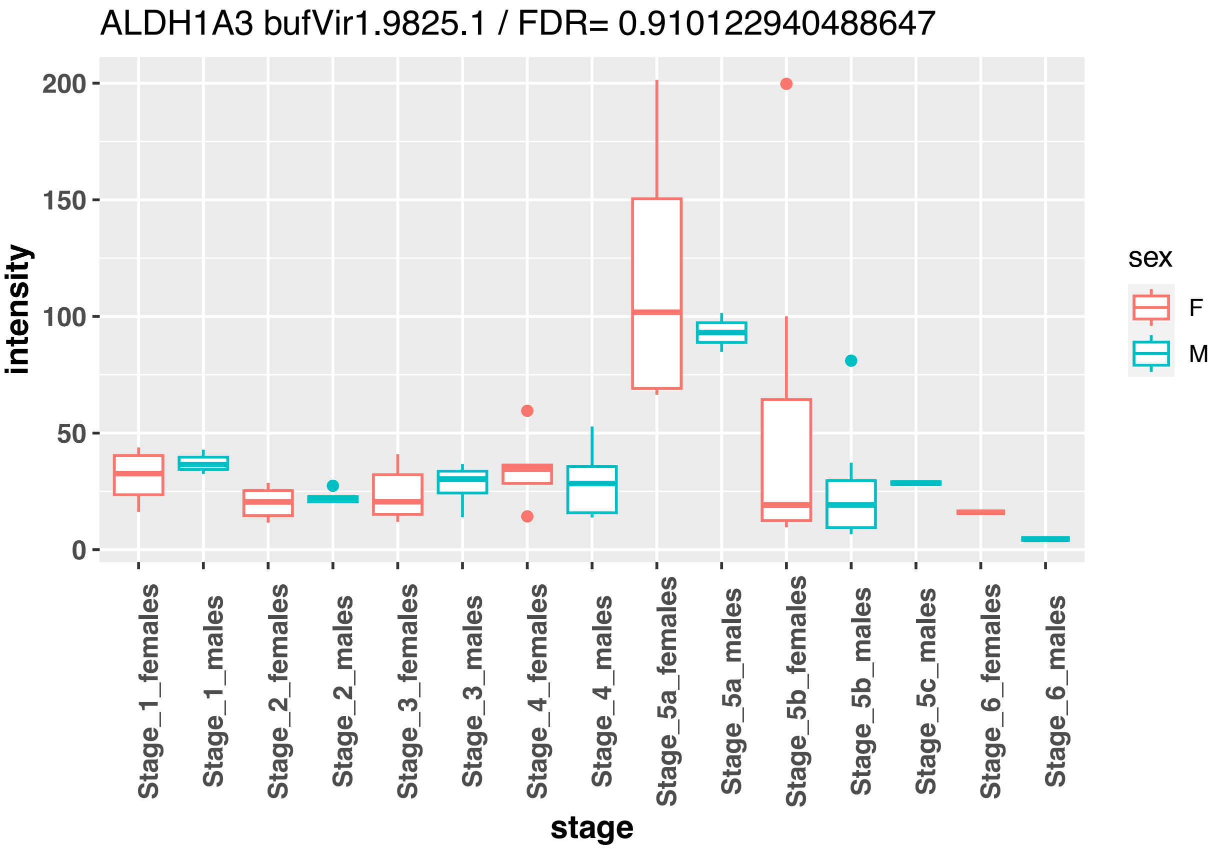
ALDH1A3 bufVir1.9825.1 / FDR= 0.910122940488647
intensity
200
150
100
50
0
stage
Stage_1_females
Stage_1_males
Stage_2_females
Stage_2_males
Stage_3_females
Stage_3_males
Stage_4_females
Stage_4_males
Stage_5a_females
Stage_5a_males
Stage_5b_females
Stage_5b_males
Stage_5c_males
Stage_6_females
Stage_6_males
sex
F
M

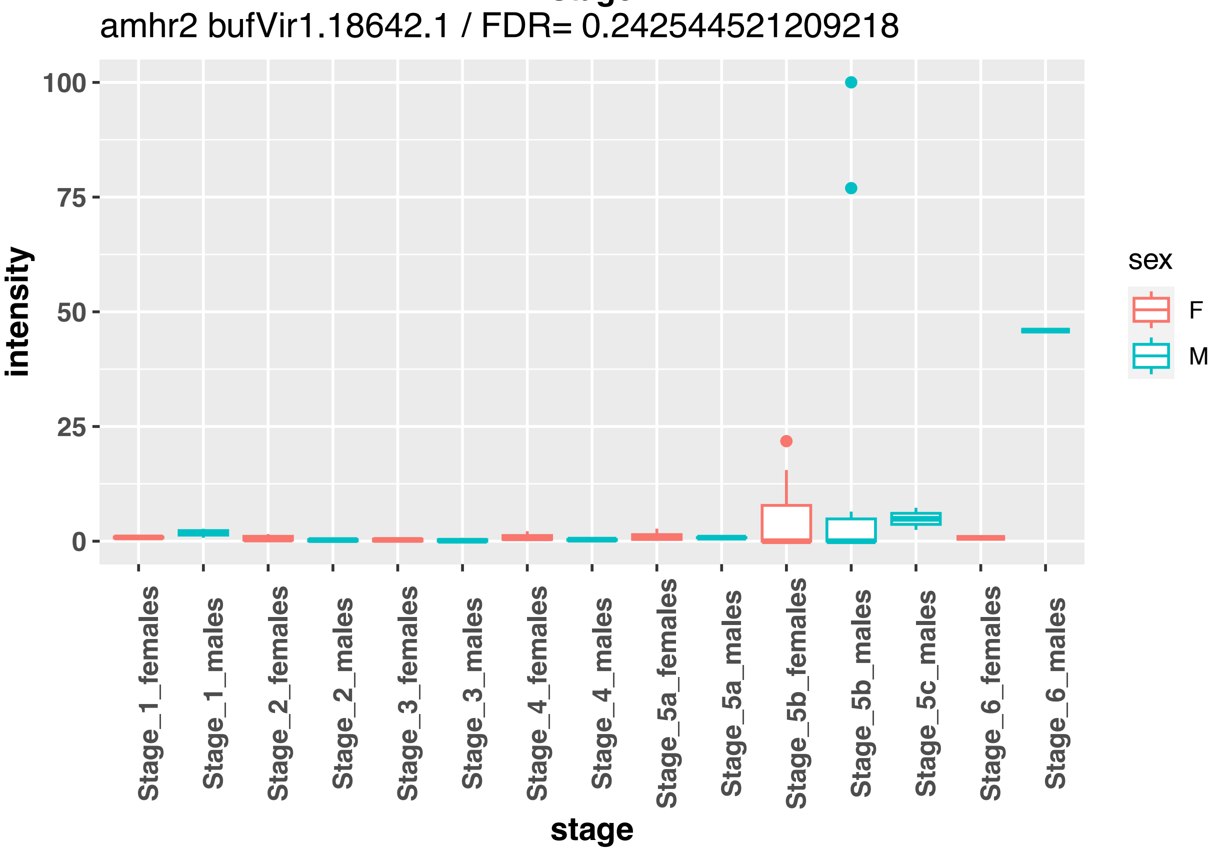
amhr2 bufVir1.18642.1 / FDR= 0.242544521209218
intensity
100
75
50
25
0
stage
Stage_1_females
Stage_1_males
Stage_2_females
Stage_2_males
Stage_3_females
Stage_3_males
Stage_4_females
Stage_4_males
Stage_5a_females
Stage_5a_males
Stage_5b_females
Stage_5b_males
Stage_5c_males
Stage_6_females
Stage_6_males
sex
F
M

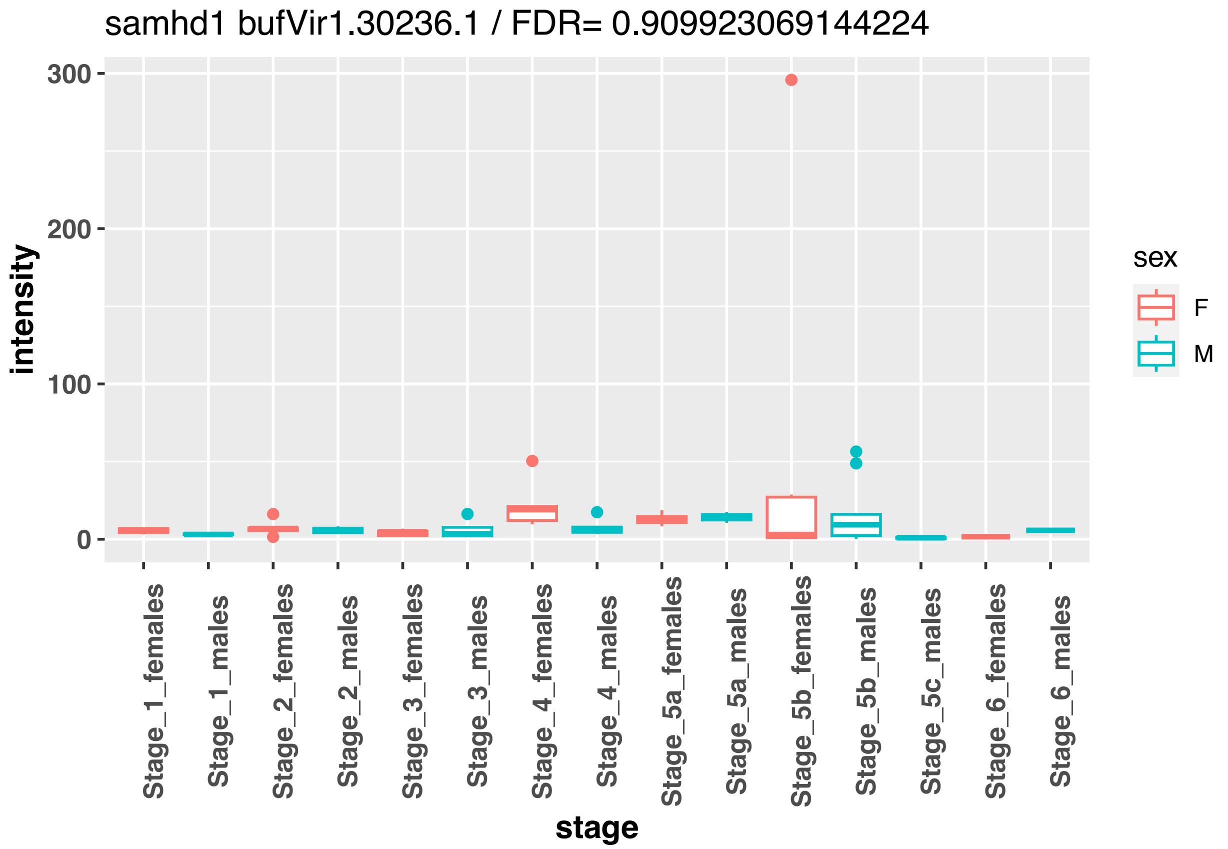
samhd1 bufVir1.30236.1 / FDR= 0.909923069144224
intensity
300
200
100
0
stage
Stage_1_females
Stage_1_males
Stage_2_females
Stage_2_males
Stage_3_females
Stage_3_males
Stage_4_females
Stage_4_males
Stage_5a_females
Stage_5a_males
Stage_5b_females
Stage_5b_males
Stage_5c_males
Stage_6_females
Stage_6_males
sex
F
M

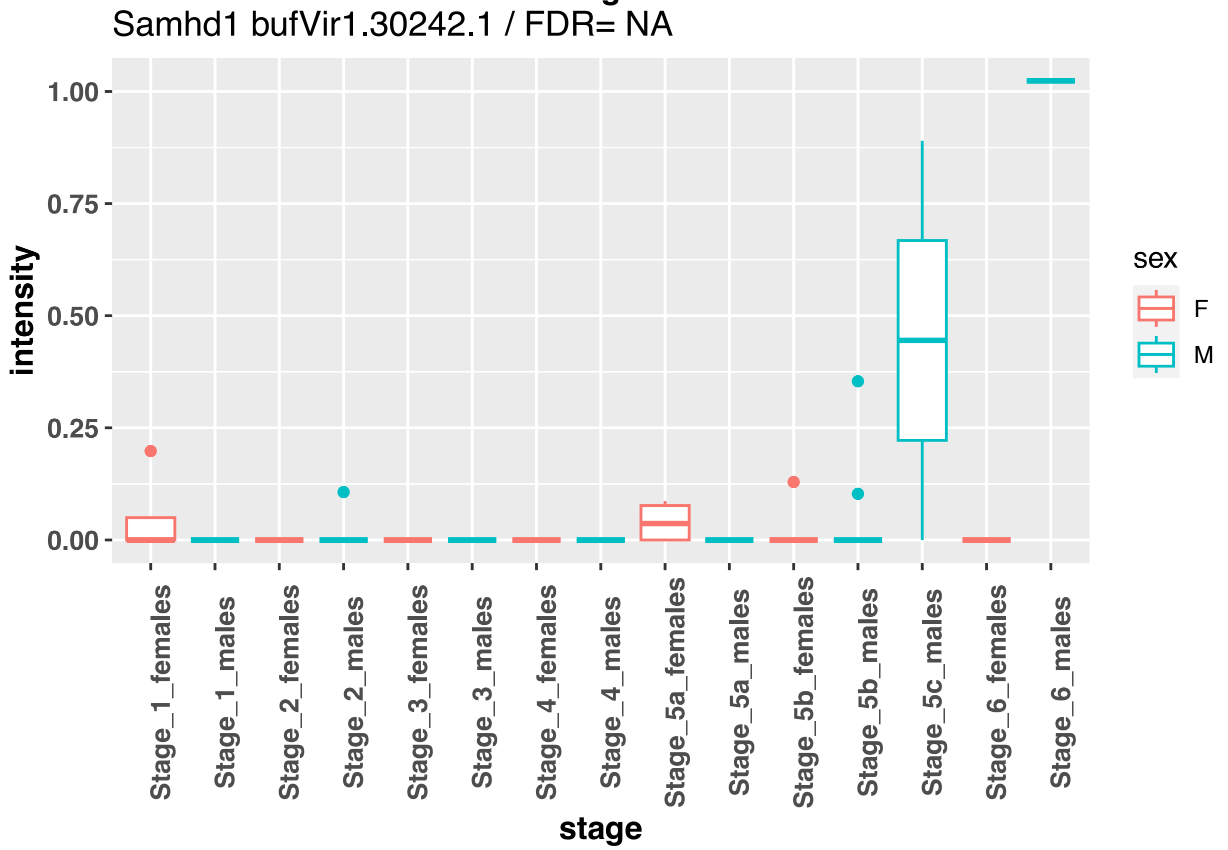
Samhd1 bufVir1.30242.1 / FDR= NA
intensity
1.00
0.75
0.50
0.25
0.00
stage
Stage_1_females
Stage_1_males
Stage_2_females
Stage_2_males
Stage_3_females
Stage_3_males
Stage_4_females
Stage_4_males
Stage_5a_females
Stage_5a_males
Stage_5b_females
Stage_5b_males
Stage_5c_males
Stage_6_females
Stage_6_males
sex
F
M

samhd1 bufVir1.30245.1 / FDR= NA

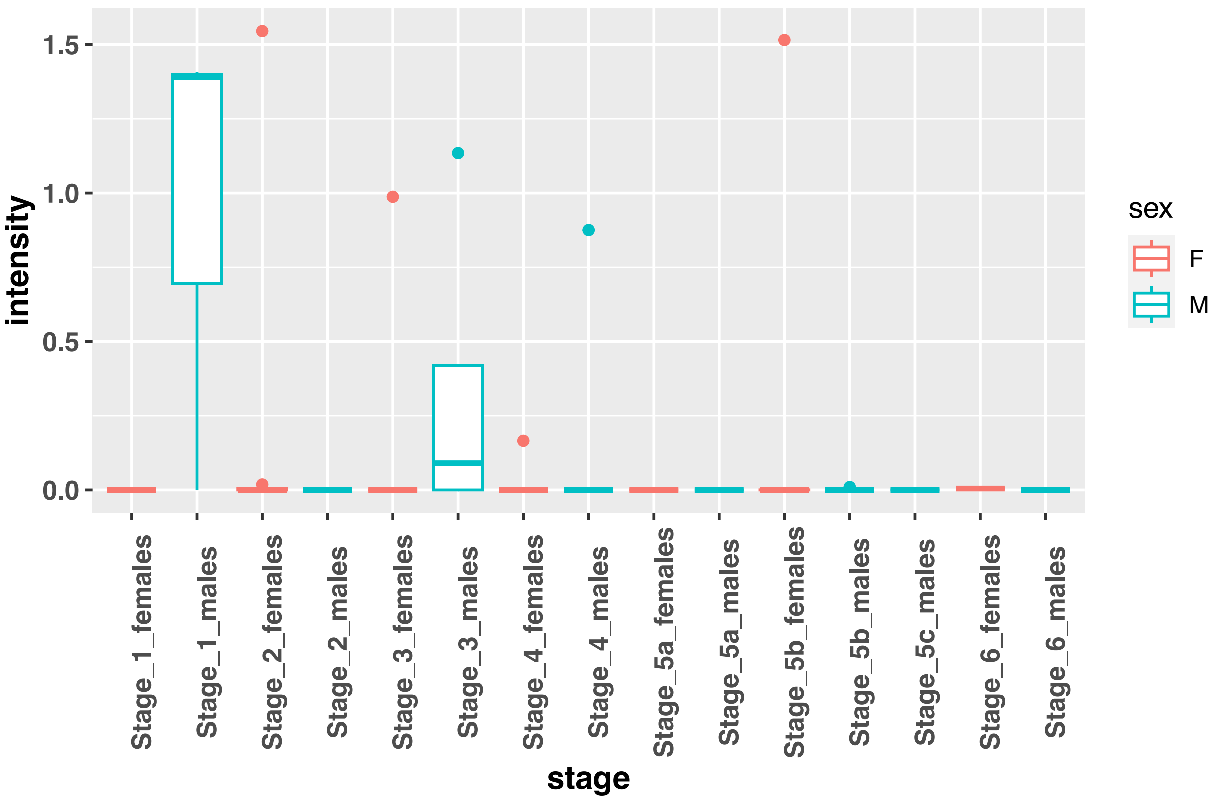


samhd1 bufVir1.30245.4 / FDR= 0.999980732933913

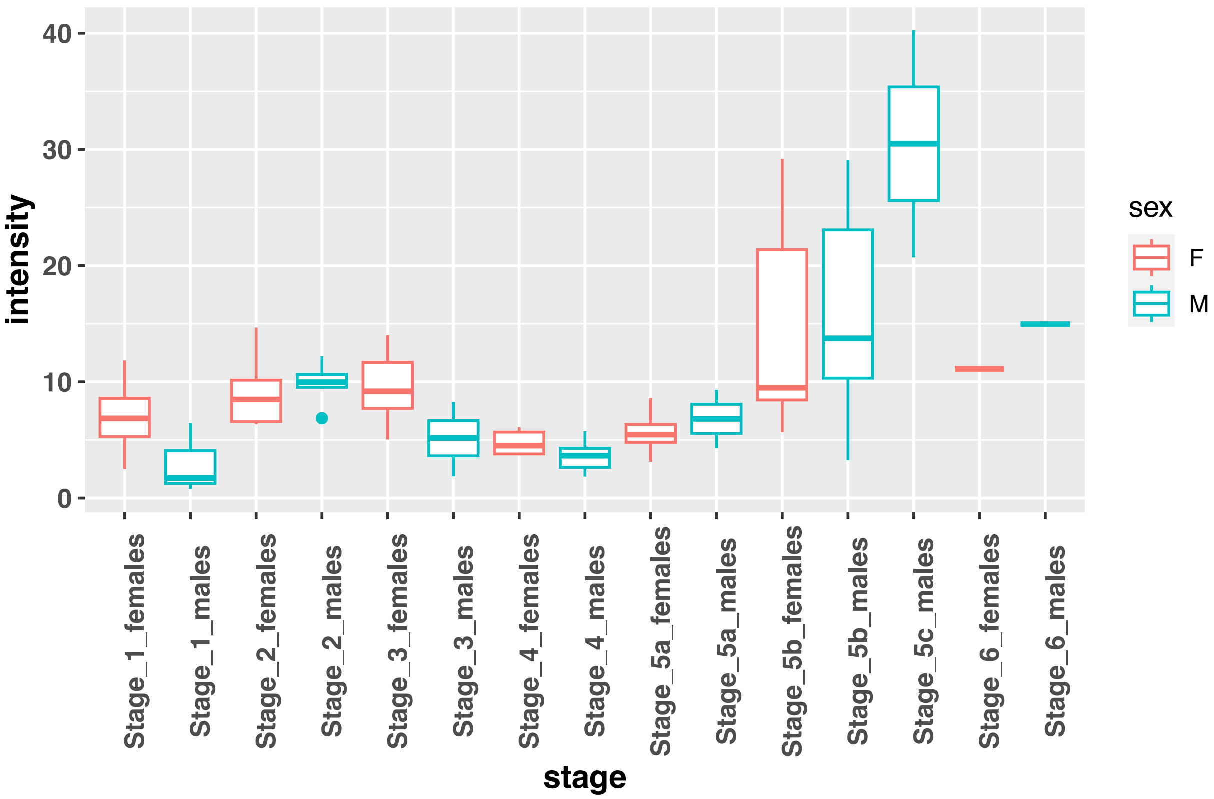

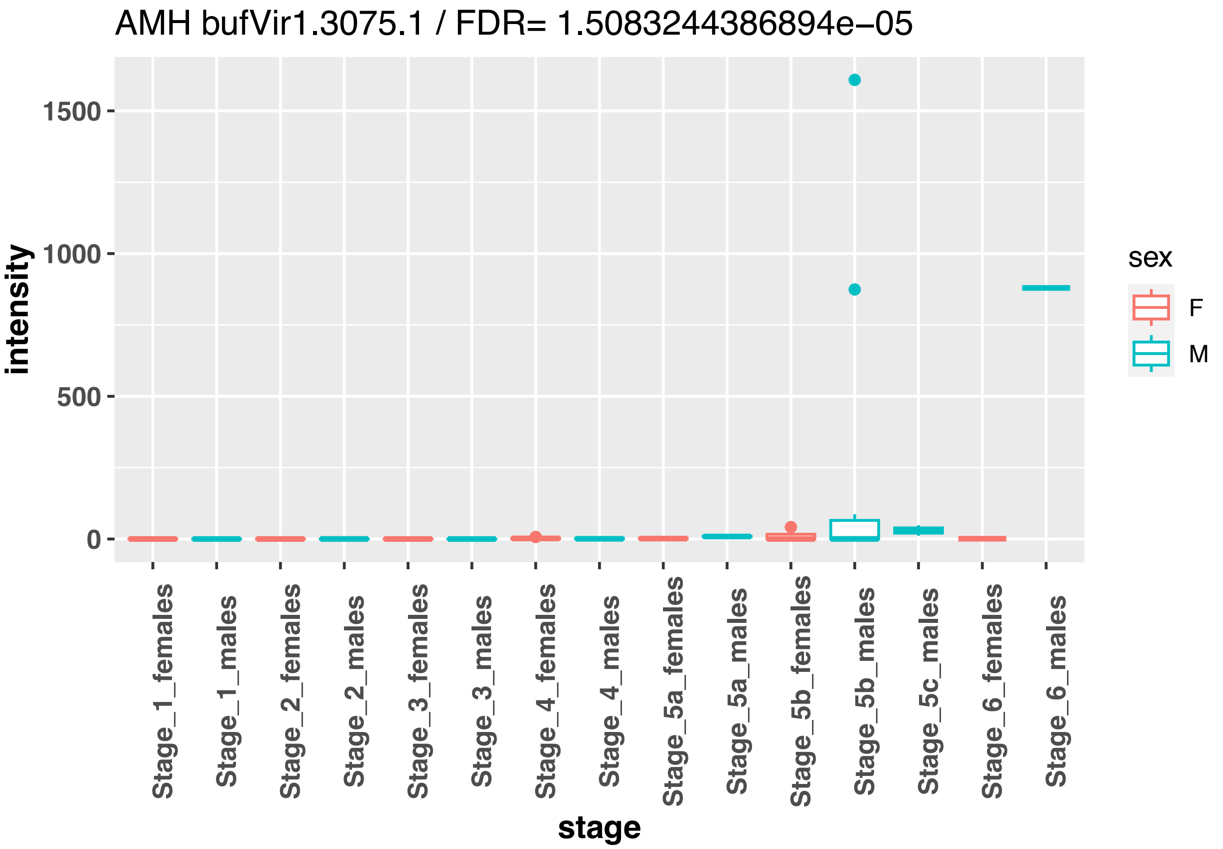
AMH bufVir1.3075.1 / FDR= 1.5083244386894e−05
intensity
1500
1000
500
0
Stage_1_females
Stage_1_males
Stage_2_females
Stage_2_males
Stage_3_females
Stage_3_males
Stage_4_females
Stage_4_males
Stage_5a_females
Stage_5a_males
Stage_5b_females
Stage_5b_males
Stage_5c_males
Stage_6_females
Stage_6_males
stage
sex
F
M

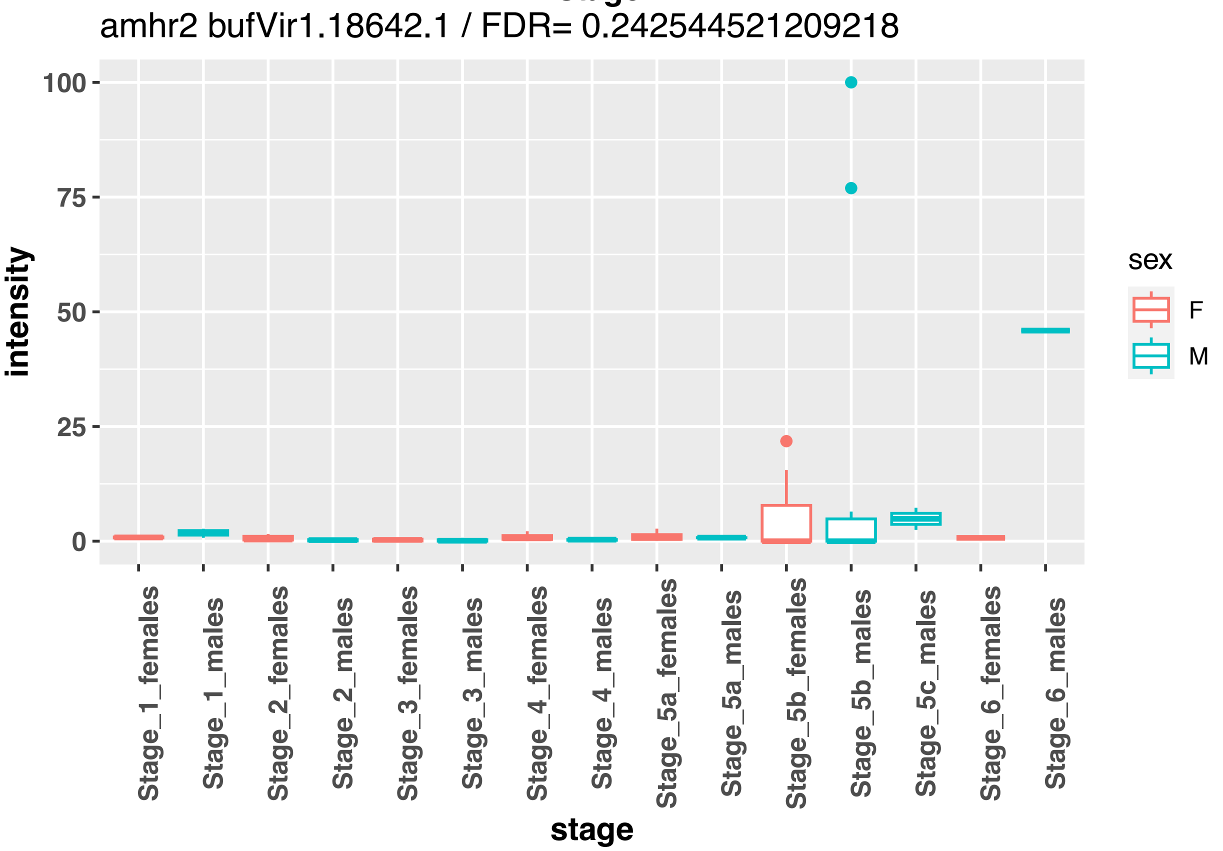
amhr2 bufVir1.18642.1 / FDR= 0.242544521209218
intensity
100
75
50
25
0
Stage_1_females
Stage_1_males
Stage_2_females
Stage_2_males
Stage_3_females
Stage_3_males
Stage_4_females
Stage_4_males
Stage_5a_females
Stage_5a_males
Stage_5b_females
Stage_5b_males
Stage_5c_males
Stage_6_females
Stage_6_males
stage
sex
F
M

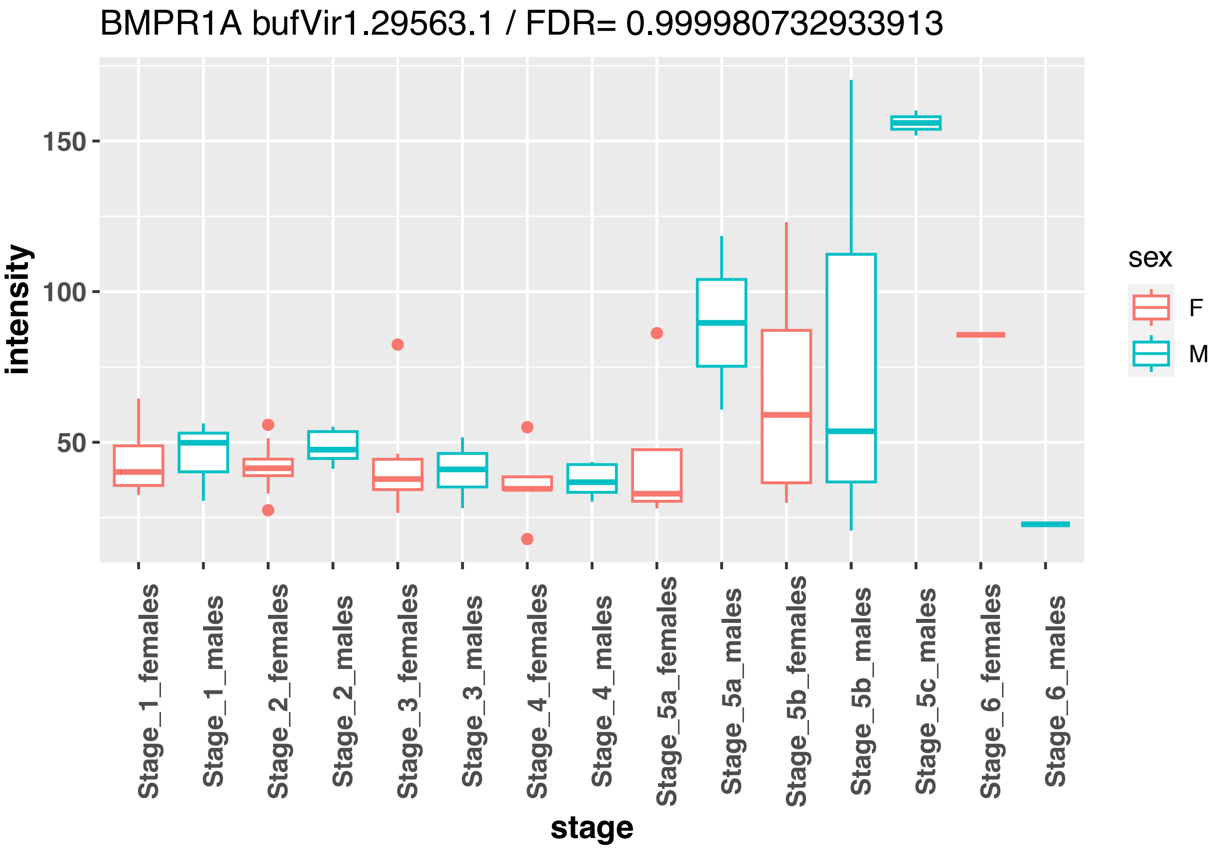
BMPR1A bufVir1.29563.1 / FDR= 0.999980732933913
intensity
150
100
50
stage
Stage_1_females
Stage_1_males
Stage_2_females
Stage_2_males
Stage_3_females
Stage_3_males
Stage_4_females
Stage_4_males
Stage_5a_females
Stage_5a_males
Stage_5b_females
Stage_5b_males
Stage_5c_males
Stage_6_females
Stage_6_males
sex
F
M

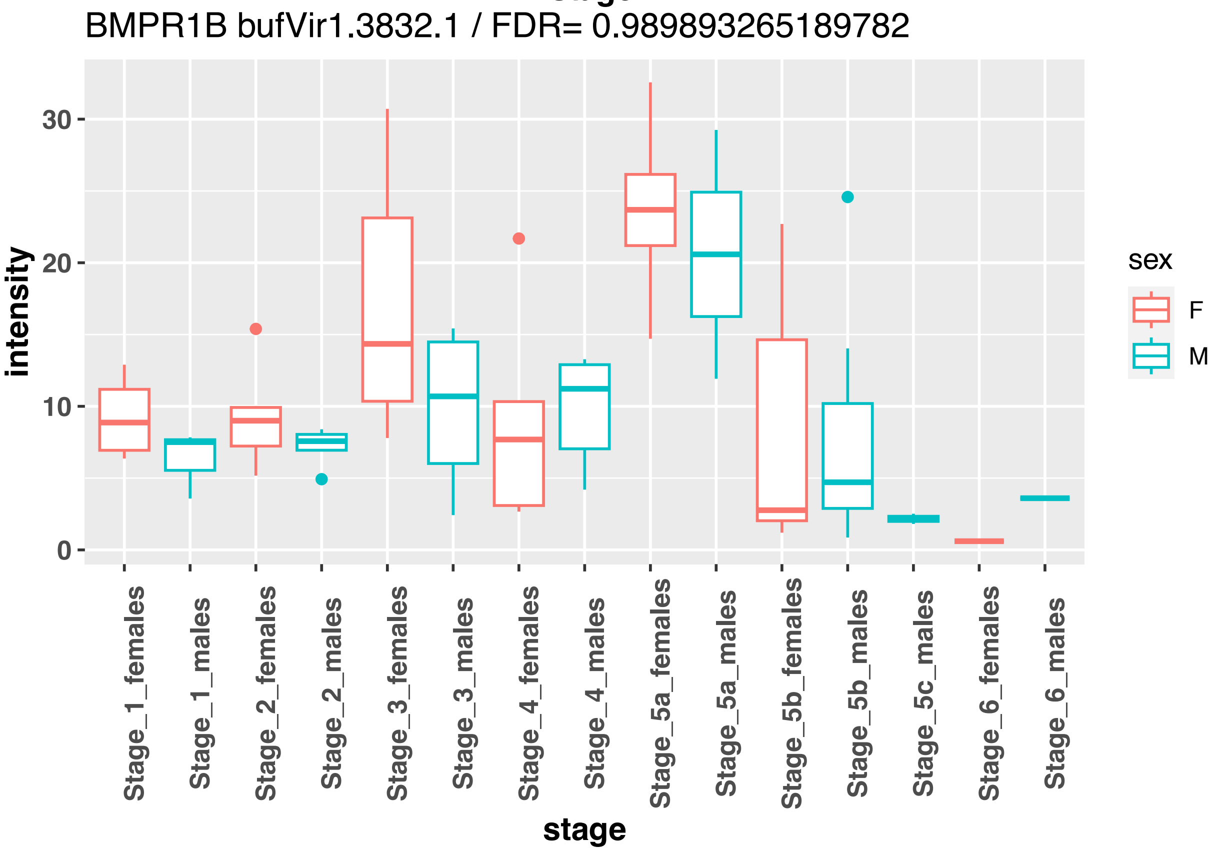
BMPR1B bufVir1.3832.1 / FDR= 0.989893265189782
intensity
30
20
10
0
stage
Stage_1_females
Stage_1_males
Stage_2_females
Stage_2_males
Stage_3_females
Stage_3_males
Stage_4_females
Stage_4_males
Stage_5a_females
Stage_5a_males
Stage_5b_females
Stage_5b_males
Stage_5c_males
Stage_6_females
Stage_6_males
sex
F
M

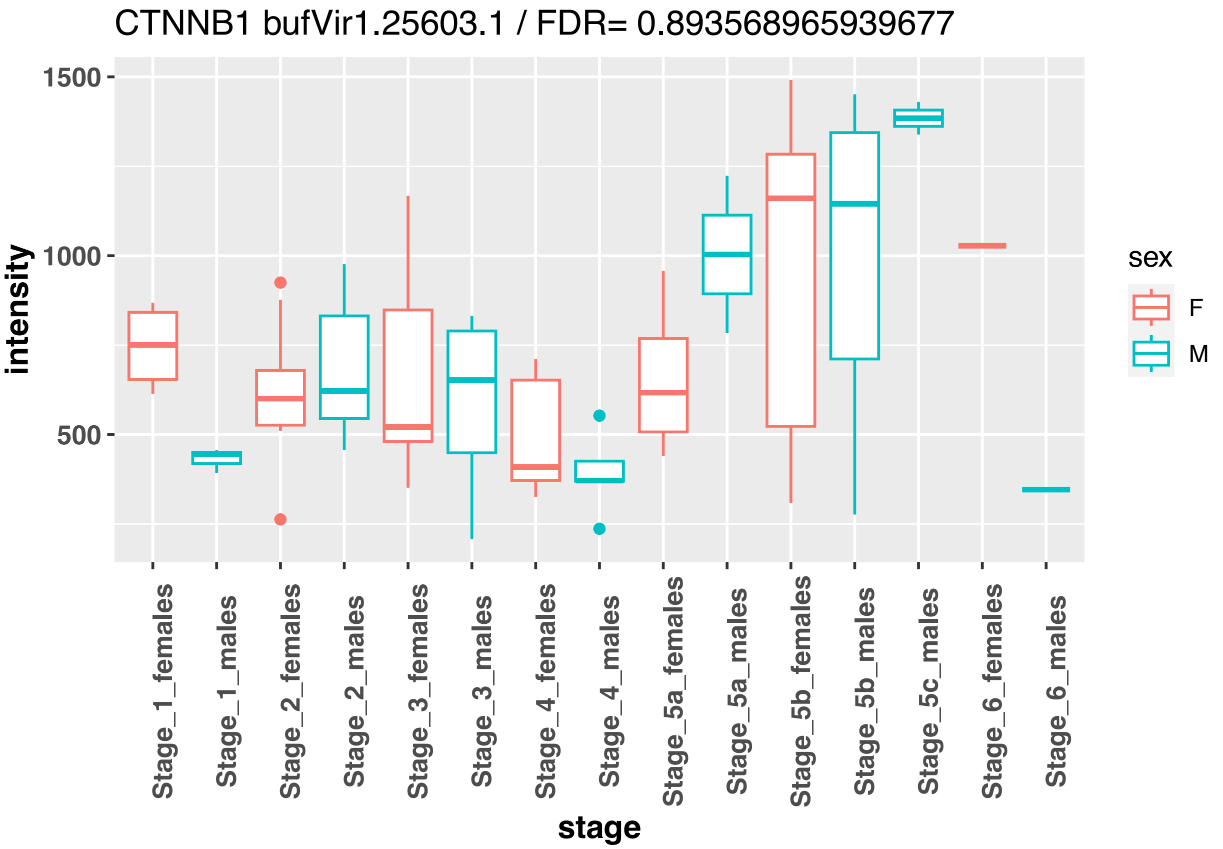
CTNNB1 bufVir1.25603.1 / FDR= 0.893568965939677
intensity
1500
1000
500
sex
F
M
Stage_1_females
Stage_1_males
Stage_2_females
Stage_2_males
Stage_3_females
Stage_3_males
Stage_4_females
Stage_4_males
Stage_5a_females
Stage_5a_males
Stage_5b_females
Stage_5b_males
Stage_5c_males
Stage_6_females
Stage_6_males
stage

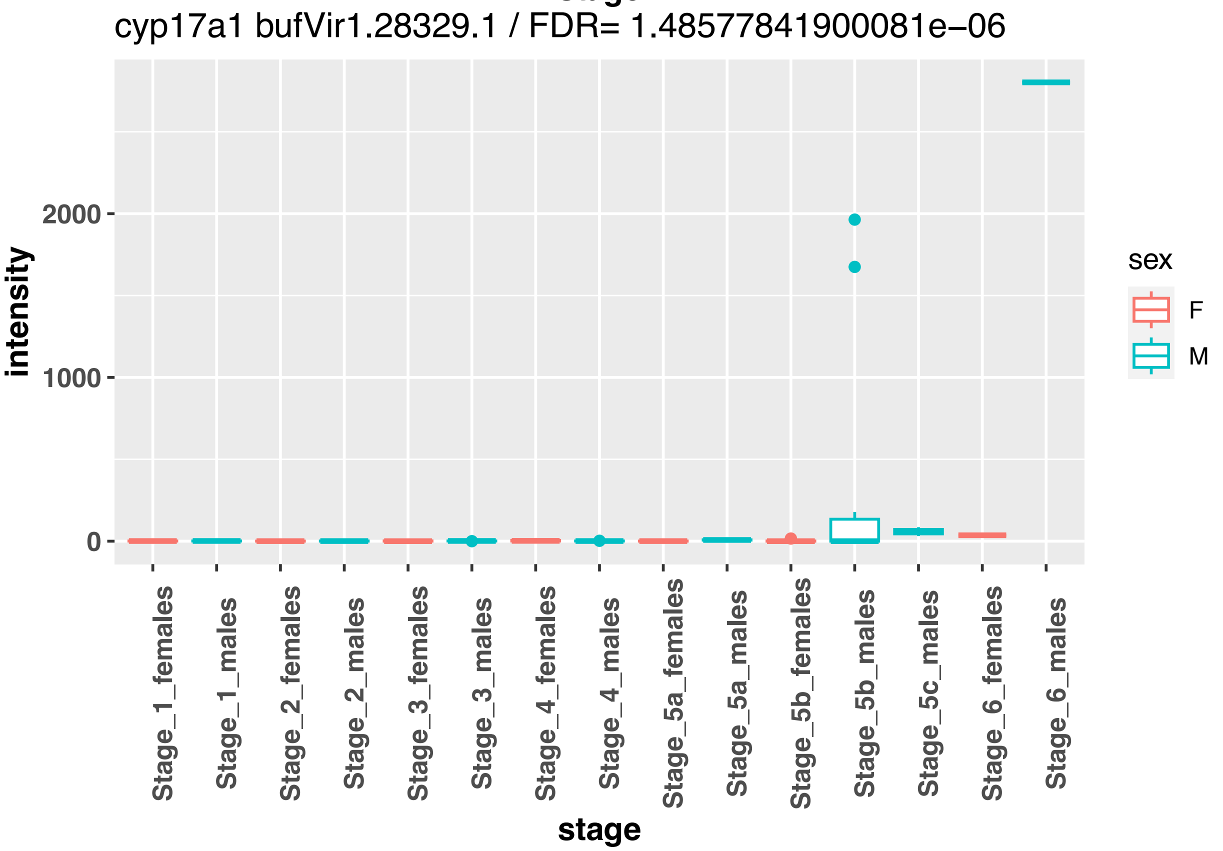
cyp17a1 bufVir1.28329.1 / FDR= 1.48577841900081e−06
intensity
2000
1000
0
sex
F
M
Stage_1_females
Stage_1_males
Stage_2_females
Stage_2_males
Stage_3_females
Stage_3_males
Stage_4_females
Stage_4_males
Stage_5a_females
Stage_5a_males
Stage_5b_females
Stage_5b_males
Stage_5c_males
Stage_6_females
Stage_6_males
stage

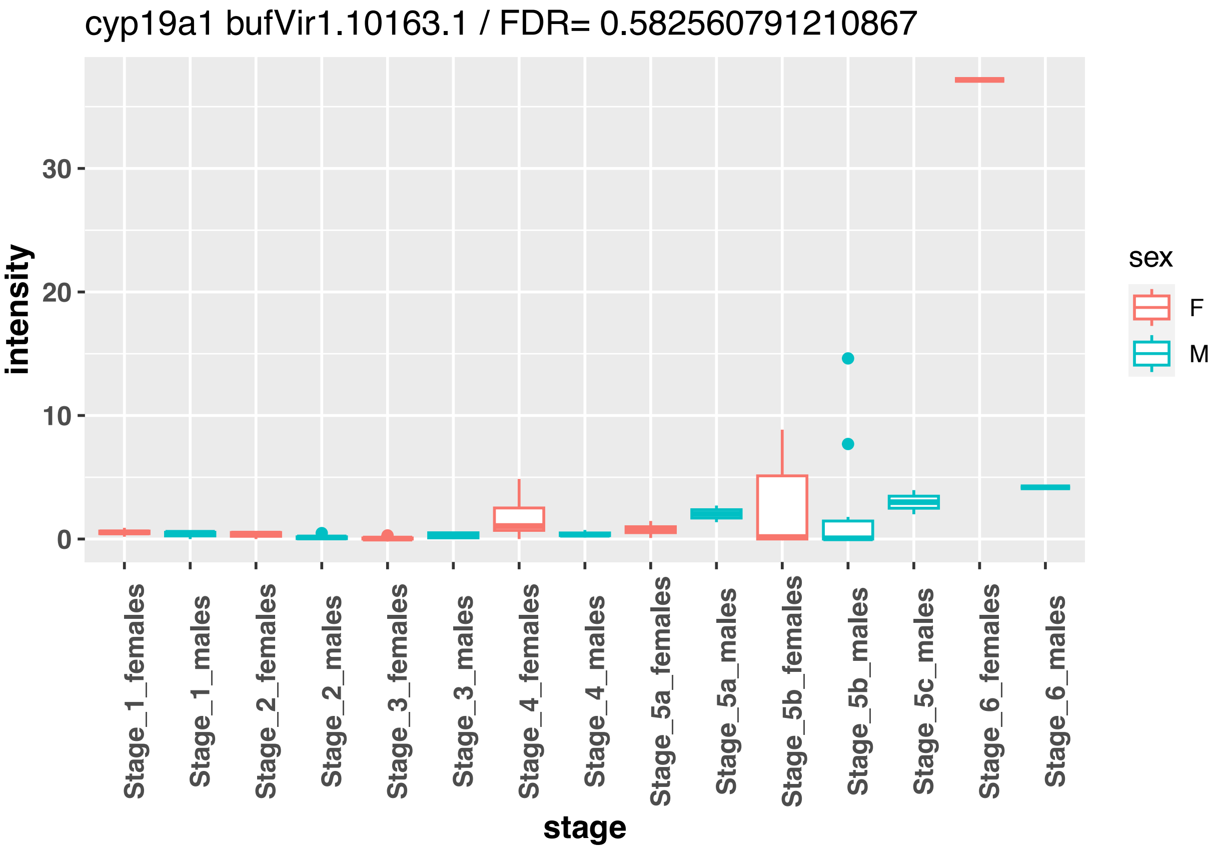
cyp19a1 bufVir1.10163.1 / FDR= 0.582560791210867
intensity
30
20
10
0
sex
F
M
Stage_1_females
Stage_1_males
Stage_2_females
Stage_2_males
Stage_3_females
Stage_3_males
Stage_4_females
Stage_4_males
Stage_5a_females
Stage_5a_males
Stage_5b_females
Stage_5b_males
Stage_5c_males
Stage_6_females
Stage_6_males
stage

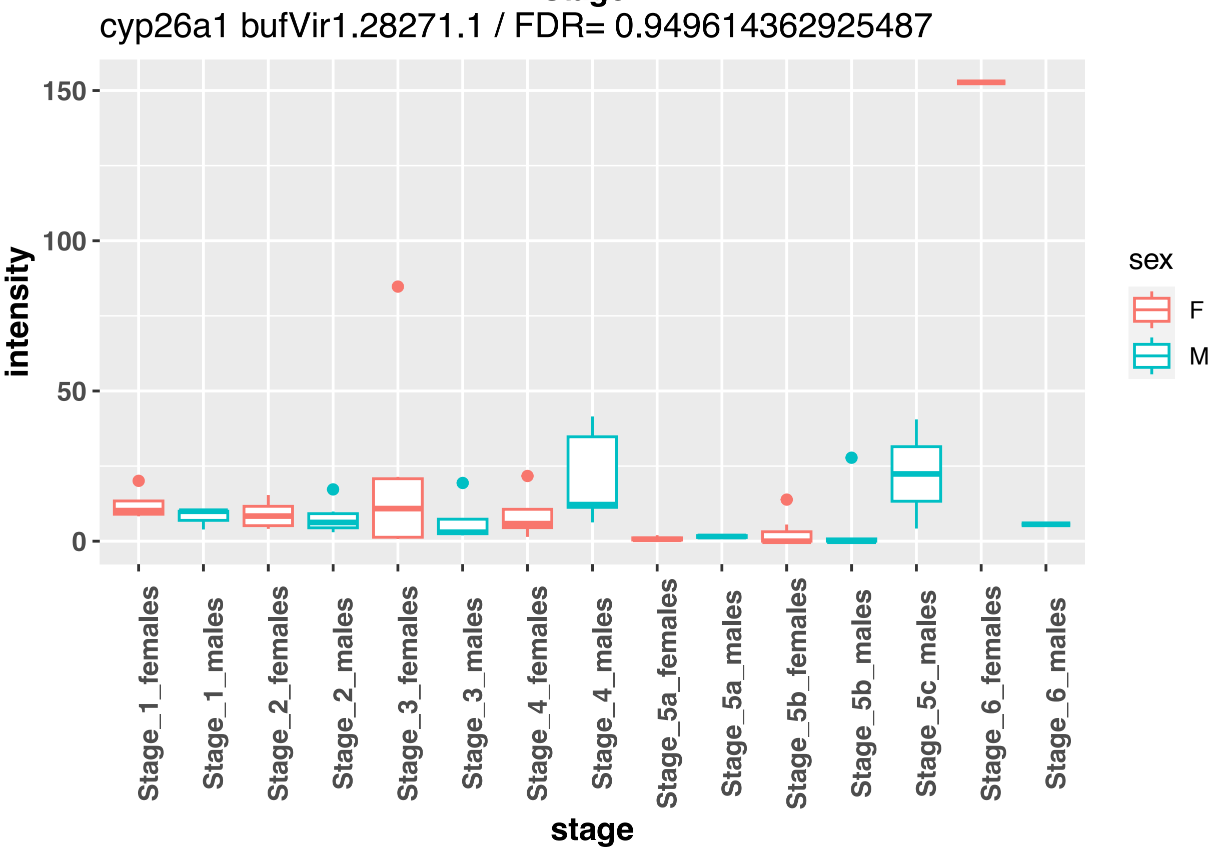
cyp26a1 bufVir1.28271.1 / FDR= 0.949614362925487
intensity
150
100
50
0
sex
F
M
Stage_1_females
Stage_1_males
Stage_2_females
Stage_2_males
Stage_3_females
Stage_3_males
Stage_4_females
Stage_4_males
Stage_5a_females
Stage_5a_males
Stage_5b_females
Stage_5b_males
Stage_5c_males
Stage_6_females
Stage_6_males
stage

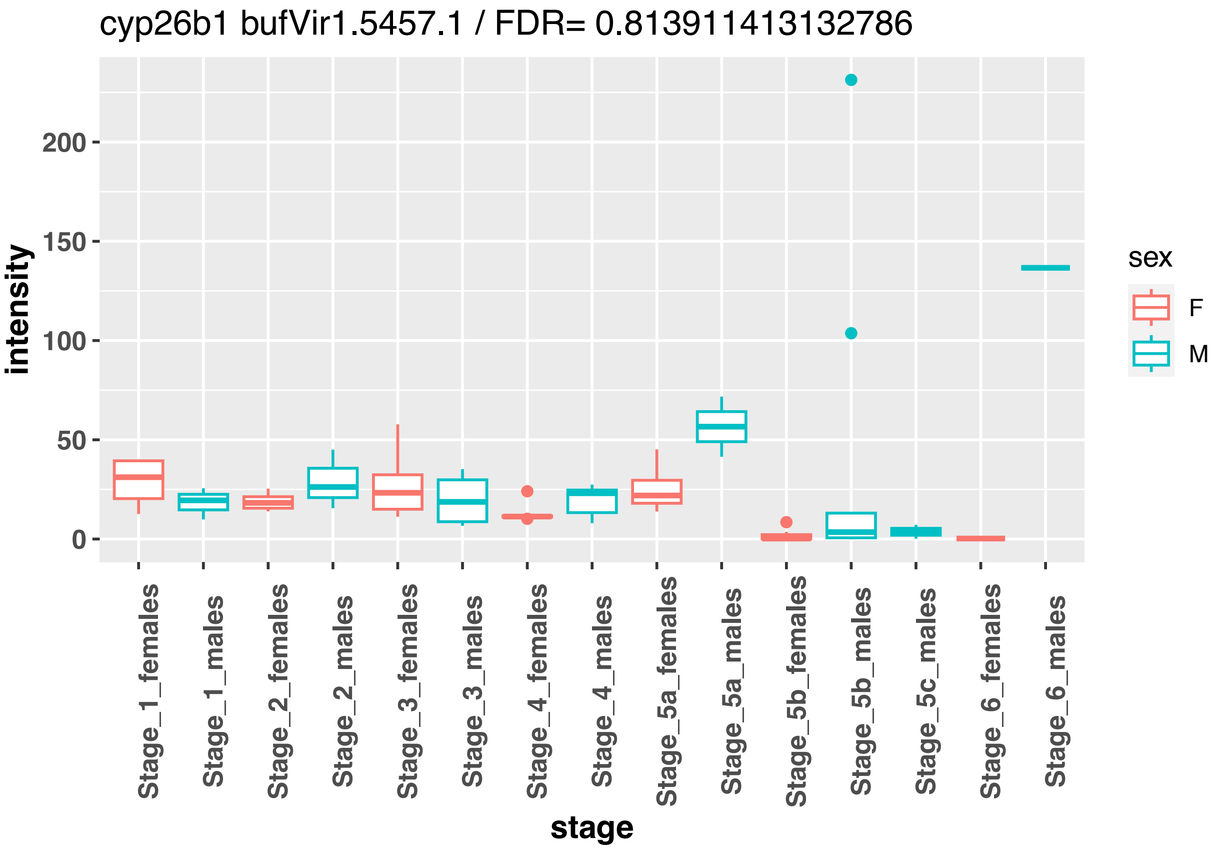
cyp26b1 bufVir1.5457.1 / FDR= 0.813911413132786
intensity
200
150
100
50
0
Stage_1_females
Stage_1_males
Stage_2_females
Stage_2_males
Stage_3_females
Stage_3_males
Stage_4_females
Stage_4_males
Stage_5a_females
Stage_5a_males
Stage_5b_females
Stage_5b_males
Stage_5c_males
Stage_6_females
Stage_6_males
stage
sex
F
M

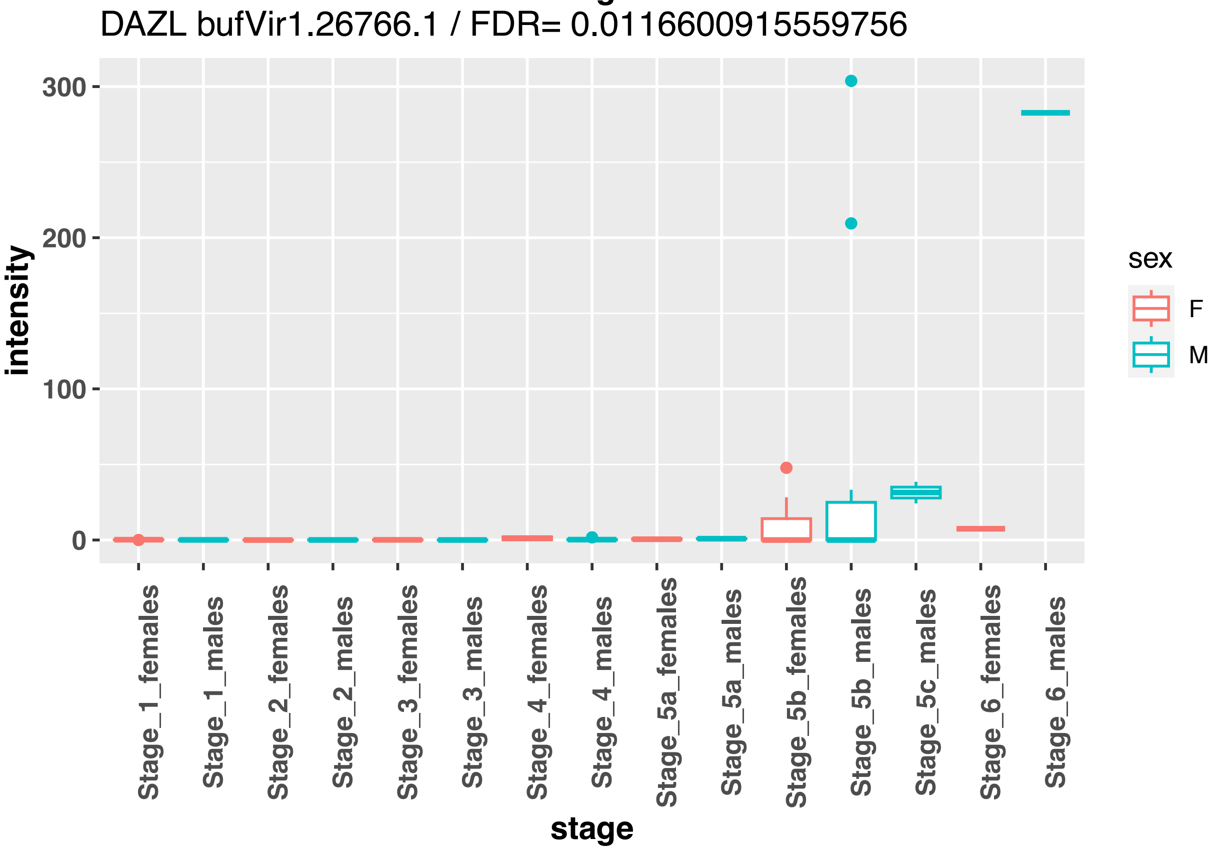
DAZL bufVir1.26766.1 / FDR= 0.0116600915559756
intensity
300
200
100
0
Stage_1_females
Stage_1_males
Stage_2_females
Stage_2_males
Stage_3_females
Stage_3_males
Stage_4_females
Stage_4_males
Stage_5a_females
Stage_5a_males
Stage_5b_females
Stage_5b_males
Stage_5c_males
Stage_6_females
Stage_6_males
stage
sex
F
M

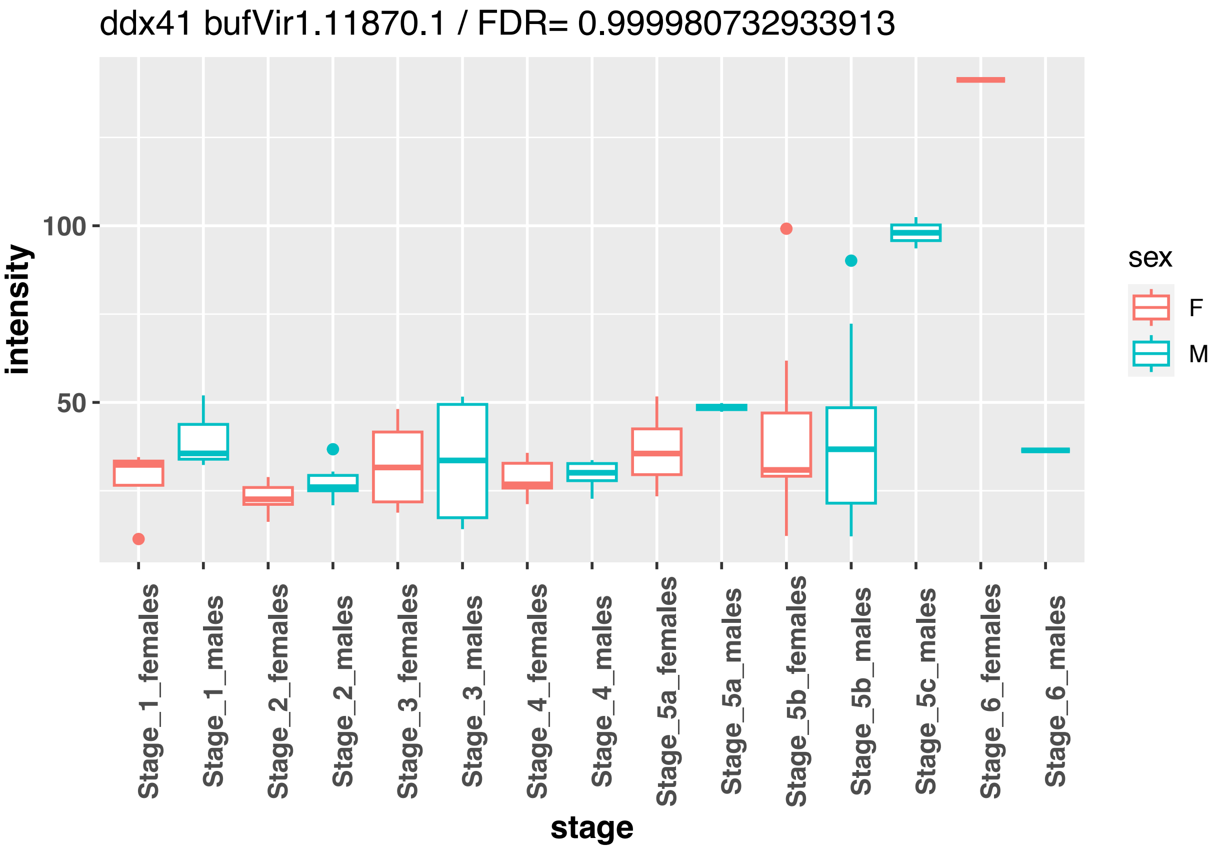
ddx41 bufVir1.11870.1 / FDR= 0.999980732933913
intensity
100
50
stage
Stage_1_females
Stage_1_males
Stage_2_females
Stage_2_males
Stage_3_females
Stage_3_males
Stage_4_females
Stage_4_males
Stage_5a_females
Stage_5a_males
Stage_5b_females
Stage_5b_males
Stage_5c_males
Stage_6_females
Stage_6_males
sex
F
M

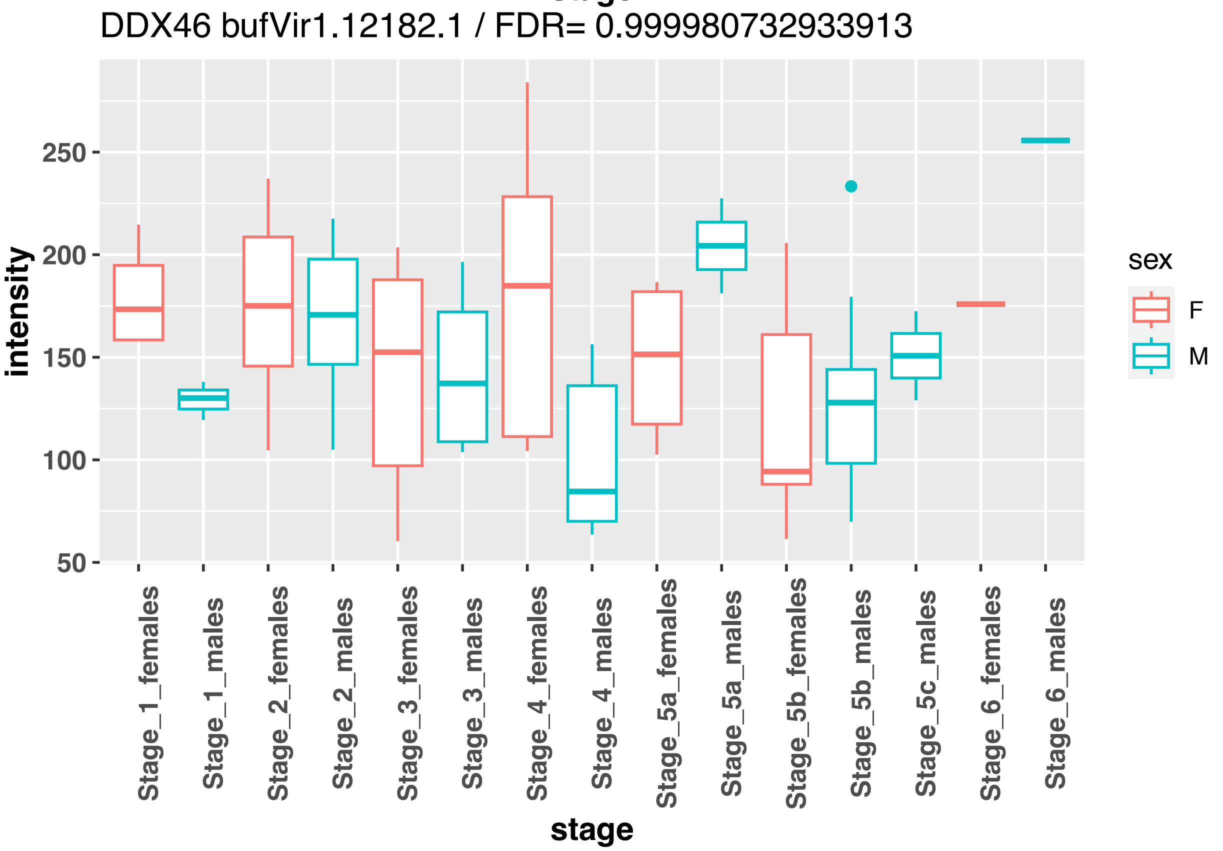
DDX46 bufVir1.12182.1 / FDR= 0.999980732933913
intensity
250
200
150
100
50
stage
Stage_1_females
Stage_1_males
Stage_2_females
Stage_2_males
Stage_3_females
Stage_3_males
Stage_4_females
Stage_4_males
Stage_5a_females
Stage_5a_males
Stage_5b_females
Stage_5b_males
Stage_5c_males
Stage_6_females
Stage_6_males
sex
F
M

DDX43 bufVir1.21784.1 / FDR= 0.561875147054289

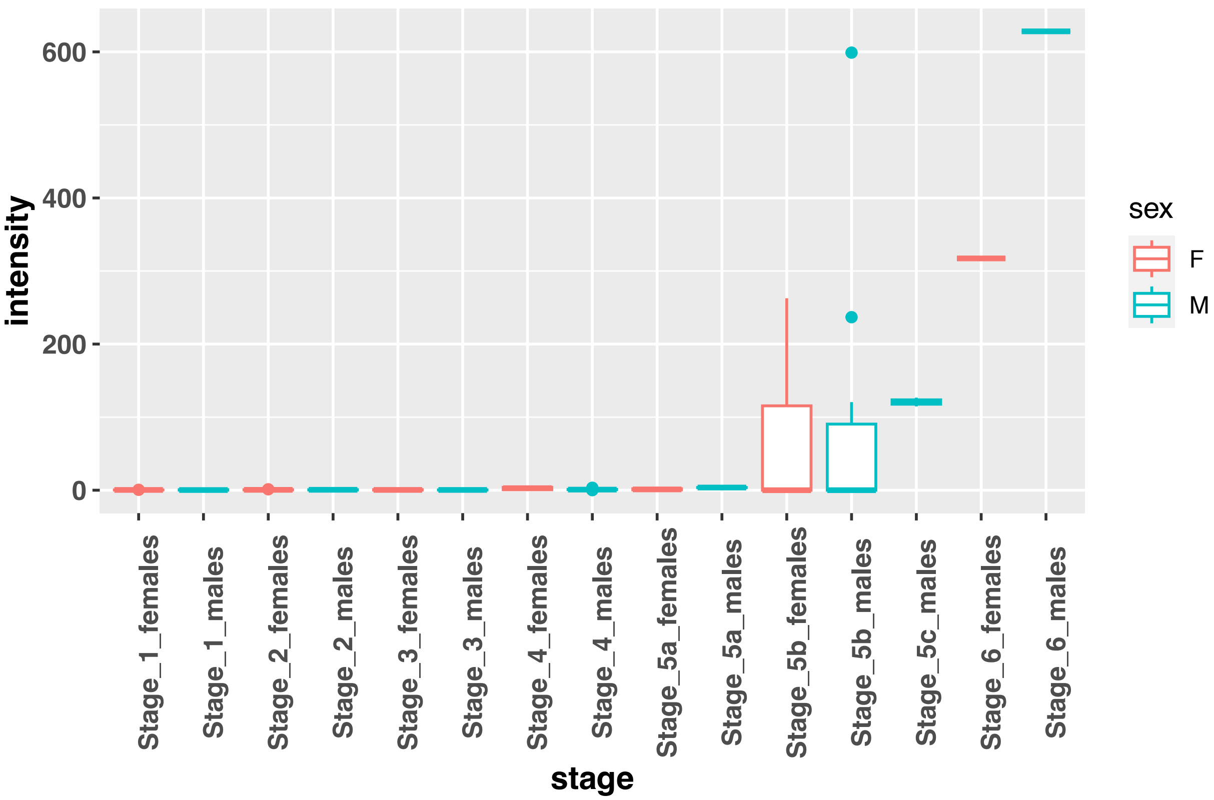


ddx49 bufVir1.2807.1 / FDR= 0.934199034864266

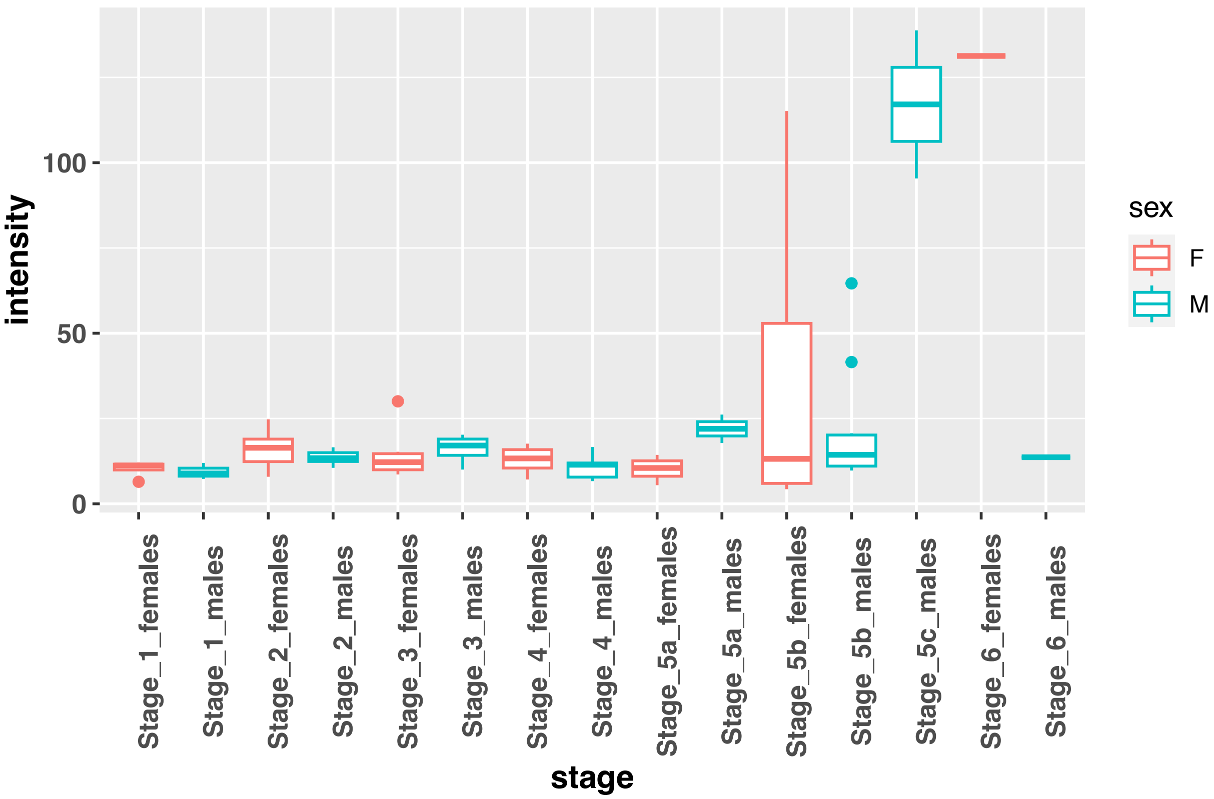

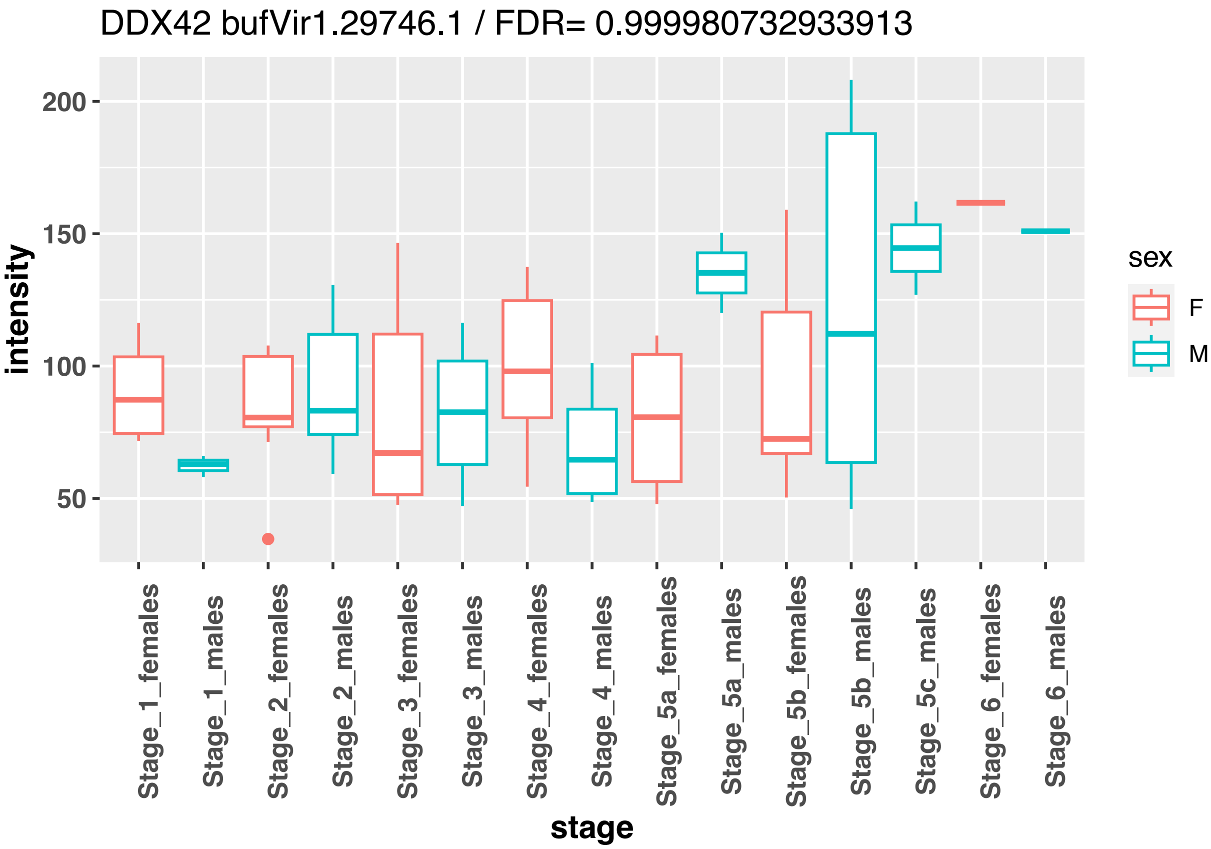
DDX42 bufVir1.29746.1 / FDR= 0.999980732933913
intensity
200
150
100
50
stage
Stage_1_females
Stage_1_males
Stage_2_females
Stage_2_males
Stage_3_females
Stage_3_males
Stage_4_females
Stage_4_males
Stage_5a_females
Stage_5a_males
Stage_5b_females
Stage_5b_males
Stage_5c_males
Stage_6_females
Stage_6_males
sex
F
M

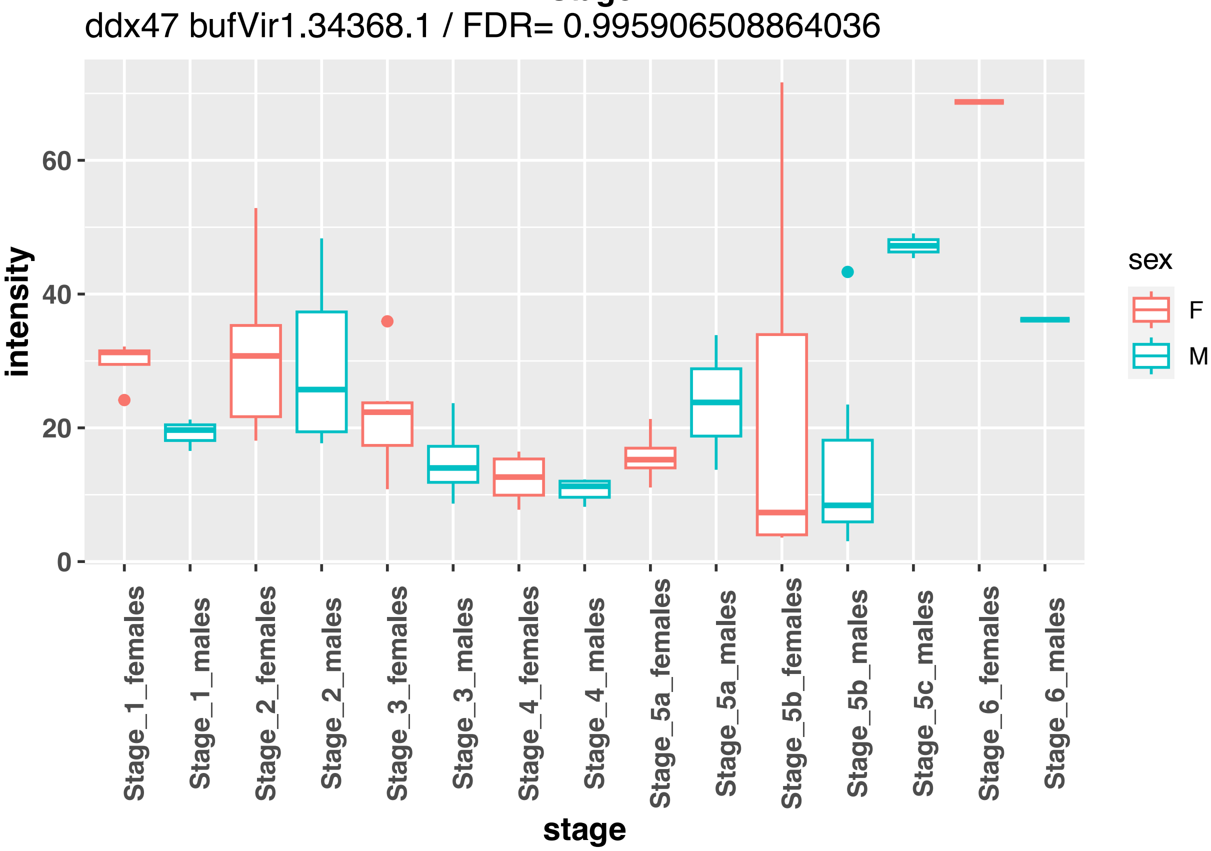
ddx47 bufVir1.34368.1 / FDR= 0.995906508864036
intensity
60
40
20
0
stage
Stage_1_females
Stage_1_males
Stage_2_females
Stage_2_males
Stage_3_females
Stage_3_males
Stage_4_females
Stage_4_males
Stage_5a_females
Stage_5a_males
Stage_5b_females
Stage_5b_males
Stage_5c_males
Stage_6_females
Stage_6_males
sex
F
M

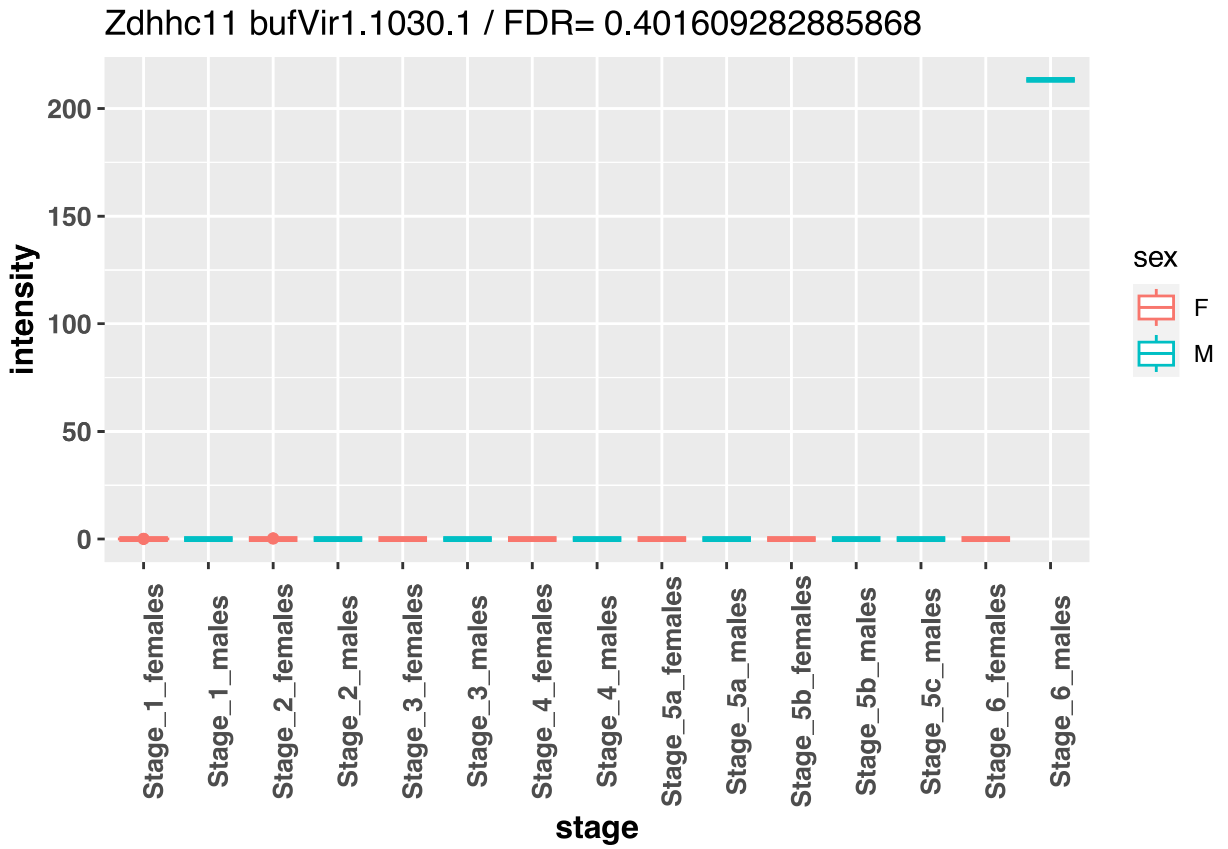
Zdhhc11 bufVir1.1030.1 / FDR= 0.401609282885868
intensity
200
150
100
50
0
stage
Stage_1_females
Stage_1_males
Stage_2_females
Stage_2_males
Stage_3_females
Stage_3_males
Stage_4_females
Stage_4_males
Stage_5a_females
Stage_5a_males
Stage_5b_females
Stage_5b_males
Stage_5c_males
Stage_6_females
Stage_6_males
sex
F
M

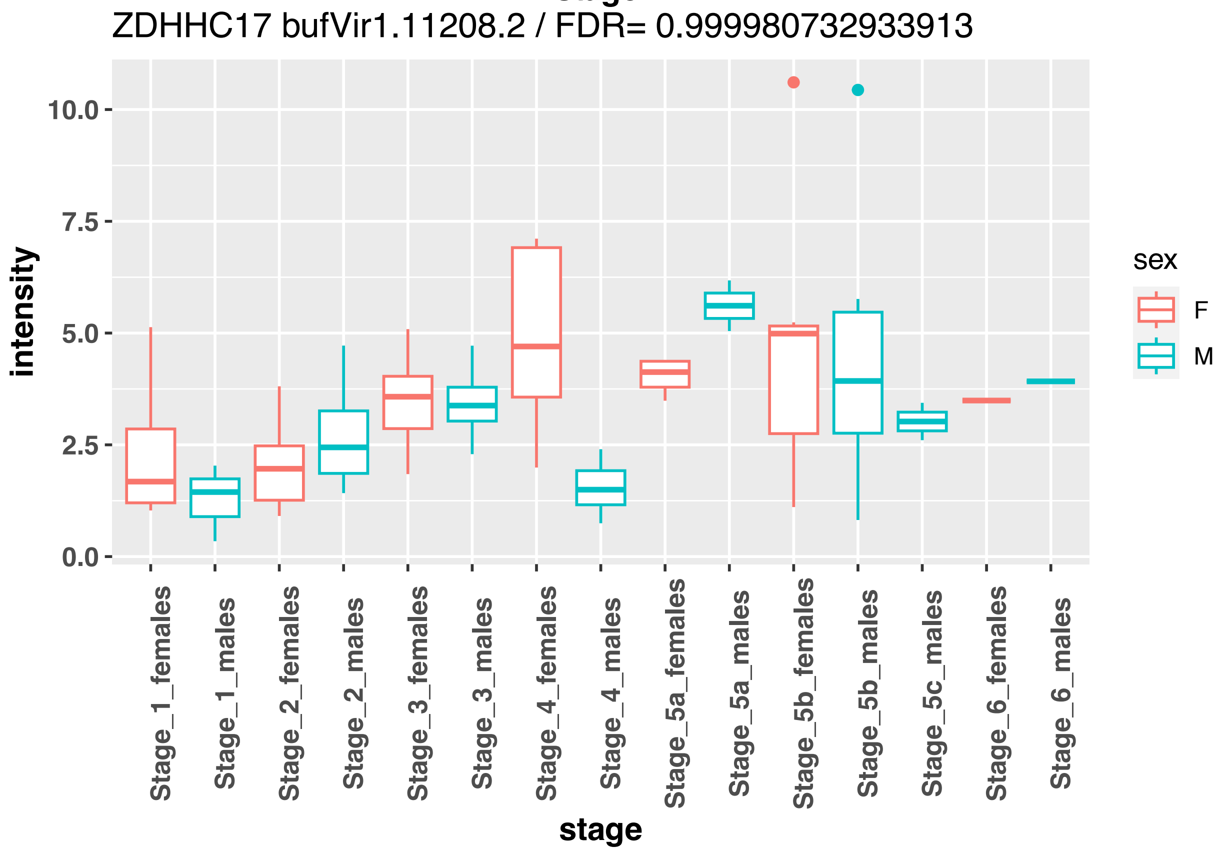
ZDHHC17 bufVir1.11208.2 / FDR= 0.999980732933913
intensity
10.0
7.5
5.0
2.5
0.0
stage
Stage_1_females
Stage_1_males
Stage_2_females
Stage_2_males
Stage_3_females
Stage_3_males
Stage_4_females
Stage_4_males
Stage_5a_females
Stage_5a_males
Stage_5b_females
Stage_5b_males
Stage_5c_males
Stage_6_females
Stage_6_males
sex
F
M

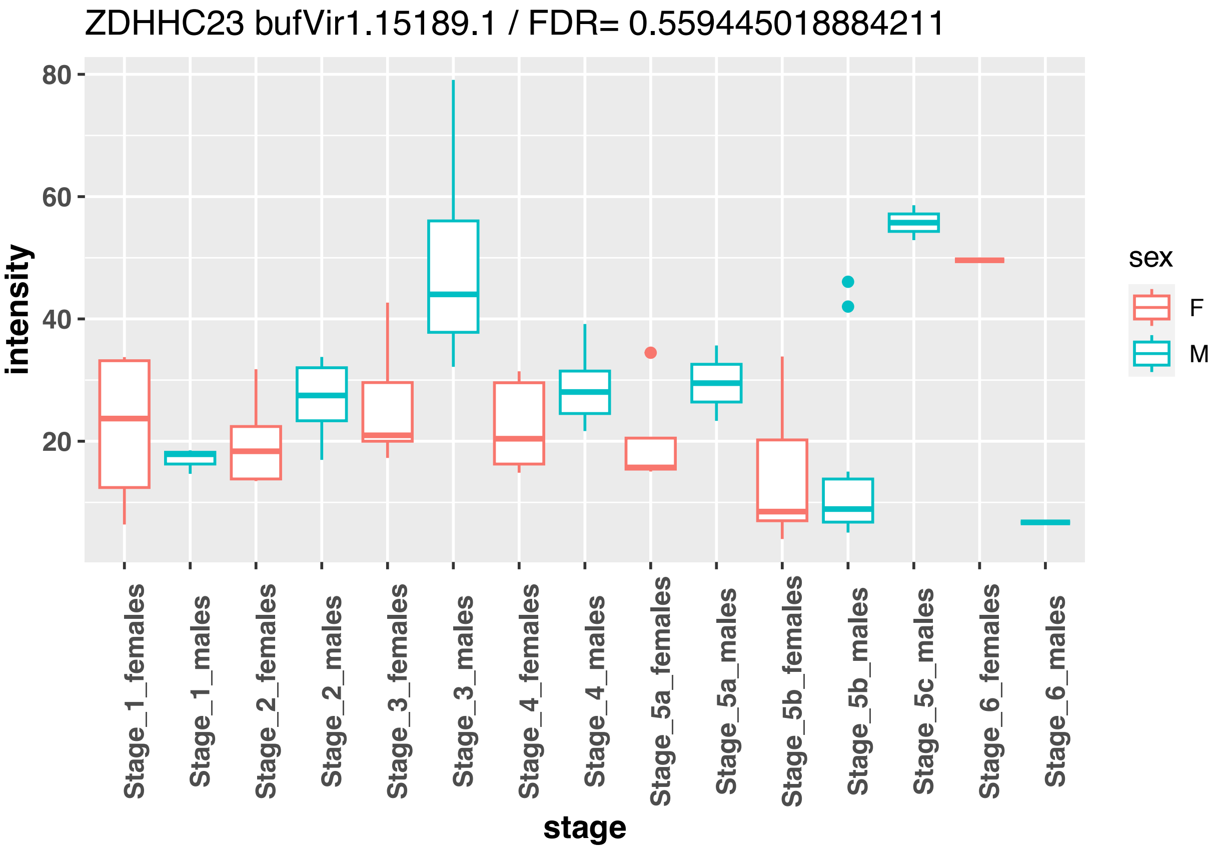
ZDHHC23 bufVir1.15189.1 / FDR= 0.559445018884211
intensity
80
60
40
20
Stage_1_females
Stage_1_males
Stage_2_females
Stage_2_males
Stage_3_females
Stage_3_males
Stage_4_females
Stage_4_males
Stage_5a_females
Stage_5a_males
Stage_5b_females
Stage_5b_males
Stage_5c_males
Stage_6_females
Stage_6_males
stage
sex
F
M

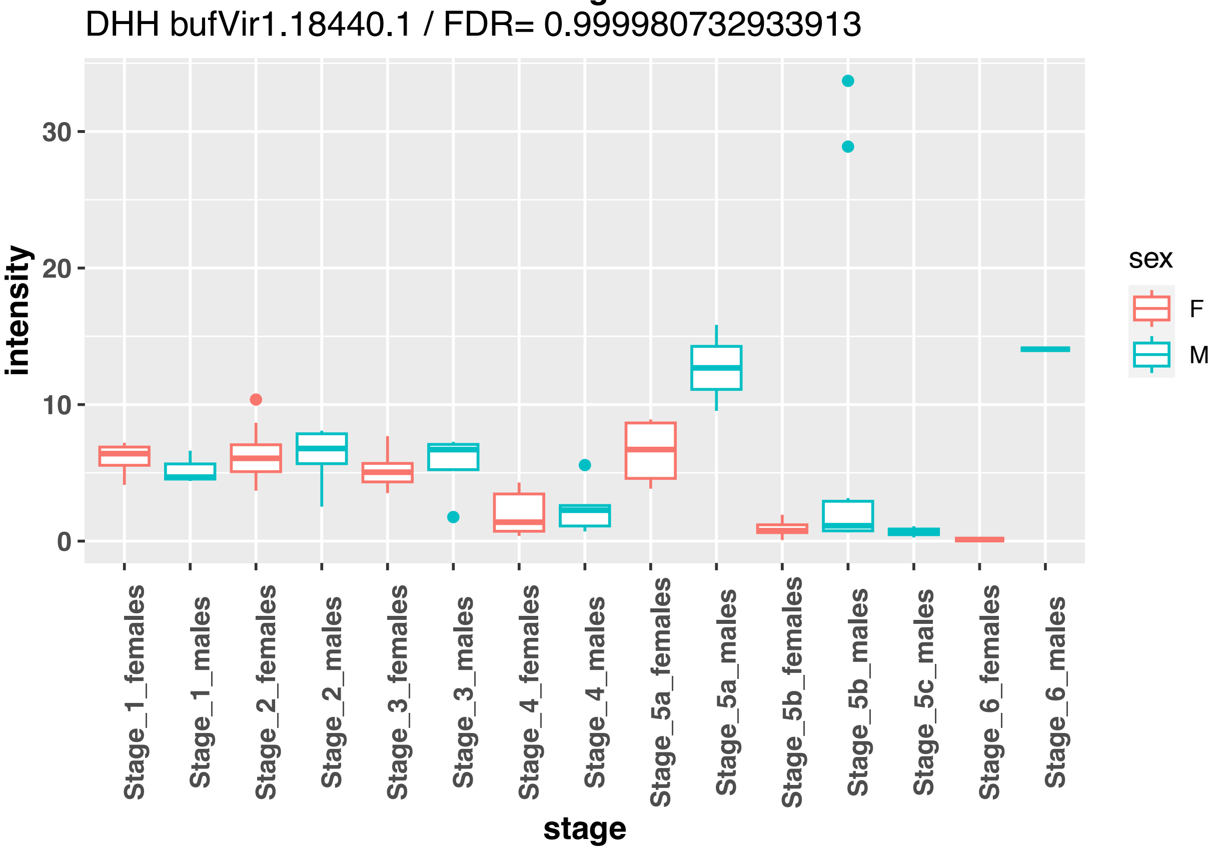
DHH bufVir1.18440.1 / FDR= 0.999980732933913
intensity
30
20
10
0
Stage_1_females
Stage_1_males
Stage_2_females
Stage_2_males
Stage_3_females
Stage_3_males
Stage_4_females
Stage_4_males
Stage_5a_females
Stage_5a_males
Stage_5b_females
Stage_5b_males
Stage_5c_males
Stage_6_females
Stage_6_males
stage
sex
F
M

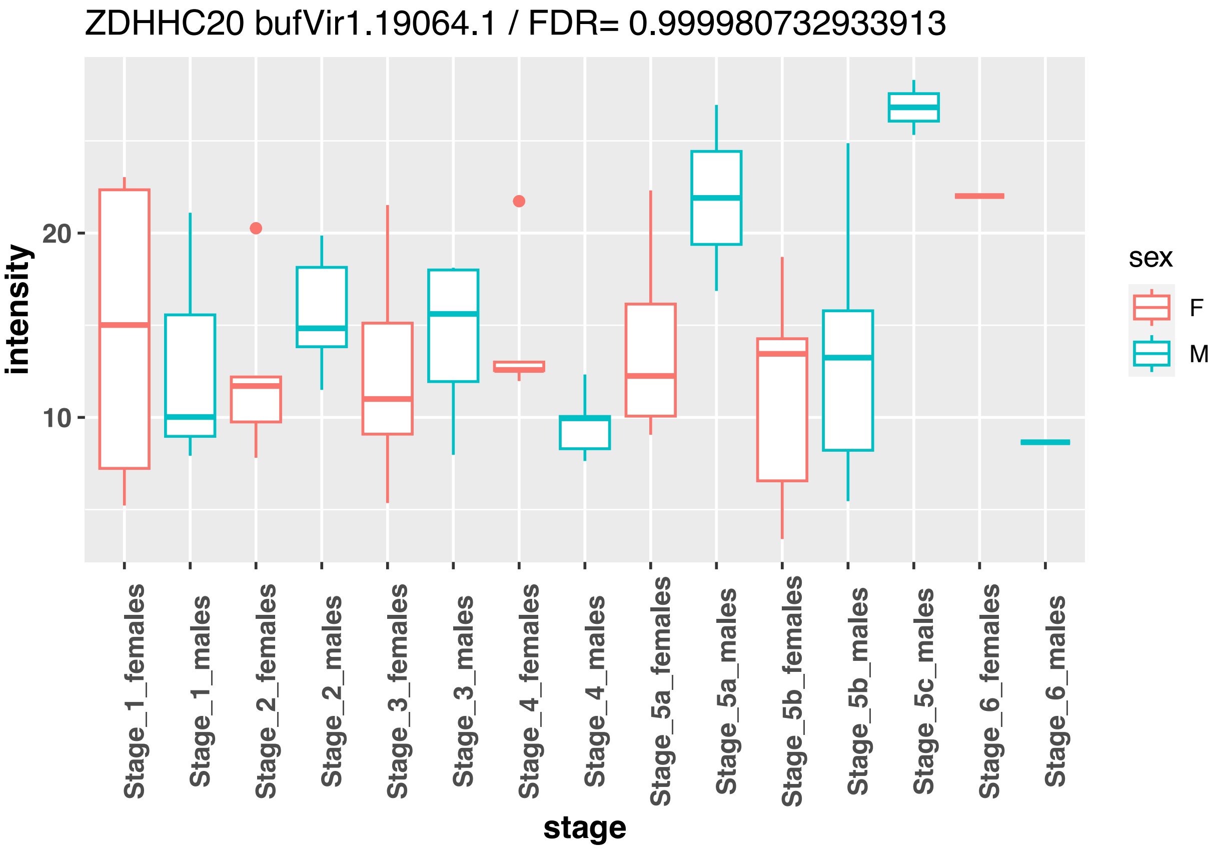
ZDHHC20 bufVir1.19064.1 / FDR= 0.999980732933913
intensity
20
10
Stage_1_females
Stage_1_males
Stage_2_females
Stage_2_males
Stage_3_females
Stage_3_males
Stage_4_females
Stage_4_males
Stage_5a_females
Stage_5a_males
Stage_5b_females
Stage_5b_males
Stage_5c_males
Stage_6_females
Stage_6_males
stage
sex
F
M

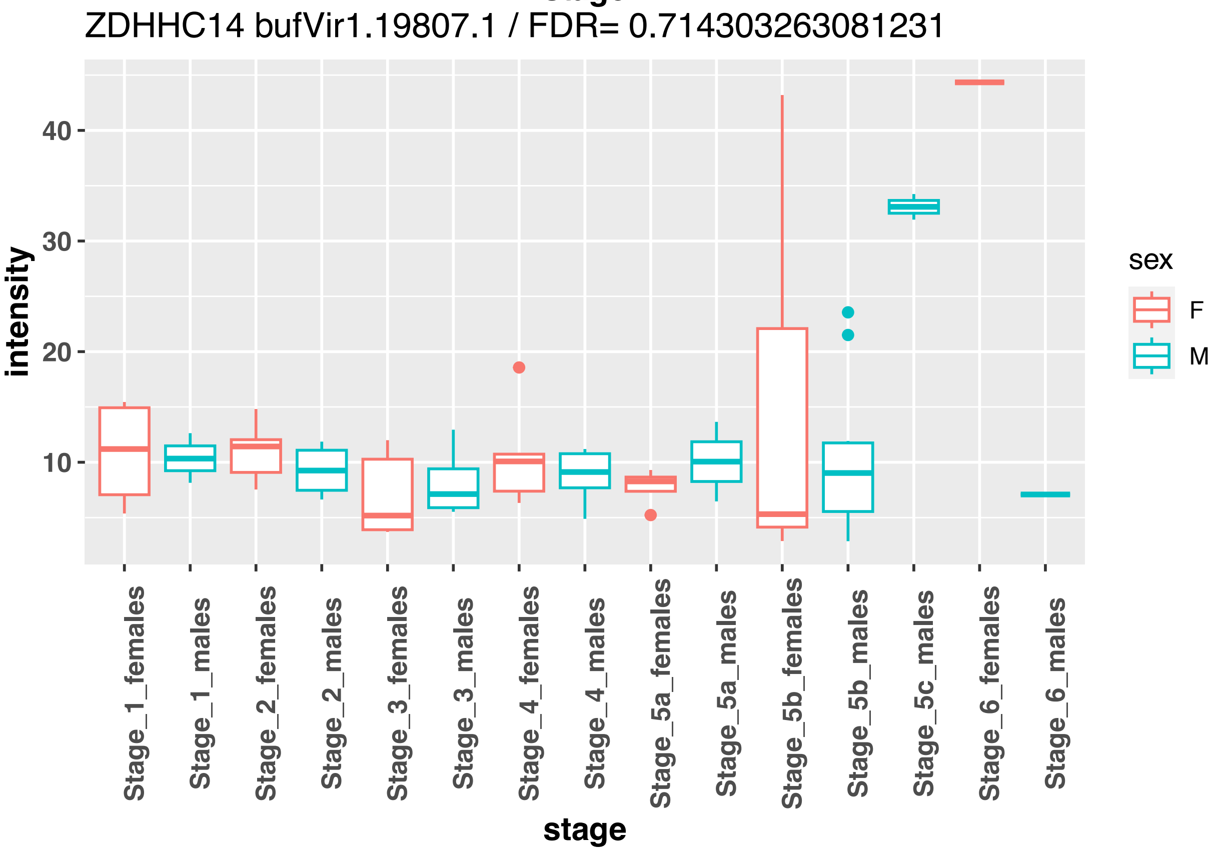
ZDHHC14 bufVir1.19807.1 / FDR= 0.714303263081231
intensity
40
30
20
10
Stage_1_females
Stage_1_males
Stage_2_females
Stage_2_males
Stage_3_females
Stage_3_males
Stage_4_females
Stage_4_males
Stage_5a_females
Stage_5a_males
Stage_5b_females
Stage_5b_males
Stage_5c_males
Stage_6_females
Stage_6_males
stage
sex
F
M

ZDHHC21 bufVir1.2459.1 / FDR= 0.957890412966163

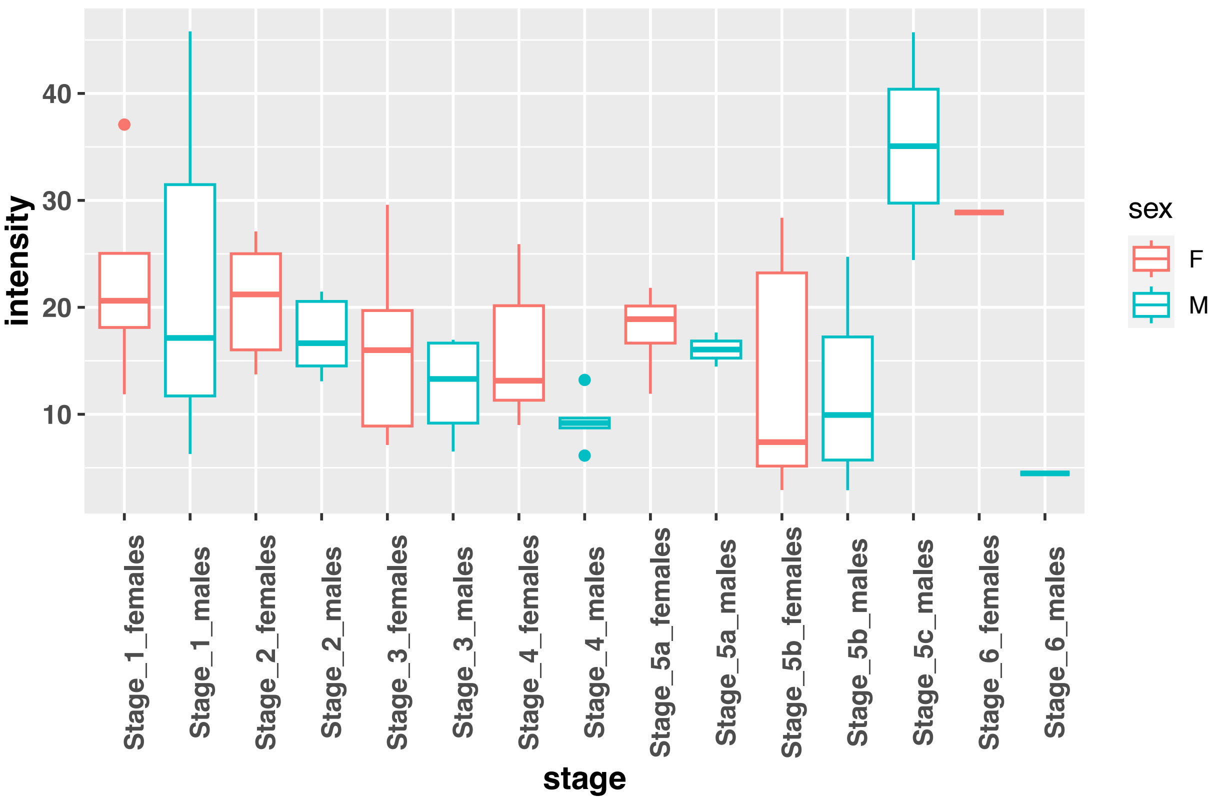


ZDHHC3 bufVir1.25611.2 / FDR= 0.999980732933913

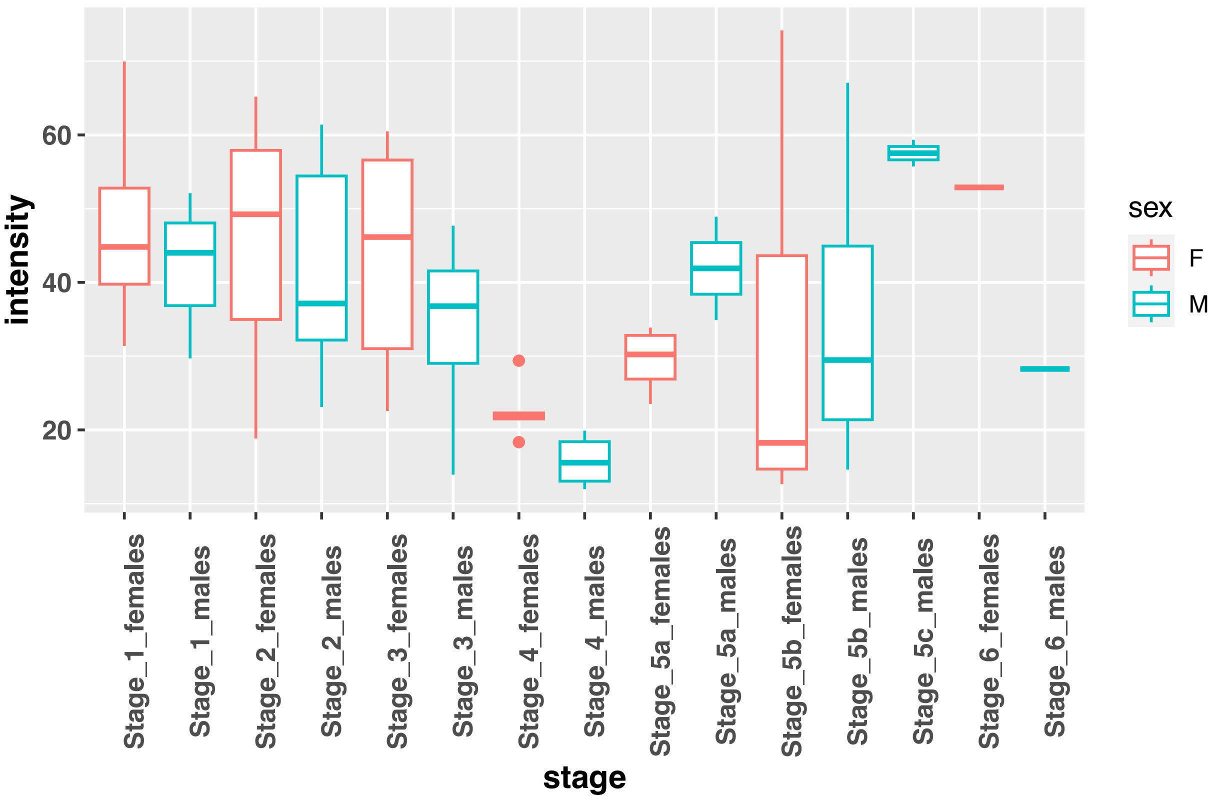

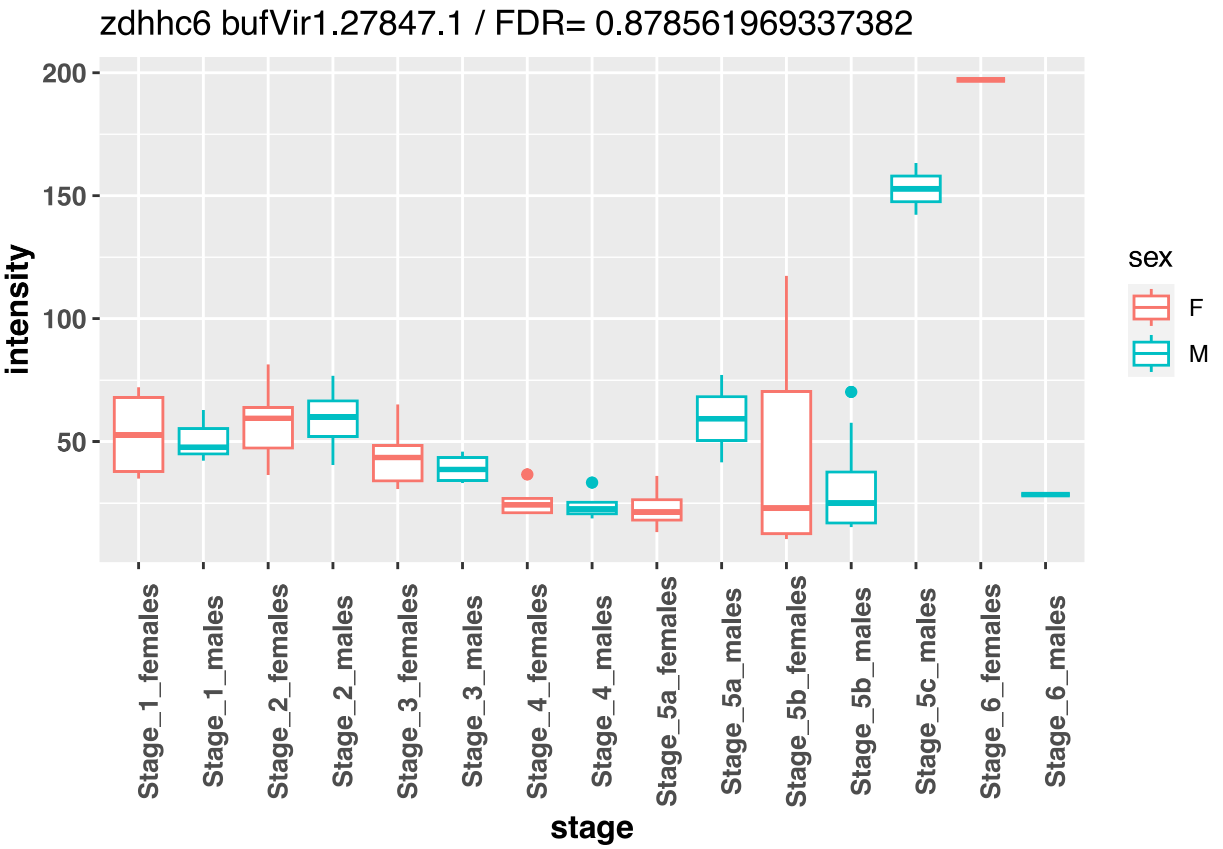
zdhhc6 bufVir1.27847.1 / FDR= 0.878561969337382
intensity
200
150
100
50
stage
Stage_1_females
Stage_1_males
Stage_2_females
Stage_2_males
Stage_3_females
Stage_3_males
Stage_4_females
Stage_4_males
Stage_5a_females
Stage_5a_males
Stage_5b_females
Stage_5b_males
Stage_5c_males
Stage_6_females
Stage_6_males
sex
F
M

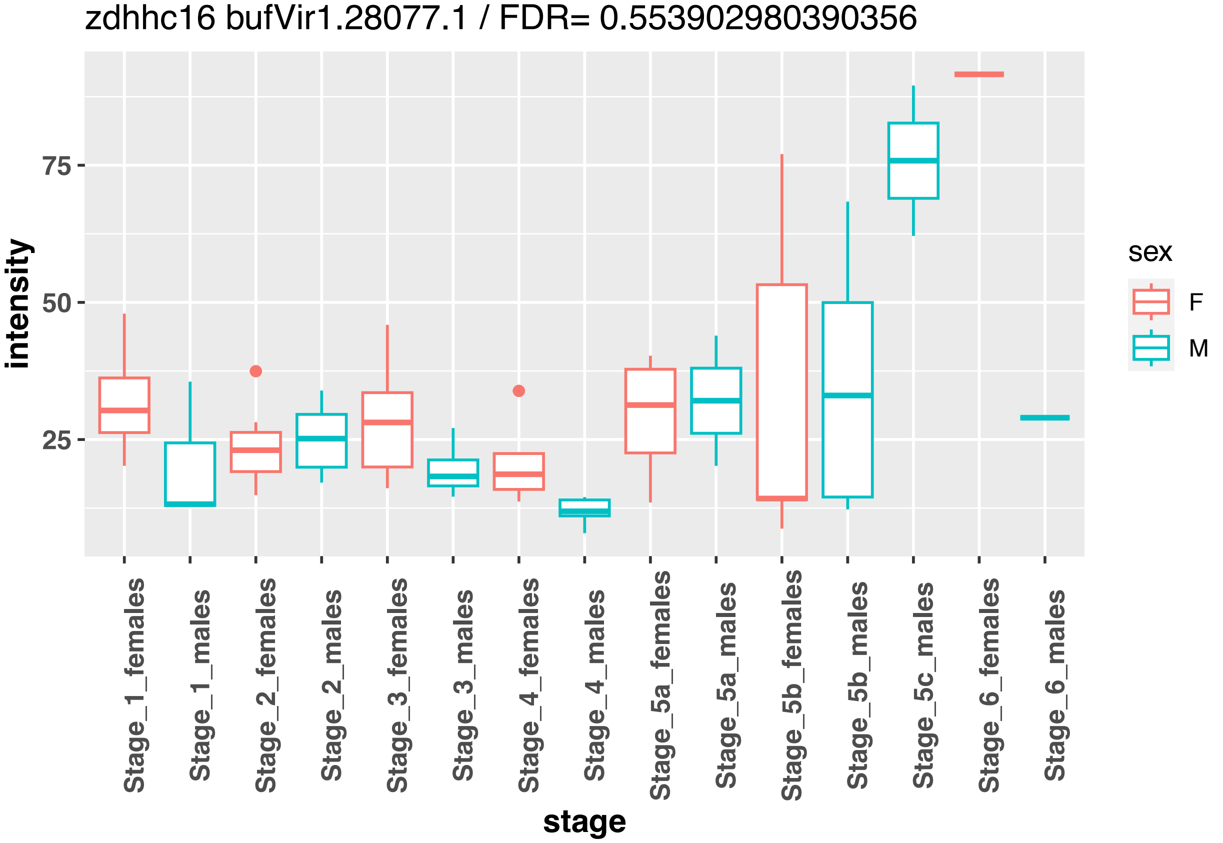
zdhhc16 bufVir1.28077.1 / FDR= 0.553902980390356
intensity
75
50
25
stage
Stage_1_females
Stage_1_males
Stage_2_females
Stage_2_males
Stage_3_females
Stage_3_males
Stage_4_females
Stage_4_males
Stage_5a_females
Stage_5a_males
Stage_5b_females
Stage_5b_males
Stage_5c_males
Stage_6_females
Stage_6_males
sex
F
M

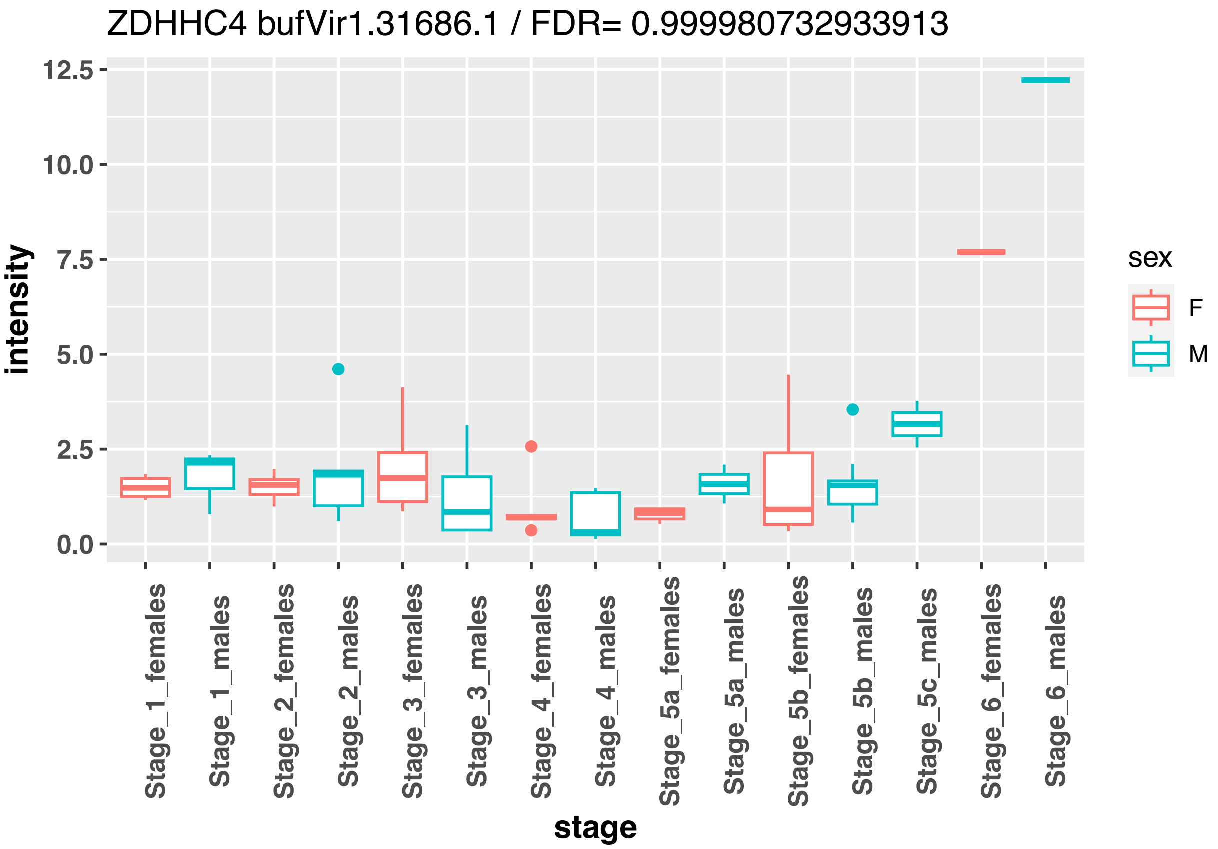
ZDHHC4 bufVir1.31686.1 / FDR= 0.999980732933913
intensity
12.5
10.0
7.5
5.0
2.5
0.0
Stage_1_females
Stage_1_males
Stage_2_females
Stage_2_males
Stage_3_females
Stage_3_males
Stage_4_females
Stage_4_males
Stage_5a_females
Stage_5a_males
Stage_5b_females
Stage_5b_males
Stage_5c_males
Stage_6_females
Stage_6_males
stage
sex
F
M

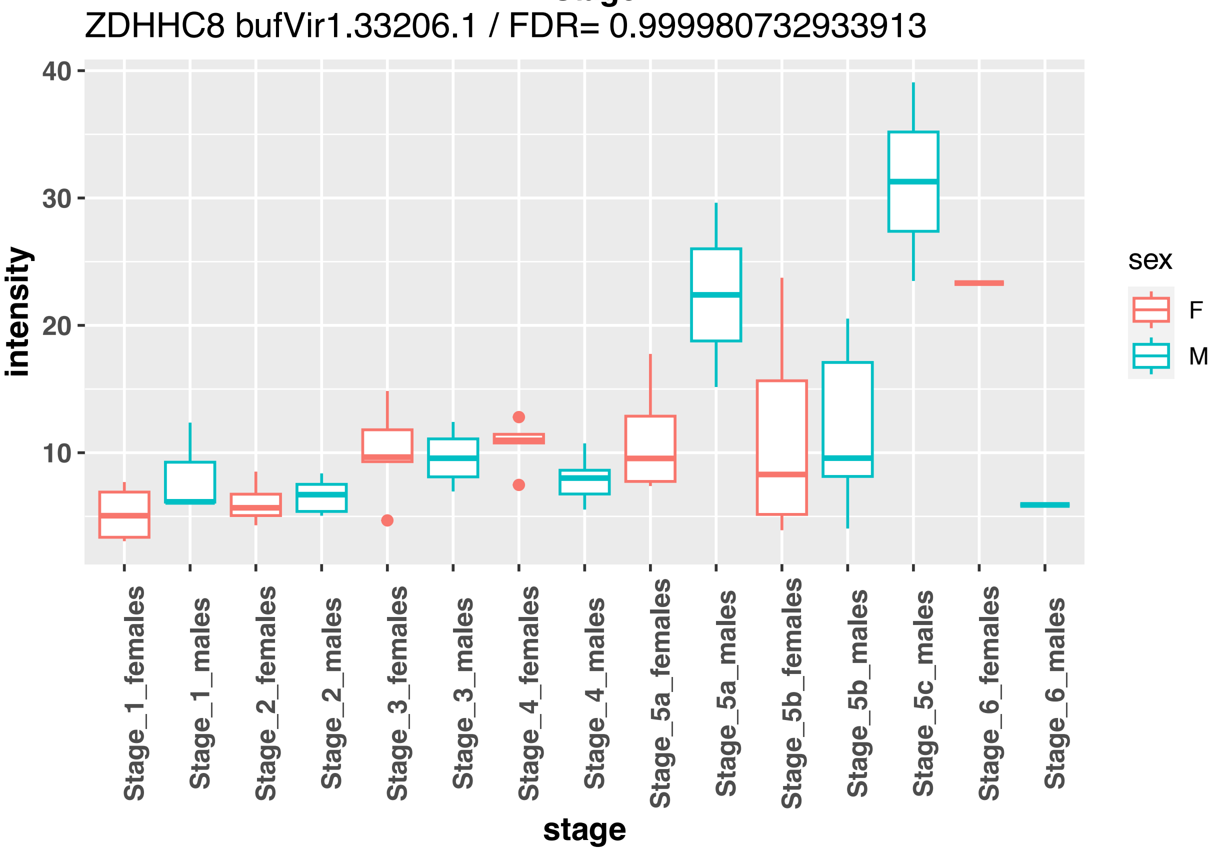
ZDHHC8 bufVir1.33206.1 / FDR= 0.999980732933913
intensity
40
30
20
10
Stage_1_females
Stage_1_males
Stage_2_females
Stage_2_males
Stage_3_females
Stage_3_males
Stage_4_females
Stage_4_males
Stage_5a_females
Stage_5a_males
Stage_5b_females
Stage_5b_males
Stage_5c_males
Stage_6_females
Stage_6_males
stage
sex
F
M

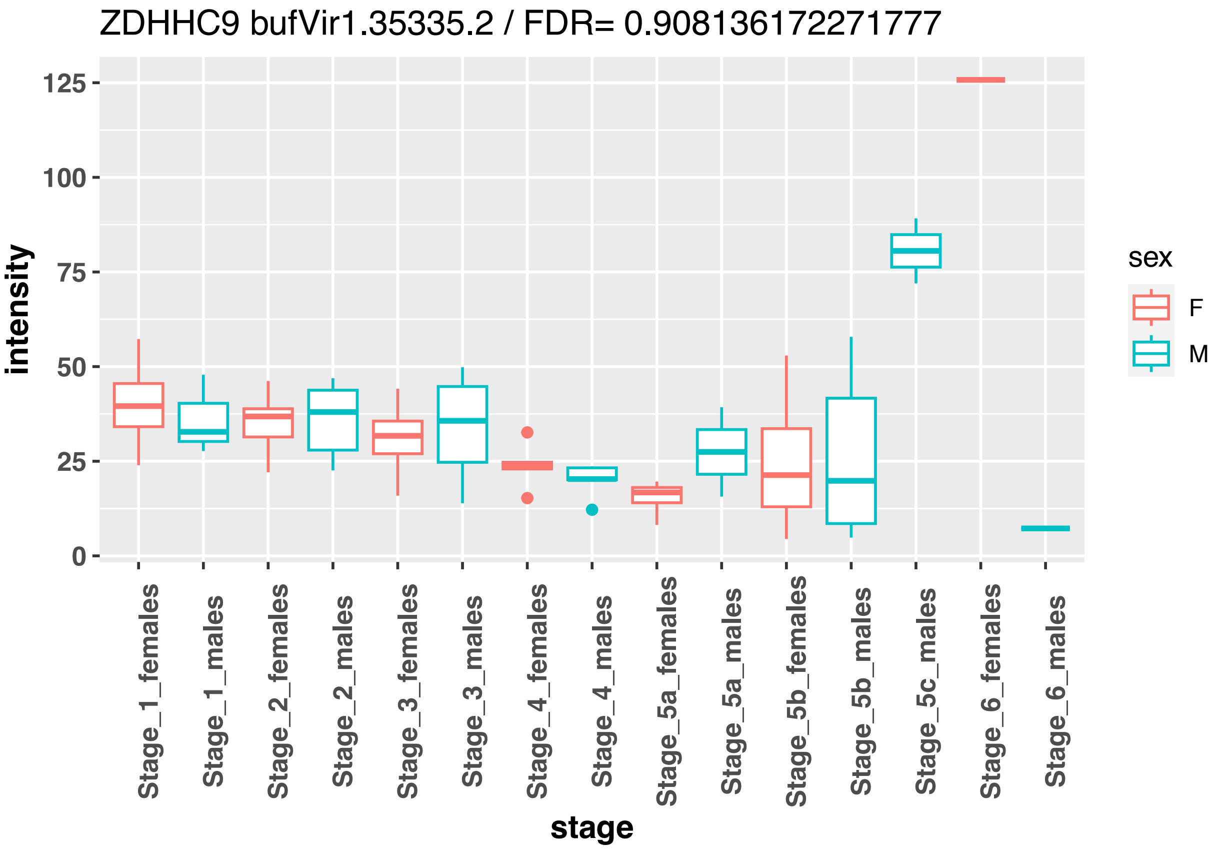
ZDHHC9 bufVir1.35335.2 / FDR= 0.908136172271777
intensity
125
100
75
50
25
0
stage
Stage_1_females
Stage_1_males
Stage_2_females
Stage_2_males
Stage_3_females
Stage_3_males
Stage_4_females
Stage_4_males
Stage_5a_females
Stage_5a_males
Stage_5b_females
Stage_5b_males
Stage_5c_males
Stage_6_females
Stage_6_males
sex
F
M

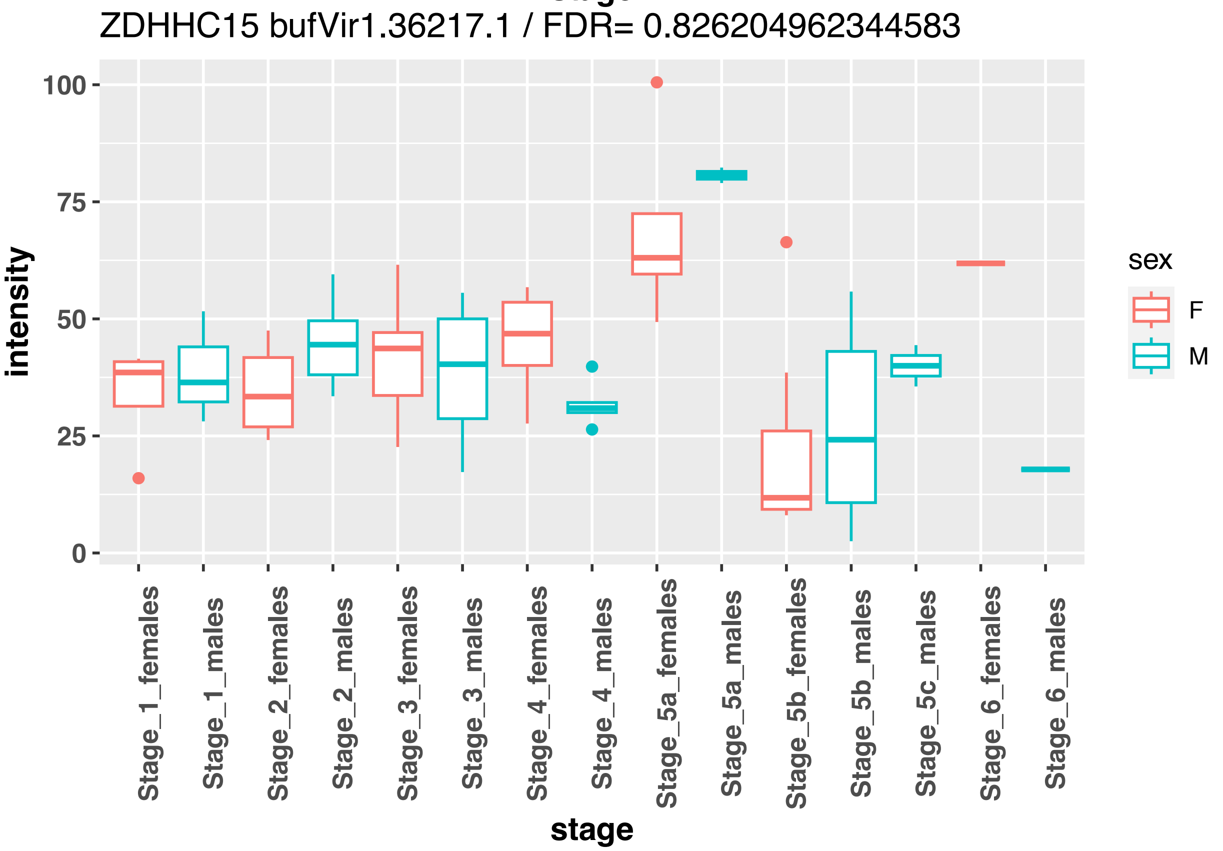
ZDHHC15 bufVir1.36217.1 / FDR= 0.826204962344583
intensity
100
75
50
25
0
stage
Stage_1_females
Stage_1_males
Stage_2_females
Stage_2_males
Stage_3_females
Stage_3_males
Stage_4_females
Stage_4_males
Stage_5a_females
Stage_5a_males
Stage_5b_females
Stage_5b_males
Stage_5c_males
Stage_6_females
Stage_6_males
sex
F
M

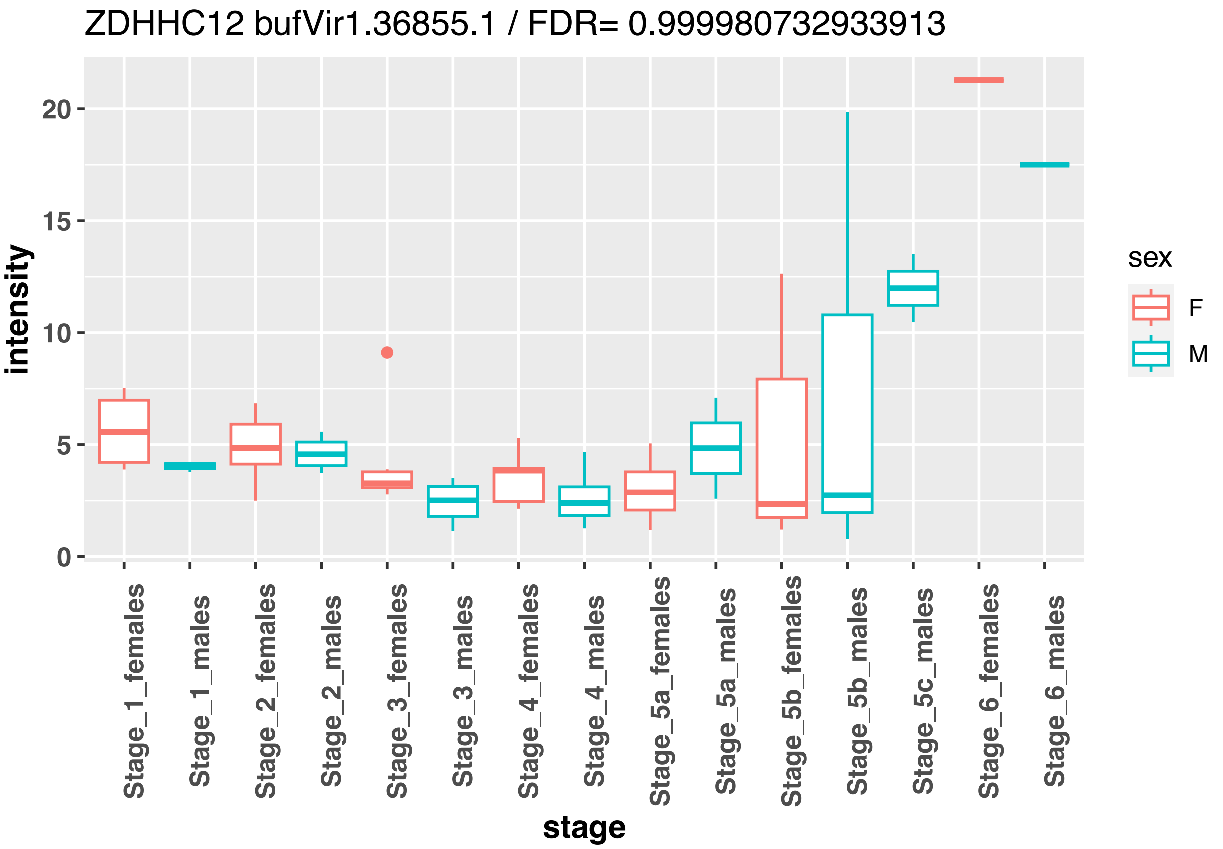
ZDHHC12 bufVir1.36855.1 / FDR= 0.999980732933913
intensity
20
15
10
5
0
Stage_1_females
Stage_1_males
Stage_2_females
Stage_2_males
Stage_3_females
Stage_3_males
Stage_4_females
Stage_4_males
Stage_5a_females
Stage_5a_males
Stage_5b_females
Stage_5b_males
Stage_5c_males
Stage_6_females
Stage_6_males
stage
sex
F
M

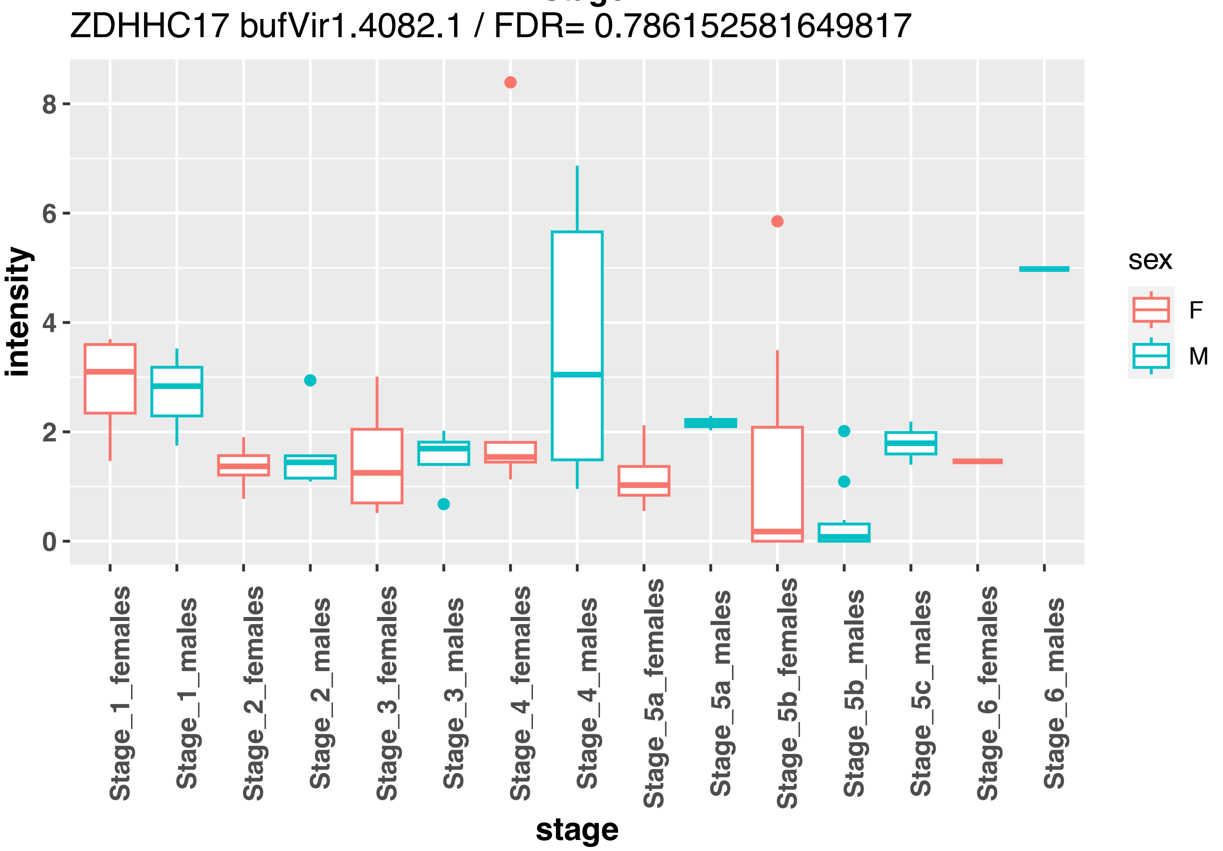
ZDHHC17 bufVir1.4082.1 / FDR= 0.786152581649817
intensity
8
6
4
2
0
Stage_1_females
Stage_1_males
Stage_2_females
Stage_2_males
Stage_3_females
Stage_3_males
Stage_4_females
Stage_4_males
Stage_5a_females
Stage_5a_males
Stage_5b_females
Stage_5b_males
Stage_5c_males
Stage_6_females
Stage_6_males
stage
sex
F
M

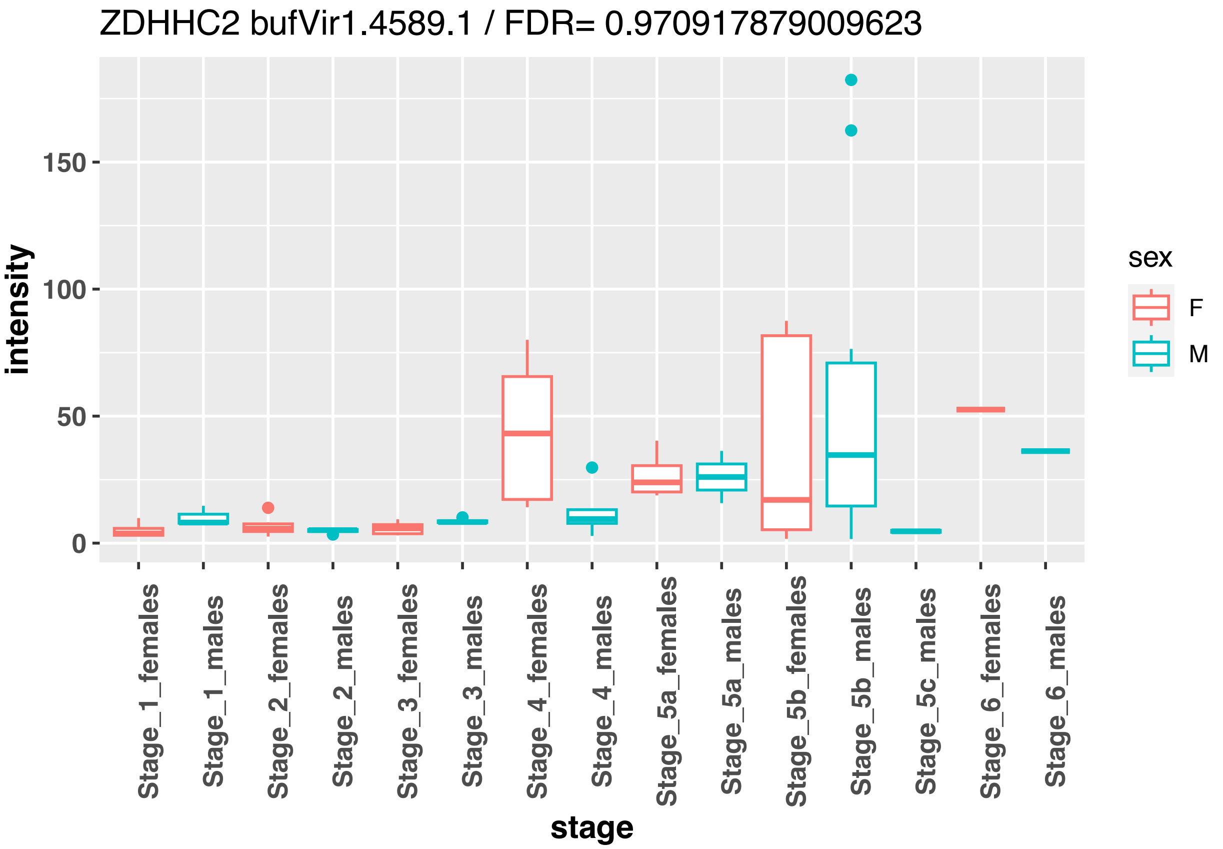
ZDHHC2 bufVir1.4589.1 / FDR= 0.970917879009623
intensity
150
100
50
0
Stage_1_females
Stage_1_males
Stage_2_females
Stage_2_males
Stage_3_females
Stage_3_males
Stage_4_females
Stage_4_males
Stage_5a_females
Stage_5a_males
Stage_5b_females
Stage_5b_males
Stage_5c_males
Stage_6_females
Stage_6_males
stage
sex
F
M

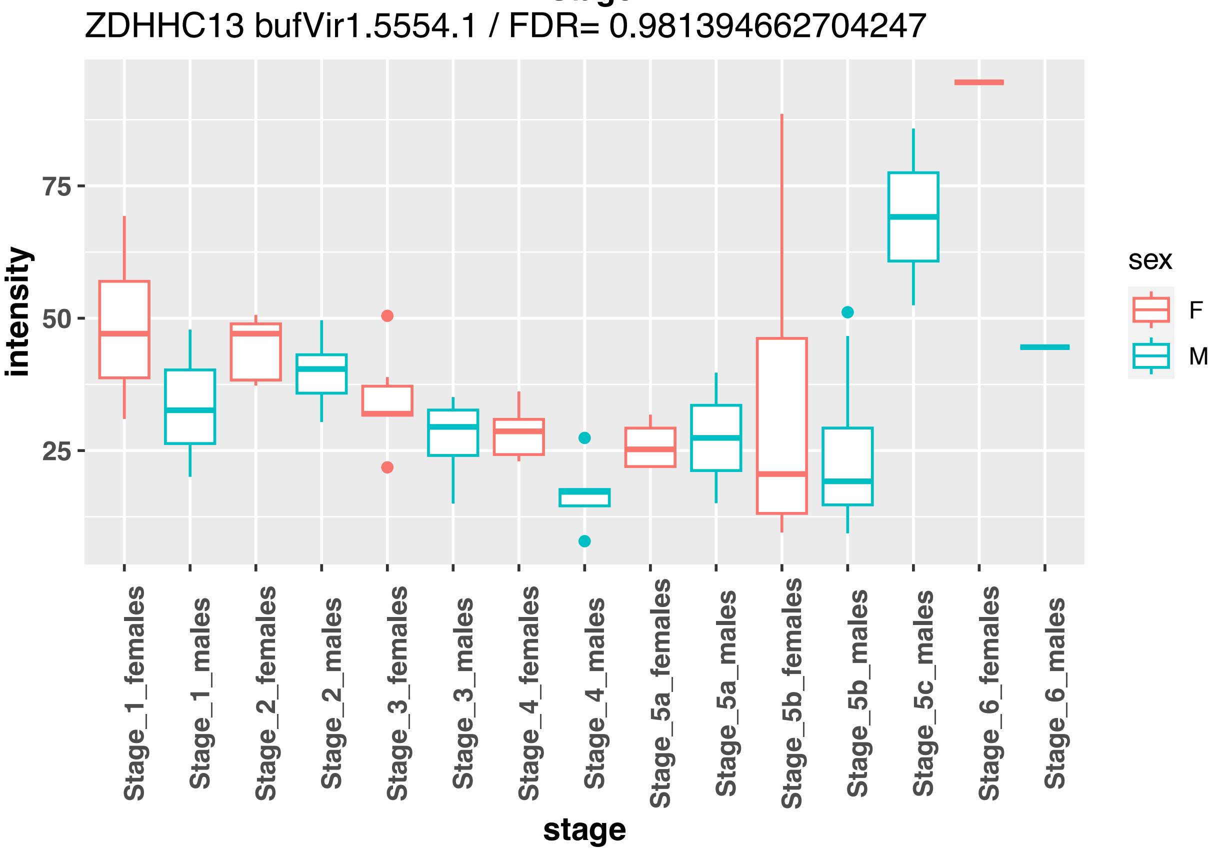
ZDHHC13 bufVir1.5554.1 / FDR= 0.981394662704247
intensity
75
50
25
Stage_1_females
Stage_1_males
Stage_2_females
Stage_2_males
Stage_3_females
Stage_3_males
Stage_4_females
Stage_4_males
Stage_5a_females
Stage_5a_males
Stage_5b_females
Stage_5b_males
Stage_5c_males
Stage_6_females
Stage_6_males
stage
sex
F
M

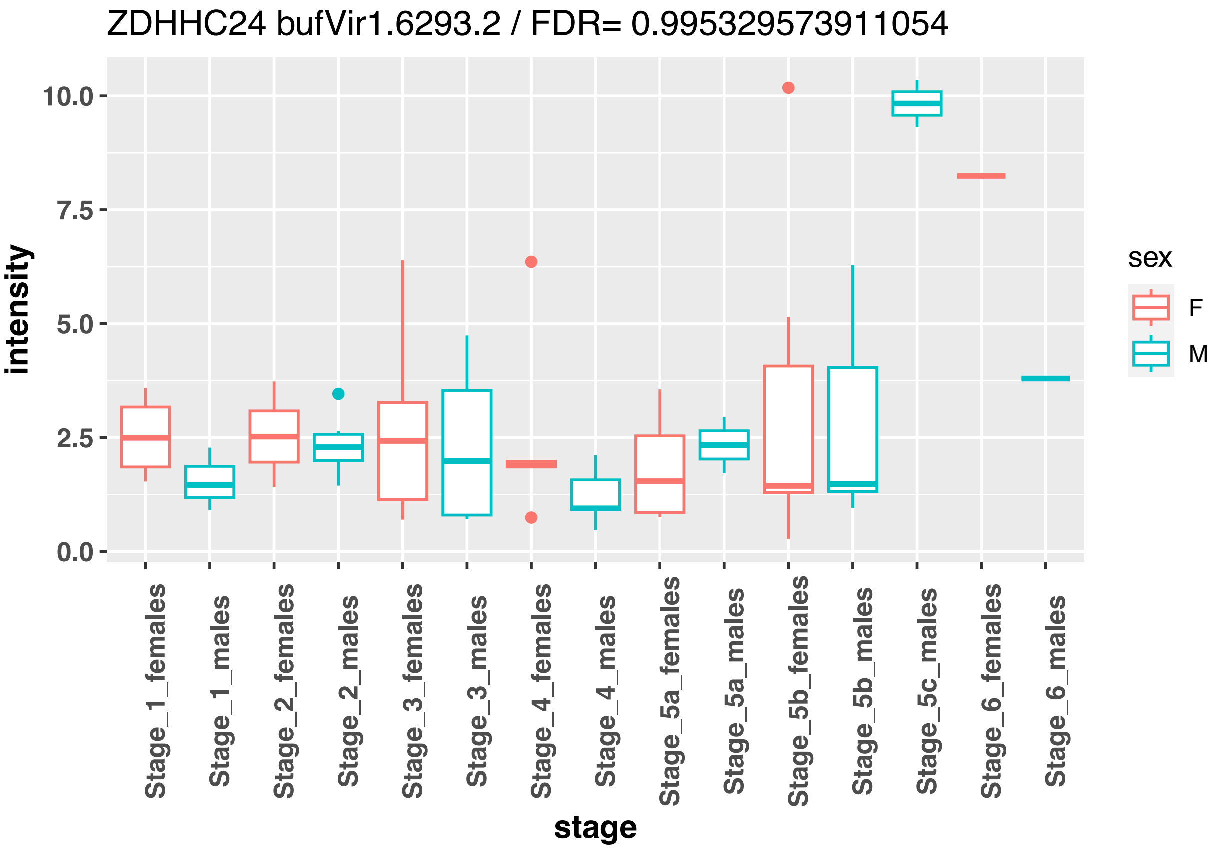
ZDHHC24 bufVir1.6293.2 / FDR= 0.995329573911054
intensity
10.0
7.5
5.0
2.5
0.0
Stage_1_females
Stage_1_males
Stage_2_females
Stage_2_males
Stage_3_females
Stage_3_males
Stage_4_females
Stage_4_males
Stage_5a_females
Stage_5a_males
Stage_5b_females
Stage_5b_males
Stage_5c_males
Stage_6_females
Stage_6_males
stage
sex
F
M

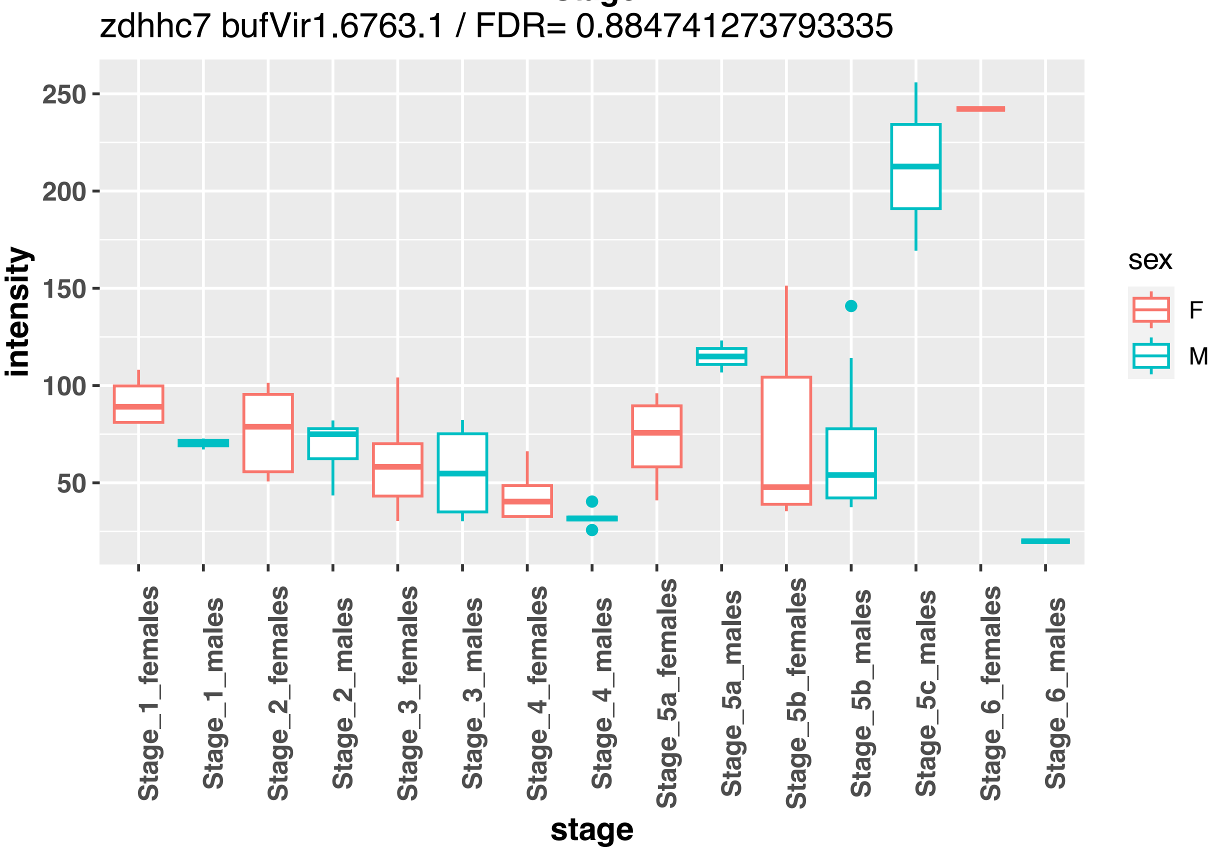
zdhhc7 bufVir1.6763.1 / FDR= 0.884741273793335
intensity
250
200
150
100
50
Stage_1_females
Stage_1_males
Stage_2_females
Stage_2_males
Stage_3_females
Stage_3_males
Stage_4_females
Stage_4_males
Stage_5a_females
Stage_5a_males
Stage_5b_females
Stage_5b_males
Stage_5c_males
Stage_6_females
Stage_6_males
stage
sex
F
M

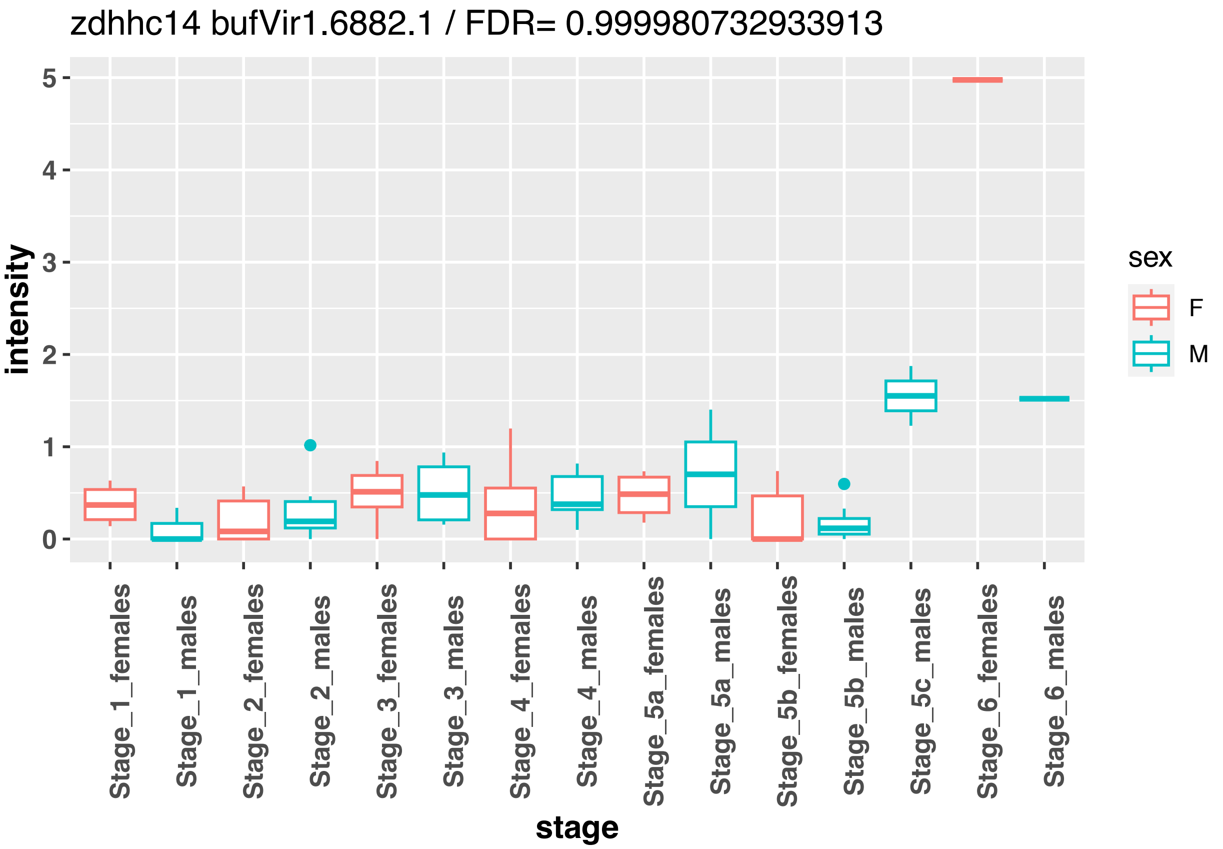
zdhhc14 bufVir1.6882.1 / FDR= 0.999980732933913
intensity
0
1
2
3
4
5
sex
F
M
Stage_1_females
Stage_1_males
Stage_2_females
Stage_2_males
Stage_3_females
Stage_3_males
Stage_4_females
Stage_4_males
Stage_5a_females
Stage_5a_males
Stage_5b_females
Stage_5b_males
Stage_5c_males
Stage_6_females
Stage_6_males
stage

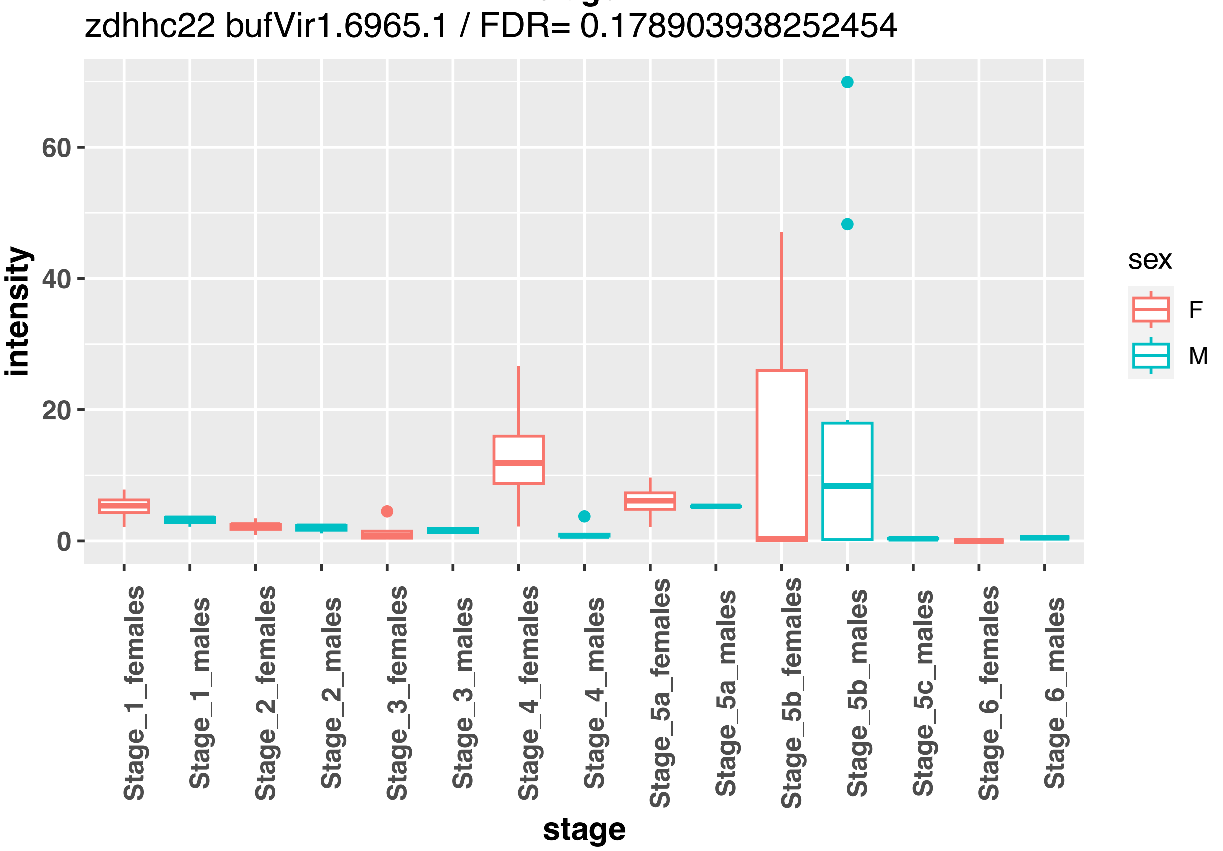
zdhhc22 bufVir1.6965.1 / FDR= 0.178903938252454
intensity
0
20
40
60
sex
F
M
Stage_1_females
Stage_1_males
Stage_2_females
Stage_2_males
Stage_3_females
Stage_3_males
Stage_4_females
Stage_4_males
Stage_5a_females
Stage_5a_males
Stage_5b_females
Stage_5b_males
Stage_5c_males
Stage_6_females
Stage_6_males
stage

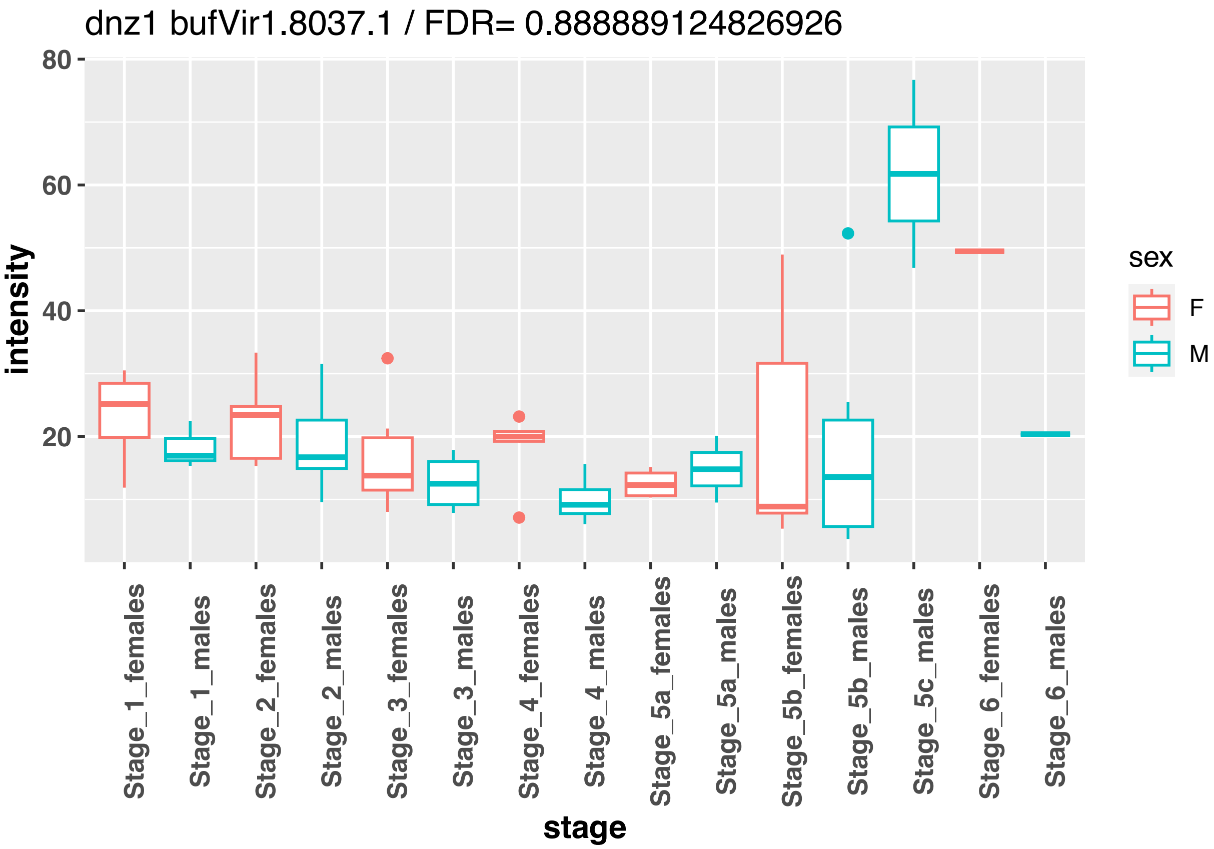
dnz1 bufVir1.8037.1 / FDR= 0.888889124826926
intensity
80
60
40
20
sex
F
M
Stage_1_females
Stage_1_males
Stage_2_females
Stage_2_males
Stage_3_females
Stage_3_males
Stage_4_females
Stage_4_males
Stage_5a_females
Stage_5a_males
Stage_5b_females
Stage_5b_males
Stage_5c_males
Stage_6_females
Stage_6_males
stage

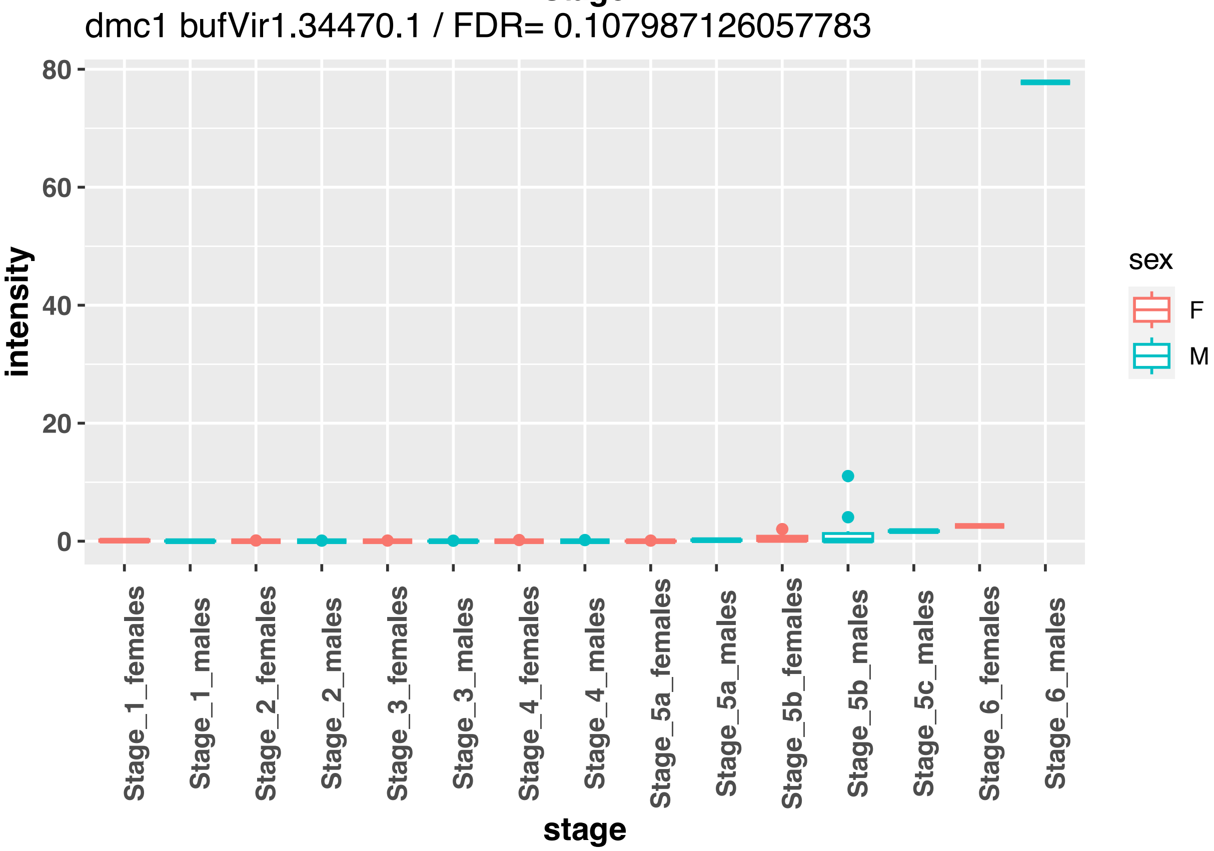
dmc1 bufVir1.34470.1 / FDR= 0.107987126057783
intensity
80
60
40
20
0
sex
F
M
Stage_1_females
Stage_1_males
Stage_2_females
Stage_2_males
Stage_3_females
Stage_3_males
Stage_4_females
Stage_4_males
Stage_5a_females
Stage_5a_males
Stage_5b_females
Stage_5b_males
Stage_5c_males
Stage_6_females
Stage_6_males
stage

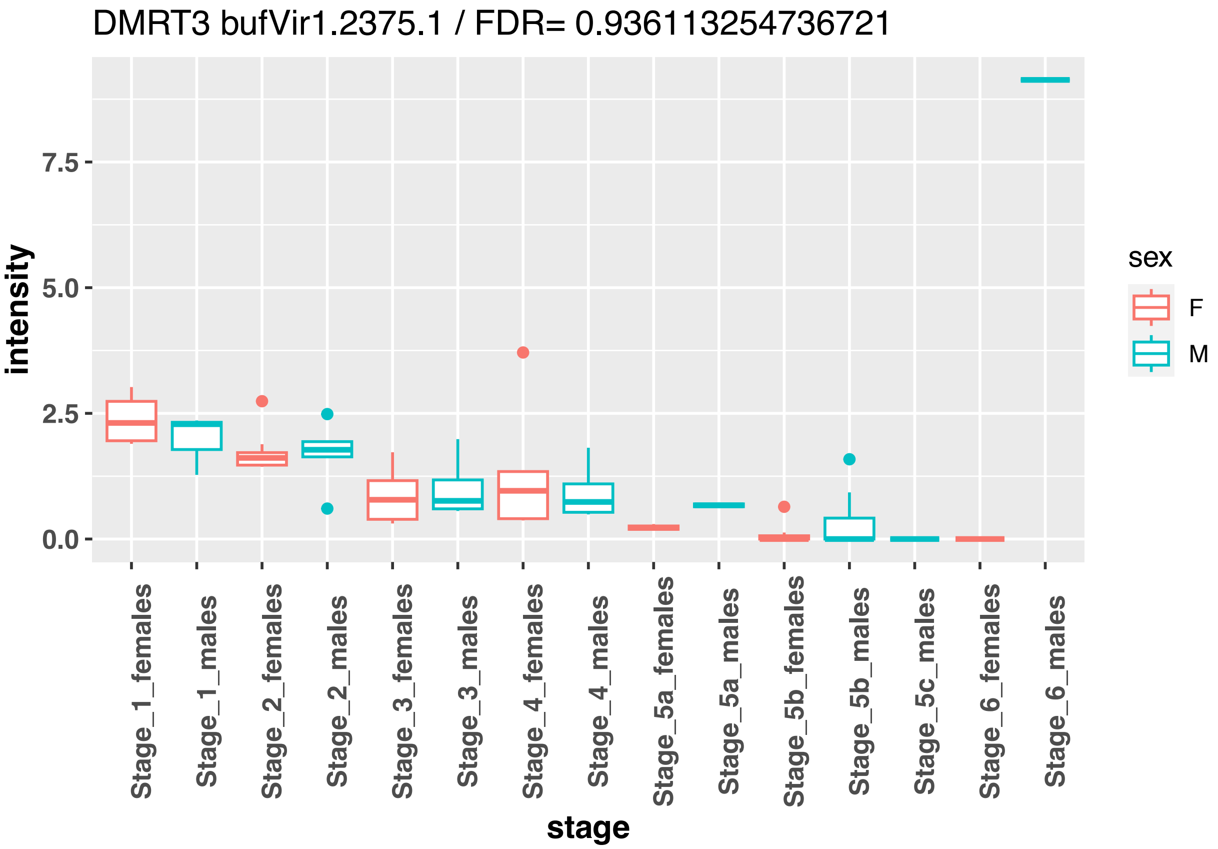
DMRT3 bufVir1.2375.1 / FDR= 0.936113254736721
intensity
7.5
5.0
2.5
0.0
Stage_1_females
Stage_1_males
Stage_2_females
Stage_2_males
Stage_3_females
Stage_3_males
Stage_4_females
Stage_4_males
Stage_5a_females
Stage_5a_males
Stage_5b_females
Stage_5b_males
Stage_5c_males
Stage_6_females
Stage_6_males
stage
sex
F
M

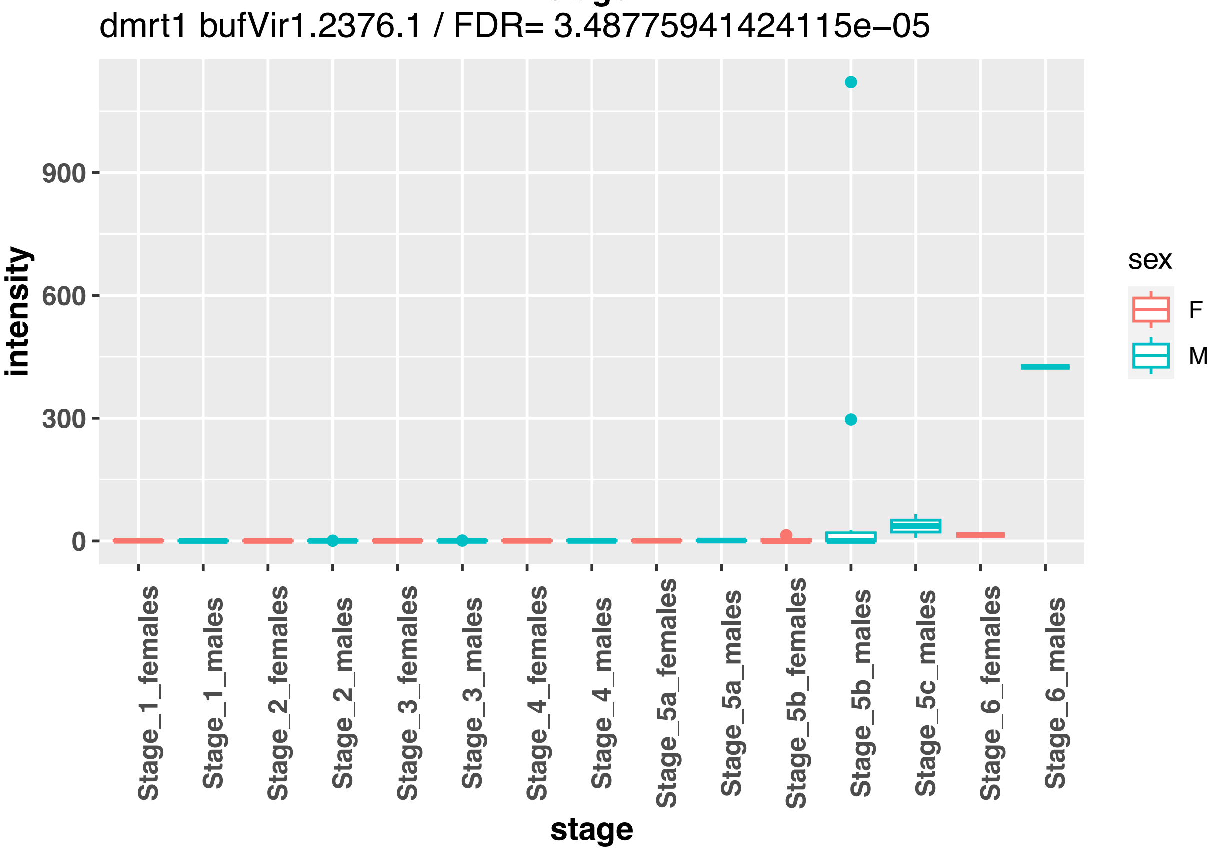
dmrt1 bufVir1.2376.1 / FDR= 3.48775941424115e−05
intensity
900
600
300
0
Stage_1_females
Stage_1_males
Stage_2_females
Stage_2_males
Stage_3_females
Stage_3_males
Stage_4_females
Stage_4_males
Stage_5a_females
Stage_5a_males
Stage_5b_females
Stage_5b_males
Stage_5c_males
Stage_6_females
Stage_6_males
stage
sex
F
M

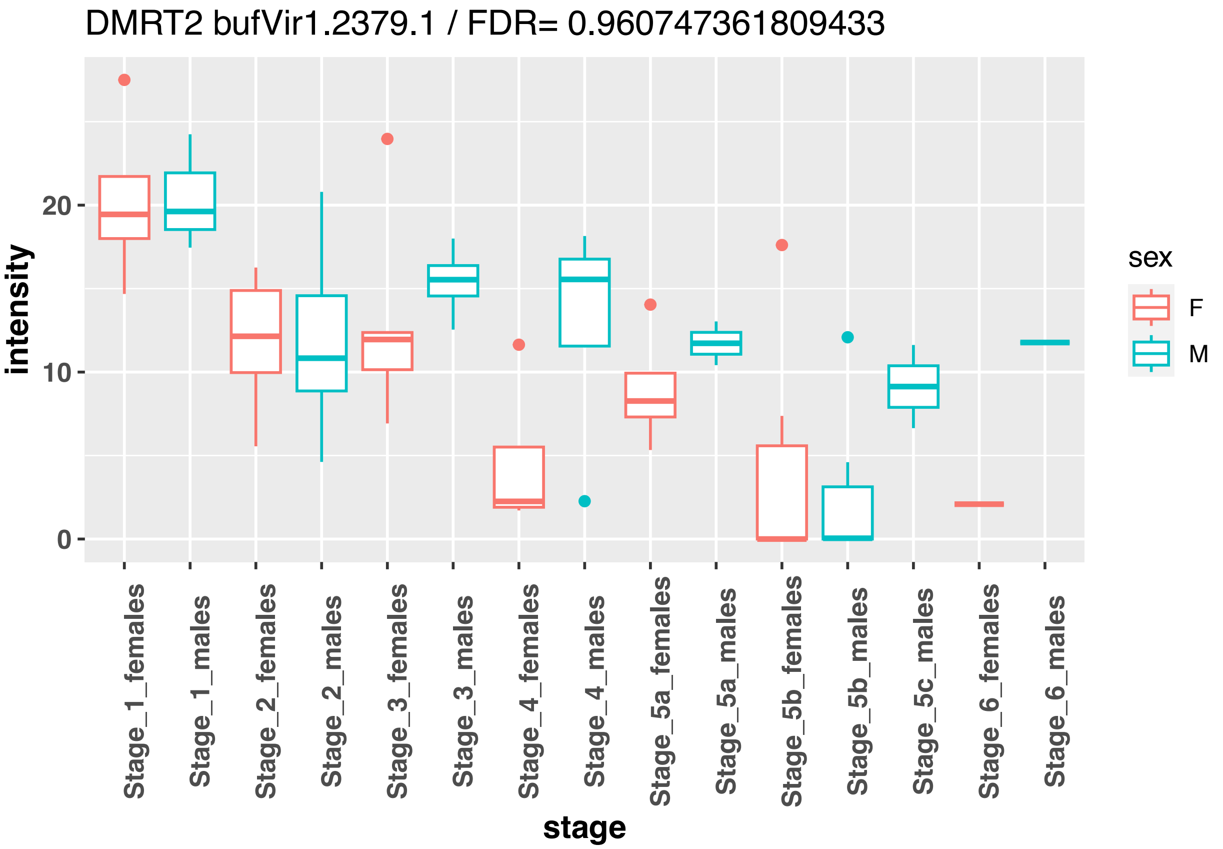
DMRT2 bufVir1.2379.1 / FDR= 0.960747361809433
intensity
20
10
0
stage
Stage_1_females
Stage_1_males
Stage_2_females
Stage_2_males
Stage_3_females
Stage_3_males
Stage_4_females
Stage_4_males
Stage_5a_females
Stage_5a_males
Stage_5b_females
Stage_5b_males
Stage_5c_males
Stage_6_females
Stage_6_males
sex
F
M

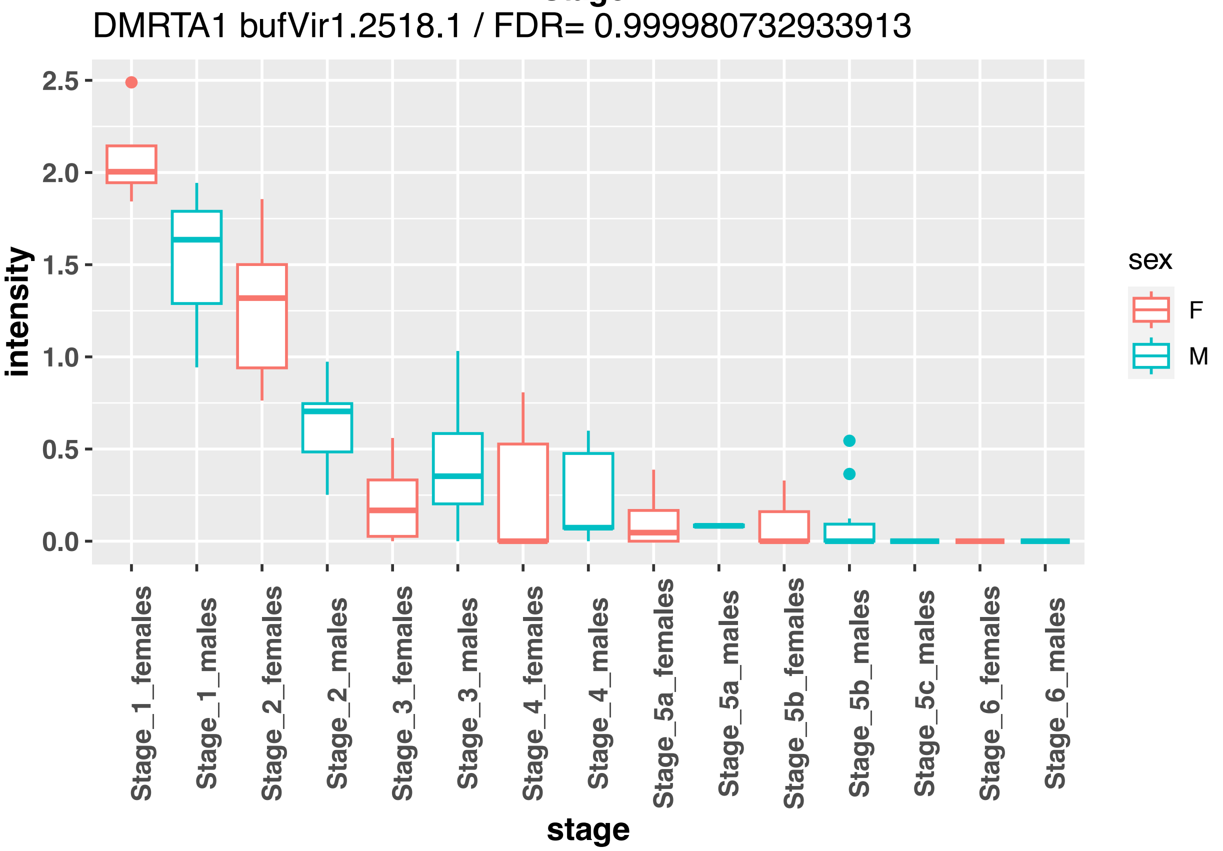
DMRTA1 bufVir1.2518.1 / FDR= 0.999980732933913
intensity
2.5
2.0
1.5
1.0
0.5
0.0
stage
Stage_1_females
Stage_1_males
Stage_2_females
Stage_2_males
Stage_3_females
Stage_3_males
Stage_4_females
Stage_4_males
Stage_5a_females
Stage_5a_males
Stage_5b_females
Stage_5b_males
Stage_5c_males
Stage_6_females
Stage_6_males
sex
F
M

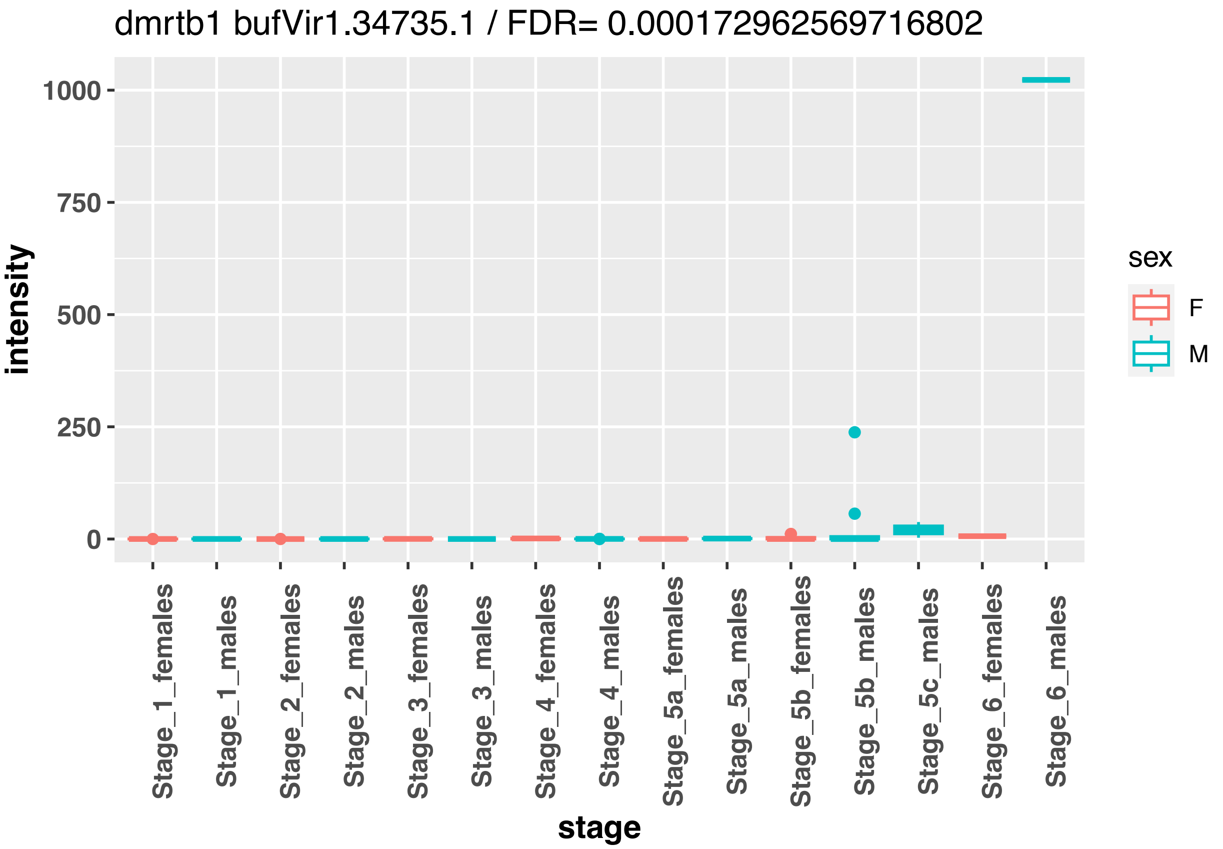
dmrtb1 bufVir1.34735.1 / FDR= 0.000172962569716802
intensity
1000
750
500
250
0
stage
Stage_1_females
Stage_1_males
Stage_2_females
Stage_2_males
Stage_3_females
Stage_3_males
Stage_4_females
Stage_4_males
Stage_5a_females
Stage_5a_males
Stage_5b_females
Stage_5b_males
Stage_5c_males
Stage_6_females
Stage_6_males
sex
F
M

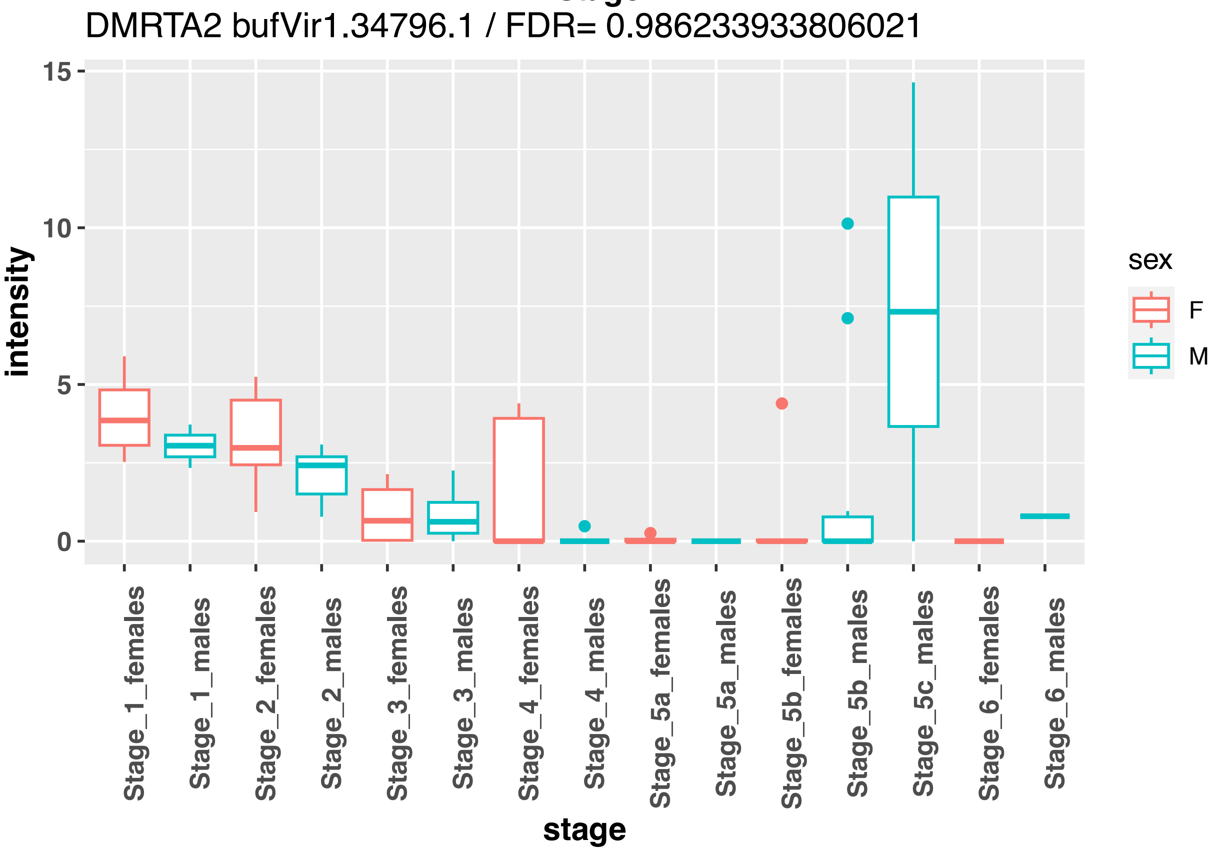
DMRTA2 bufVir1.34796.1 / FDR= 0.986233933806021
intensity
15
10
5
0
stage
Stage_1_females
Stage_1_males
Stage_2_females
Stage_2_males
Stage_3_females
Stage_3_males
Stage_4_females
Stage_4_males
Stage_5a_females
Stage_5a_males
Stage_5b_females
Stage_5b_males
Stage_5c_males
Stage_6_females
Stage_6_males
sex
F
M

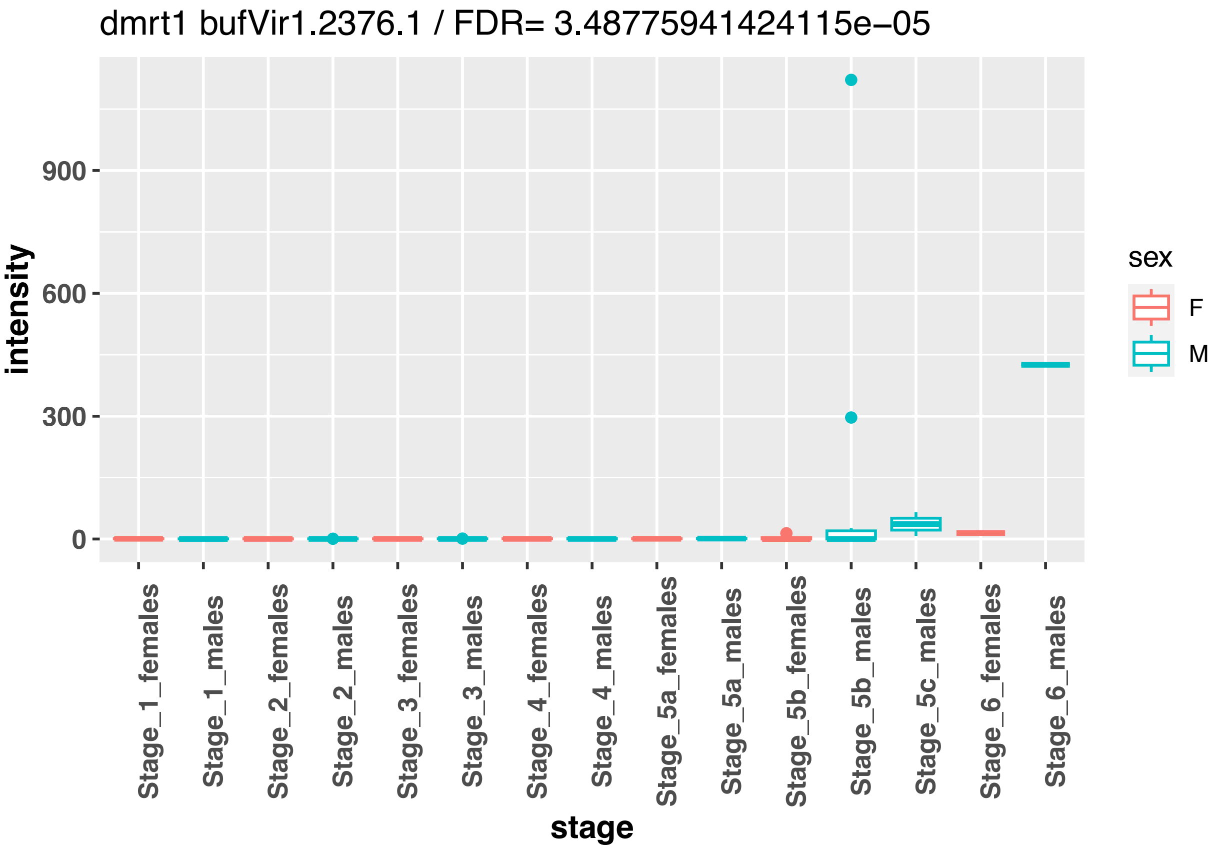
dmrt1 bufVir1.2376.1 / FDR= 3.48775941424115e−05
intensity
900
600
300
0
stage
Stage_1_females
Stage_1_males
Stage_2_females
Stage_2_males
Stage_3_females
Stage_3_males
Stage_4_females
Stage_4_males
Stage_5a_females
Stage_5a_males
Stage_5b_females
Stage_5b_males
Stage_5c_males
Stage_6_females
Stage_6_males
sex
F
M

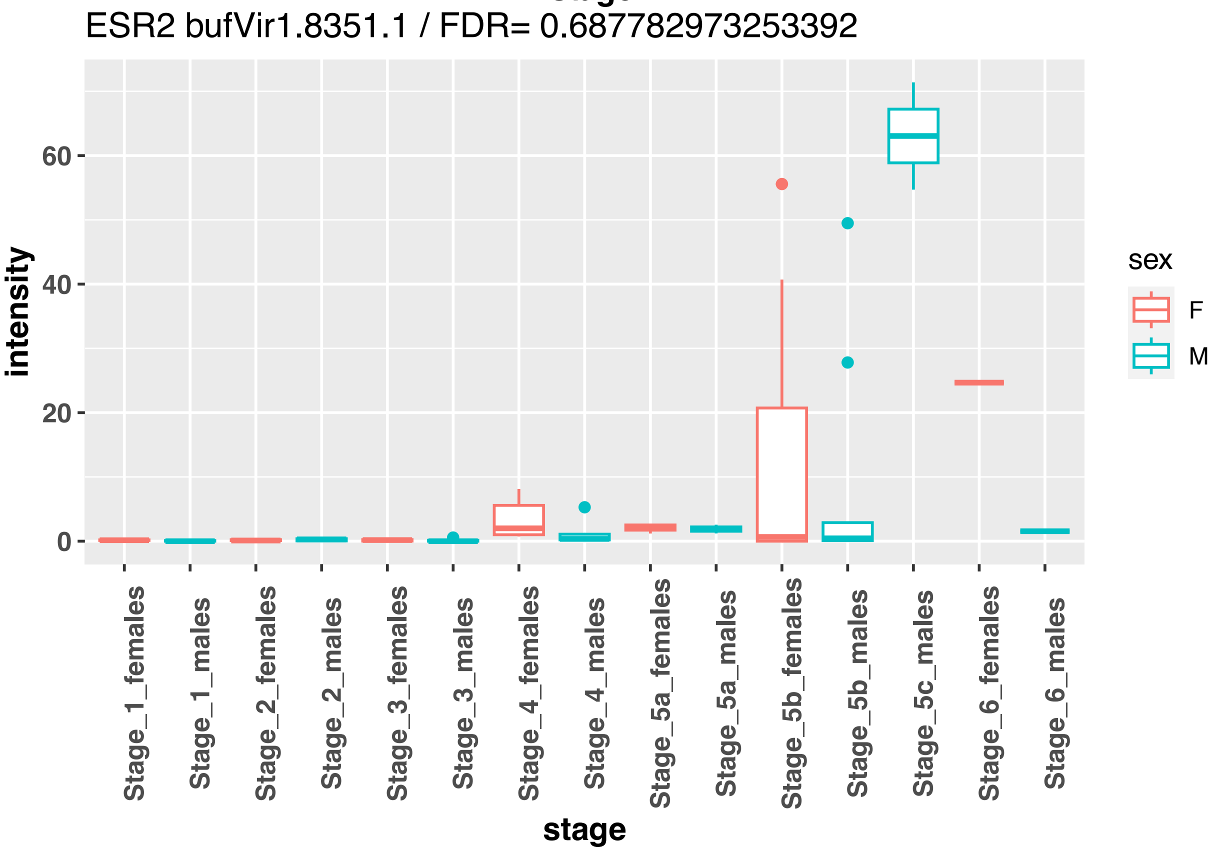
ESR2 bufVir1.8351.1 / FDR= 0.687782973253392
intensity
60
40
20
0
stage
Stage_1_females
Stage_1_males
Stage_2_females
Stage_2_males
Stage_3_females
Stage_3_males
Stage_4_females
Stage_4_males
Stage_5a_females
Stage_5a_males
Stage_5b_females
Stage_5b_males
Stage_5c_males
Stage_6_females
Stage_6_males
sex
F
M

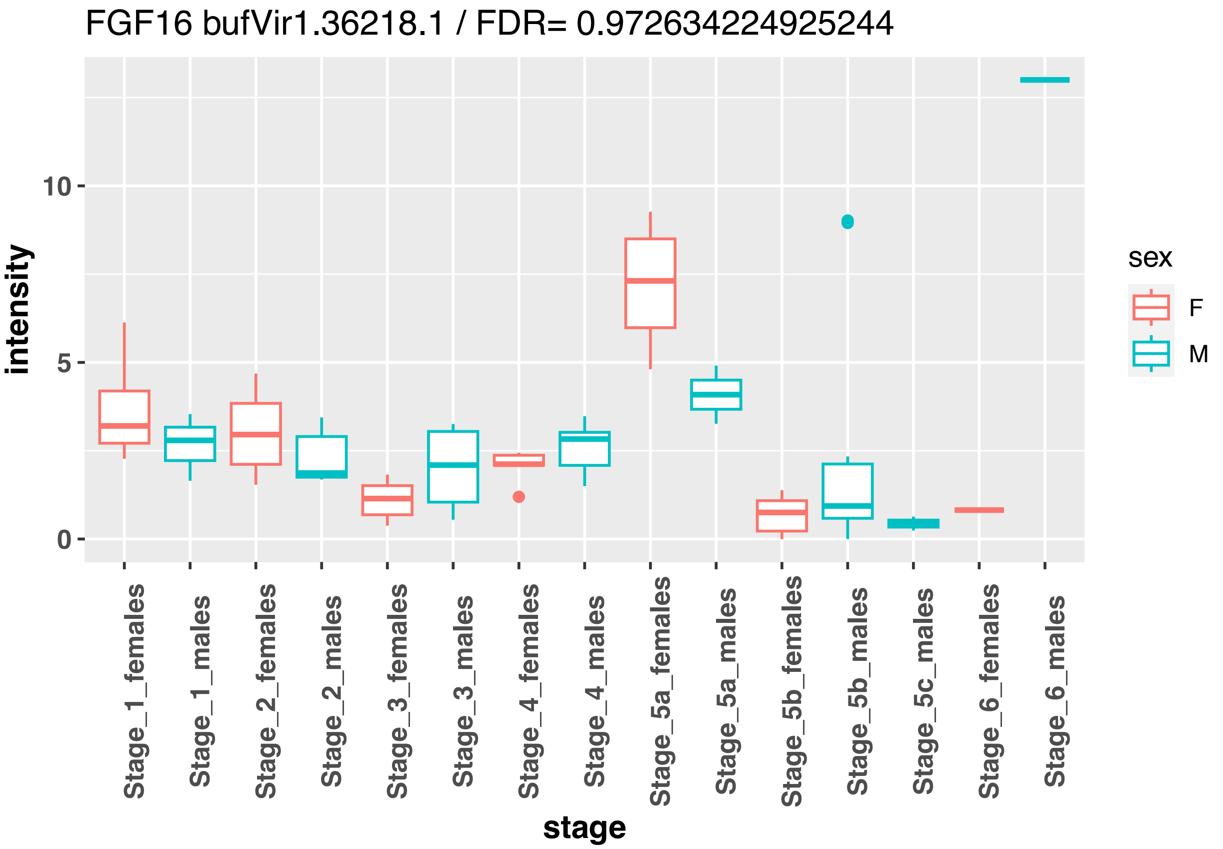
FGF16 bufVir1.36218.1 / FDR= 0.972634224925244
intensity
stage
sex
F
M
0
5
10
Stage_1_females
Stage_1_males
Stage_2_females
Stage_2_males
Stage_3_females
Stage_3_males
Stage_4_females
Stage_4_males
Stage_5a_females
Stage_5a_males
Stage_5b_females
Stage_5b_males
Stage_5c_males
Stage_6_females
Stage_6_males

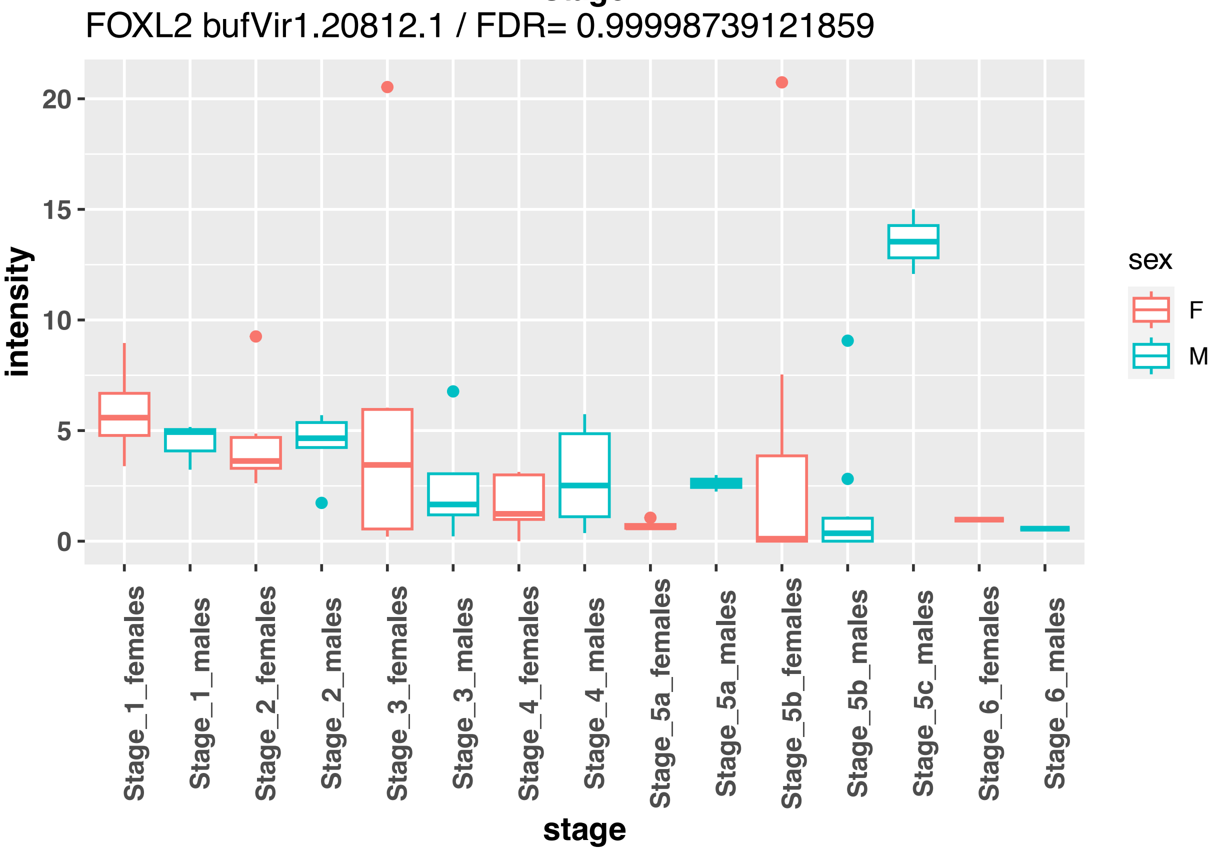
FOXL2 bufVir1.20812.1 / FDR= 0.99998739121859
intensity
stage
sex
F
M
0
5
10
15
20
Stage_1_females
Stage_1_males
Stage_2_females
Stage_2_males
Stage_3_females
Stage_3_males
Stage_4_females
Stage_4_males
Stage_5a_females
Stage_5a_males
Stage_5b_females
Stage_5b_males
Stage_5c_males
Stage_6_females
Stage_6_males

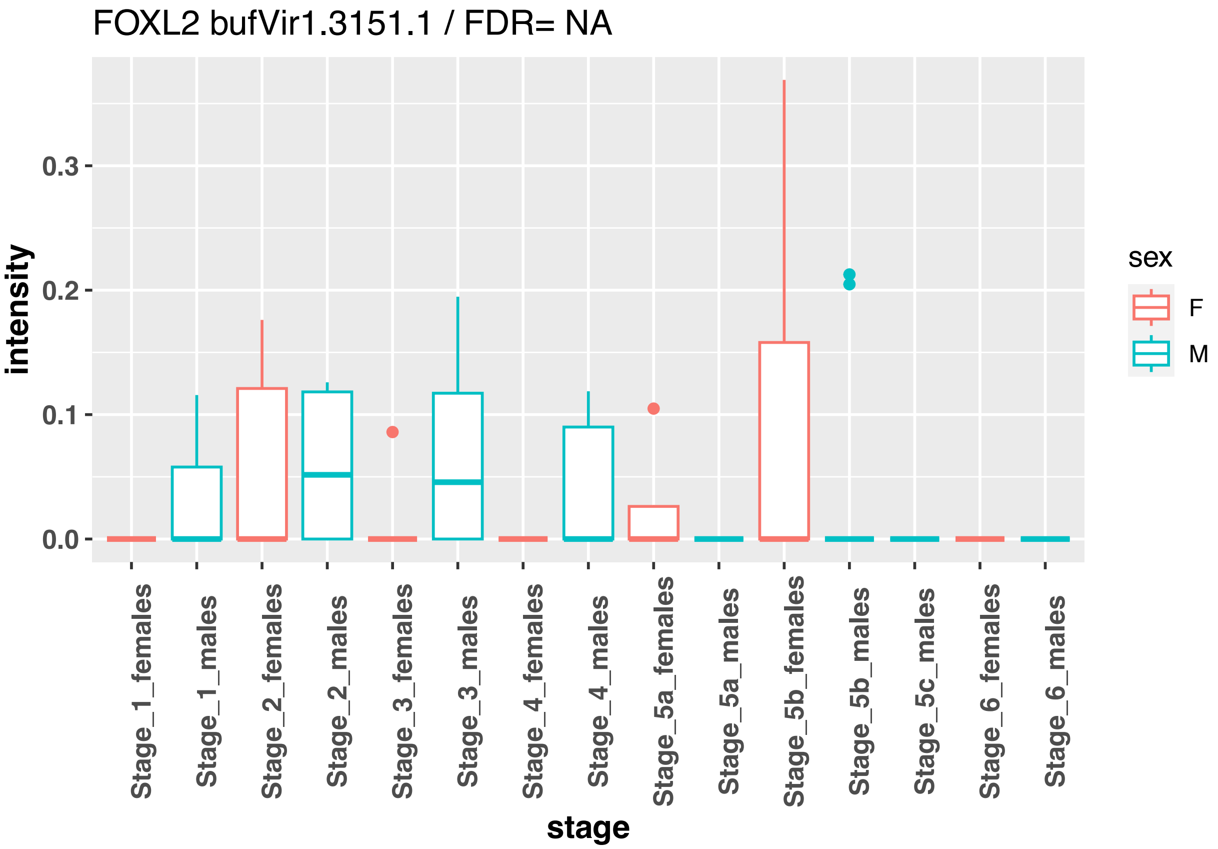
FOXL2 bufVir1.3151.1 / FDR= NA
intensity
0.3
0.2
0.1
0.0
sex
F
M
Stage_1_females
Stage_1_males
Stage_2_females
Stage_2_males
Stage_3_females
Stage_3_males
Stage_4_females
Stage_4_males
Stage_5a_females
Stage_5a_males
Stage_5b_females
Stage_5b_males
Stage_5c_males
Stage_6_females
Stage_6_males
stage

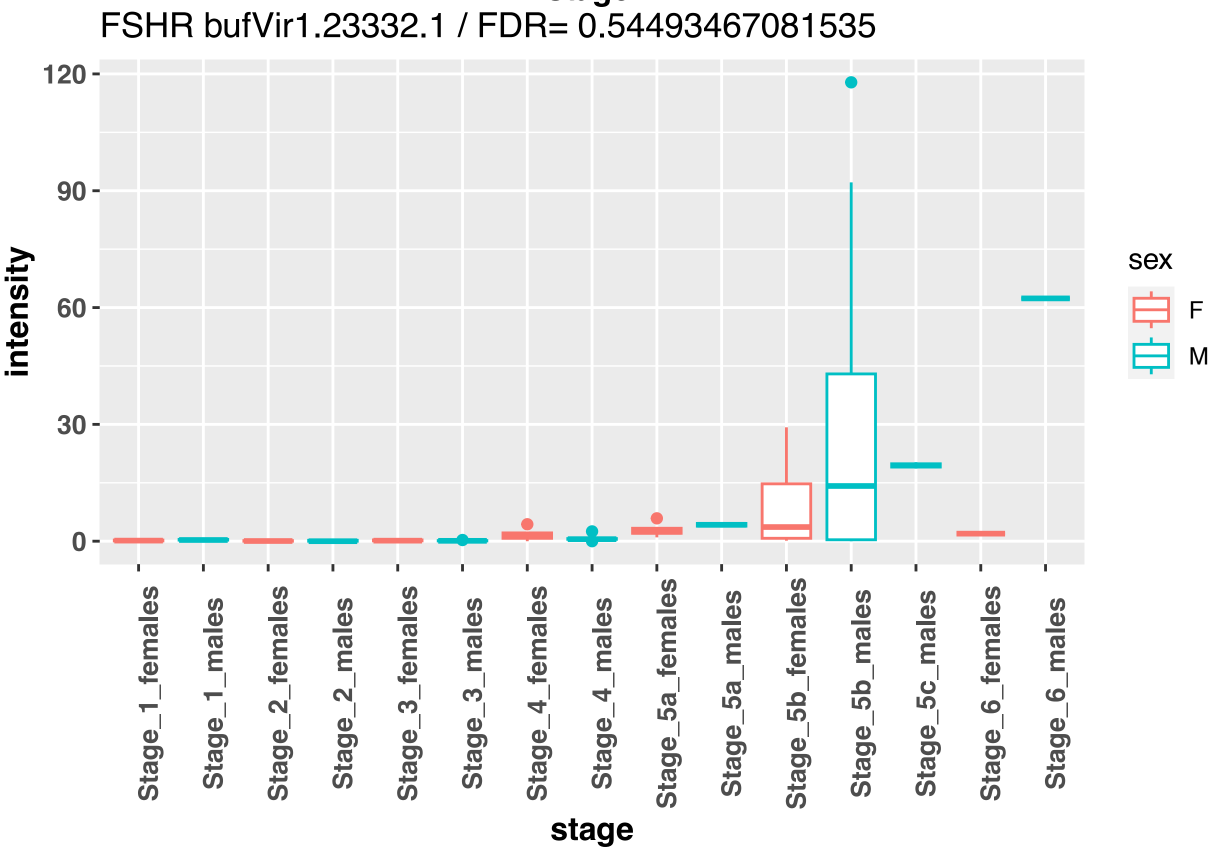
FSHR bufVir1.23332.1 / FDR= 0.54493467081535
intensity
120
90
60
30
0
sex
F
M
Stage_1_females
Stage_1_males
Stage_2_females
Stage_2_males
Stage_3_females
Stage_3_males
Stage_4_females
Stage_4_males
Stage_5a_females
Stage_5a_males
Stage_5b_females
Stage_5b_males
Stage_5c_males
Stage_6_females
Stage_6_males
stage

FZD7 bufVir1.32536.1 / FDR= 0.995329573911054

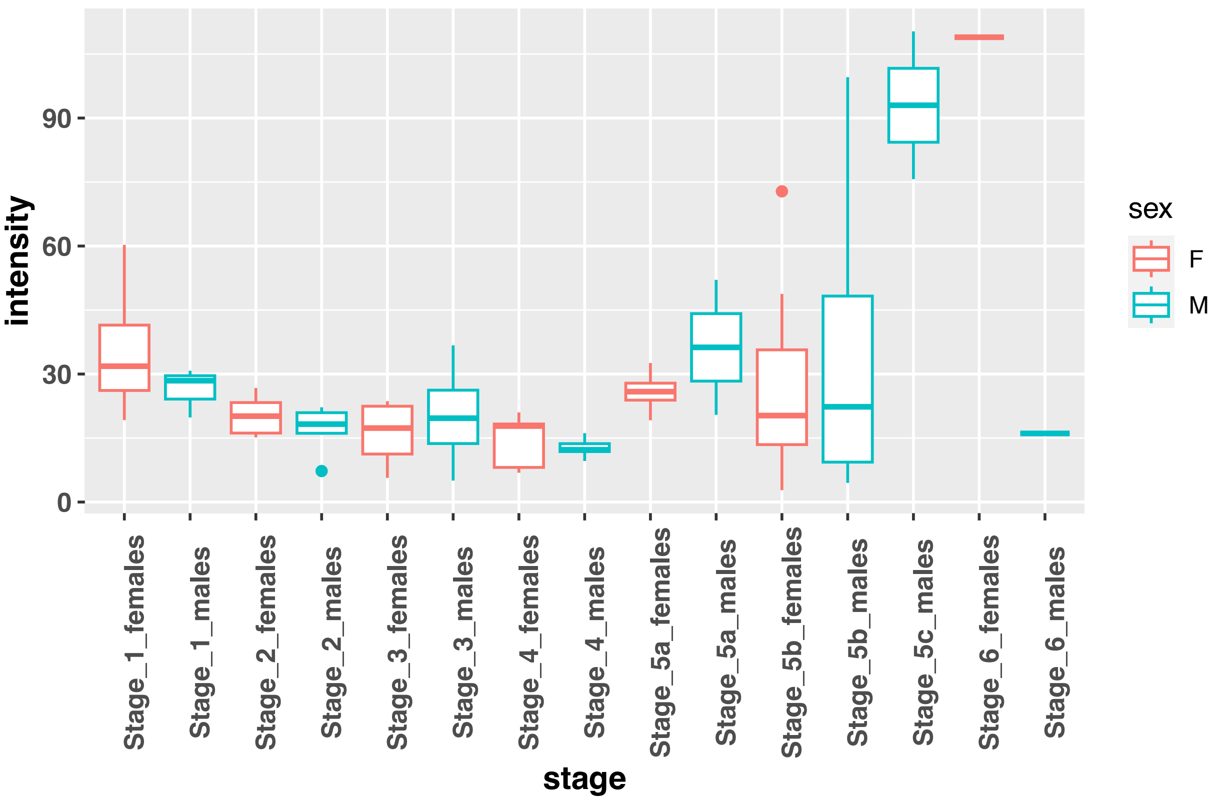


GATA4 bufVir1.20146.1 / FDR= 0.999980732933913

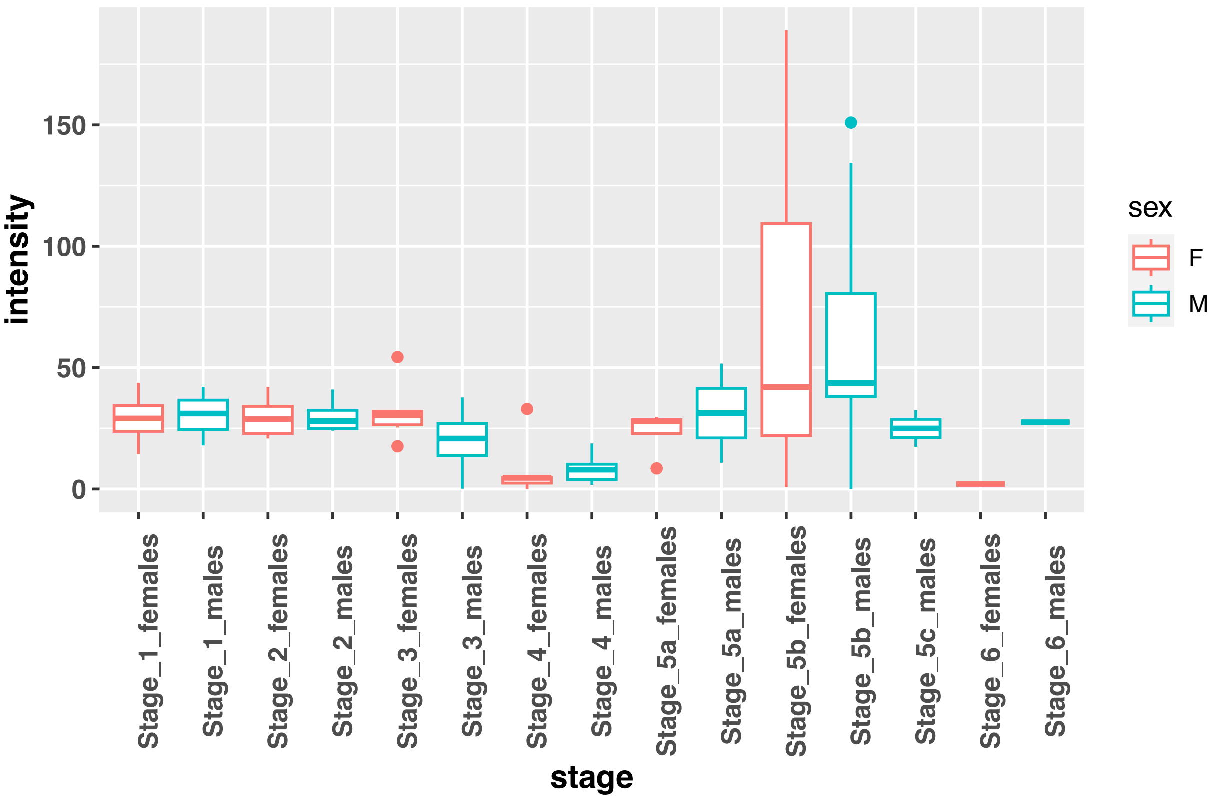

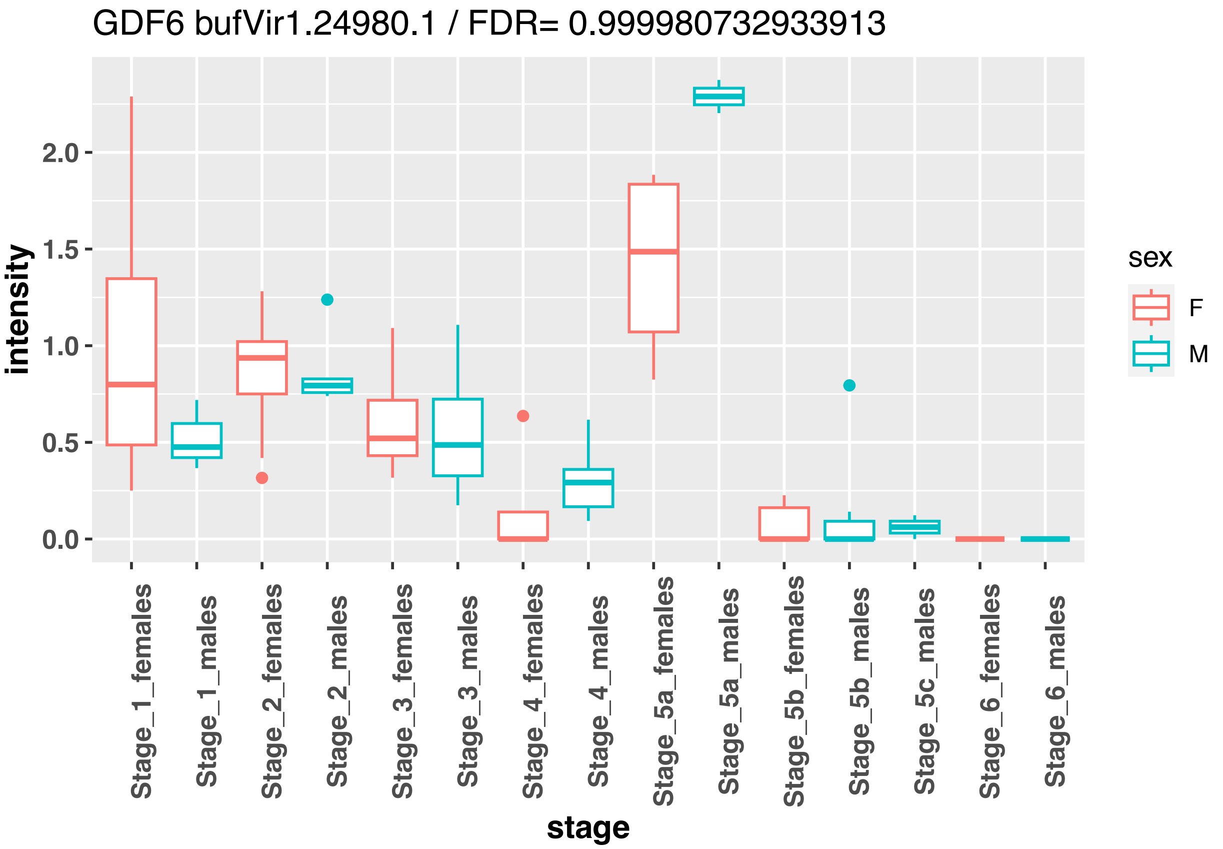
GDF6 bufVir1.24980.1 / FDR= 0.999980732933913
intensity
2.0
1.5
1.0
0.5
0.0
sex
F
M
Stage_1_females
Stage_1_males
Stage_2_females
Stage_2_males
Stage_3_females
Stage_3_males
Stage_4_females
Stage_4_males
Stage_5a_females
Stage_5a_males
Stage_5b_females
Stage_5b_males
Stage_5c_males
Stage_6_females
Stage_6_males
stage

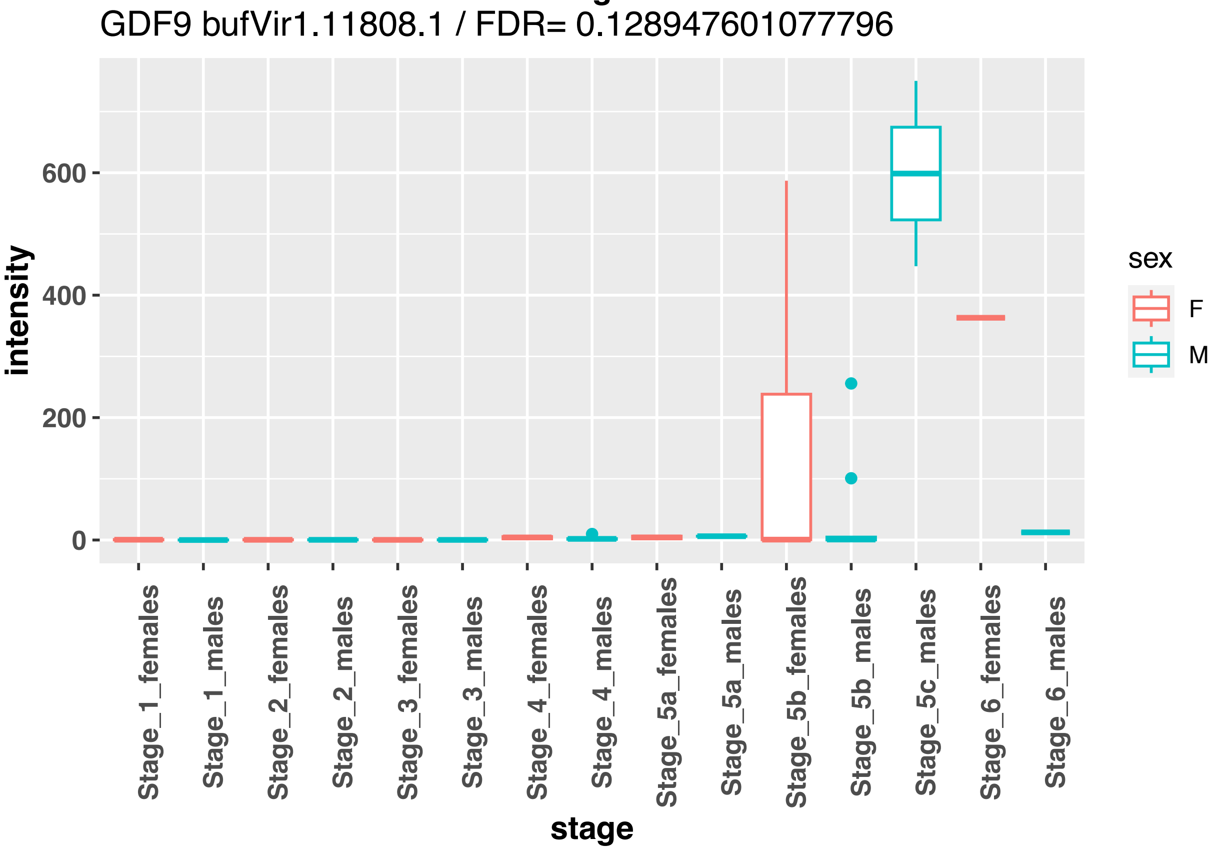
GDF9 bufVir1.11808.1 / FDR= 0.128947601077796
intensity
600
400
200
0
sex
F
M
Stage_1_females
Stage_1_males
Stage_2_females
Stage_2_males
Stage_3_females
Stage_3_males
Stage_4_females
Stage_4_males
Stage_5a_females
Stage_5a_males
Stage_5b_females
Stage_5b_males
Stage_5c_males
Stage_6_females
Stage_6_males
stage

GLI3 bufVir1.25677.1 / FDR= 0.999980732933913

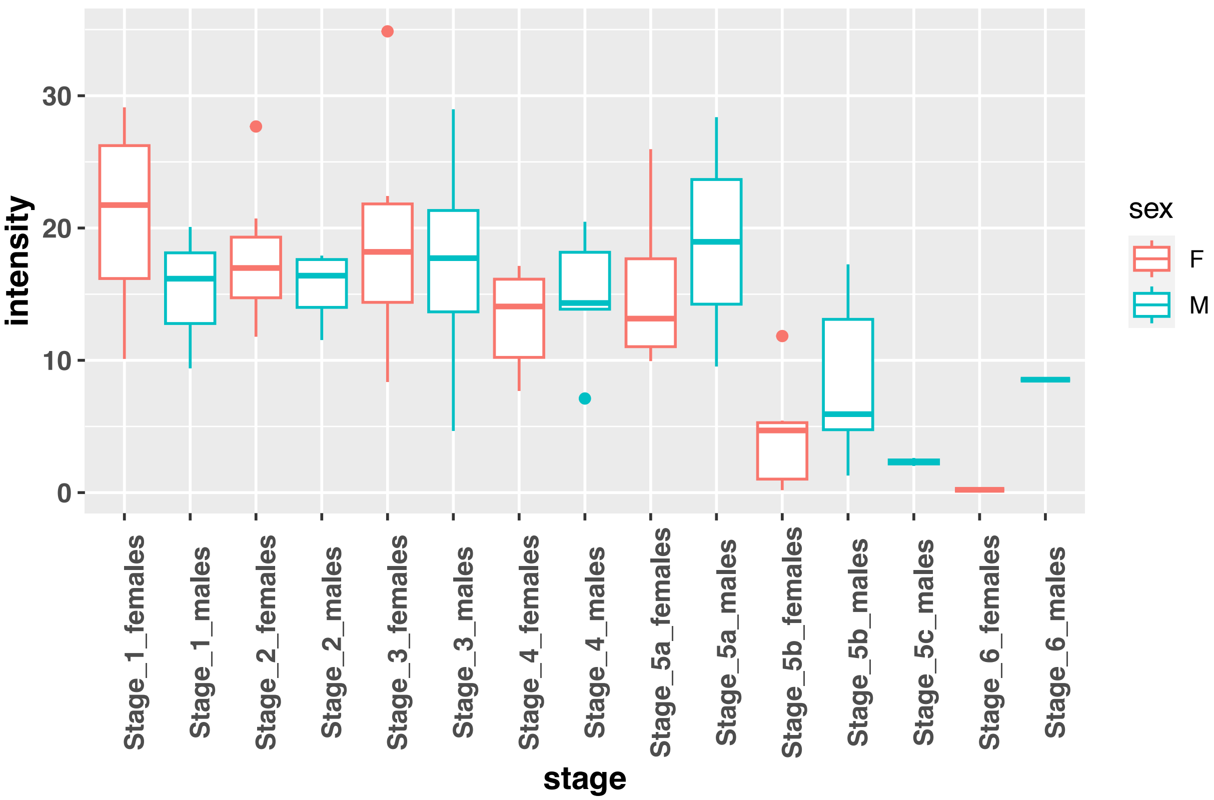


HHIPL2 bufVir1.23474.1 / FDR= 0.999980732933913

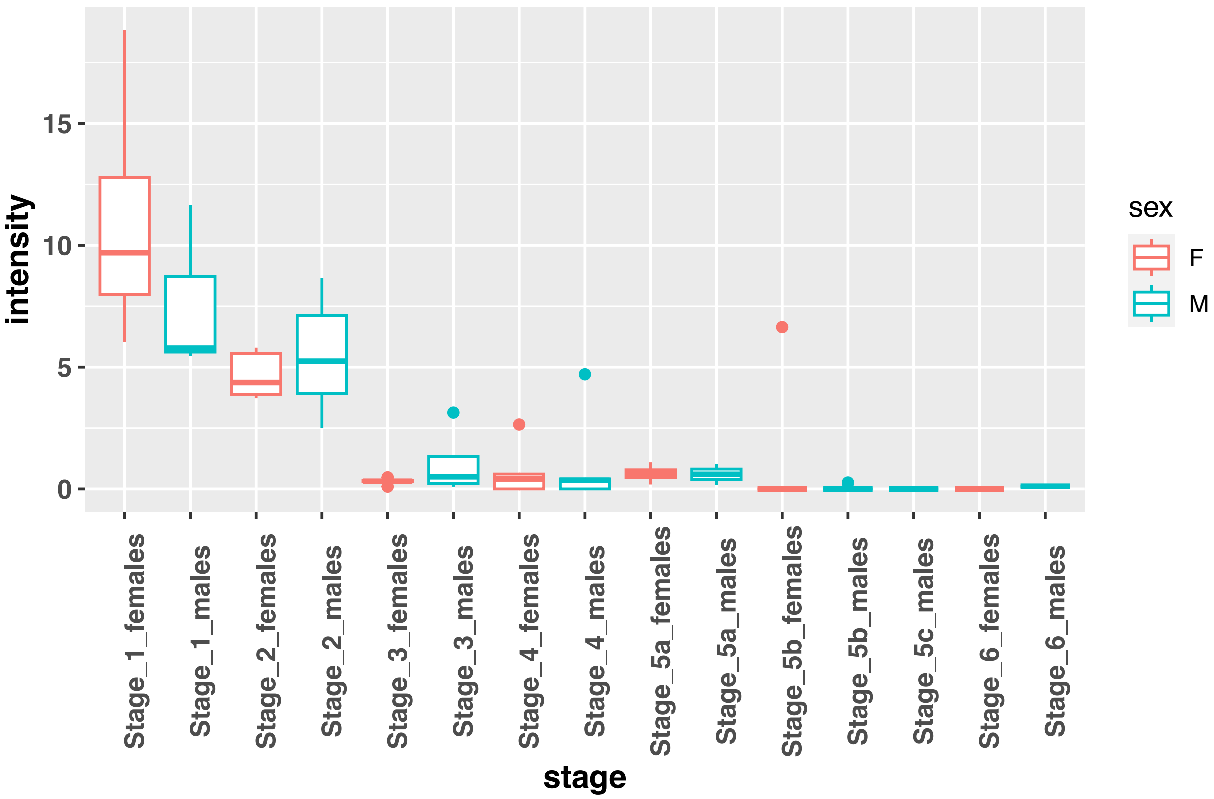

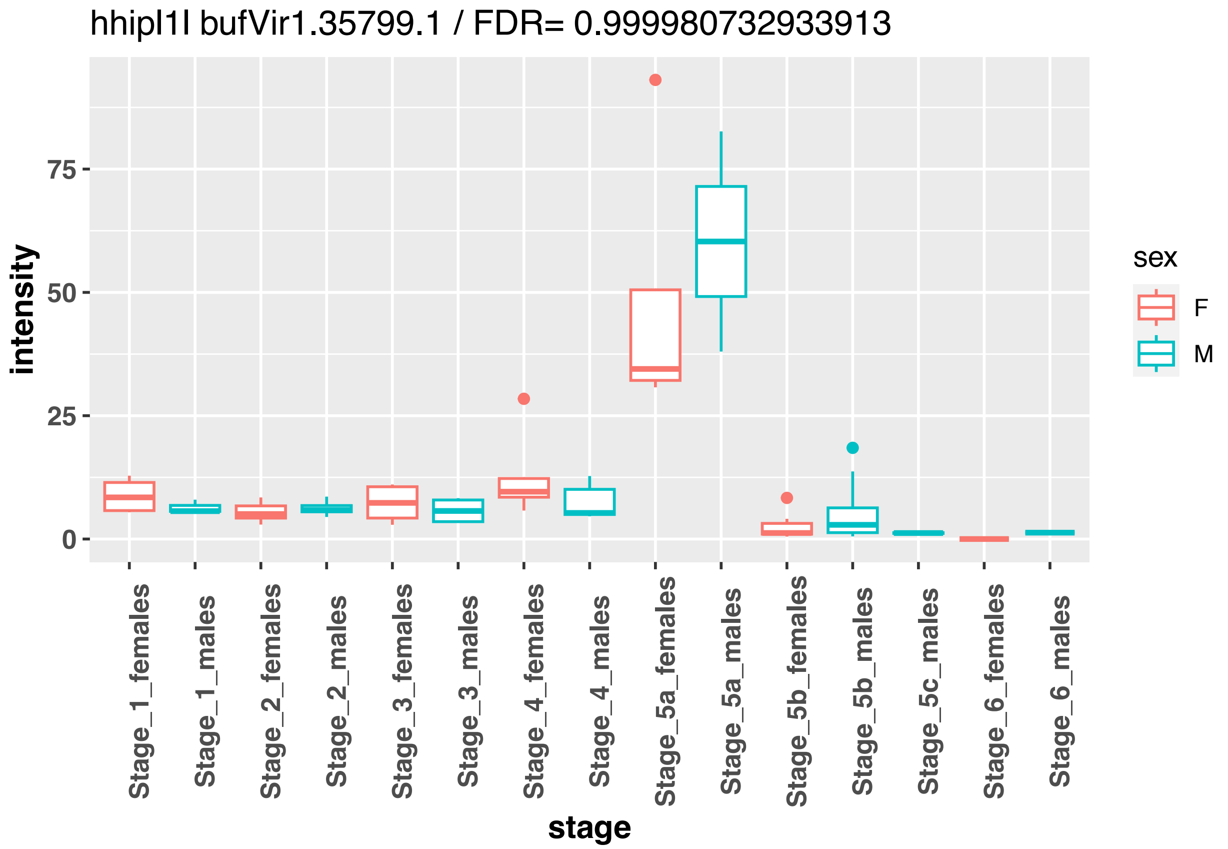
hhipl1l bufVir1.35799.1 / FDR= 0.999980732933913
intensity
75
50
25
0
Stage_1_females
Stage_1_males
Stage_2_females
Stage_2_males
Stage_3_females
Stage_3_males
Stage_4_females
Stage_4_males
Stage_5a_females
Stage_5a_males
Stage_5b_females
Stage_5b_males
Stage_5c_males
Stage_6_females
Stage_6_males
stage
sex
F
M

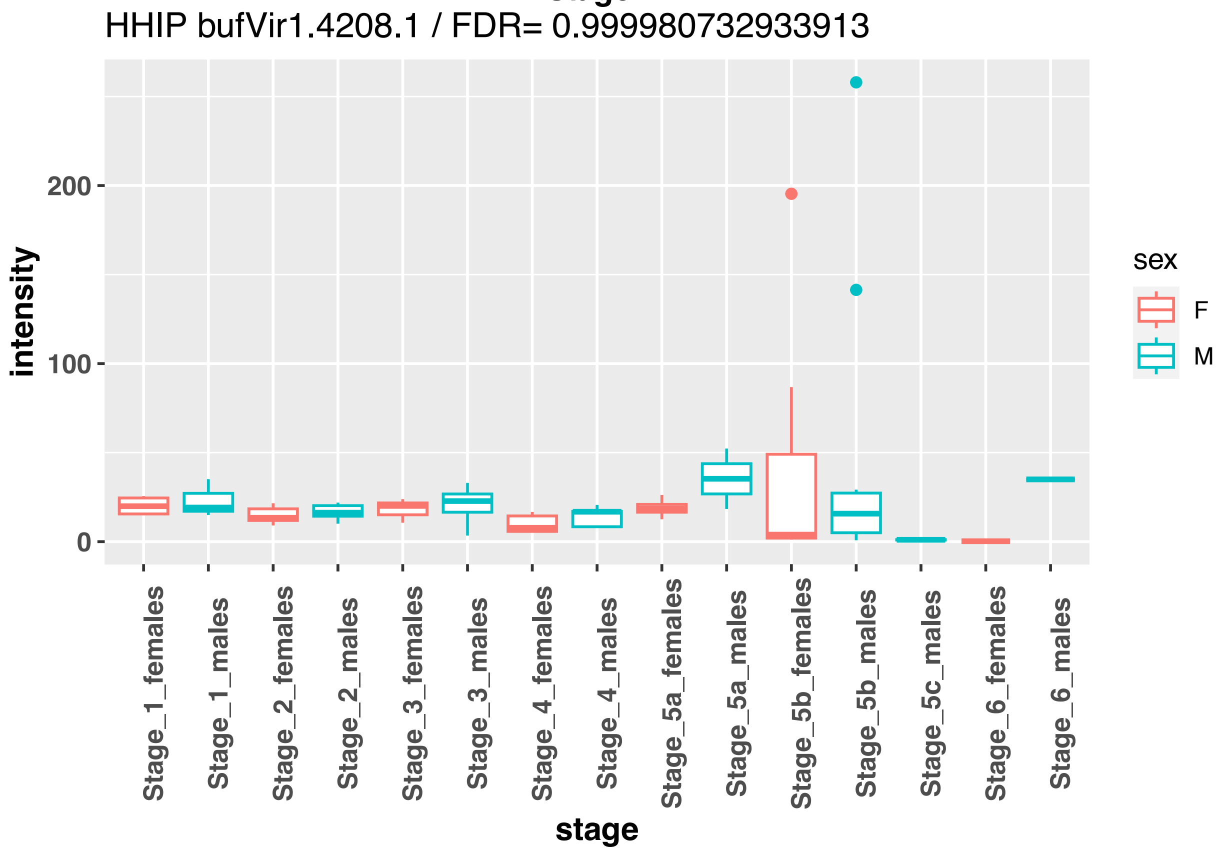
HHIP bufVir1.4208.1 / FDR= 0.999980732933913
intensity
200
100
0
Stage_1_females
Stage_1_males
Stage_2_females
Stage_2_males
Stage_3_females
Stage_3_males
Stage_4_females
Stage_4_males
Stage_5a_females
Stage_5a_males
Stage_5b_females
Stage_5b_males
Stage_5c_males
Stage_6_females
Stage_6_males
stage
sex
F
M

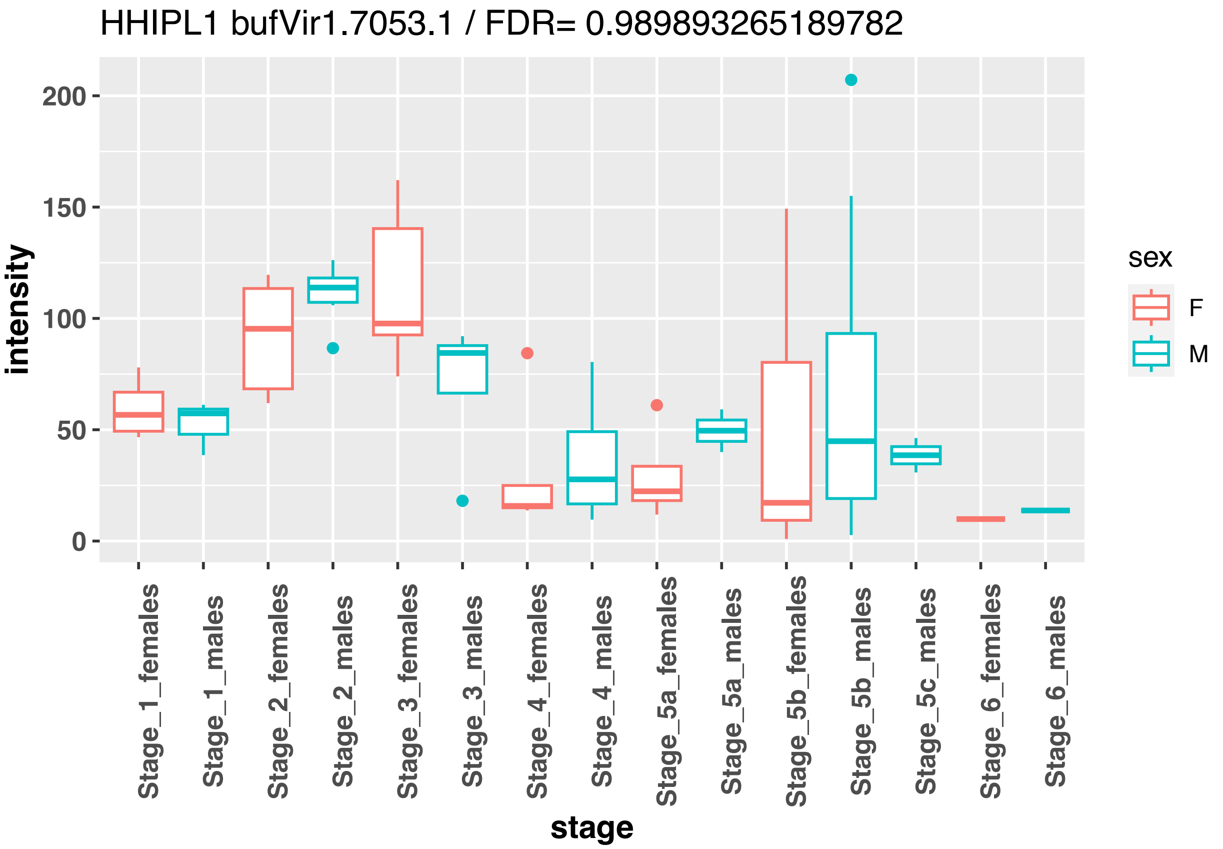
HHIPL1 bufVir1.7053.1 / FDR= 0.989893265189782
intensity
200
150
100
50
0
Stage_1_females
Stage_1_males
Stage_2_females
Stage_2_males
Stage_3_females
Stage_3_males
Stage_4_females
Stage_4_males
Stage_5a_females
Stage_5a_males
Stage_5b_females
Stage_5b_males
Stage_5c_males
Stage_6_females
Stage_6_males
stage
sex
F
M

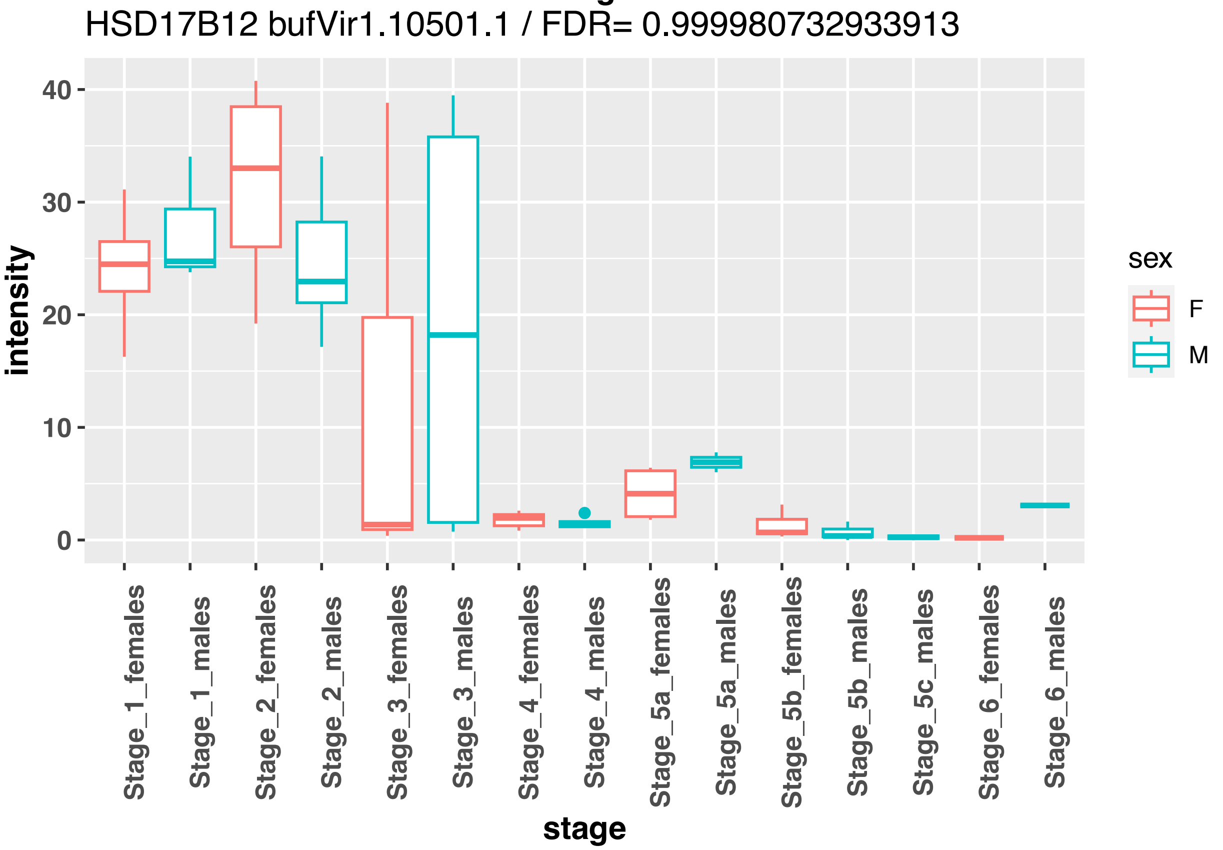
HSD17B12 bufVir1.10501.1 / FDR= 0.999980732933913
intensity
40
30
20
10
0
Stage_1_females
Stage_1_males
Stage_2_females
Stage_2_males
Stage_3_females
Stage_3_males
Stage_4_females
Stage_4_males
Stage_5a_females
Stage_5a_males
Stage_5b_females
Stage_5b_males
Stage_5c_males
Stage_6_females
Stage_6_males
stage
sex
F
M

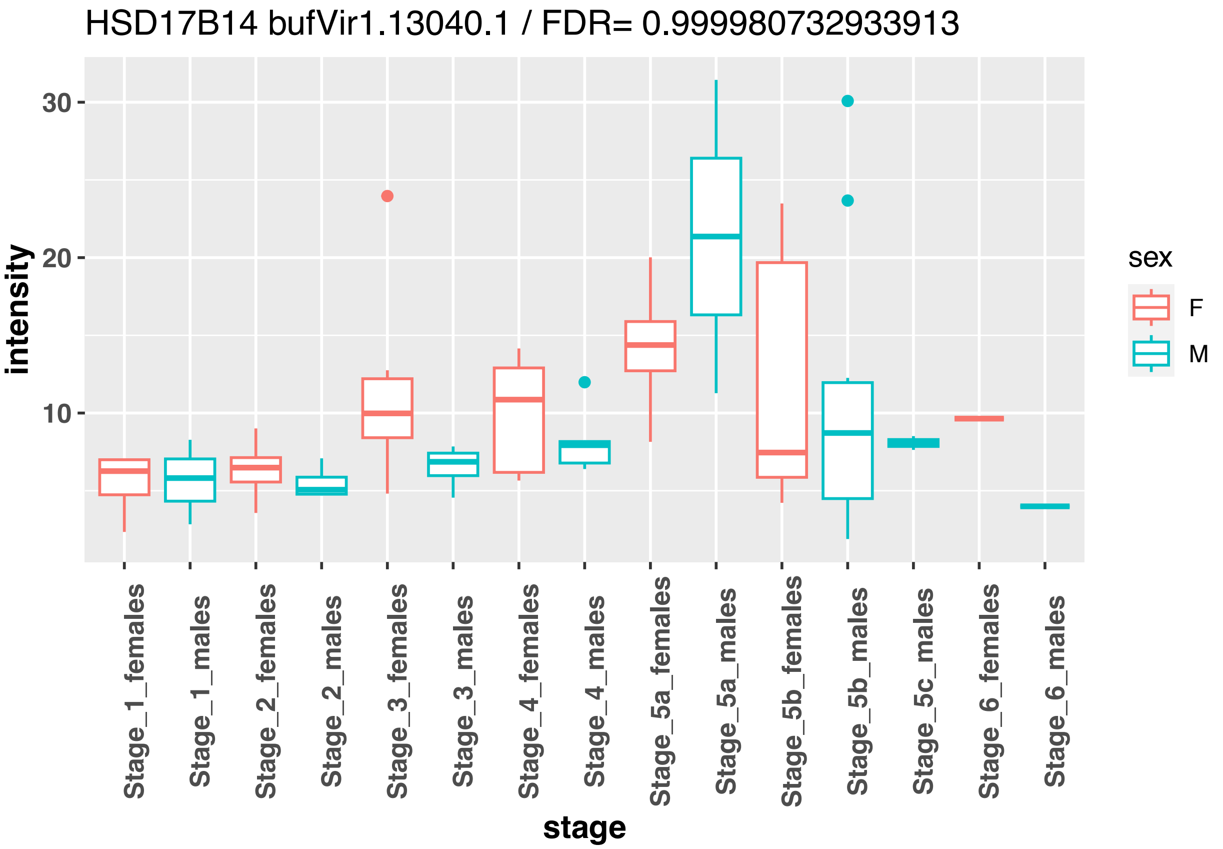
HSD17B14 bufVir1.13040.1 / FDR= 0.999980732933913
30
20
10
intensity
Stage_1_females
Stage_1_males
Stage_2_females
Stage_2_males
Stage_3_females
Stage_3_males
Stage_4_females
Stage_4_males
Stage_5a_females
Stage_5a_males
Stage_5b_females
Stage_5b_males
Stage_5c_males
Stage_6_females
Stage_6_males
stage
sex
F
M

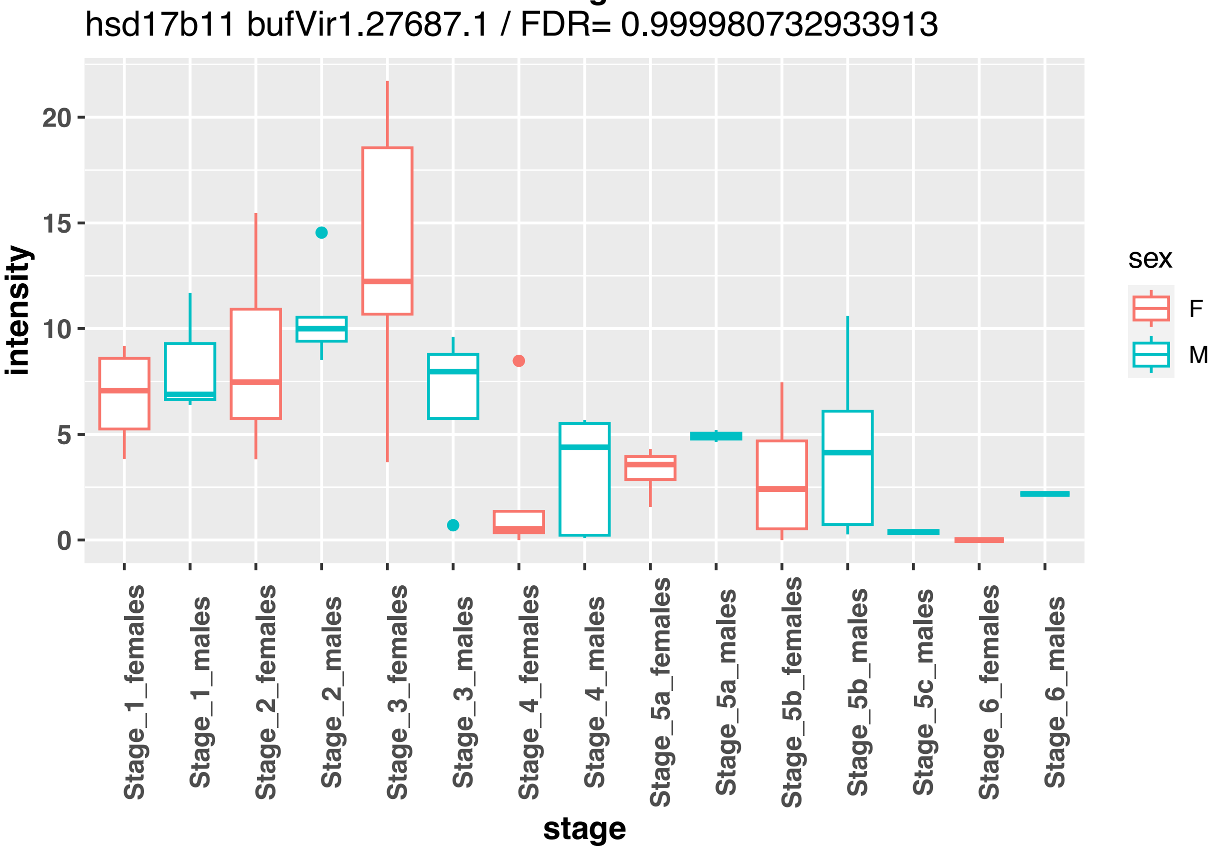
hsd17b11 bufVir1.27687.1 / FDR= 0.999980732933913
20
15
10
5
0
intensity
Stage_1_females
Stage_1_males
Stage_2_females
Stage_2_males
Stage_3_females
Stage_3_males
Stage_4_females
Stage_4_males
Stage_5a_females
Stage_5a_males
Stage_5b_females
Stage_5b_males
Stage_5c_males
Stage_6_females
Stage_6_males
stage
sex
F
M

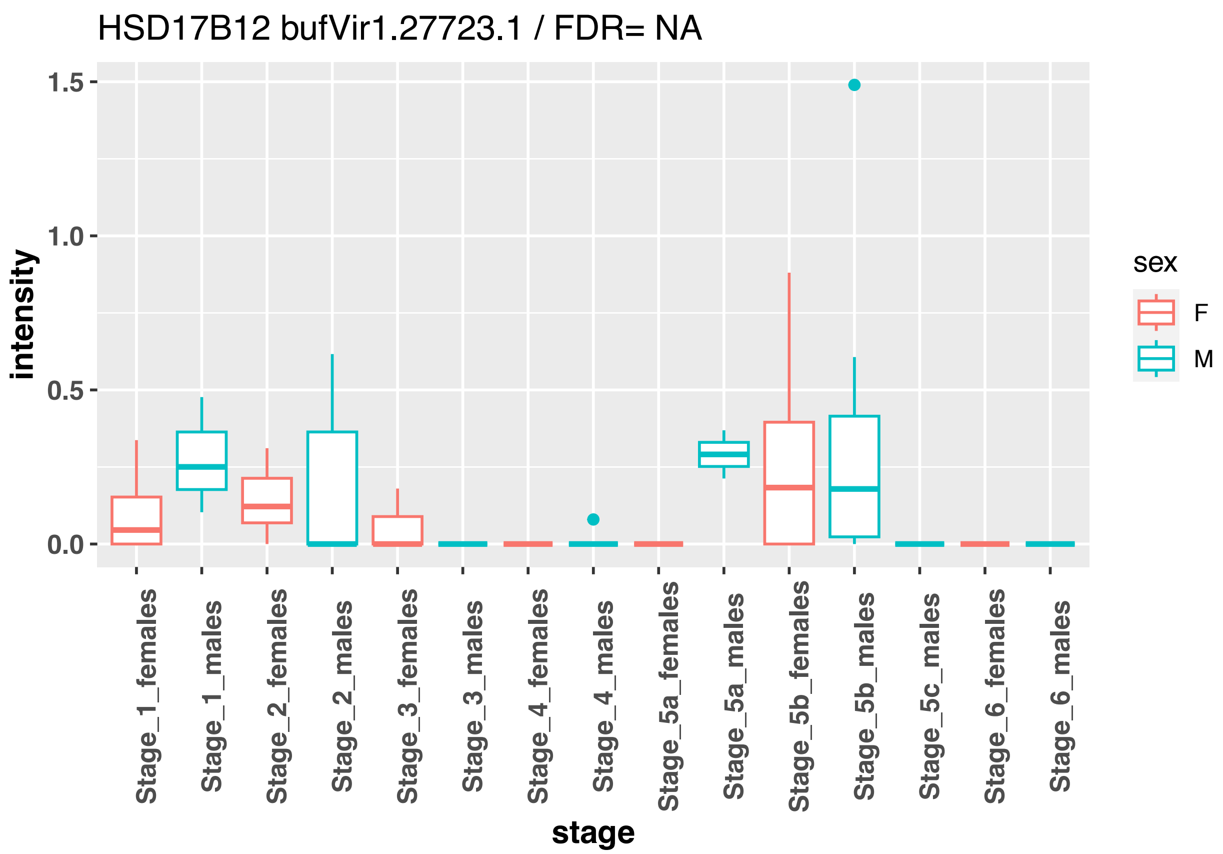
HSD17B12 bufVir1.27723.1 / FDR= NA
intensity
1.5
1.0
0.5
0.0
Stage_1_females
Stage_1_males
Stage_2_females
Stage_2_males
Stage_3_females
Stage_3_males
Stage_4_females
Stage_4_males
Stage_5a_females
Stage_5a_males
Stage_5b_females
Stage_5b_males
Stage_5c_males
Stage_6_females
Stage_6_males
stage
sex
F
M

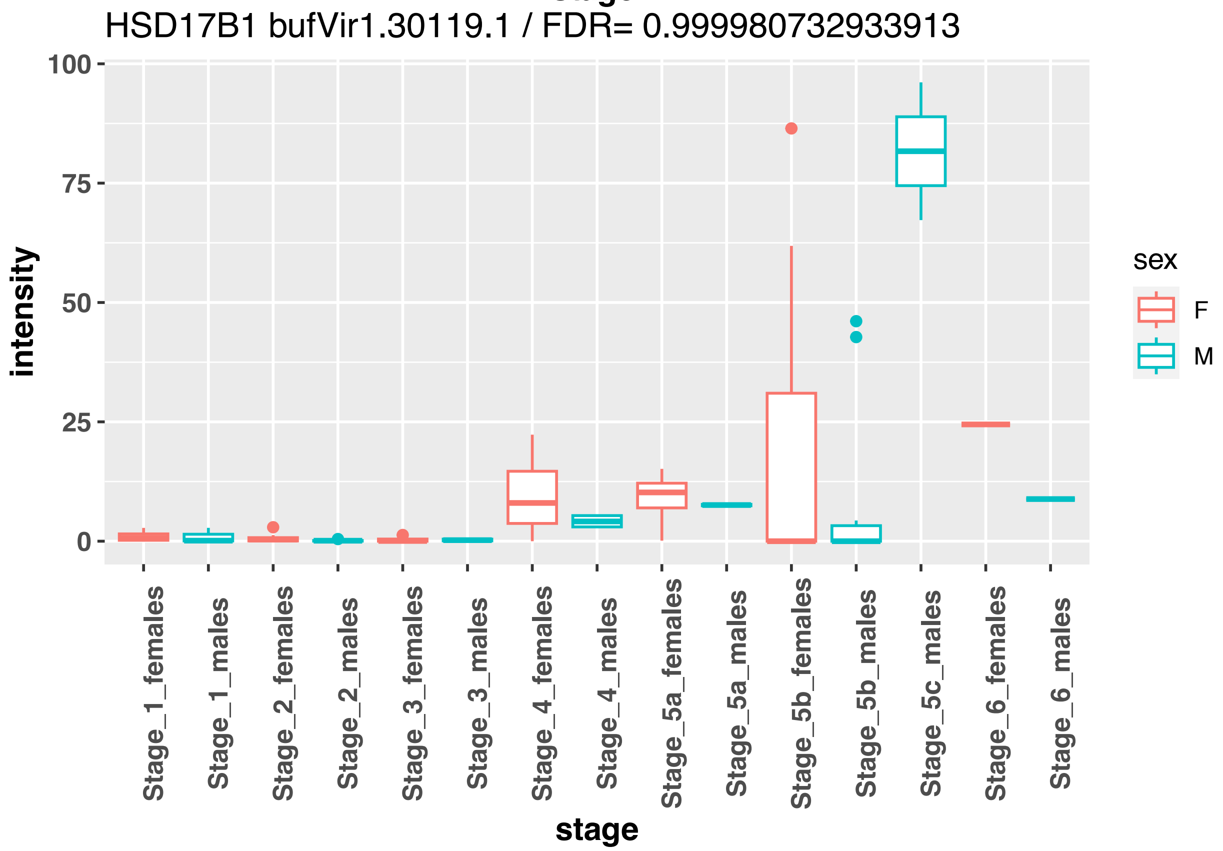
HSD17B1 bufVir1.30119.1 / FDR= 0.999980732933913
intensity
100
75
50
25
0
Stage_1_females
Stage_1_males
Stage_2_females
Stage_2_males
Stage_3_females
Stage_3_males
Stage_4_females
Stage_4_males
Stage_5a_females
Stage_5a_males
Stage_5b_females
Stage_5b_males
Stage_5c_males
Stage_6_females
Stage_6_males
stage
sex
F
M

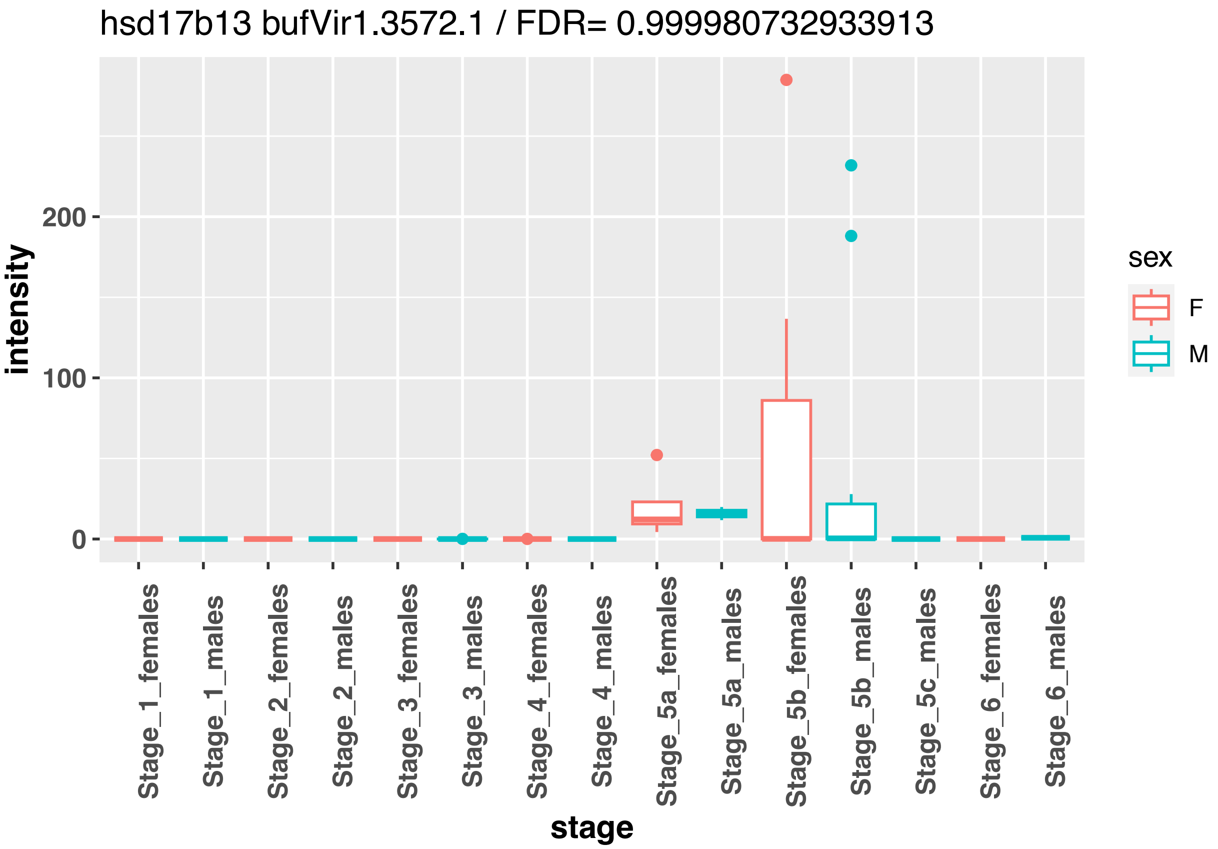
hsd17b13 bufVir1.3572.1 / FDR= 0.999980732933913
intensity
200
100
0
sex
F
M
Stage_1_females
Stage_1_males
Stage_2_females
Stage_2_males
Stage_3_females
Stage_3_males
Stage_4_females
Stage_4_males
Stage_5a_females
Stage_5a_males
Stage_5b_females
Stage_5b_males
Stage_5c_males
Stage_6_females
Stage_6_males
stage

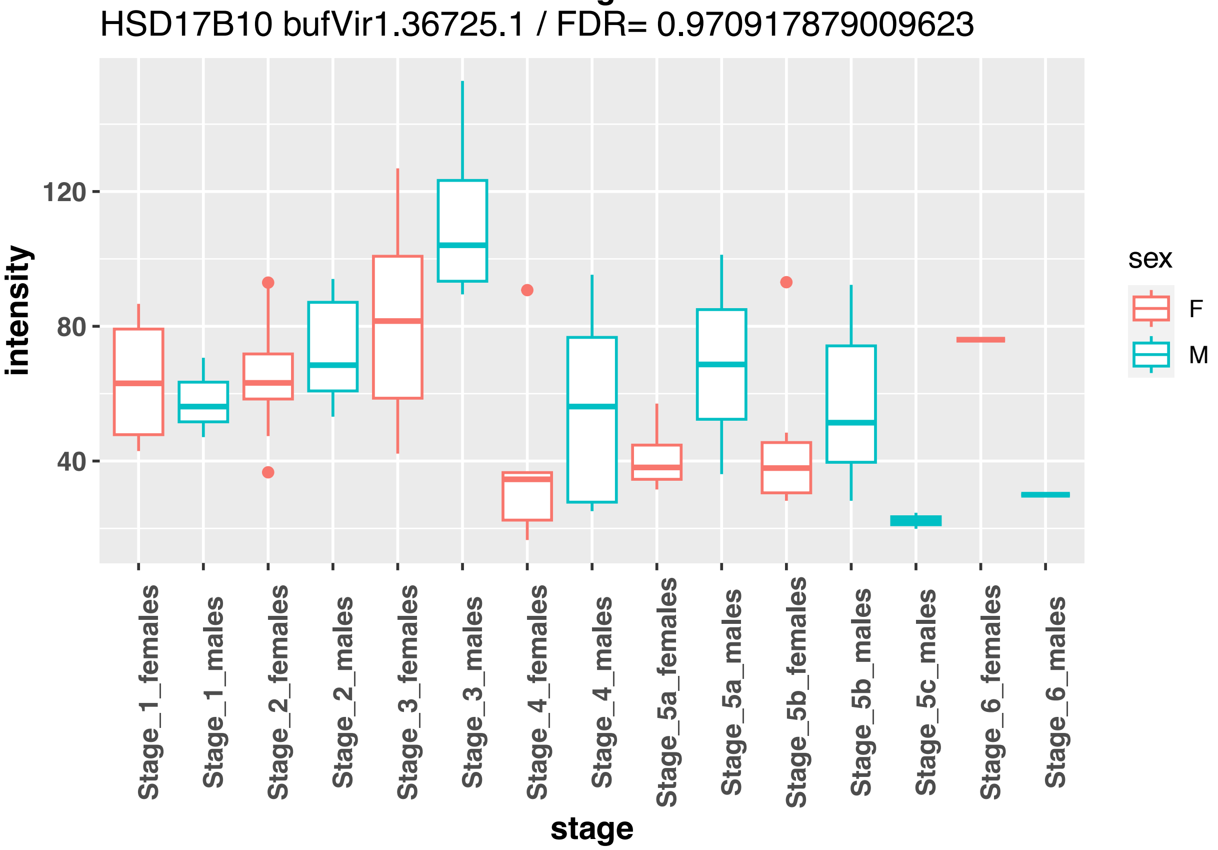
HSD17B10 bufVir1.36725.1 / FDR= 0.970917879009623
intensity
120
80
40
sex
F
M
Stage_1_females
Stage_1_males
Stage_2_females
Stage_2_males
Stage_3_females
Stage_3_males
Stage_4_females
Stage_4_males
Stage_5a_females
Stage_5a_males
Stage_5b_females
Stage_5b_males
Stage_5c_males
Stage_6_females
Stage_6_males
stage

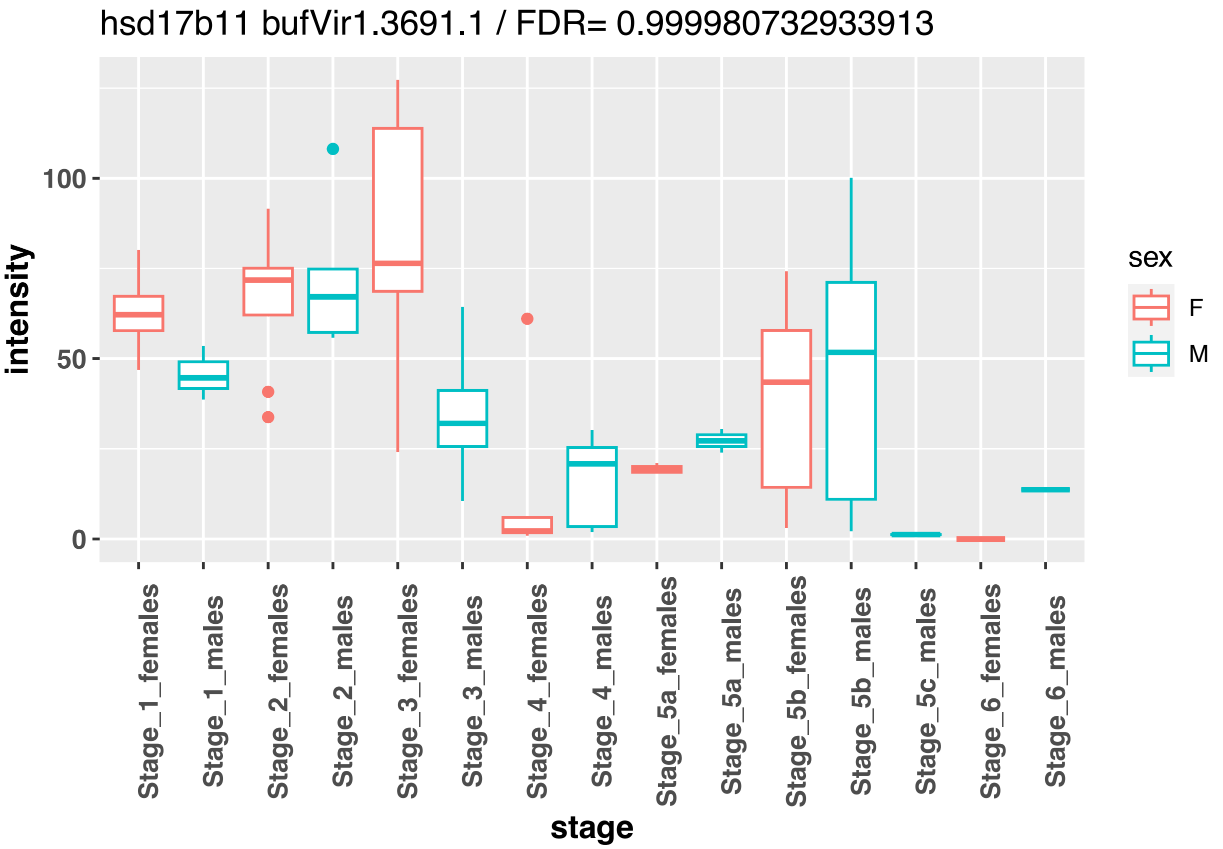

hsd17b11 bufVir1.3691.1 / FDR= 0.999980732933913
intensity
100
50
0
Stage_1_females
Stage_1_males
Stage_2_females
Stage_2_males
Stage_3_females
Stage_3_males
Stage_4_females
Stage_4_males
Stage_5a_females
Stage_5a_males
Stage_5b_females
Stage_5b_males
Stage_5c_males
Stage_6_females
Stage_6_males
stage
sex
F
M


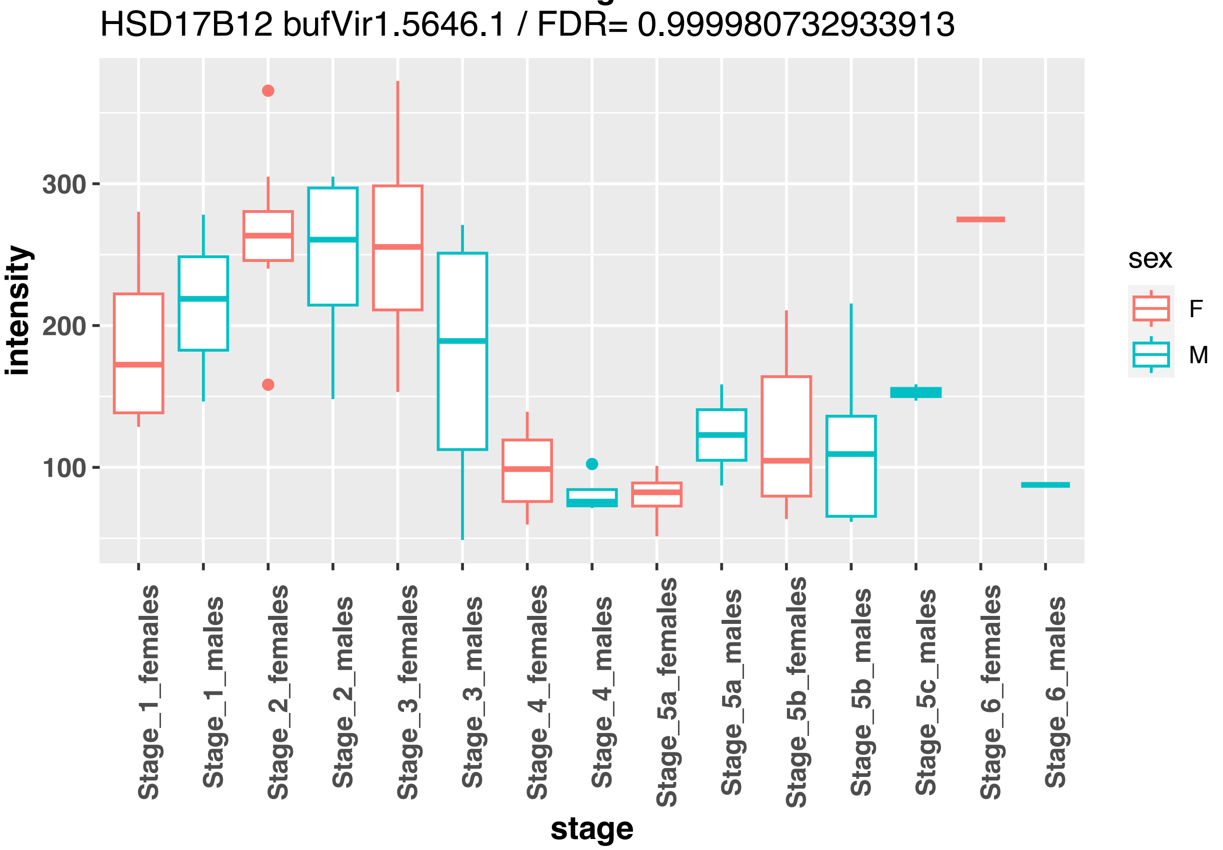

HSD17B12 bufVir1.5646.1 / FDR= 0.999980732933913
intensity
300
200
100
Stage_1_females
Stage_1_males
Stage_2_females
Stage_2_males
Stage_3_females
Stage_3_males
Stage_4_females
Stage_4_males
Stage_5a_females
Stage_5a_males
Stage_5b_females
Stage_5b_males
Stage_5c_males
Stage_6_females
Stage_6_males
stage
sex
F
M

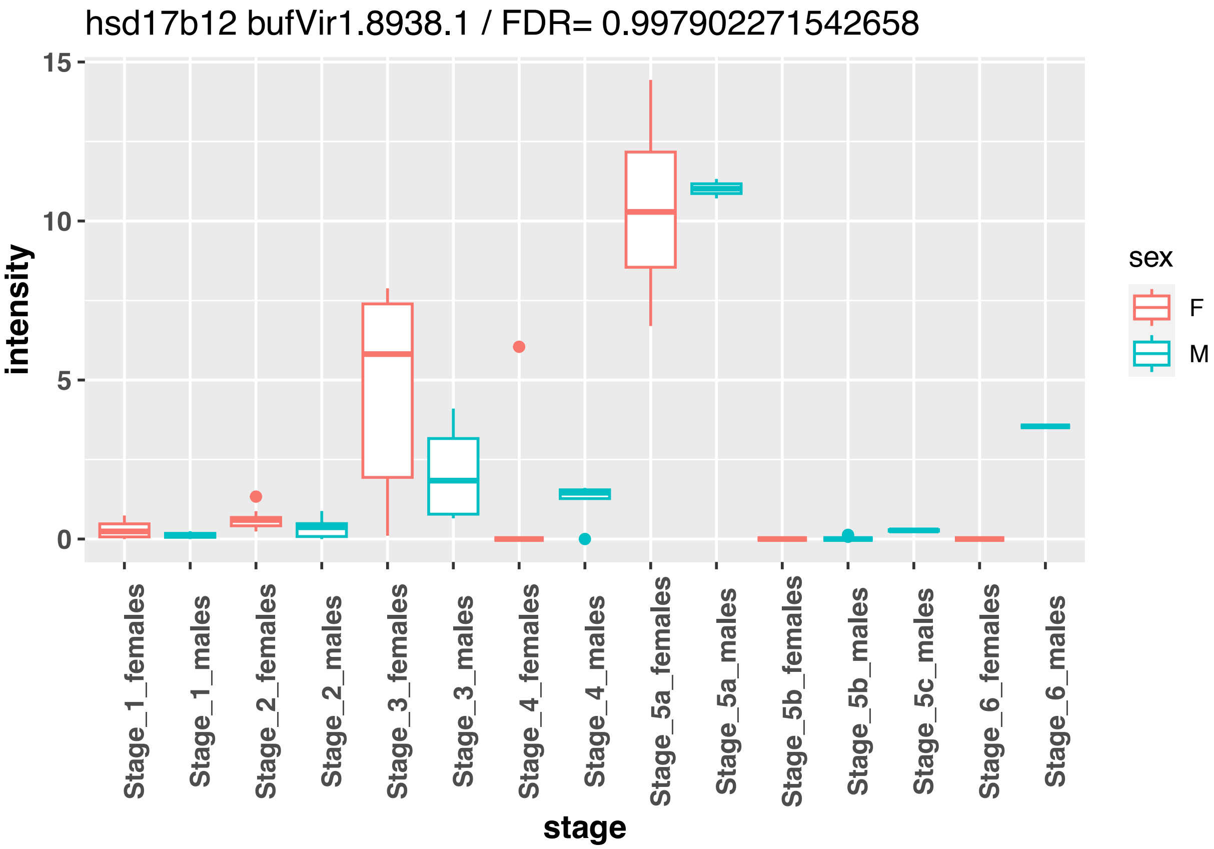
hsd17b12 bufVir1.8938.1 / FDR= 0.997902271542658
intensity
stage
sex
F
M
Stage_1_females
Stage_1_males
Stage_2_females
Stage_2_males
Stage_3_females
Stage_3_males
Stage_4_females
Stage_4_males
Stage_5a_females
Stage_5a_males
Stage_5b_females
Stage_5b_males
Stage_5c_males
Stage_6_females
Stage_6_males

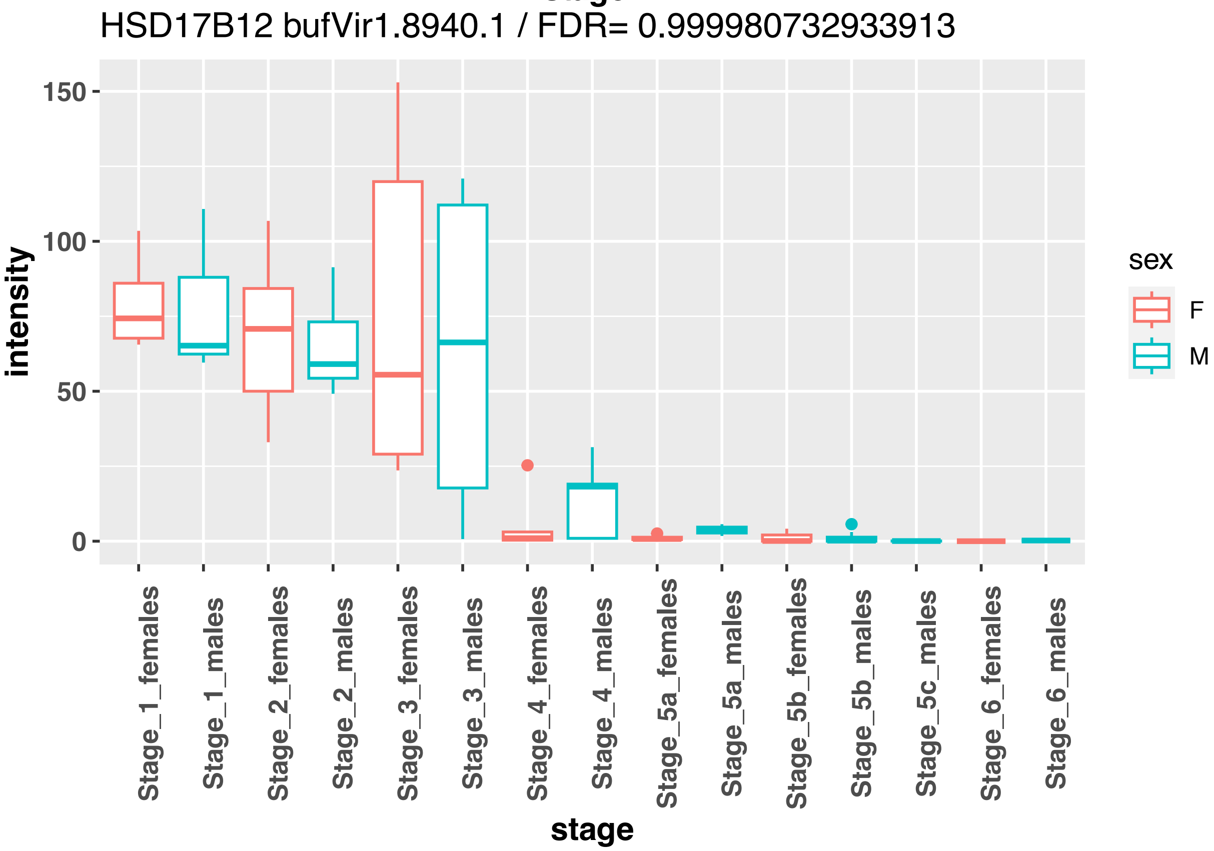
HSD17B12 bufVir1.8940.1 / FDR= 0.999980732933913
intensity
stage
sex
F
M
Stage_1_females
Stage_1_males
Stage_2_females
Stage_2_males
Stage_3_females
Stage_3_males
Stage_4_females
Stage_4_males
Stage_5a_females
Stage_5a_males
Stage_5b_females
Stage_5b_males
Stage_5c_males
Stage_6_females
Stage_6_males

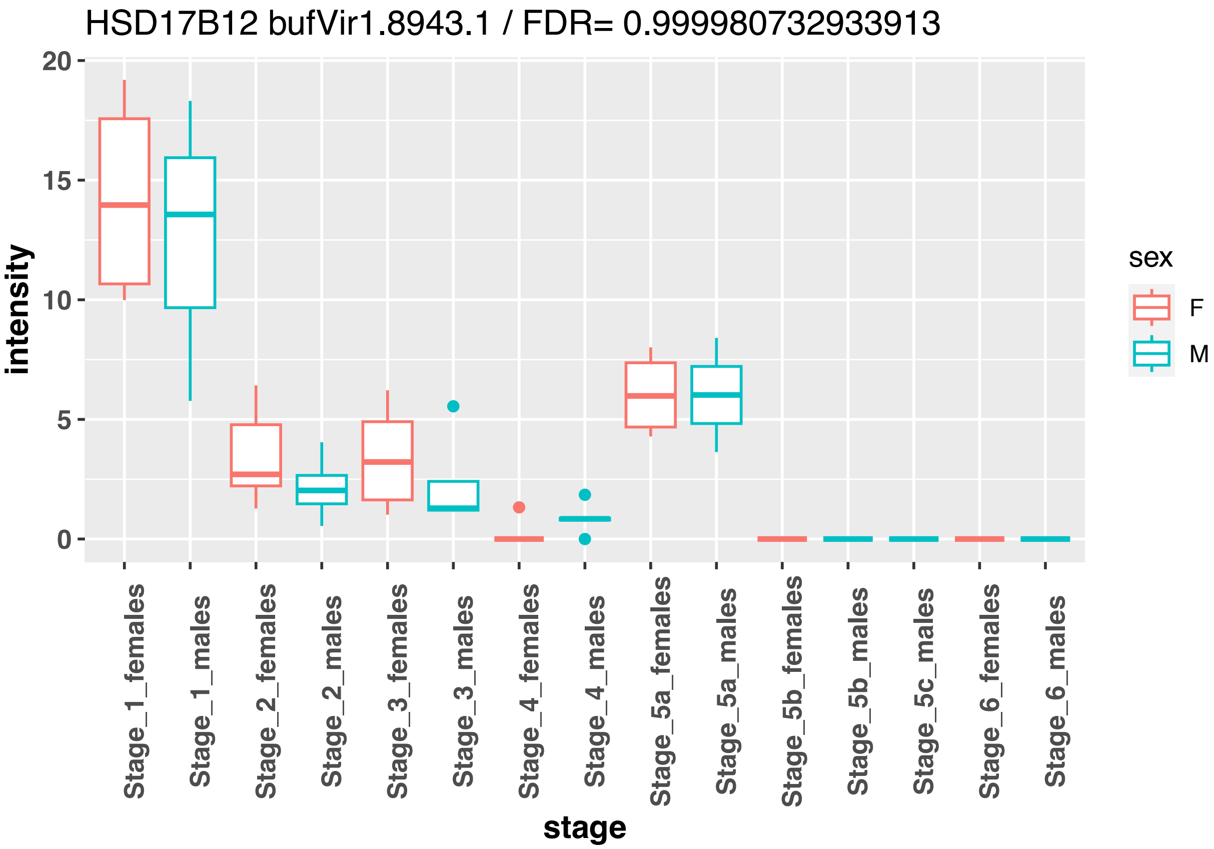

HSD17B12 bufVir1.8943.1 / FDR= 0.999980732933913
intensity
20
15
10
5
0
Stage_1_females
Stage_1_males
Stage_2_females
Stage_2_males
Stage_3_females
Stage_3_males
Stage_4_females
Stage_4_males
Stage_5a_females
Stage_5a_males
Stage_5b_females
Stage_5b_males
Stage_5c_males
Stage_6_females
Stage_6_males
stage
sex
F
M


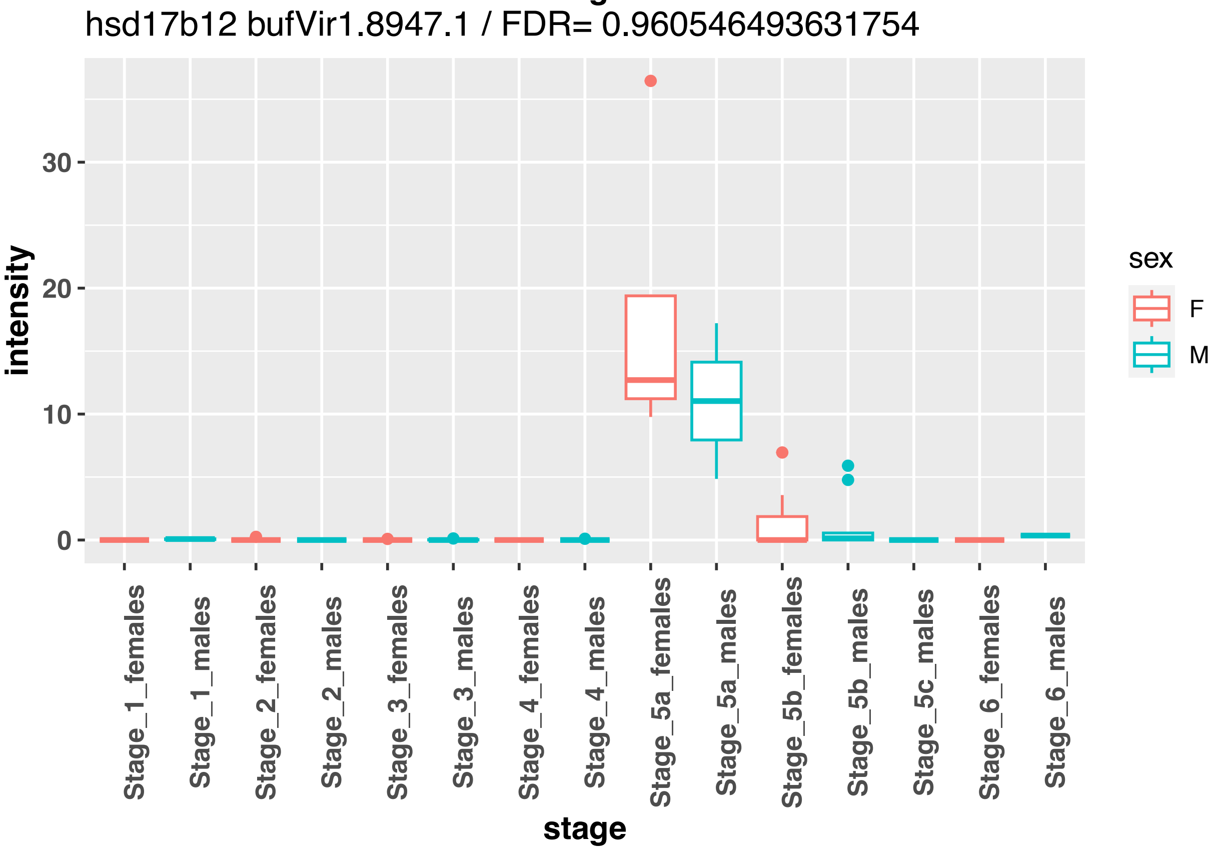

hsd17b12 bufVir1.8947.1 / FDR= 0.960546493631754
intensity
30
20
10
0
Stage_1_females
Stage_1_males
Stage_2_females
Stage_2_males
Stage_3_females
Stage_3_males
Stage_4_females
Stage_4_males
Stage_5a_females
Stage_5a_males
Stage_5b_females
Stage_5b_males
Stage_5c_males
Stage_6_females
Stage_6_males
stage
sex
F
M

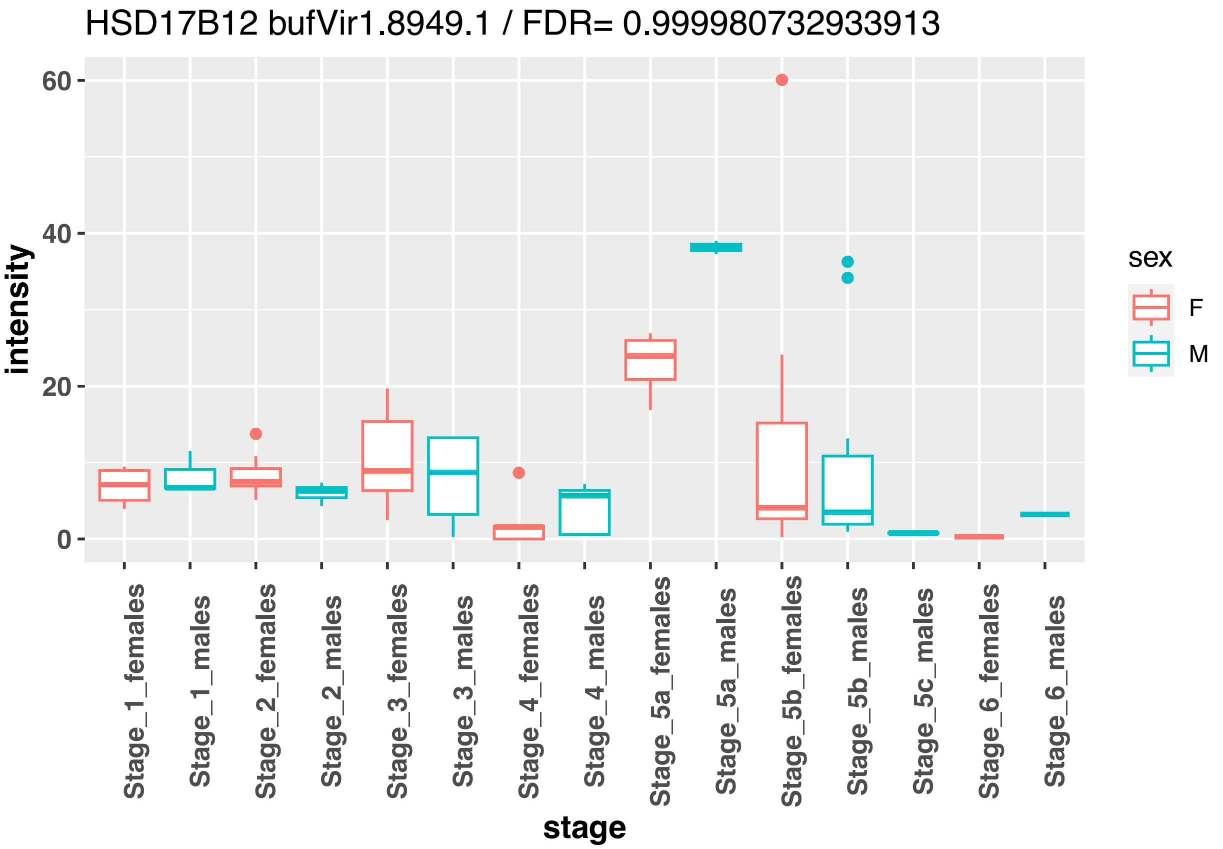
HSD17B12 bufVir1.8949.1 / FDR= 0.999980732933913
intensity
0
20
40
60
Stage_1_females
Stage_1_males
Stage_2_females
Stage_2_males
Stage_3_females
Stage_3_males
Stage_4_females
Stage_4_males
Stage_5a_females
Stage_5a_males
Stage_5b_females
Stage_5b_males
Stage_5c_males
Stage_6_females
Stage_6_males
stage
sex
F
M

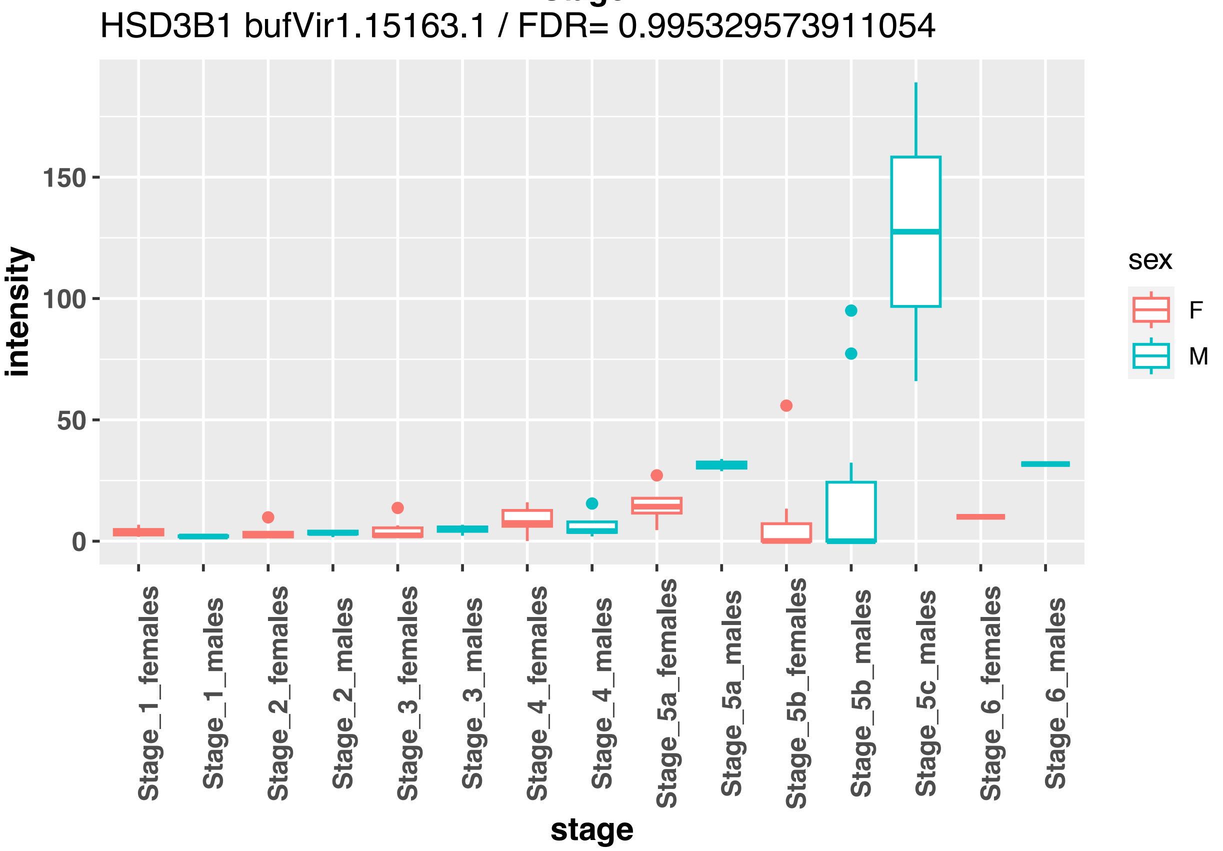
HSD3B1 bufVir1.15163.1 / FDR= 0.995329573911054
intensity
0
50
100
150
Stage_1_females
Stage_1_males
Stage_2_females
Stage_2_males
Stage_3_females
Stage_3_males
Stage_4_females
Stage_4_males
Stage_5a_females
Stage_5a_males
Stage_5b_females
Stage_5b_males
Stage_5c_males
Stage_6_females
Stage_6_males
stage
sex
F
M

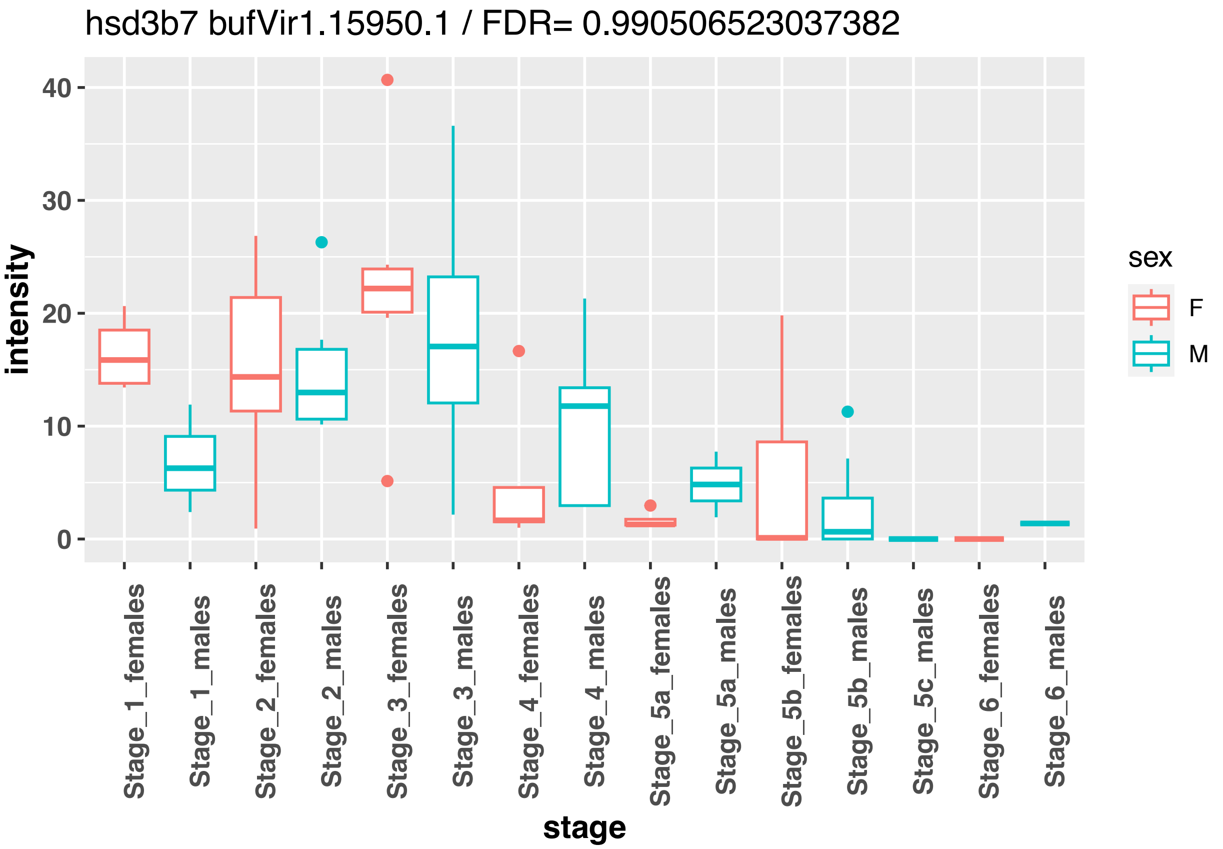
hsd3b7 bufVir1.15950.1 / FDR= 0.990506523037382
intensity
stage
sex
F
M
Stage_1_females
Stage_1_males
Stage_2_females
Stage_2_males
Stage_3_females
Stage_3_males
Stage_4_females
Stage_4_males
Stage_5a_females
Stage_5a_males
Stage_5b_females
Stage_5b_males
Stage_5c_males
Stage_6_females
Stage_6_males

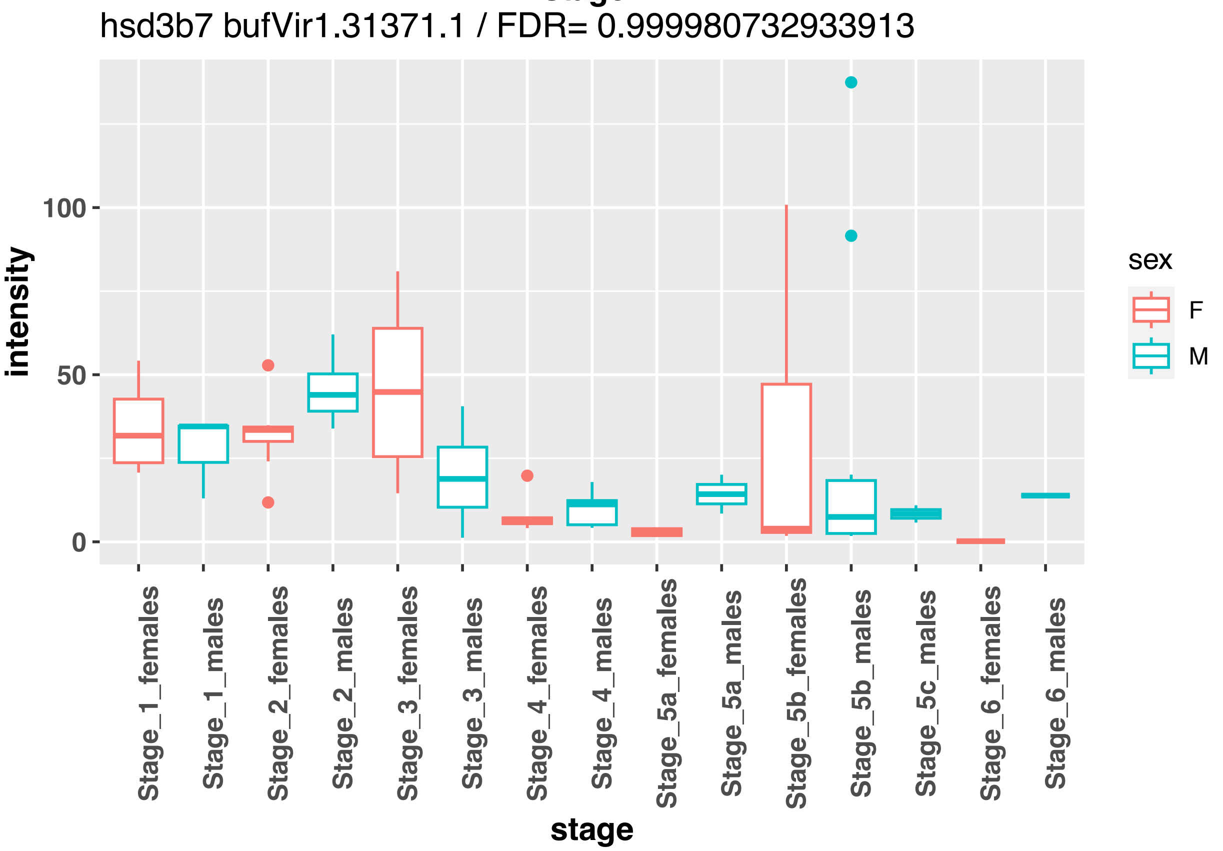
hsd3b7 bufVir1.31371.1 / FDR= 0.999980732933913
intensity
stage
sex
F
M
Stage_1_females
Stage_1_males
Stage_2_females
Stage_2_males
Stage_3_females
Stage_3_males
Stage_4_females
Stage_4_males
Stage_5a_females
Stage_5a_males
Stage_5b_females
Stage_5b_males
Stage_5c_males
Stage_6_females
Stage_6_males

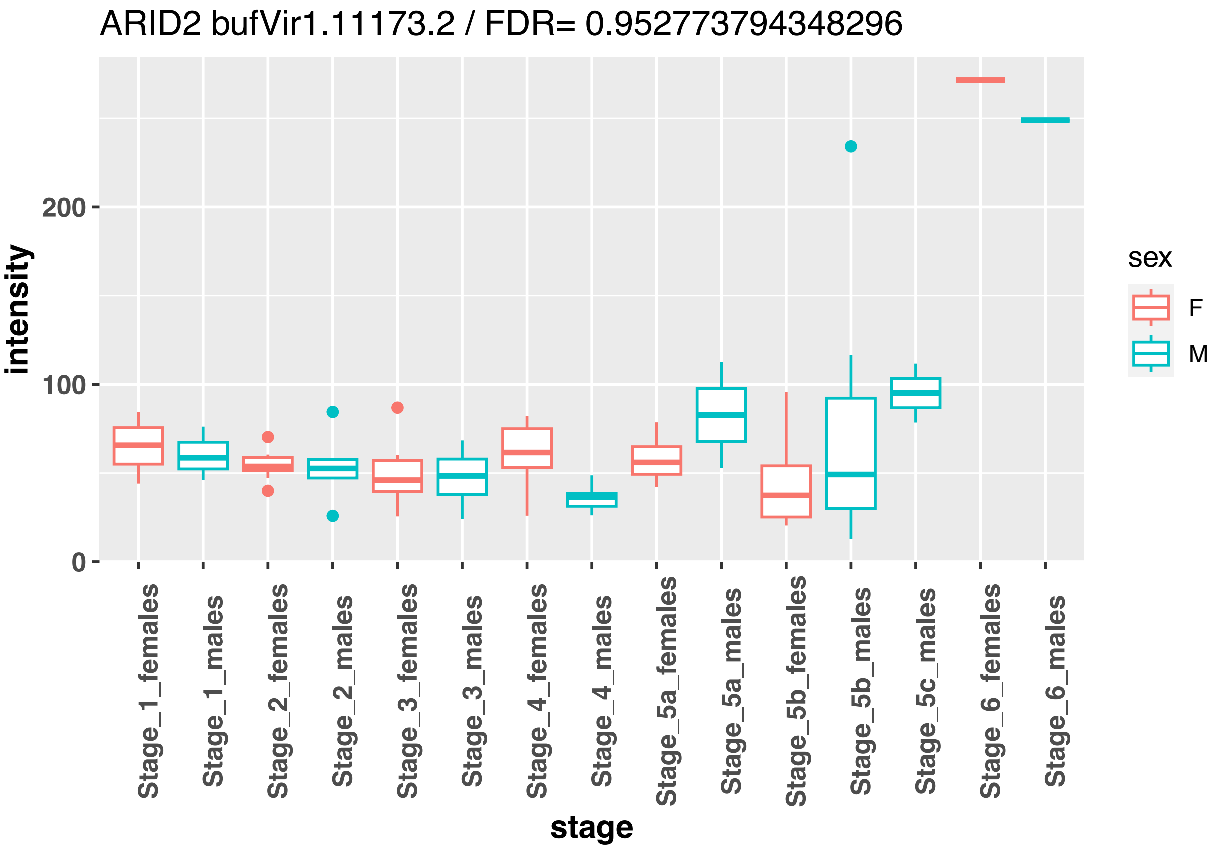
ARID2 bufVir1.11173.2 / FDR= 0.952773794348296
intensity
0
100
200
stage
Stage_1_females
Stage_1_males
Stage_2_females
Stage_2_males
Stage_3_females
Stage_3_males
Stage_4_females
Stage_4_males
Stage_5a_females
Stage_5a_males
Stage_5b_females
Stage_5b_males
Stage_5c_males
Stage_6_females
Stage_6_males
sex
F
M

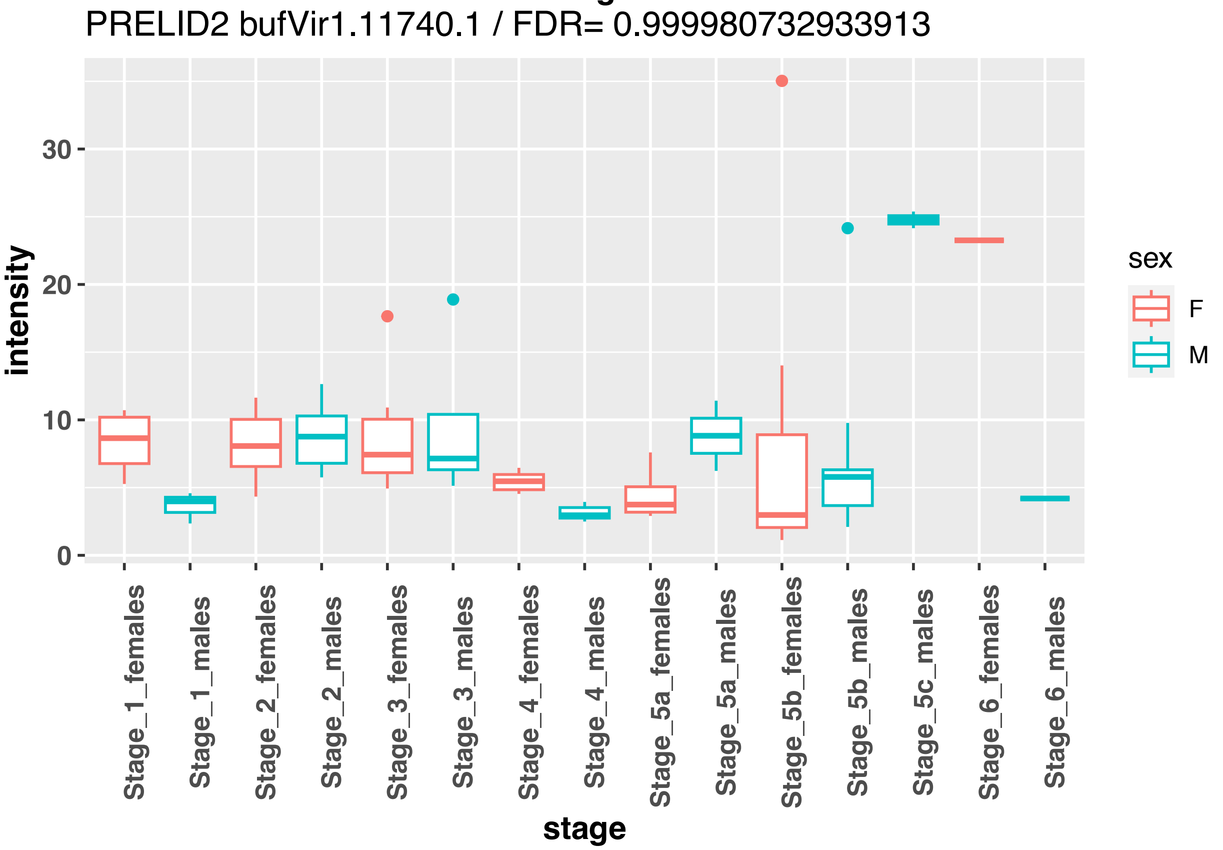
PRELID2 bufVir1.11740.1 / FDR= 0.999980732933913
intensity
0
10
20
30
stage
Stage_1_females
Stage_1_males
Stage_2_females
Stage_2_males
Stage_3_females
Stage_3_males
Stage_4_females
Stage_4_males
Stage_5a_females
Stage_5a_males
Stage_5b_females
Stage_5b_males
Stage_5c_males
Stage_6_females
Stage_6_males
sex
F
M

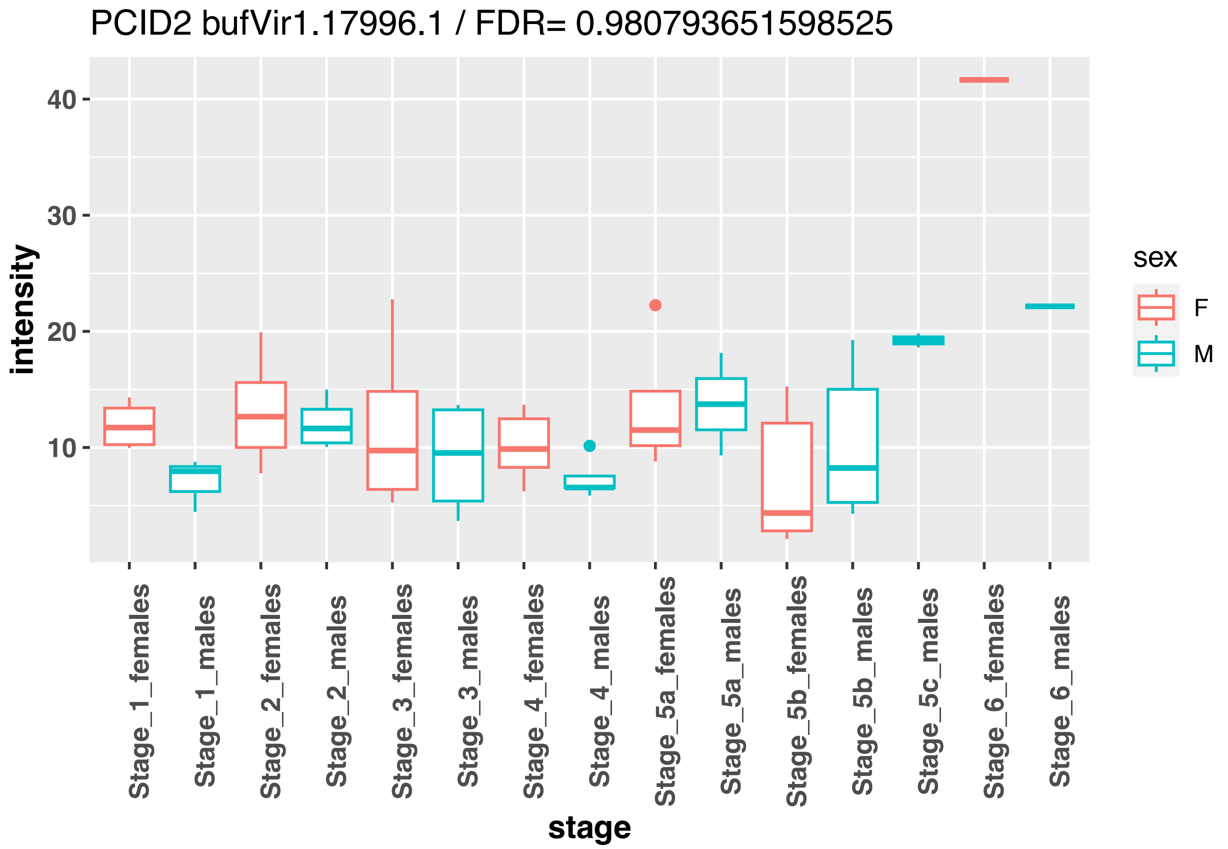
PCID2 bufVir1.17996.1 / FDR= 0.980793651598525
intensity
40
30
20
10
Stage_1_females
Stage_1_males
Stage_2_females
Stage_2_males
Stage_3_females
Stage_3_males
Stage_4_females
Stage_4_males
Stage_5a_females
Stage_5a_males
Stage_5b_females
Stage_5b_males
Stage_5c_males
Stage_6_females
Stage_6_males
stage
sex
F
M

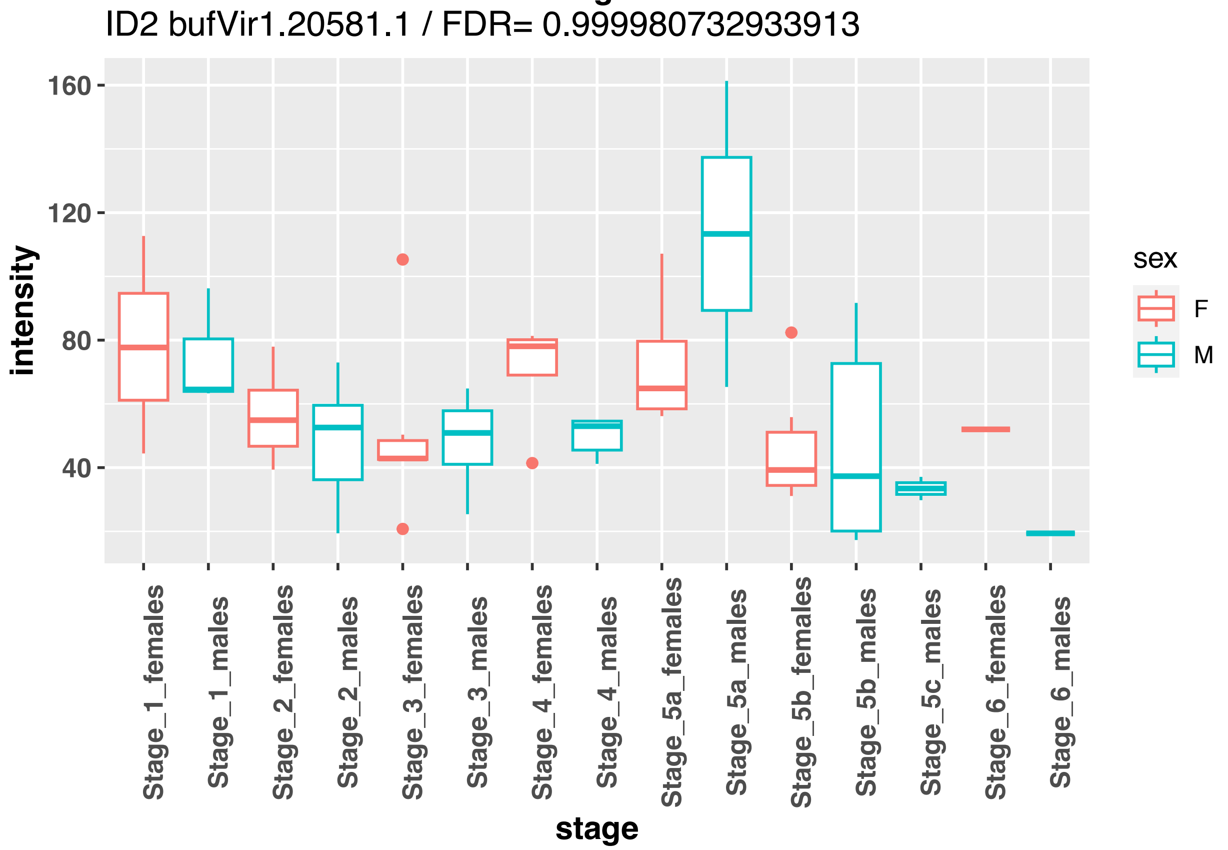
ID2 bufVir1.20581.1 / FDR= 0.999980732933913
intensity
160
120
80
40
Stage_1_females
Stage_1_males
Stage_2_females
Stage_2_males
Stage_3_females
Stage_3_males
Stage_4_females
Stage_4_males
Stage_5a_females
Stage_5a_males
Stage_5b_females
Stage_5b_males
Stage_5c_males
Stage_6_females
Stage_6_males
stage
sex
F
M

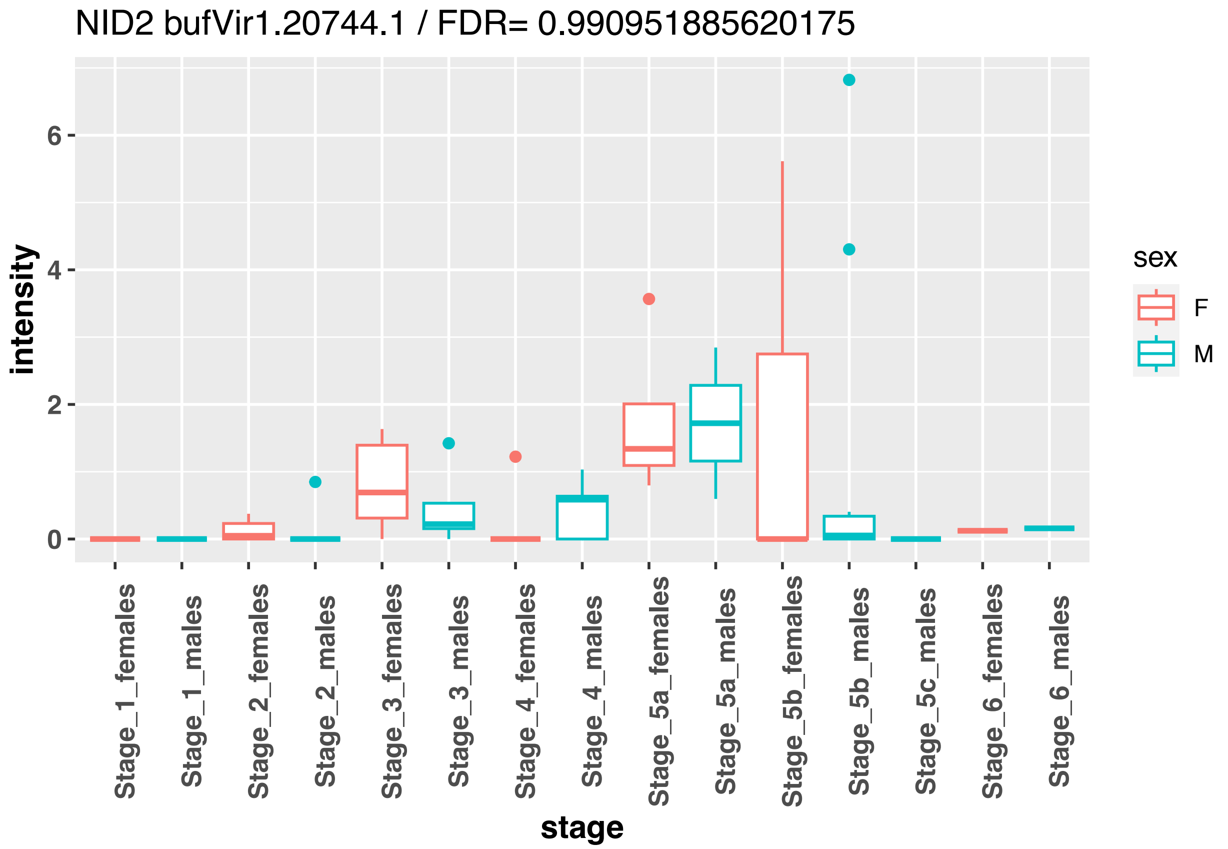
NID2 bufVir1.20744.1 / FDR= 0.990951885620175
intensity
6
4
2
0
stage
Stage_1_females
Stage_1_males
Stage_2_females
Stage_2_males
Stage_3_females
Stage_3_males
Stage_4_females
Stage_4_males
Stage_5a_females
Stage_5a_males
Stage_5b_females
Stage_5b_males
Stage_5c_males
Stage_6_females
Stage_6_males
sex
F
M

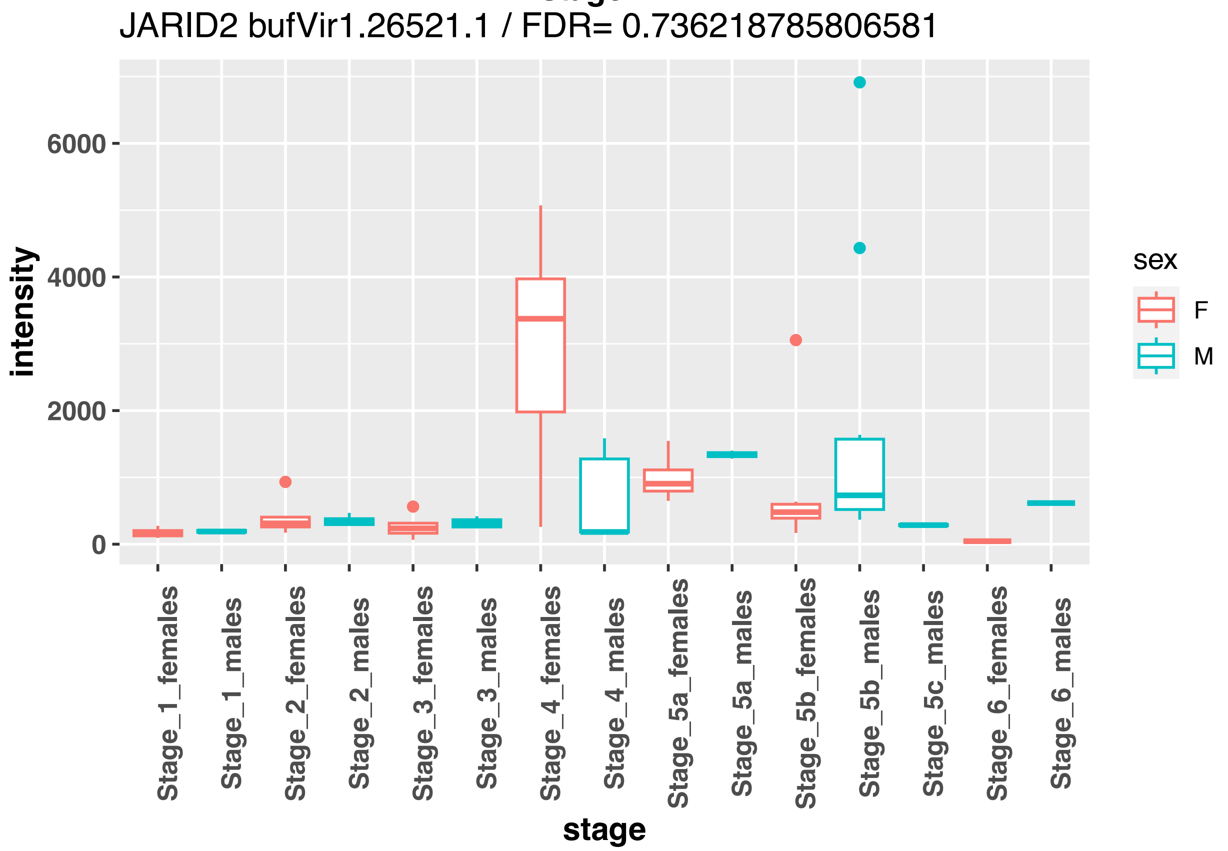
JARID2 bufVir1.26521.1 / FDR= 0.736218785806581
intensity
6000
4000
2000
0
stage
Stage_1_females
Stage_1_males
Stage_2_females
Stage_2_males
Stage_3_females
Stage_3_males
Stage_4_females
Stage_4_males
Stage_5a_females
Stage_5a_males
Stage_5b_females
Stage_5b_males
Stage_5c_males
Stage_6_females
Stage_6_males
sex
F
M

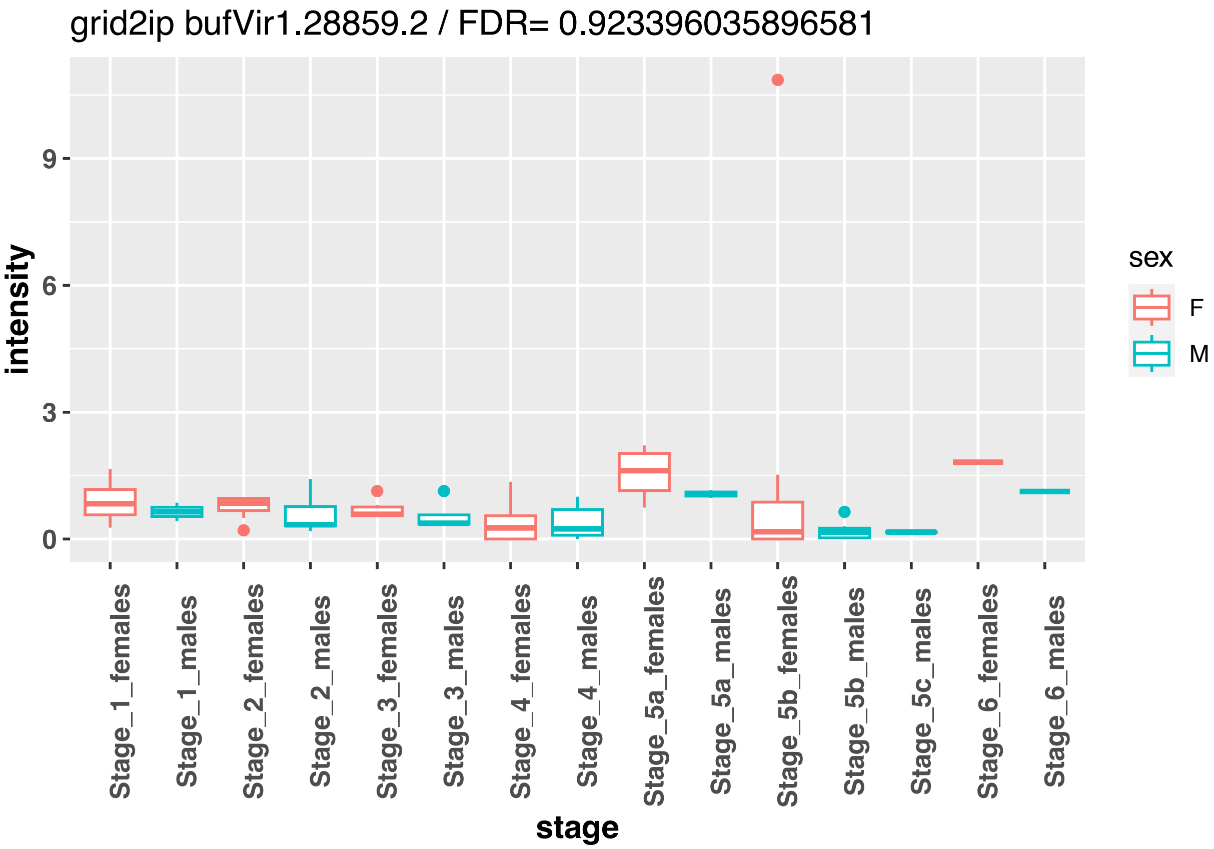
grid2ip bufVir1.28859.2 / FDR= 0.923396035896581
intensity
0
3
6
9
Stage_1_females
Stage_1_males
Stage_2_females
Stage_2_males
Stage_3_females
Stage_3_males
Stage_4_females
Stage_4_males
Stage_5a_females
Stage_5a_males
Stage_5b_females
Stage_5b_males
Stage_5c_males
Stage_6_females
Stage_6_males
stage
sex
F
M

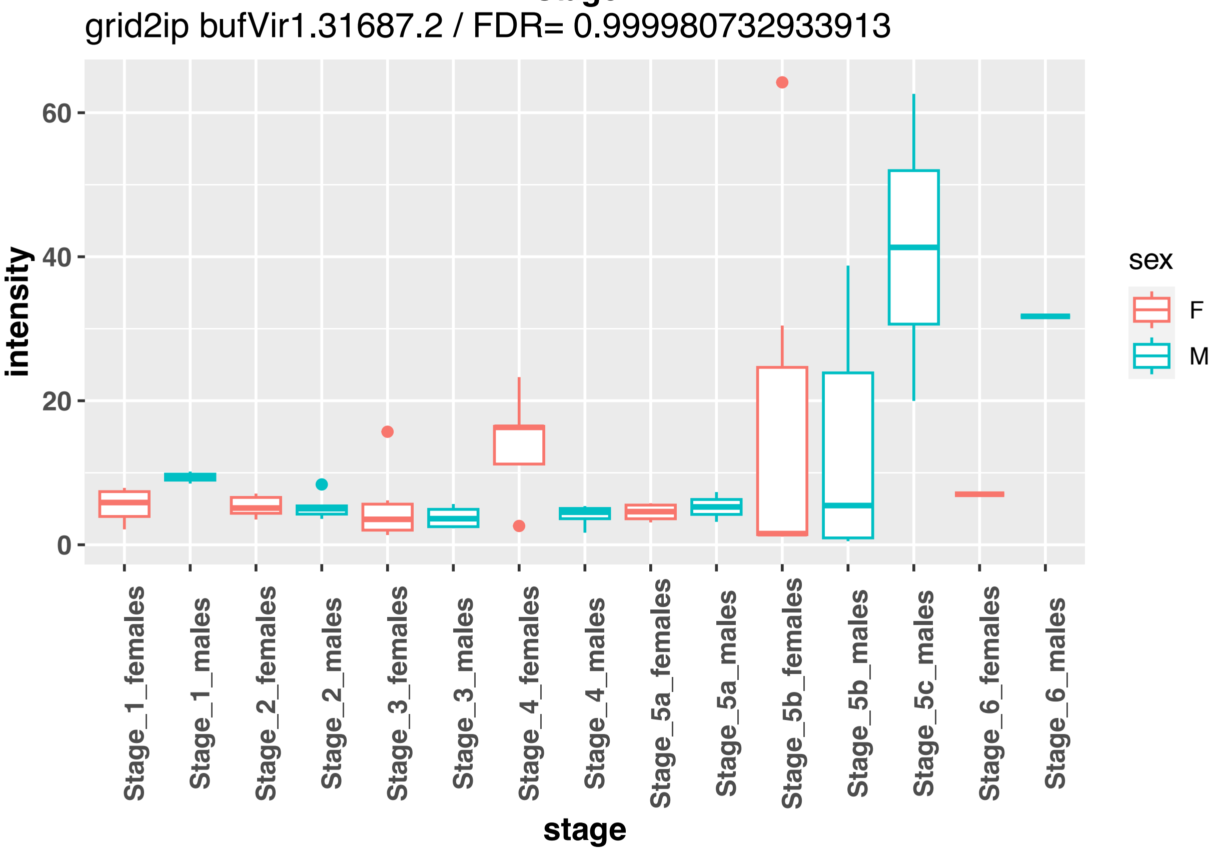
grid2ip bufVir1.31687.2 / FDR= 0.999980732933913
intensity
0
20
40
60
Stage_1_females
Stage_1_males
Stage_2_females
Stage_2_males
Stage_3_females
Stage_3_males
Stage_4_females
Stage_4_males
Stage_5a_females
Stage_5a_males
Stage_5b_females
Stage_5b_males
Stage_5c_males
Stage_6_females
Stage_6_males
stage
sex
F
M

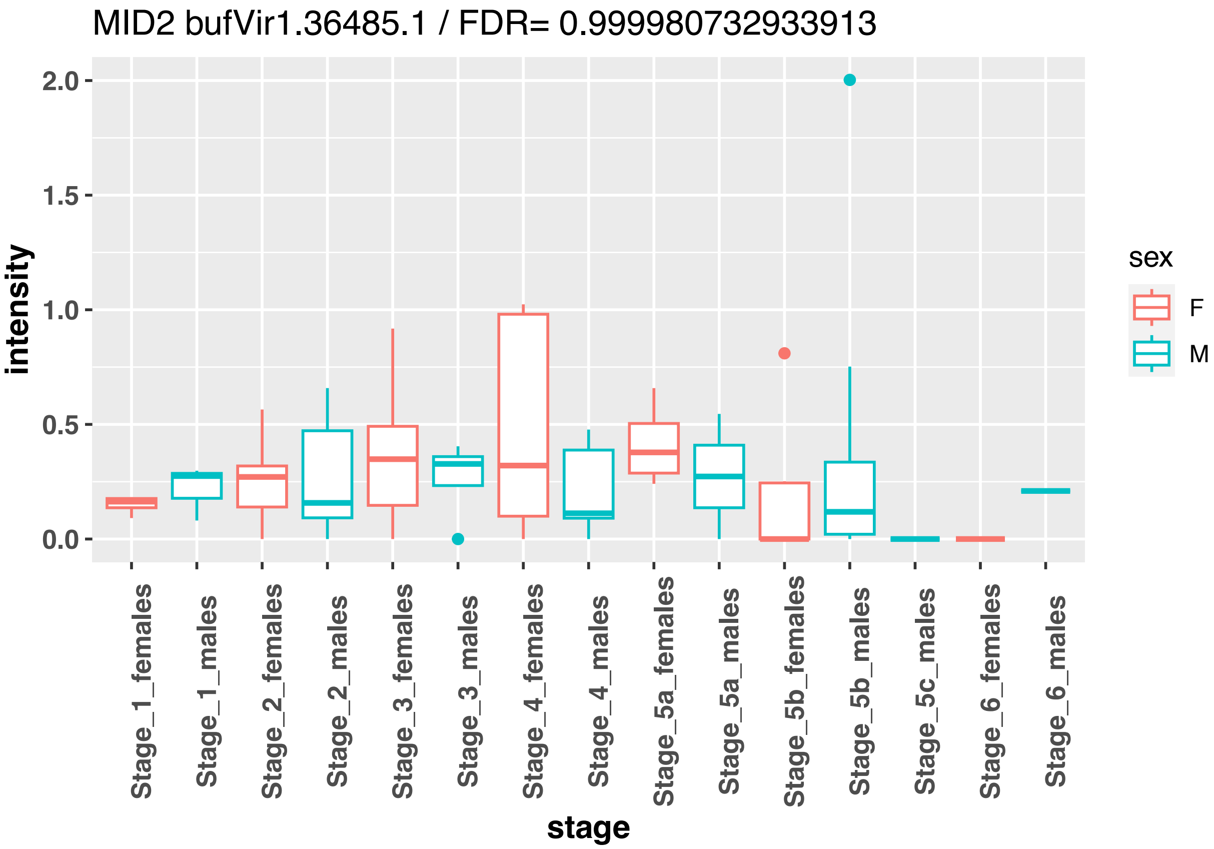
MID2 bufVir1.36485.1 / FDR= 0.999980732933913
intensity
2.0
1.5
1.0
0.5
0.0
stage
Stage_1_females
Stage_1_males
Stage_2_females
Stage_2_males
Stage_3_females
Stage_3_males
Stage_4_females
Stage_4_males
Stage_5a_females
Stage_5a_males
Stage_5b_females
Stage_5b_males
Stage_5c_males
Stage_6_females
Stage_6_males
sex
F
M

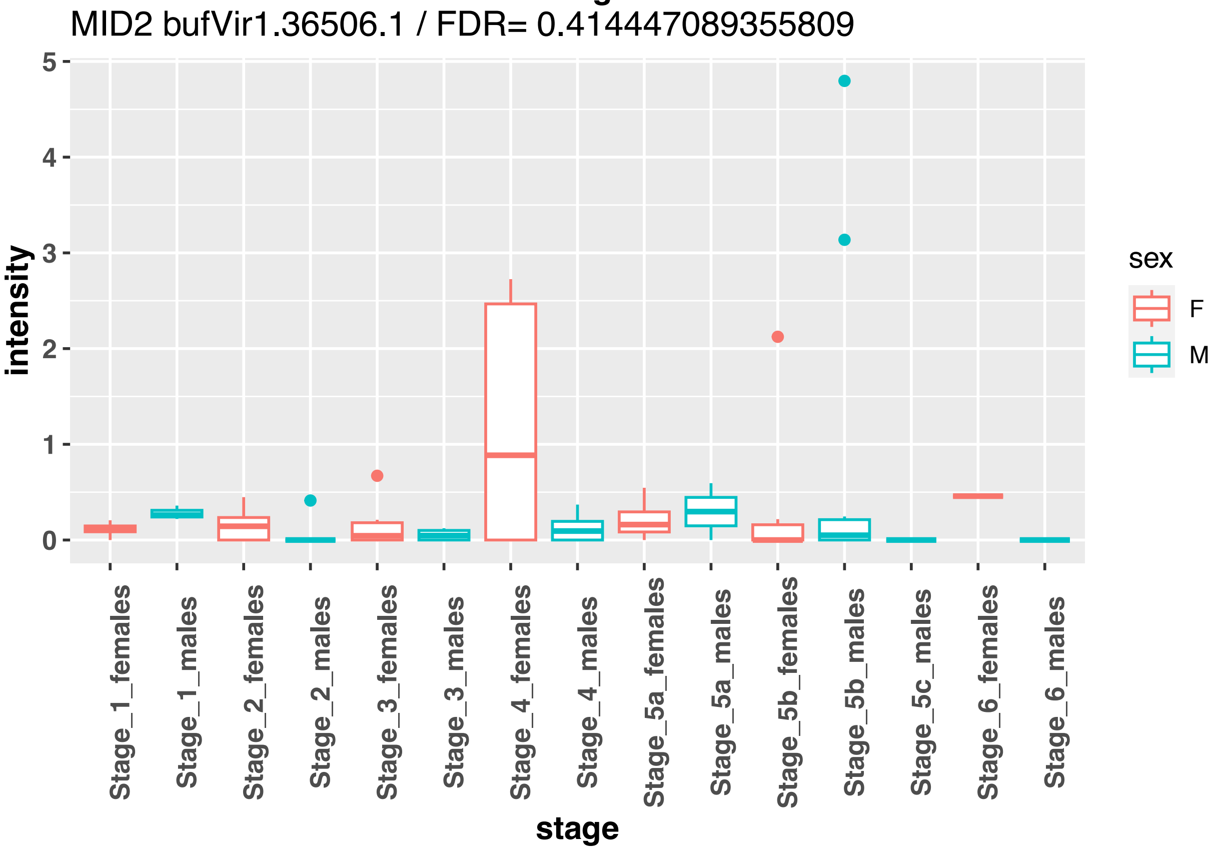
MID2 bufVir1.36506.1 / FDR= 0.414447089355809
intensity
5
4
3
2
1
0
stage
Stage_1_females
Stage_1_males
Stage_2_females
Stage_2_males
Stage_3_females
Stage_3_males
Stage_4_females
Stage_4_males
Stage_5a_females
Stage_5a_males
Stage_5b_females
Stage_5b_males
Stage_5c_males
Stage_6_females
Stage_6_males
sex
F
M

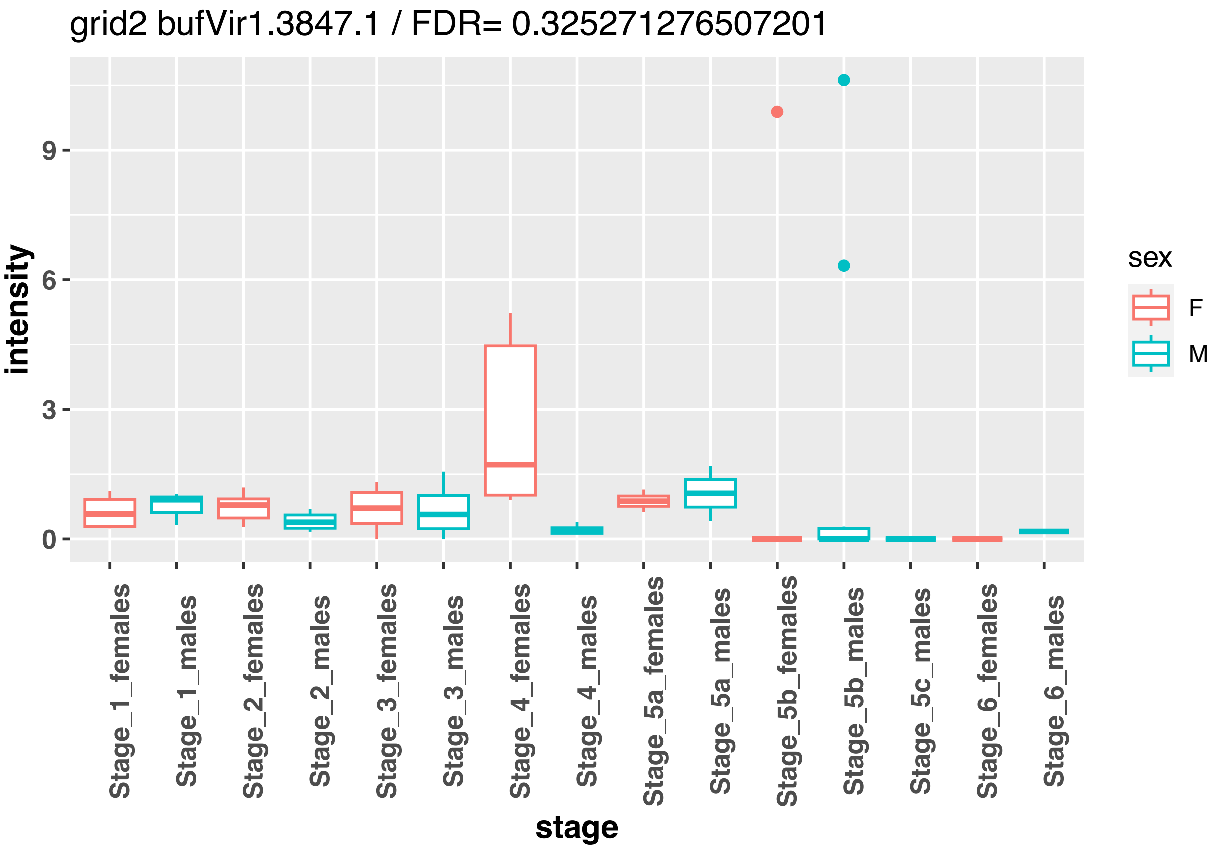

grid2 bufVir1.3847.1 / FDR= 0.325271276507201
intensity
9
6
3
0
stage
Stage_1_females
Stage_1_males
Stage_2_females
Stage_2_males
Stage_3_females
Stage_3_males
Stage_4_females
Stage_4_males
Stage_5a_females
Stage_5a_males
Stage_5b_females
Stage_5b_males
Stage_5c_males
Stage_6_females
Stage_6_males
sex
F
M


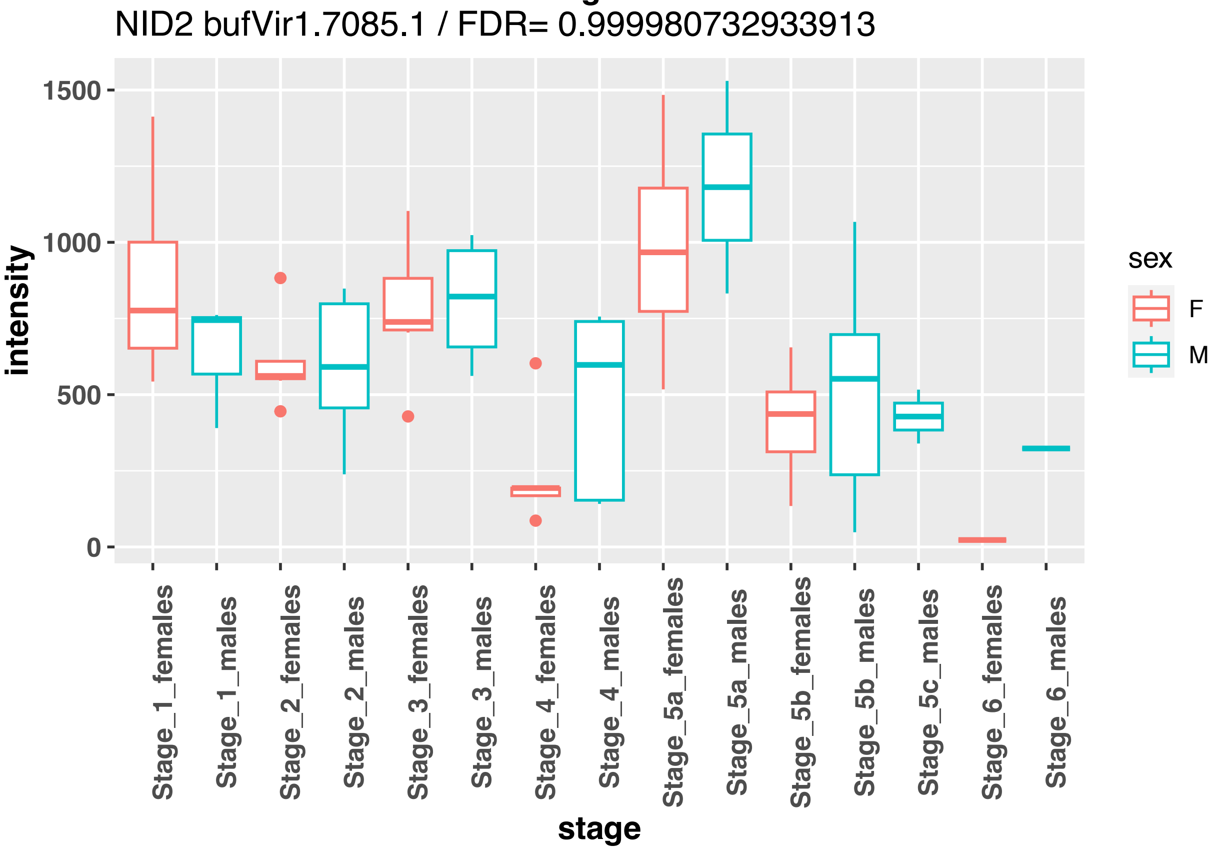

NID2 bufVir1.7085.1 / FDR= 0.999980732933913
intensity
1500
1000
500
0
stage
Stage_1_females
Stage_1_males
Stage_2_females
Stage_2_males
Stage_3_females
Stage_3_males
Stage_4_females
Stage_4_males
Stage_5a_females
Stage_5a_males
Stage_5b_females
Stage_5b_males
Stage_5c_males
Stage_6_females
Stage_6_males
sex
F
M

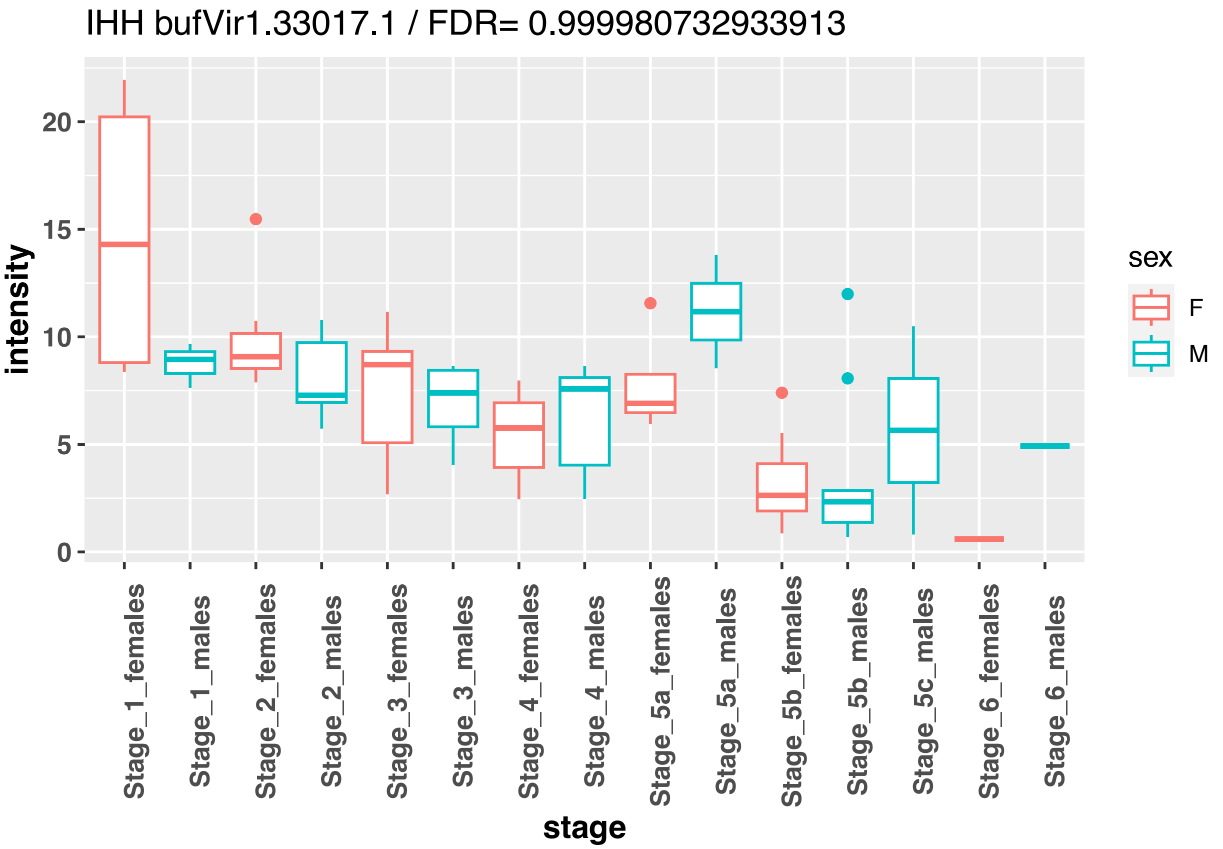
IHH bufVir1.33017.1 / FDR= 0.999980732933913
intensity
stage
sex
F
M
Stage_1_females
Stage_1_males
Stage_2_females
Stage_2_males
Stage_3_females
Stage_3_males
Stage_4_females
Stage_4_males
Stage_5a_females
Stage_5a_males
Stage_5b_females
Stage_5b_males
Stage_5c_males
Stage_6_females
Stage_6_males

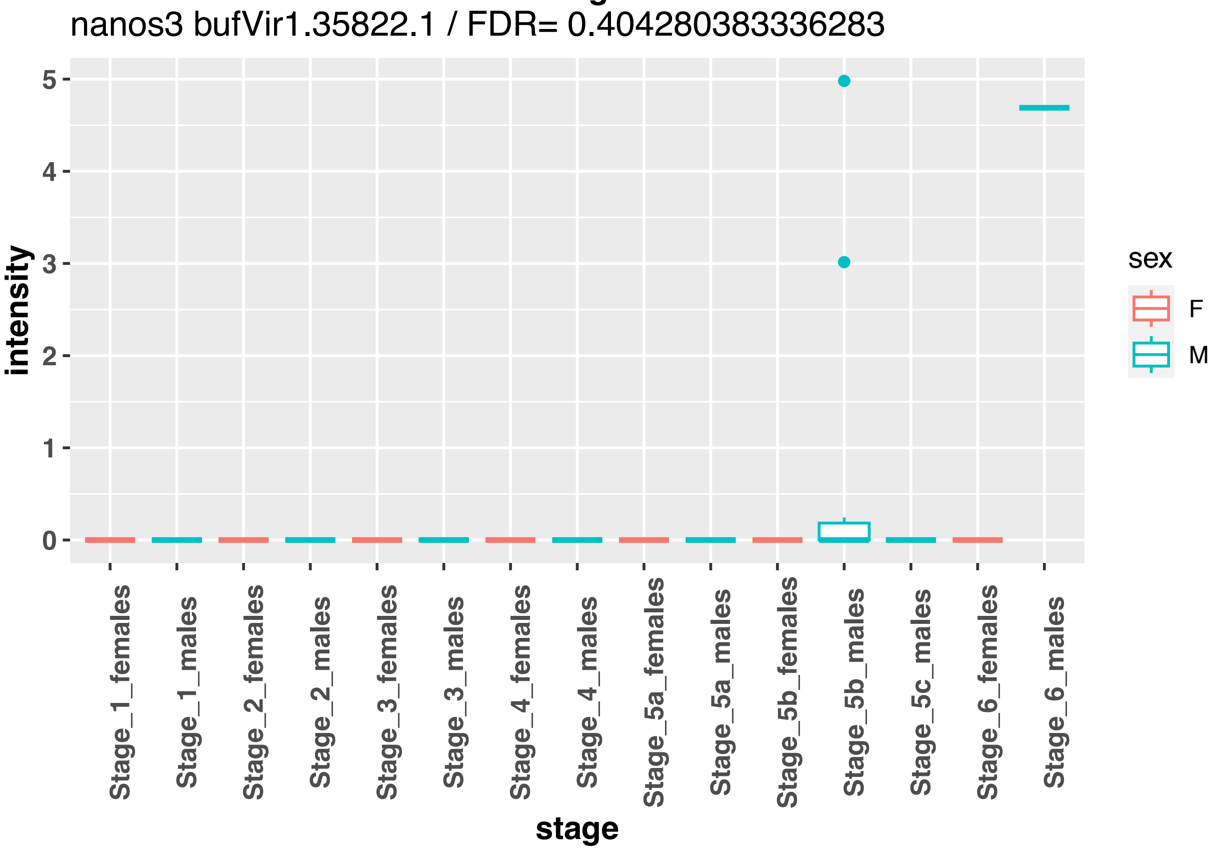
nanos3 bufVir1.35822.1 / FDR= 0.404280383336283
intensity
stage
sex
F
M
Stage_1_females
Stage_1_males
Stage_2_females
Stage_2_males
Stage_3_females
Stage_3_males
Stage_4_females
Stage_4_males
Stage_5a_females
Stage_5a_males
Stage_5b_females
Stage_5b_males
Stage_5c_males
Stage_6_females
Stage_6_males

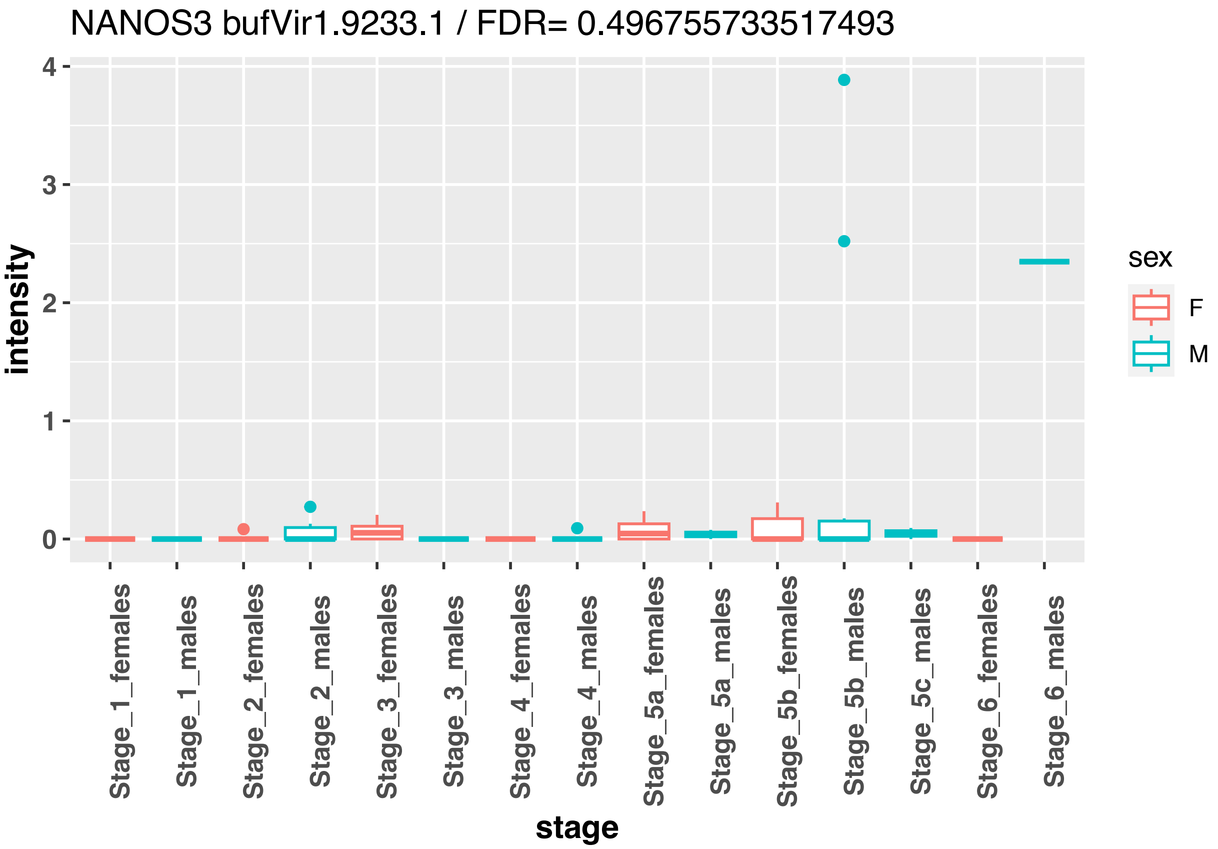
NANOS3 bufVir1.9233.1 / FDR= 0.496755733517493
intensity
0
1
2
3
4
stage
Stage_1_females
Stage_1_males
Stage_2_females
Stage_2_males
Stage_3_females
Stage_3_males
Stage_4_females
Stage_4_males
Stage_5a_females
Stage_5a_males
Stage_5b_females
Stage_5b_males
Stage_5c_males
Stage_6_females
Stage_6_males
sex
F
M

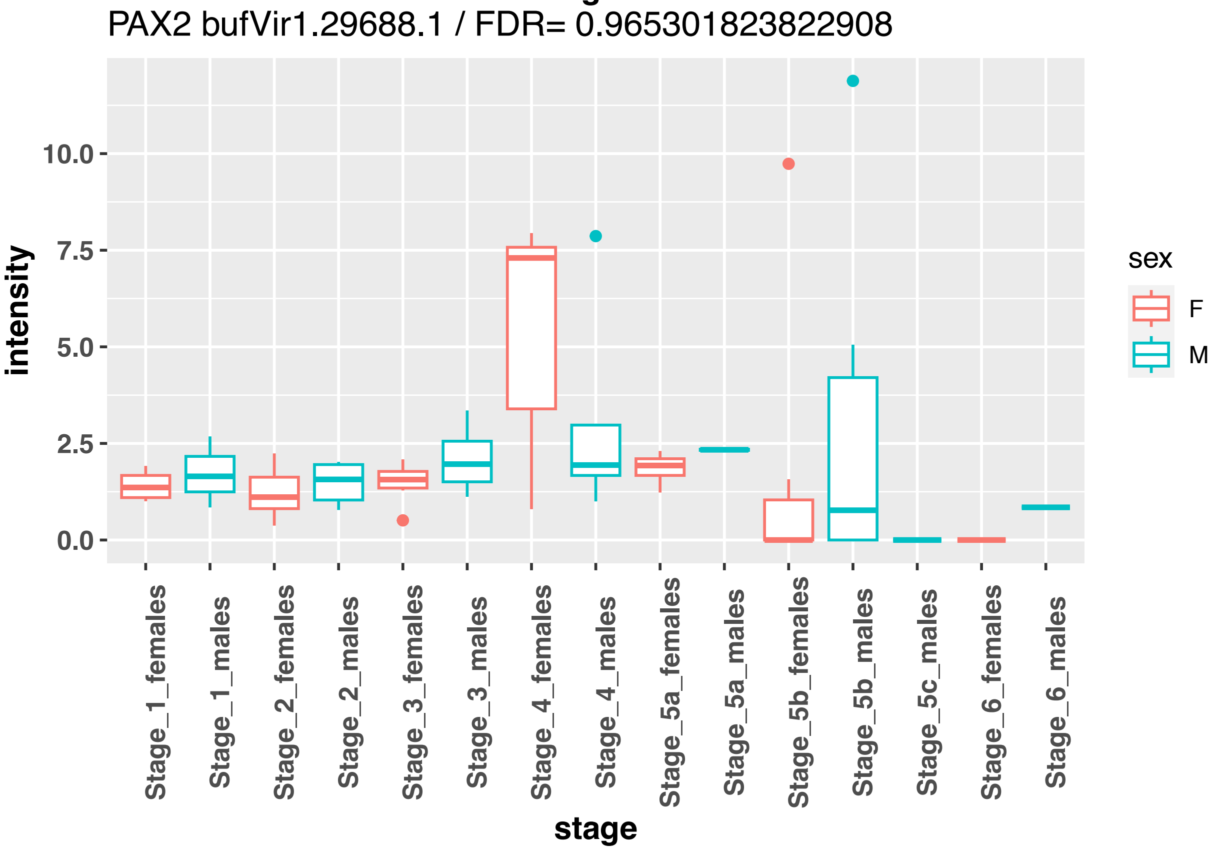
PAX2 bufVir1.29688.1 / FDR= 0.965301823822908
intensity
0.0
2.5
5.0
7.5
10.0
stage
Stage_1_females
Stage_1_males
Stage_2_females
Stage_2_males
Stage_3_females
Stage_3_males
Stage_4_females
Stage_4_males
Stage_5a_females
Stage_5a_males
Stage_5b_females
Stage_5b_males
Stage_5c_males
Stage_6_females
Stage_6_males
sex
F
M

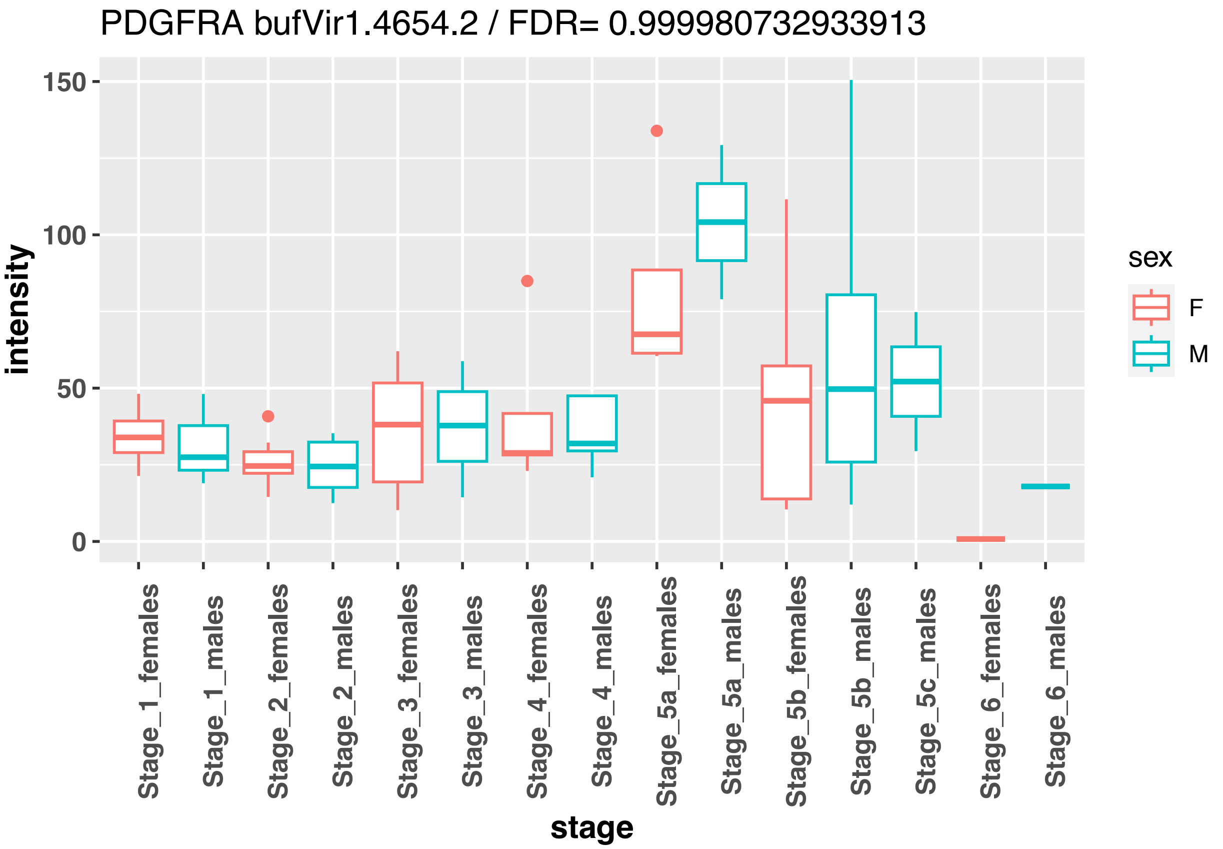
PDGFRA bufVir1.4654.2 / FDR= 0.999980732933913
intensity
150
100
50
0
stage
Stage_1_females
Stage_1_males
Stage_2_females
Stage_2_males
Stage_3_females
Stage_3_males
Stage_4_females
Stage_4_males
Stage_5a_females
Stage_5a_males
Stage_5b_females
Stage_5b_males
Stage_5c_males
Stage_6_females
Stage_6_males
sex
F
M

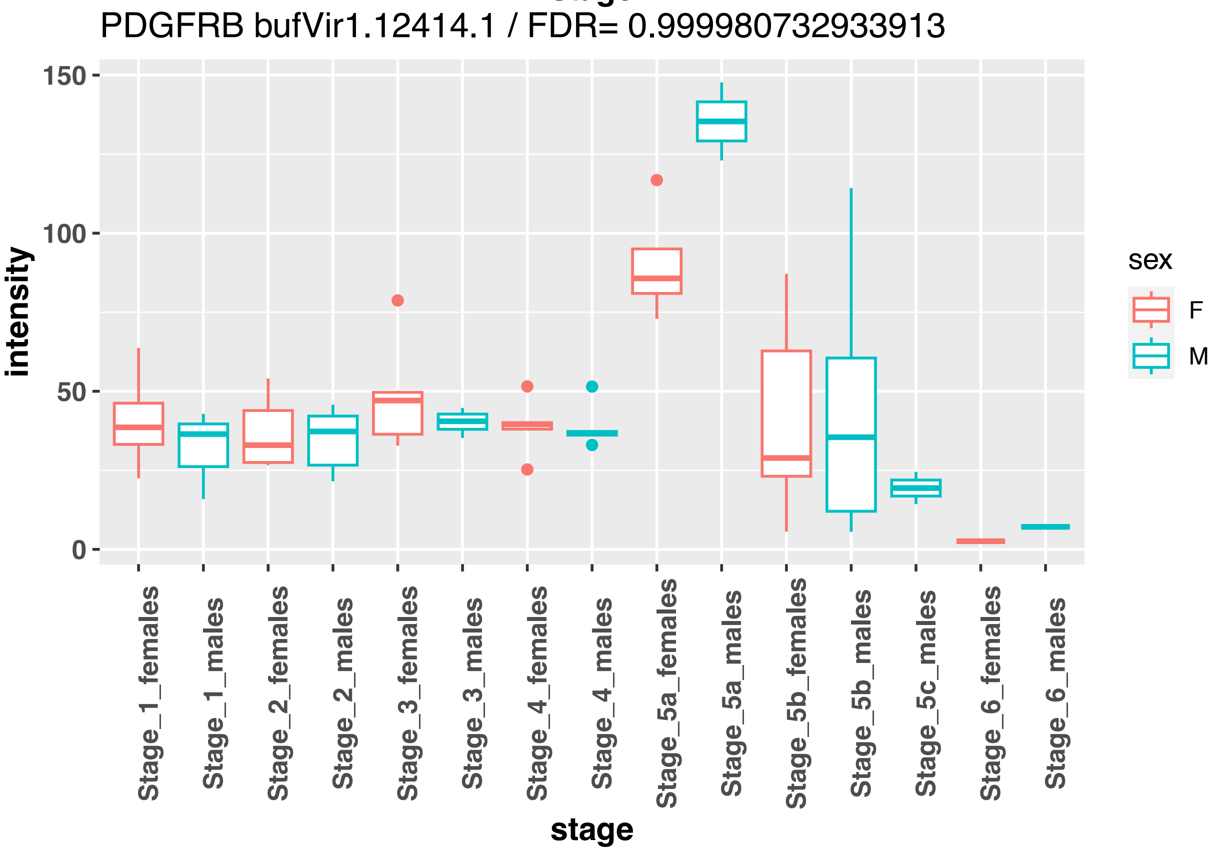
PDGFRB bufVir1.12414.1 / FDR= 0.999980732933913
intensity
150
100
50
0
stage
Stage_1_females
Stage_1_males
Stage_2_females
Stage_2_males
Stage_3_females
Stage_3_males
Stage_4_females
Stage_4_males
Stage_5a_females
Stage_5a_males
Stage_5b_females
Stage_5b_males
Stage_5c_males
Stage_6_females
Stage_6_males
sex
F
M

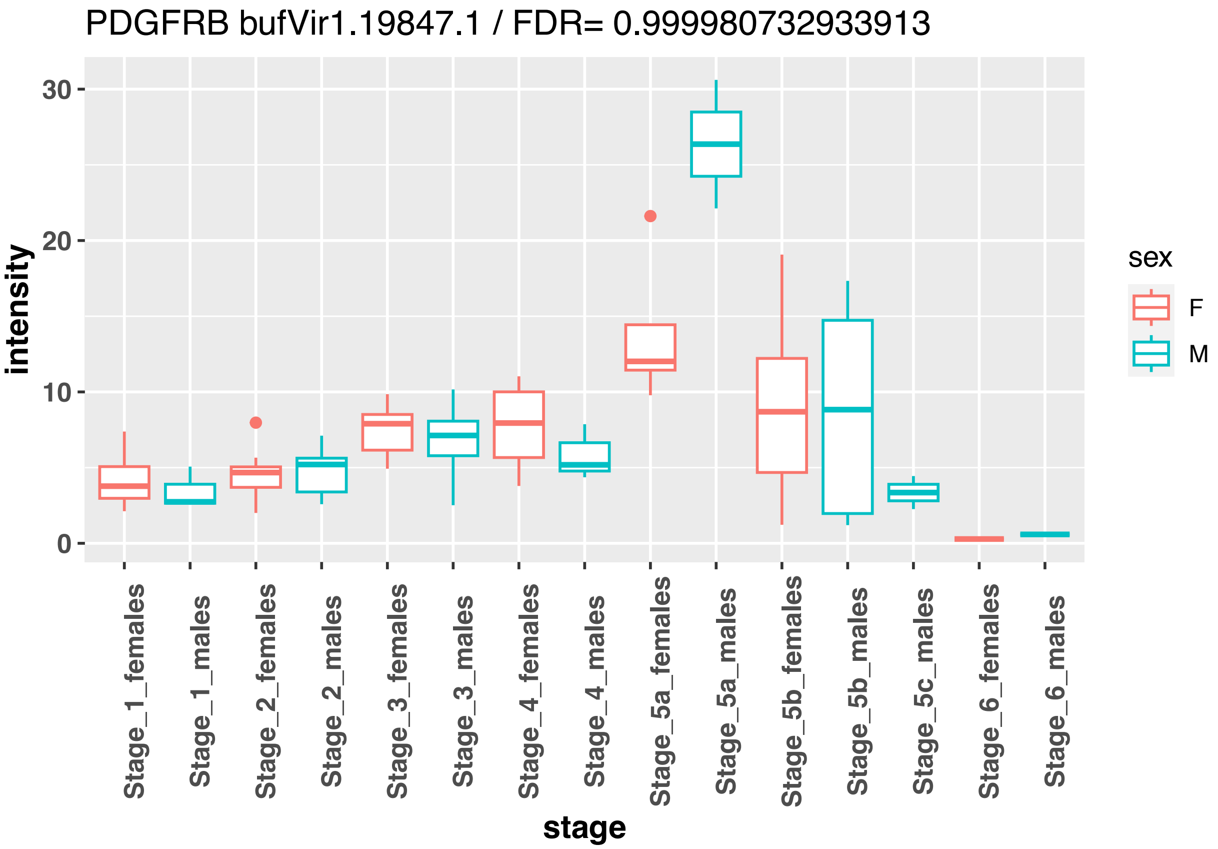
PDGFRB bufVir1.19847.1 / FDR= 0.999980732933913
intensity
30
20
10
0
stage
Stage_1_females
Stage_1_males
Stage_2_females
Stage_2_males
Stage_3_females
Stage_3_males
Stage_4_females
Stage_4_males
Stage_5a_females
Stage_5a_males
Stage_5b_females
Stage_5b_males
Stage_5c_males
Stage_6_females
Stage_6_males
sex
F
M

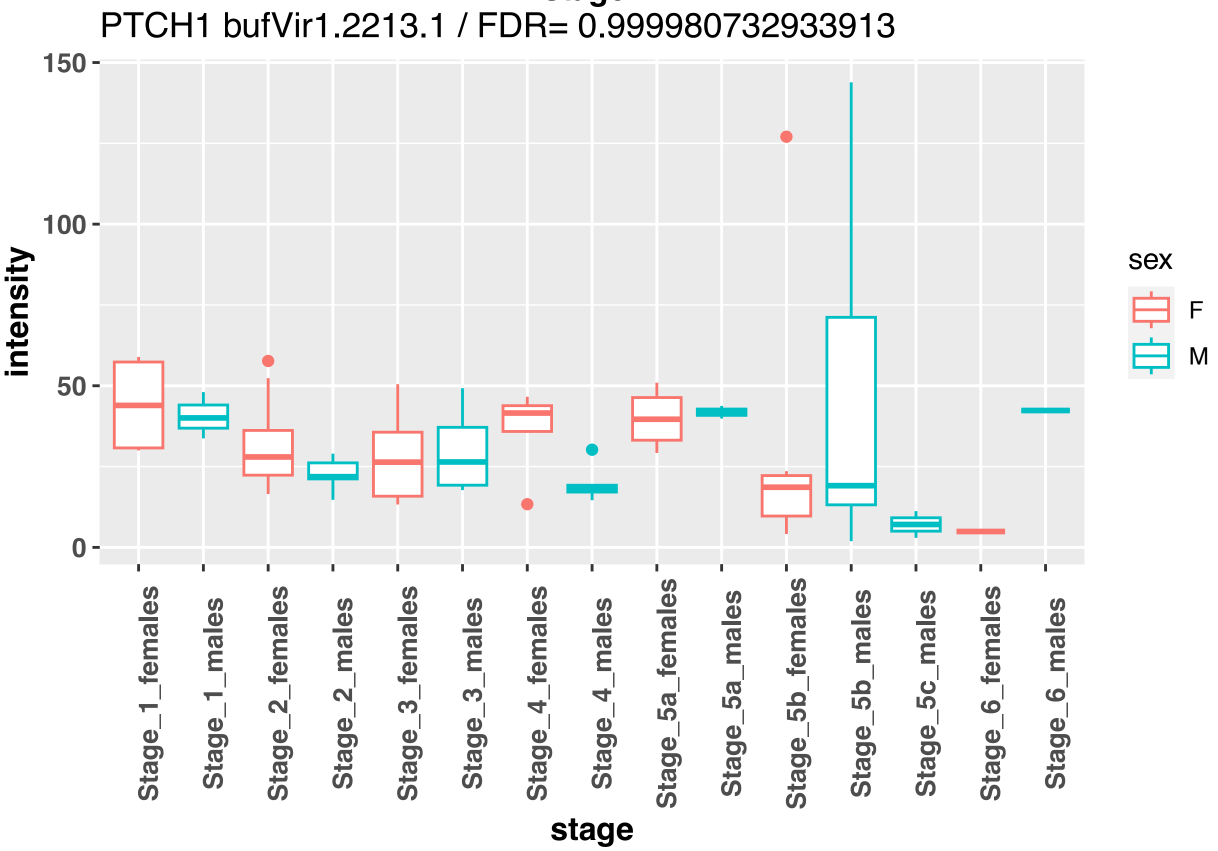
PTCH1 bufVir1.2213.1 / FDR= 0.999980732933913
intensity
150
100
50
0
stage
Stage_1_females
Stage_1_males
Stage_2_females
Stage_2_males
Stage_3_females
Stage_3_males
Stage_4_females
Stage_4_males
Stage_5a_females
Stage_5a_males
Stage_5b_females
Stage_5b_males
Stage_5c_males
Stage_6_females
Stage_6_males
sex
F
M

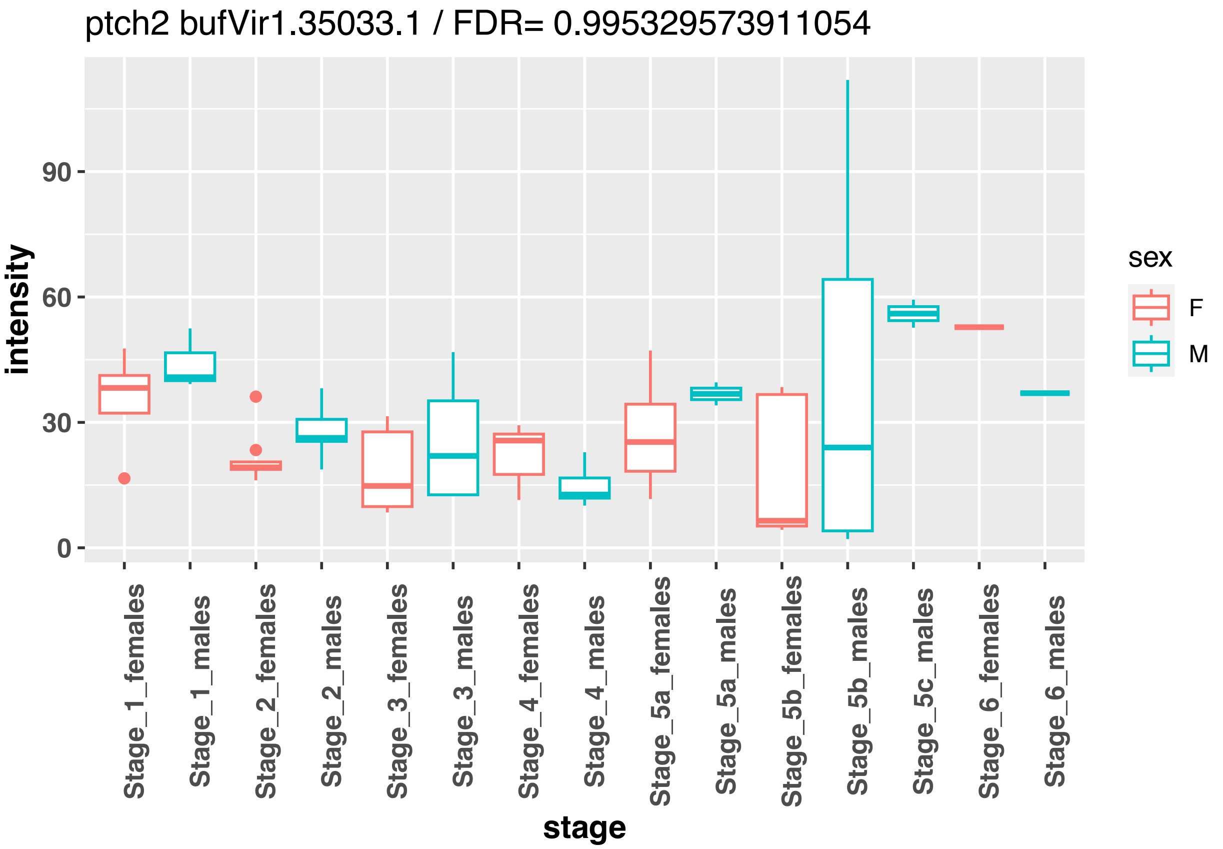

ptch2 bufVir1.35033.1 / FDR= 0.995329573911054
intensity
90
60
30
0
Stage_1_females
Stage_1_males
Stage_2_females
Stage_2_males
Stage_3_females
Stage_3_males
Stage_4_females
Stage_4_males
Stage_5a_females
Stage_5a_males
Stage_5b_females
Stage_5b_males
Stage_5c_males
Stage_6_females
Stage_6_males
stage
sex
F
M


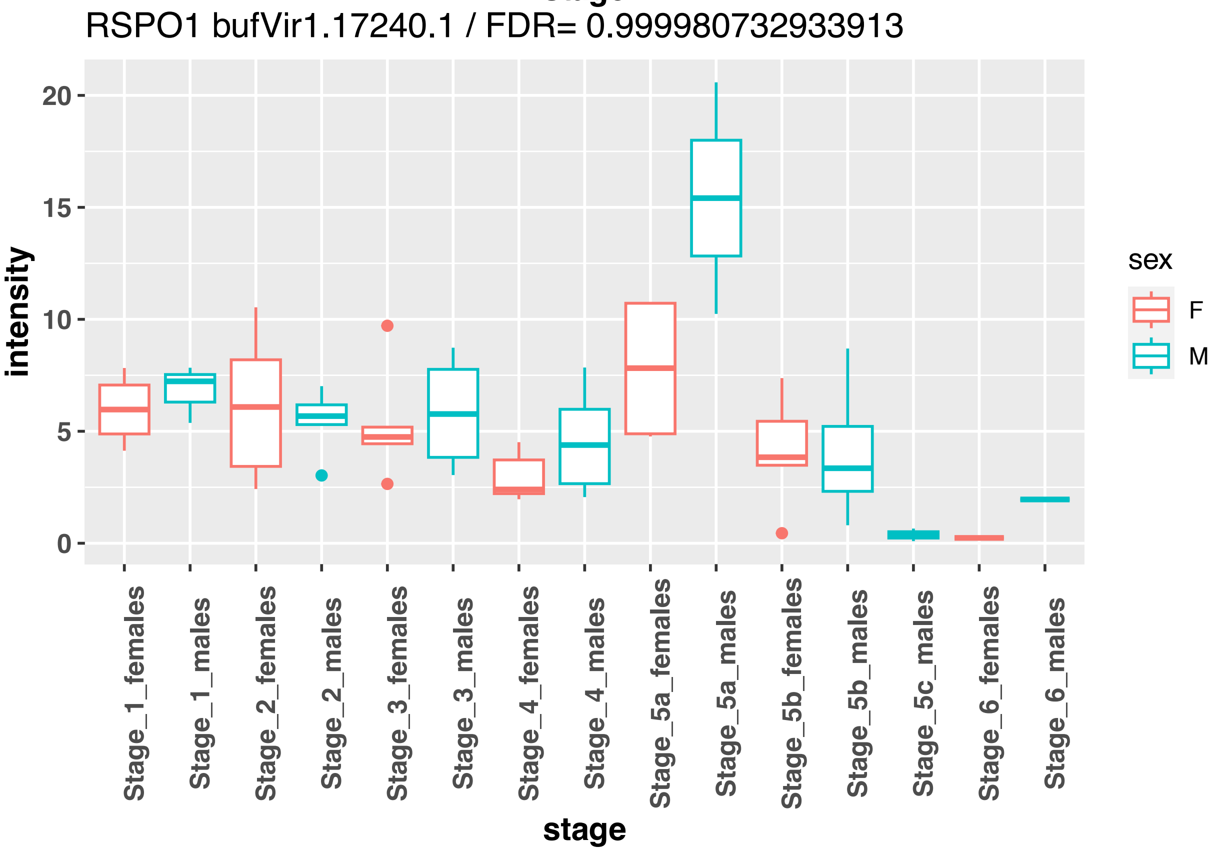

RSPO1 bufVir1.17240.1 / FDR= 0.999980732933913
intensity
20
15
10
5
0
Stage_1_females
Stage_1_males
Stage_2_females
Stage_2_males
Stage_3_females
Stage_3_males
Stage_4_females
Stage_4_males
Stage_5a_females
Stage_5a_males
Stage_5b_females
Stage_5b_males
Stage_5c_males
Stage_6_females
Stage_6_males
stage
sex
F
M

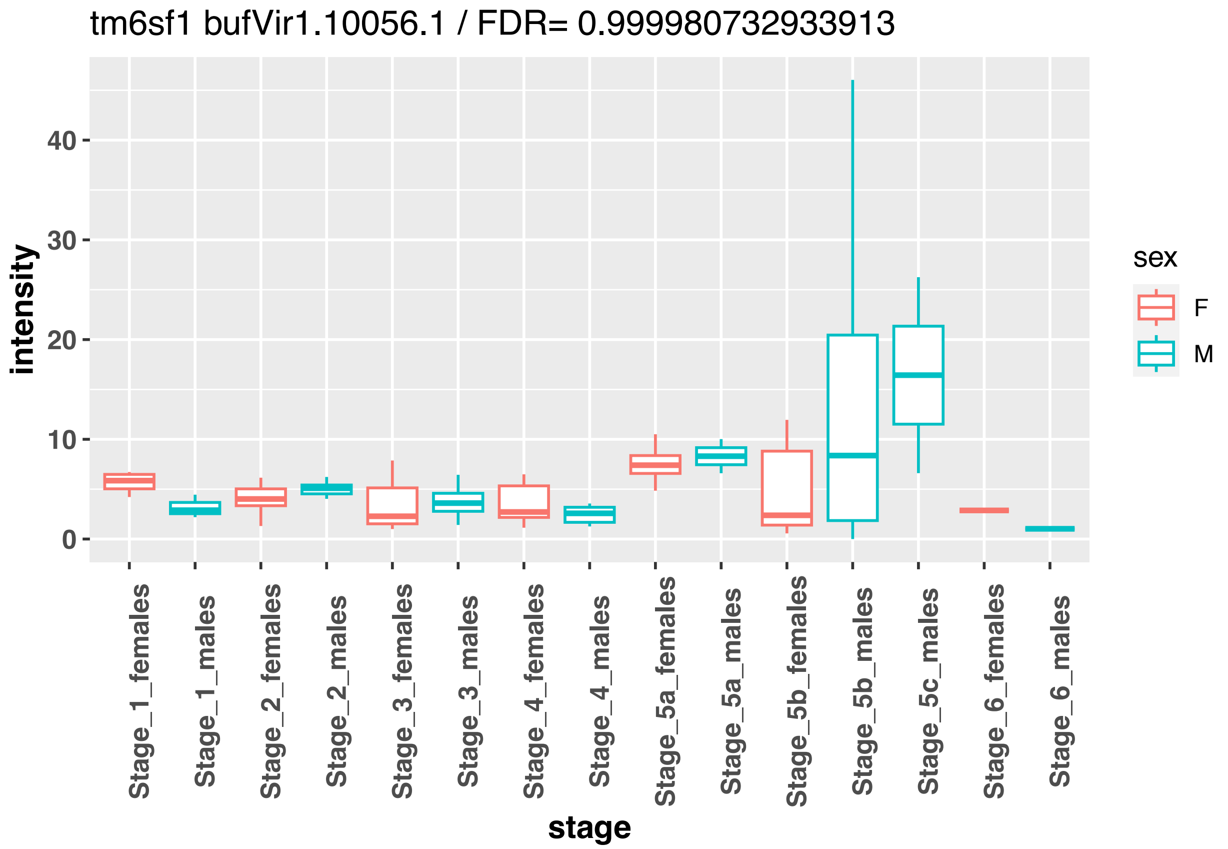
tm6sf1 bufVir1.10056.1 / FDR= 0.999980732933913
intensity
0
10
20
30
40
sex
F
M
Stage_1_females
Stage_1_males
Stage_2_females
Stage_2_males
Stage_3_females
Stage_3_males
Stage_4_females
Stage_4_males
Stage_5a_females
Stage_5a_males
Stage_5b_females
Stage_5b_males
Stage_5c_males
Stage_6_females
Stage_6_males
stage

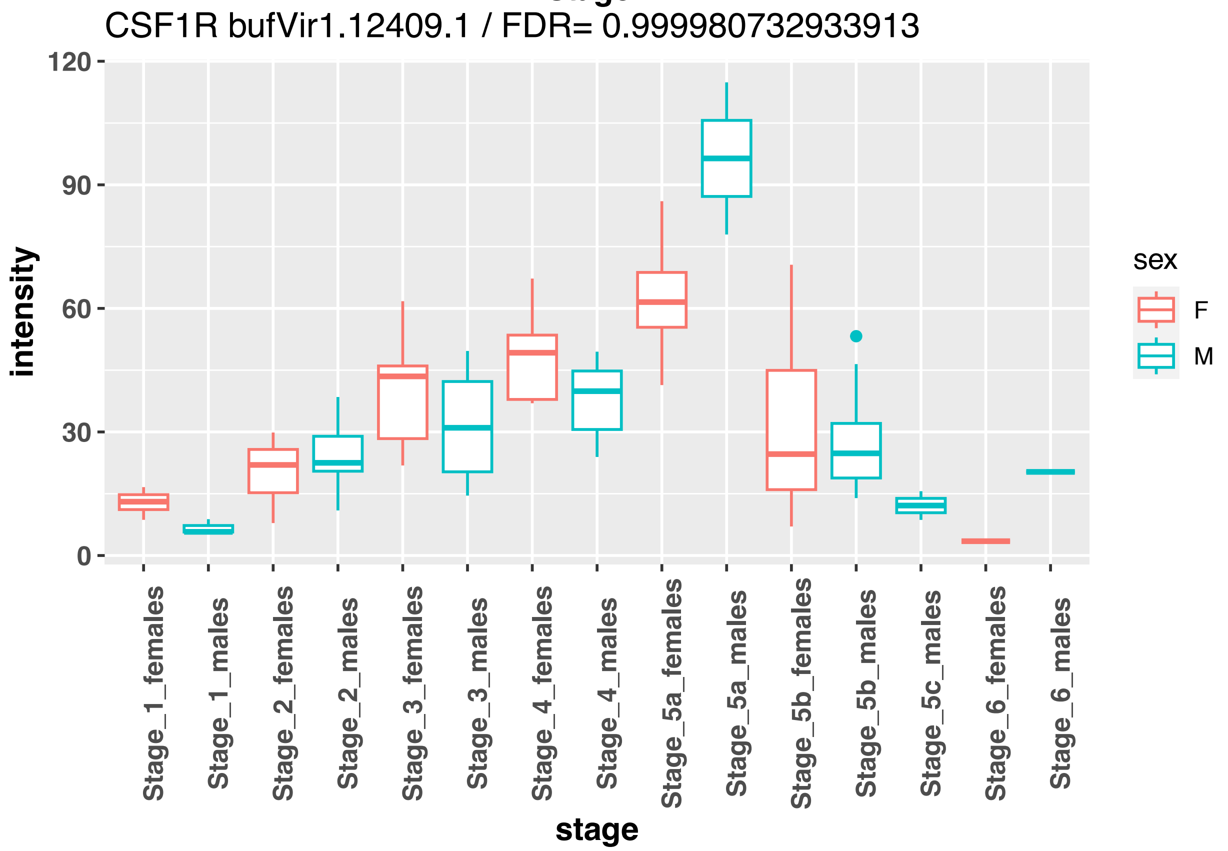
CSF1R bufVir1.12409.1 / FDR= 0.999980732933913
intensity
0
30
60
90
120
sex
F
M
Stage_1_females
Stage_1_males
Stage_2_females
Stage_2_males
Stage_3_females
Stage_3_males
Stage_4_females
Stage_4_males
Stage_5a_females
Stage_5a_males
Stage_5b_females
Stage_5b_males
Stage_5c_males
Stage_6_females
Stage_6_males
stage

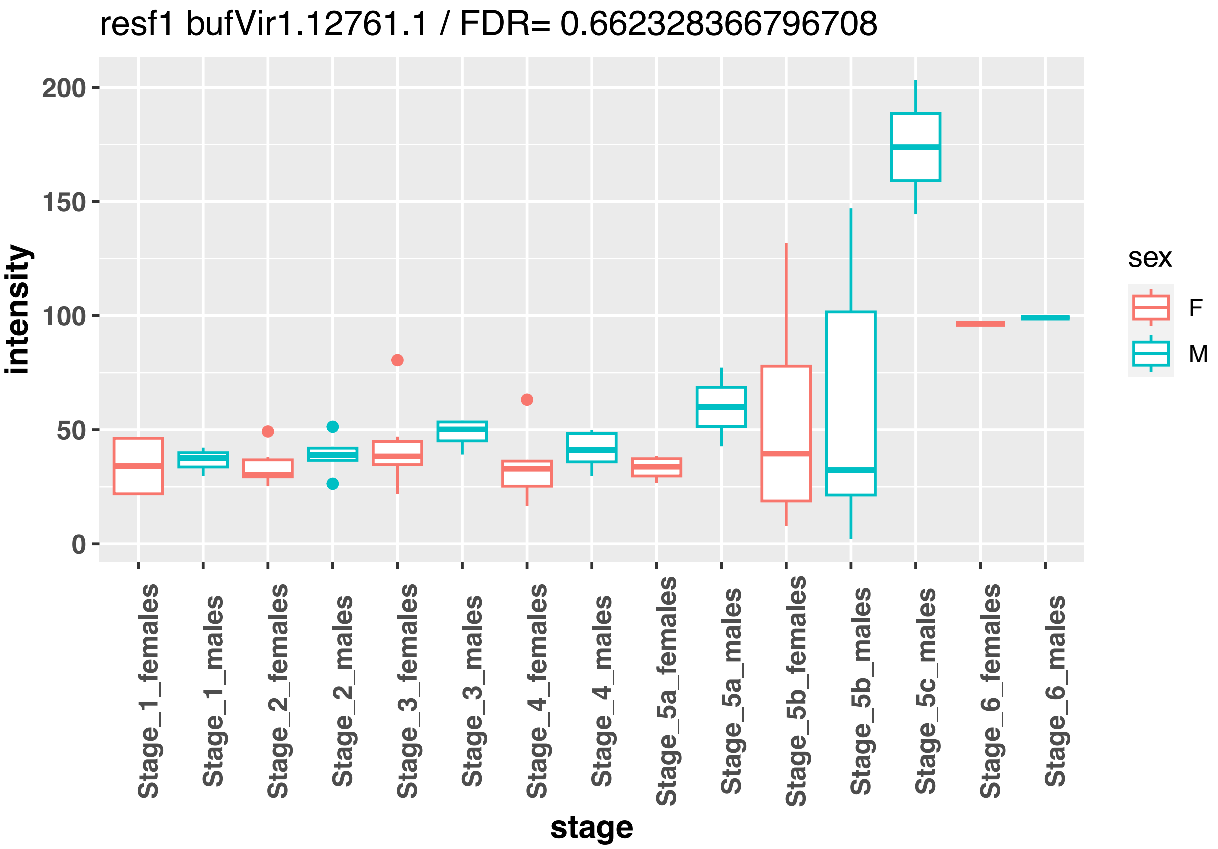

resf1 bufVir1.12761.1 / FDR= 0.662328366796708
intensity
0
50
100
150
200
stage
Stage_1_females
Stage_1_males
Stage_2_females
Stage_2_males
Stage_3_females
Stage_3_males
Stage_4_females
Stage_4_males
Stage_5a_females
Stage_5a_males
Stage_5b_females
Stage_5b_males
Stage_5c_males
Stage_6_females
Stage_6_males
sex
F
M


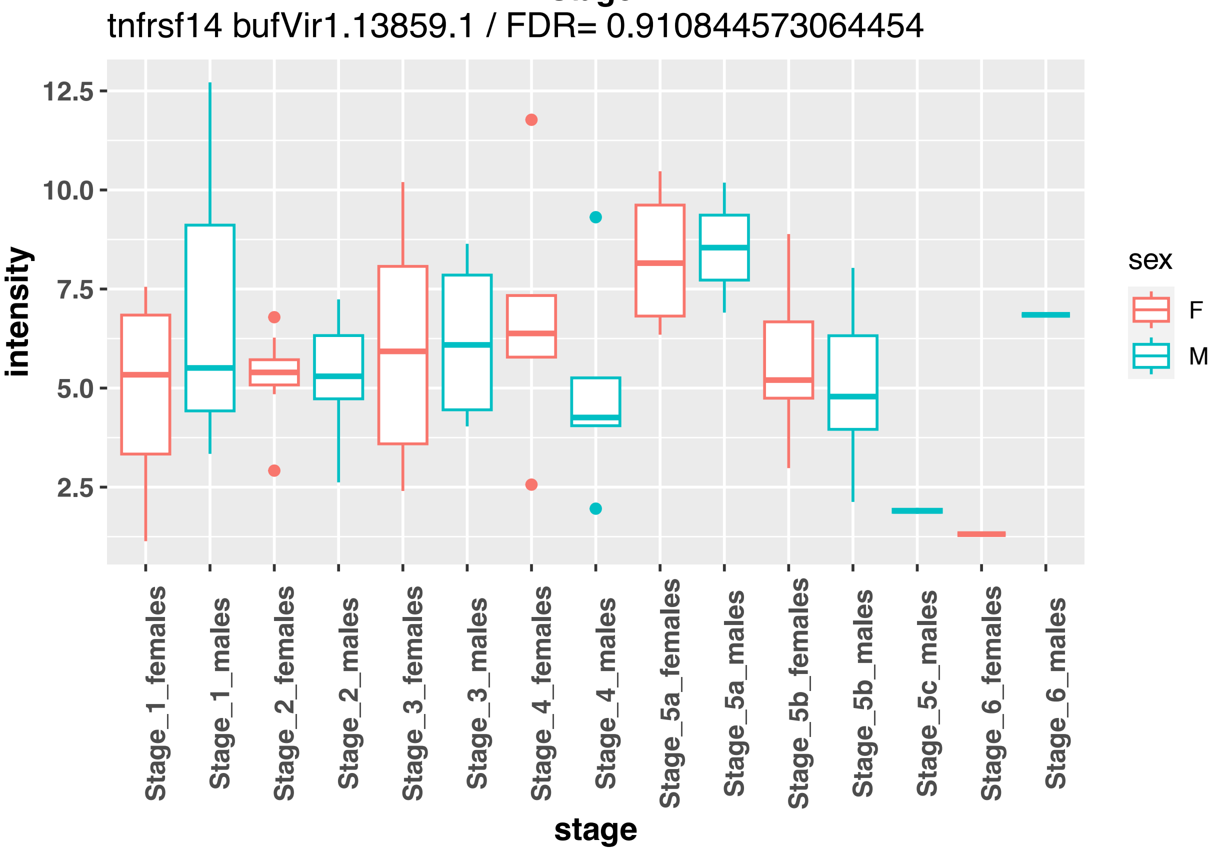

tnfrsf14 bufVir1.13859.1 / FDR= 0.910844573064454
intensity
2.5
5.0
7.5
10.0
12.5
stage
Stage_1_females
Stage_1_males
Stage_2_females
Stage_2_males
Stage_3_females
Stage_3_males
Stage_4_females
Stage_4_males
Stage_5a_females
Stage_5a_males
Stage_5b_females
Stage_5b_males
Stage_5c_males
Stage_6_females
Stage_6_males
sex
F
M

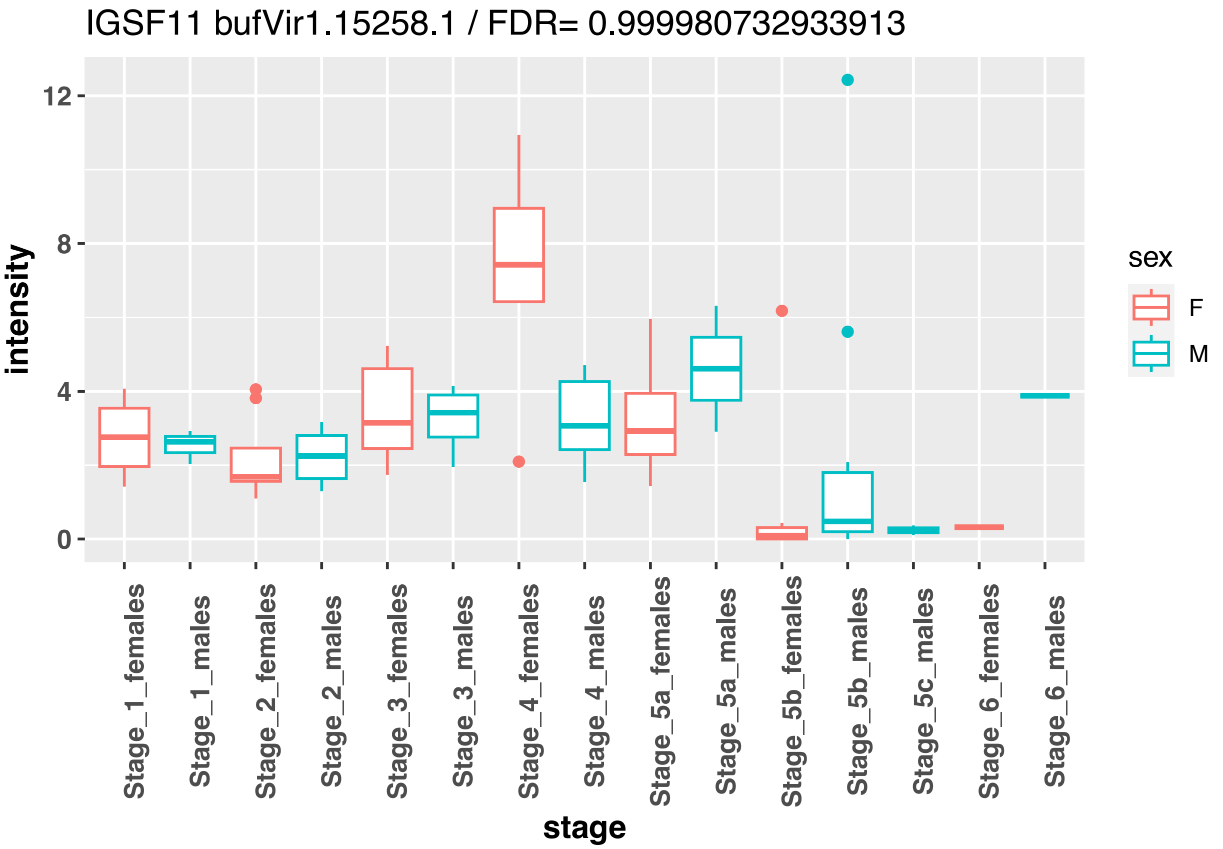
IGSF11 bufVir1.15258.1 / FDR= 0.999980732933913
intensity
12
8
4
0
stage
Stage_1_females
Stage_1_males
Stage_2_females
Stage_2_males
Stage_3_females
Stage_3_males
Stage_4_females
Stage_4_males
Stage_5a_females
Stage_5a_males
Stage_5b_females
Stage_5b_males
Stage_5c_males
Stage_6_females
Stage_6_males
sex
F
M

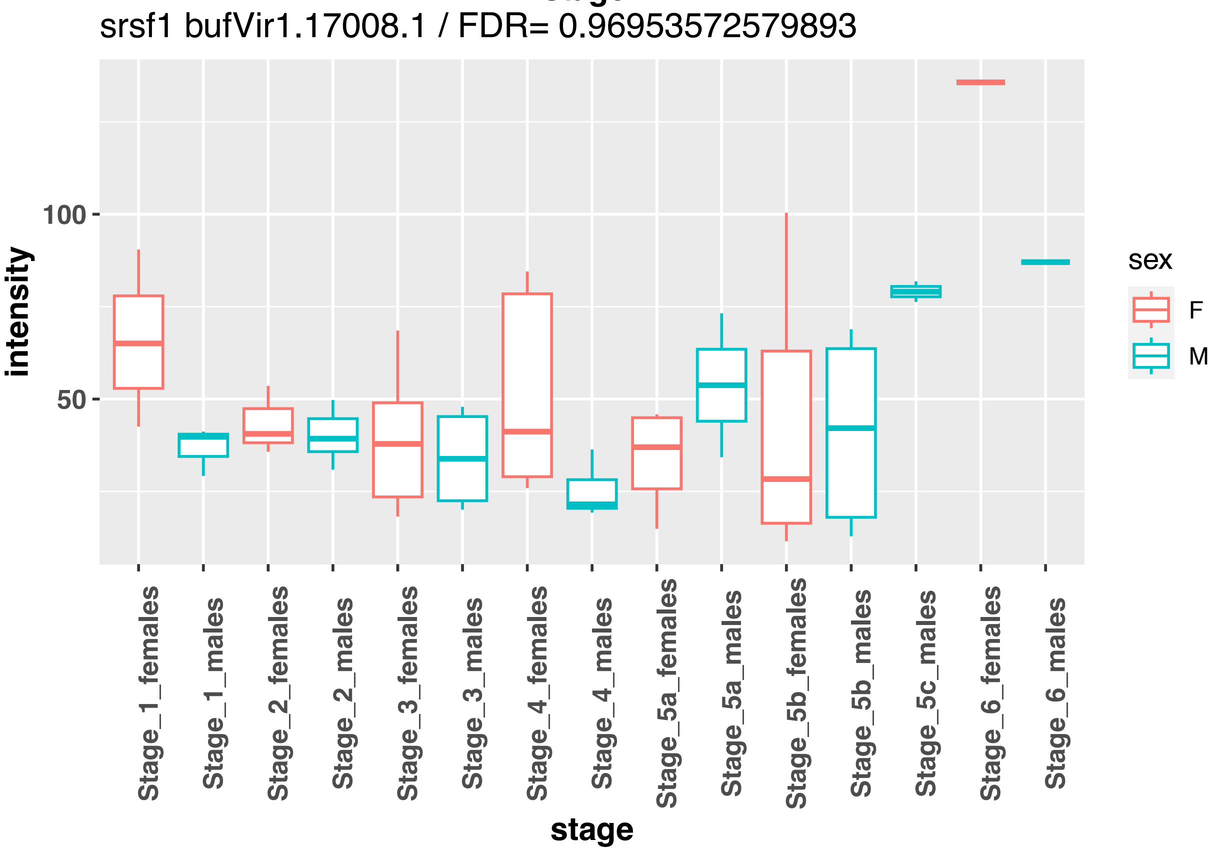
srsf1 bufVir1.17008.1 / FDR= 0.96953572579893
intensity
100
50
stage
Stage_1_females
Stage_1_males
Stage_2_females
Stage_2_males
Stage_3_females
Stage_3_males
Stage_4_females
Stage_4_males
Stage_5a_females
Stage_5a_males
Stage_5b_females
Stage_5b_males
Stage_5c_males
Stage_6_females
Stage_6_males
sex
F
M

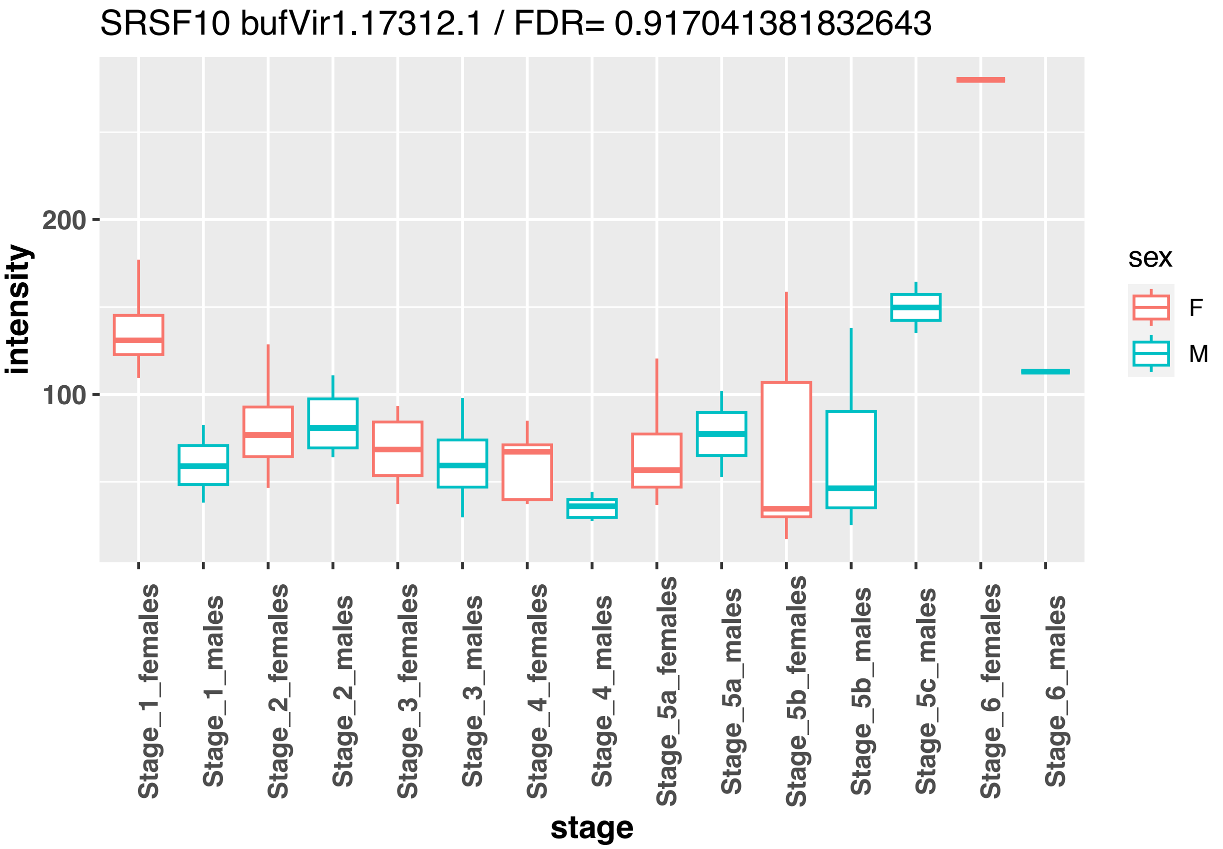
SRSF10 bufVir1.17312.1 / FDR= 0.917041381832643
intensity
200
100
stage
Stage_1_females
Stage_1_males
Stage_2_females
Stage_2_males
Stage_3_females
Stage_3_males
Stage_4_females
Stage_4_males
Stage_5a_females
Stage_5a_males
Stage_5b_females
Stage_5b_males
Stage_5c_males
Stage_6_females
Stage_6_males
sex
F
M

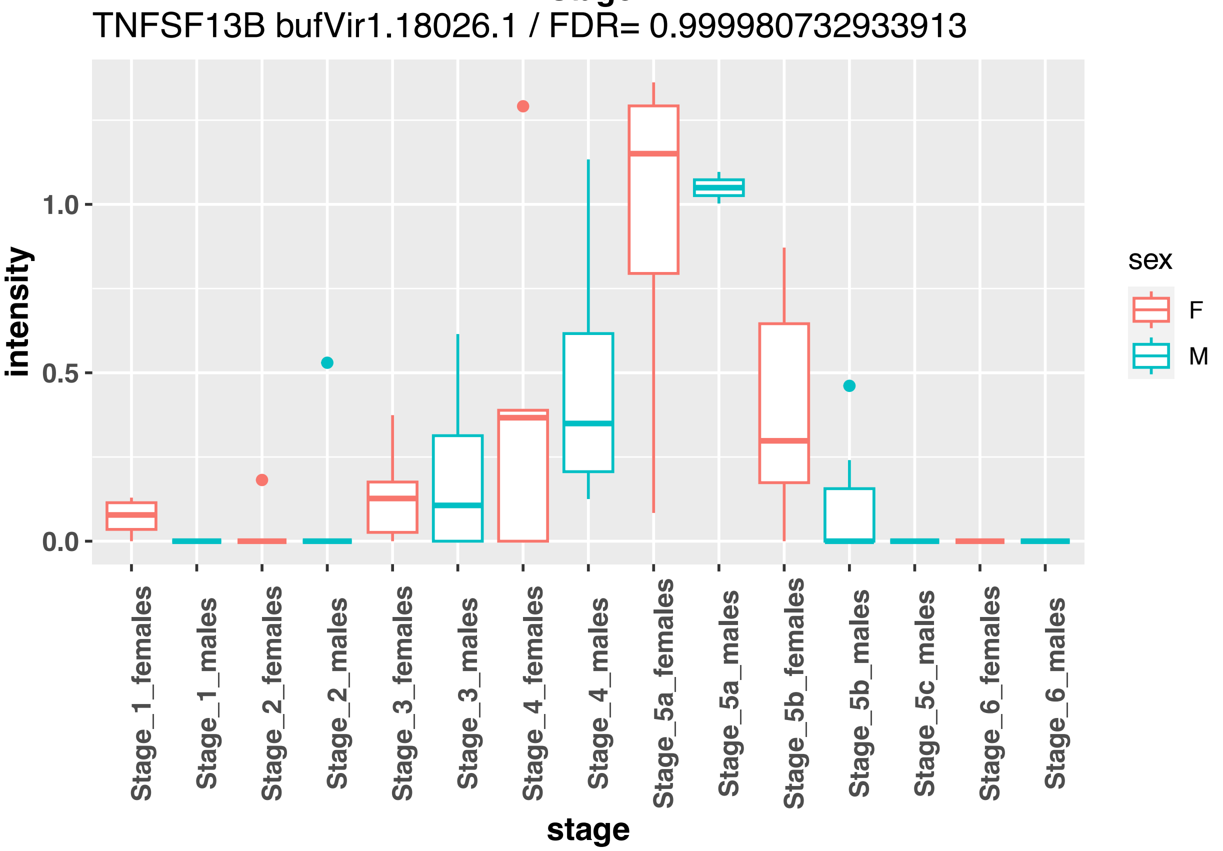
TNFSF13B bufVir1.18026.1 / FDR= 0.999980732933913
intensity
1.0
0.5
0.0
stage
Stage_1_females
Stage_1_males
Stage_2_females
Stage_2_males
Stage_3_females
Stage_3_males
Stage_4_females
Stage_4_males
Stage_5a_females
Stage_5a_males
Stage_5b_females
Stage_5b_males
Stage_5c_males
Stage_6_females
Stage_6_males
sex
F
M

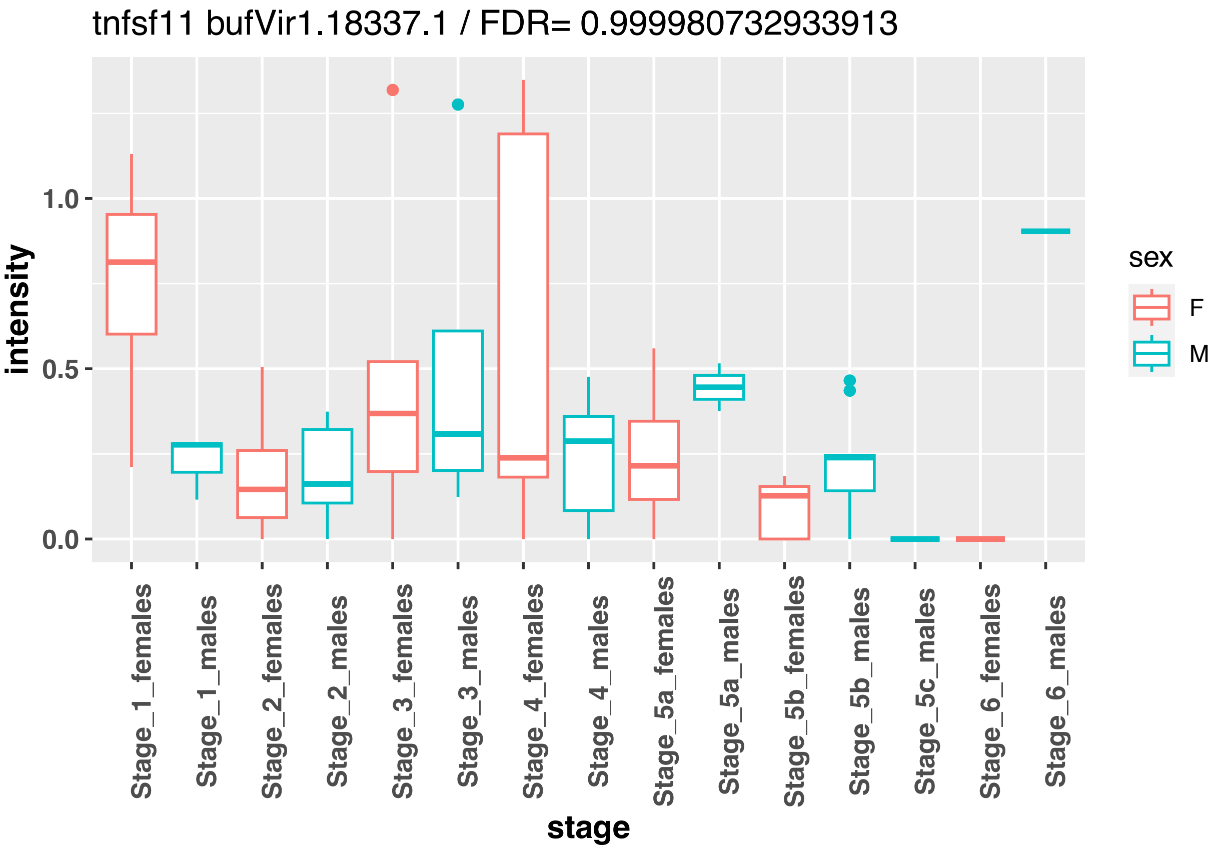

tnfsf11 bufVir1.18337.1 / FDR= 0.999980732933913
intensity
1.0
0.5
0.0
stage
Stage_1_females
Stage_1_males
Stage_2_females
Stage_2_males
Stage_3_females
Stage_3_males
Stage_4_females
Stage_4_males
Stage_5a_females
Stage_5a_males
Stage_5b_females
Stage_5b_males
Stage_5c_males
Stage_6_females
Stage_6_males
sex
F
M


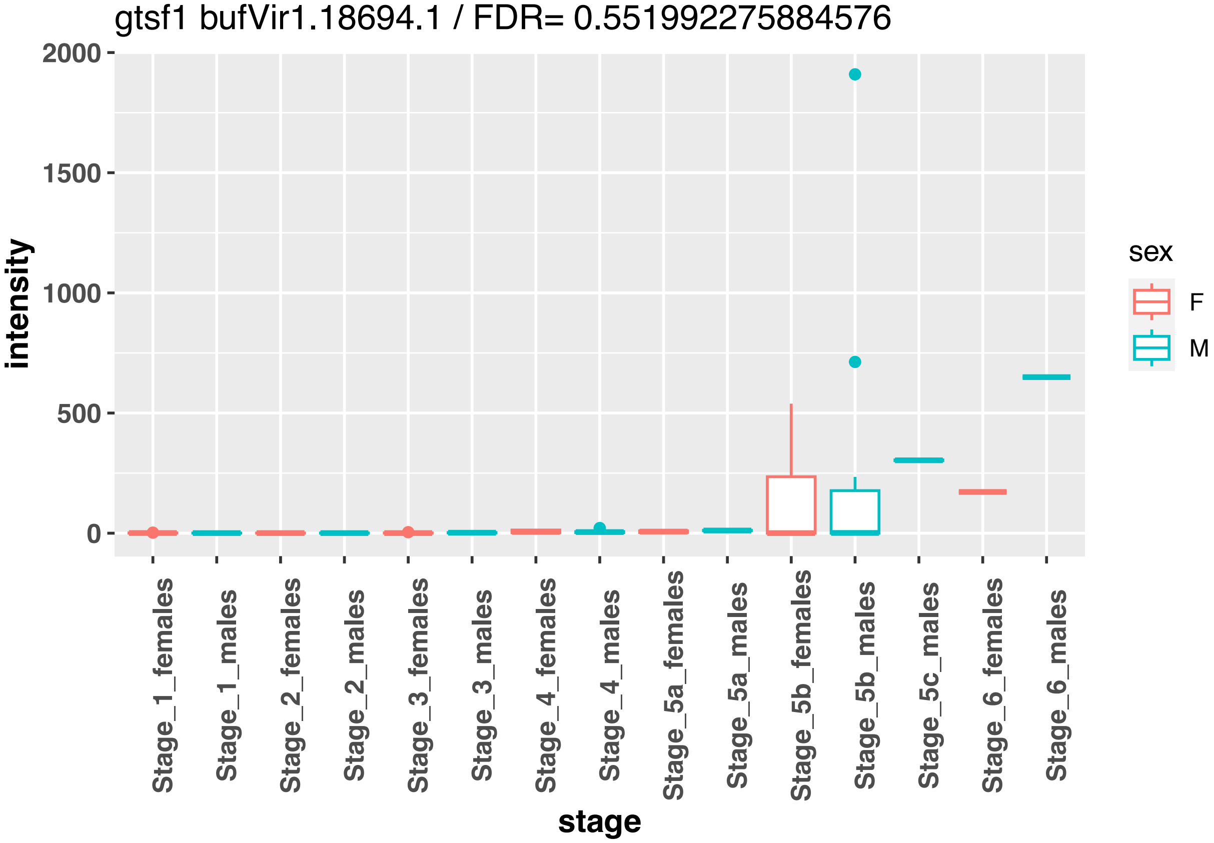

gtsf1 bufVir1.18694.1 / FDR= 0.551992275884576
intensity
2000
1500
1000
500
0
stage
Stage_1_females
Stage_1_males
Stage_2_females
Stage_2_males
Stage_3_females
Stage_3_males
Stage_4_females
Stage_4_males
Stage_5a_females
Stage_5a_males
Stage_5b_females
Stage_5b_males
Stage_5c_males
Stage_6_females
Stage_6_males
sex
F
M

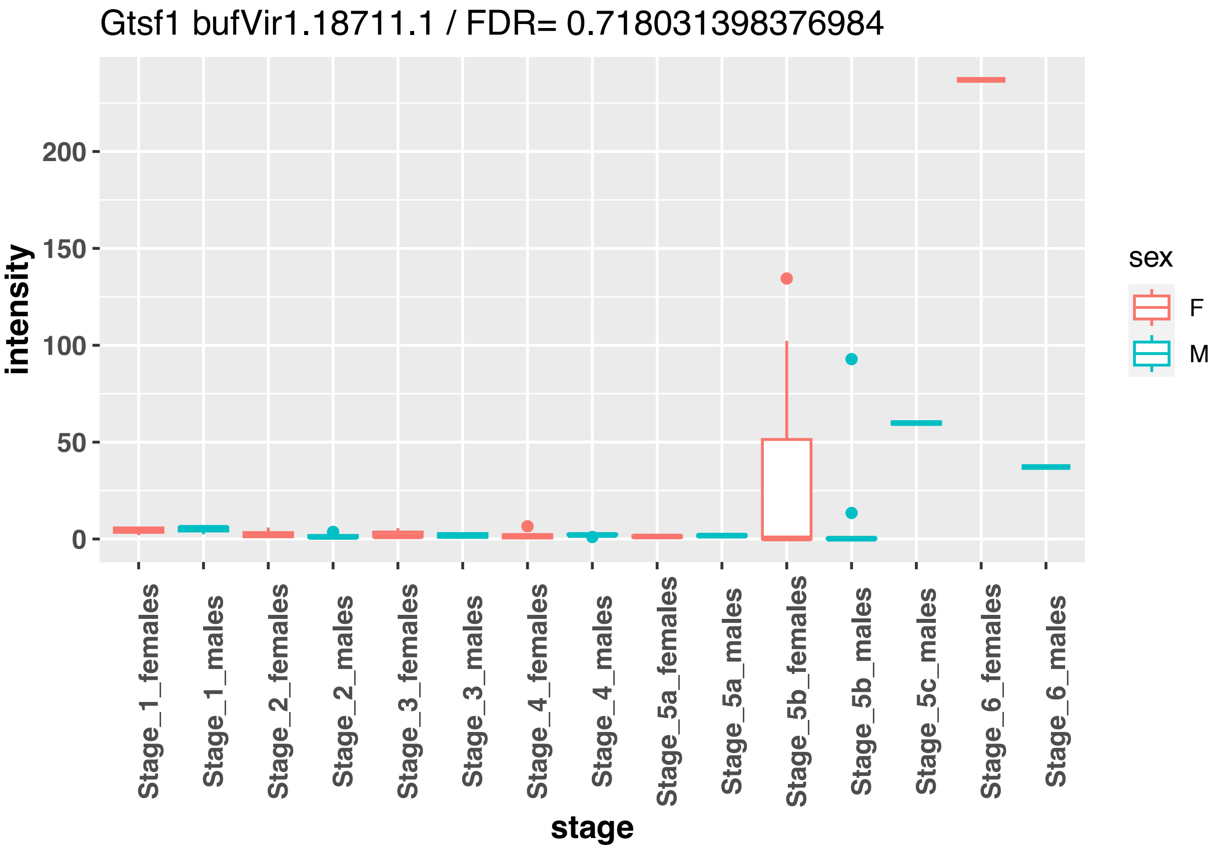
Gtsf1 bufVir1.18711.1 / FDR= 0.718031398376984
intensity
200
150
100
50
0
Stage_1_females
Stage_1_males
Stage_2_females
Stage_2_males
Stage_3_females
Stage_3_males
Stage_4_females
Stage_4_males
Stage_5a_females
Stage_5a_males
Stage_5b_females
Stage_5b_males
Stage_5c_males
Stage_6_females
Stage_6_males
stage
sex
F
M

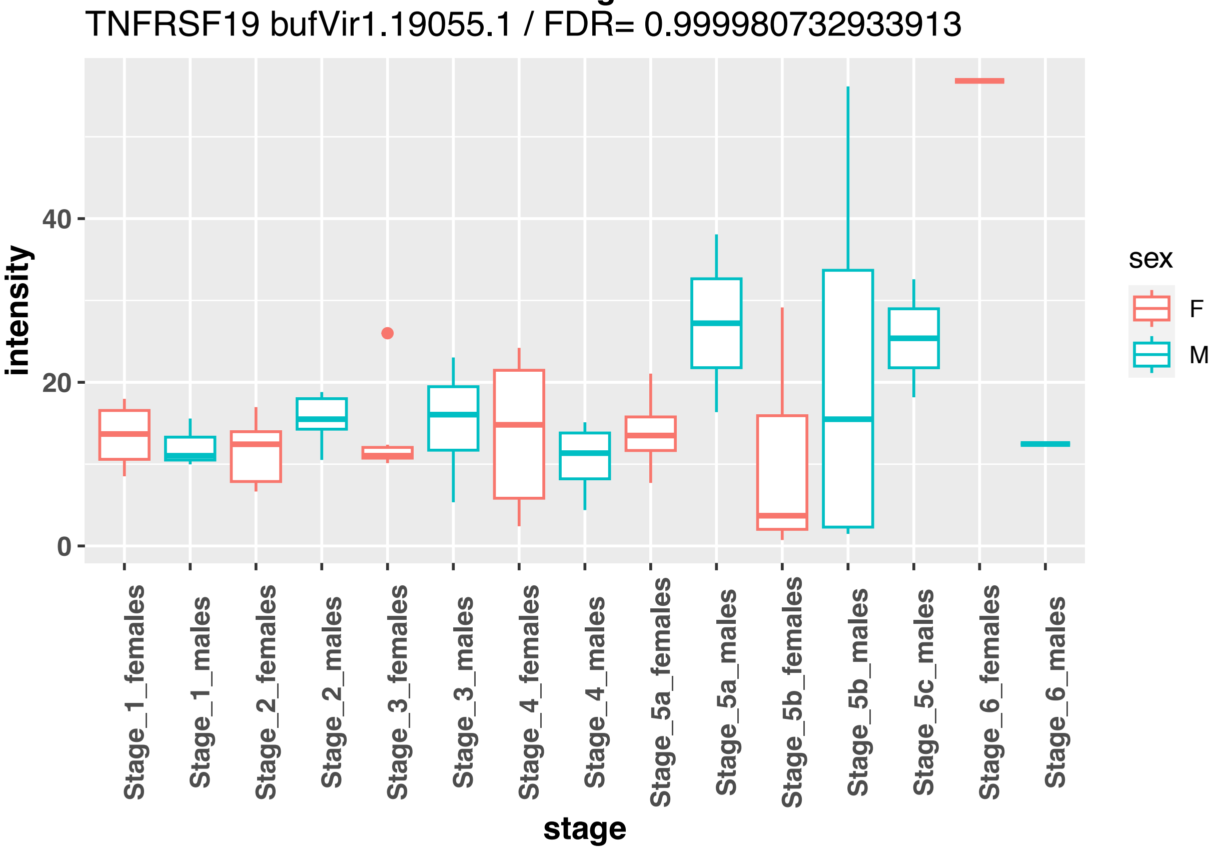
TNFRSF19 bufVir1.19055.1 / FDR= 0.999980732933913
intensity
40
20
0
Stage_1_females
Stage_1_males
Stage_2_females
Stage_2_males
Stage_3_females
Stage_3_males
Stage_4_females
Stage_4_males
Stage_5a_females
Stage_5a_males
Stage_5b_females
Stage_5b_males
Stage_5c_males
Stage_6_females
Stage_6_males
stage
sex
F
M

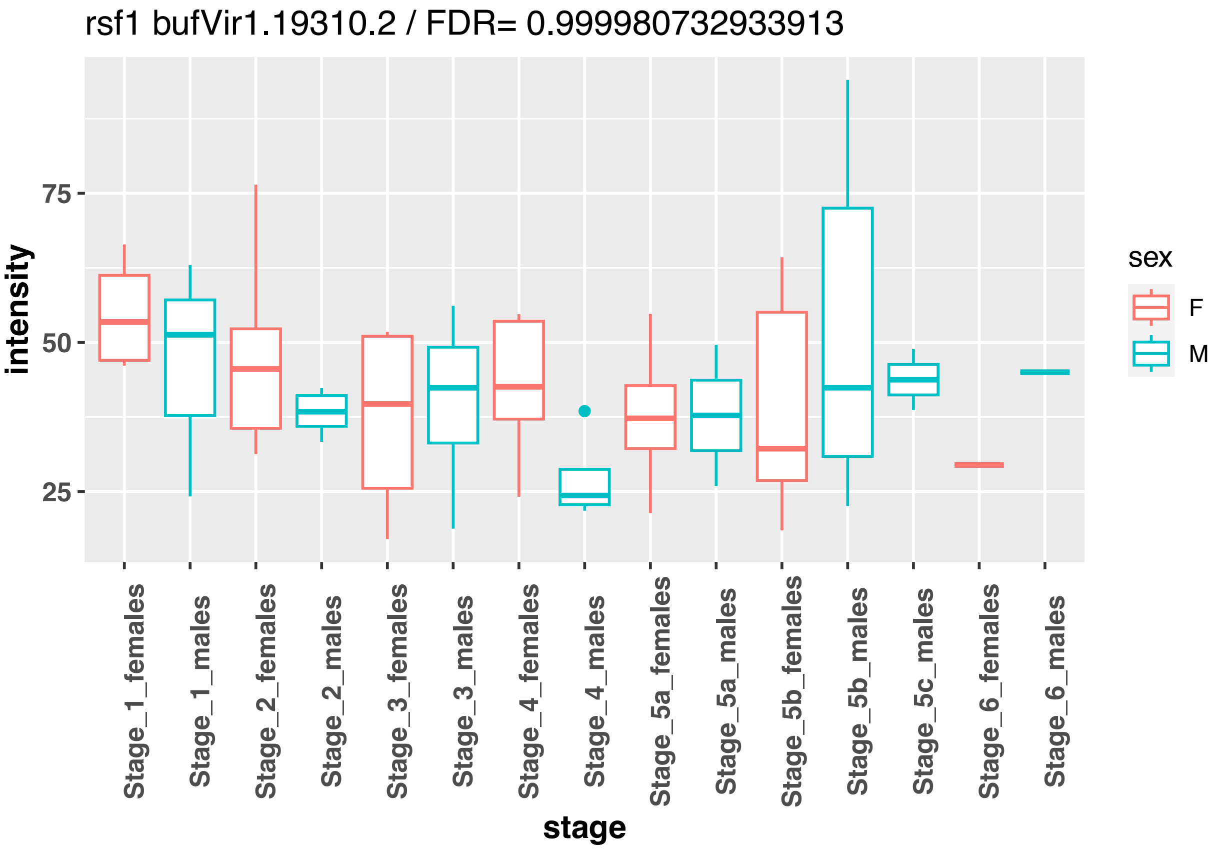
rsf1 bufVir1.19310.2 / FDR= 0.999980732933913
intensity
75
50
25
stage
Stage_1_females
Stage_1_males
Stage_2_females
Stage_2_males
Stage_3_females
Stage_3_males
Stage_4_females
Stage_4_males
Stage_5a_females
Stage_5a_males
Stage_5b_females
Stage_5b_males
Stage_5c_males
Stage_6_females
Stage_6_males
sex
F
M

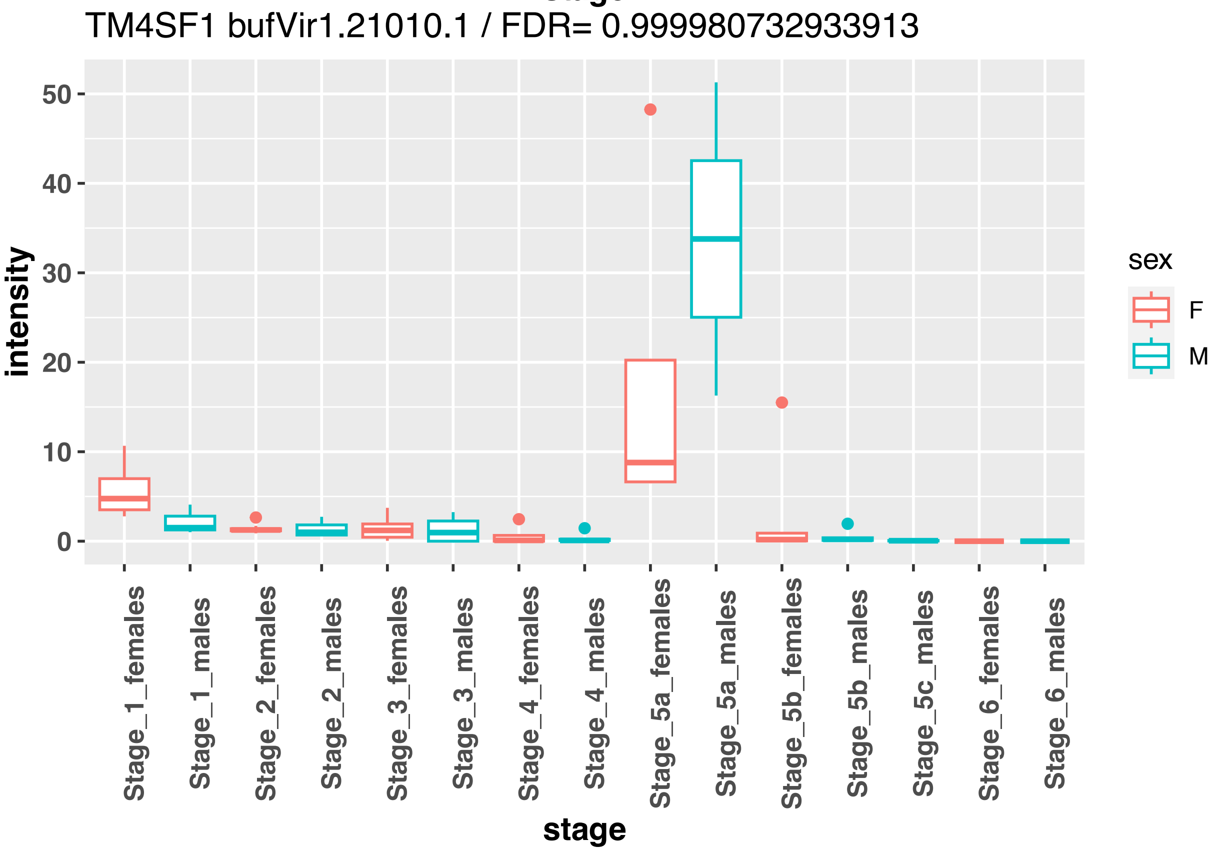
TM4SF1 bufVir1.21010.1 / FDR= 0.999980732933913
intensity
50
40
30
20
10
0
stage
Stage_1_females
Stage_1_males
Stage_2_females
Stage_2_males
Stage_3_females
Stage_3_males
Stage_4_females
Stage_4_males
Stage_5a_females
Stage_5a_males
Stage_5b_females
Stage_5b_males
Stage_5c_males
Stage_6_females
Stage_6_males
sex
F
M

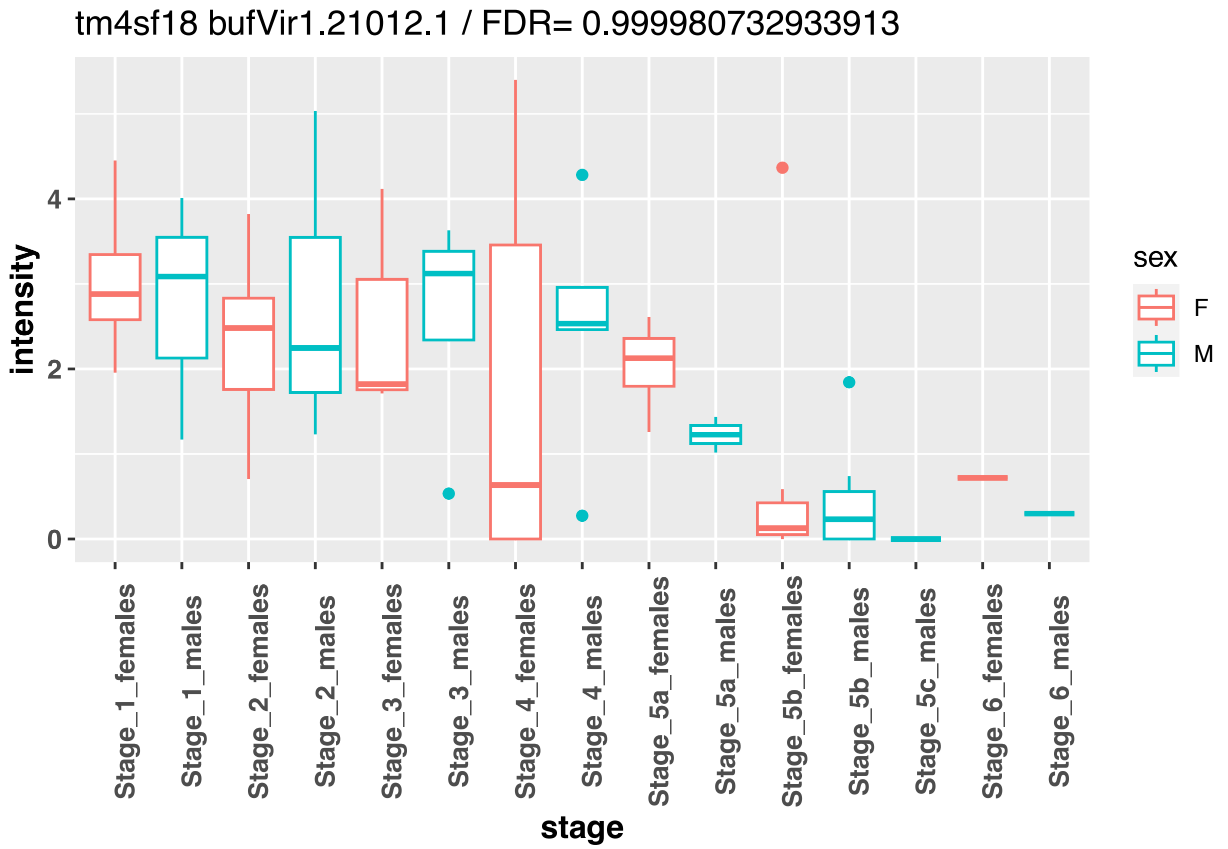
tm4sf18 bufVir1.21012.1 / FDR= 0.999980732933913
intensity
0
2
4
Stage_1_females
Stage_1_males
Stage_2_females
Stage_2_males
Stage_3_females
Stage_3_males
Stage_4_females
Stage_4_males
Stage_5a_females
Stage_5a_males
Stage_5b_females
Stage_5b_males
Stage_5c_males
Stage_6_females
Stage_6_males
stage
sex
F
M

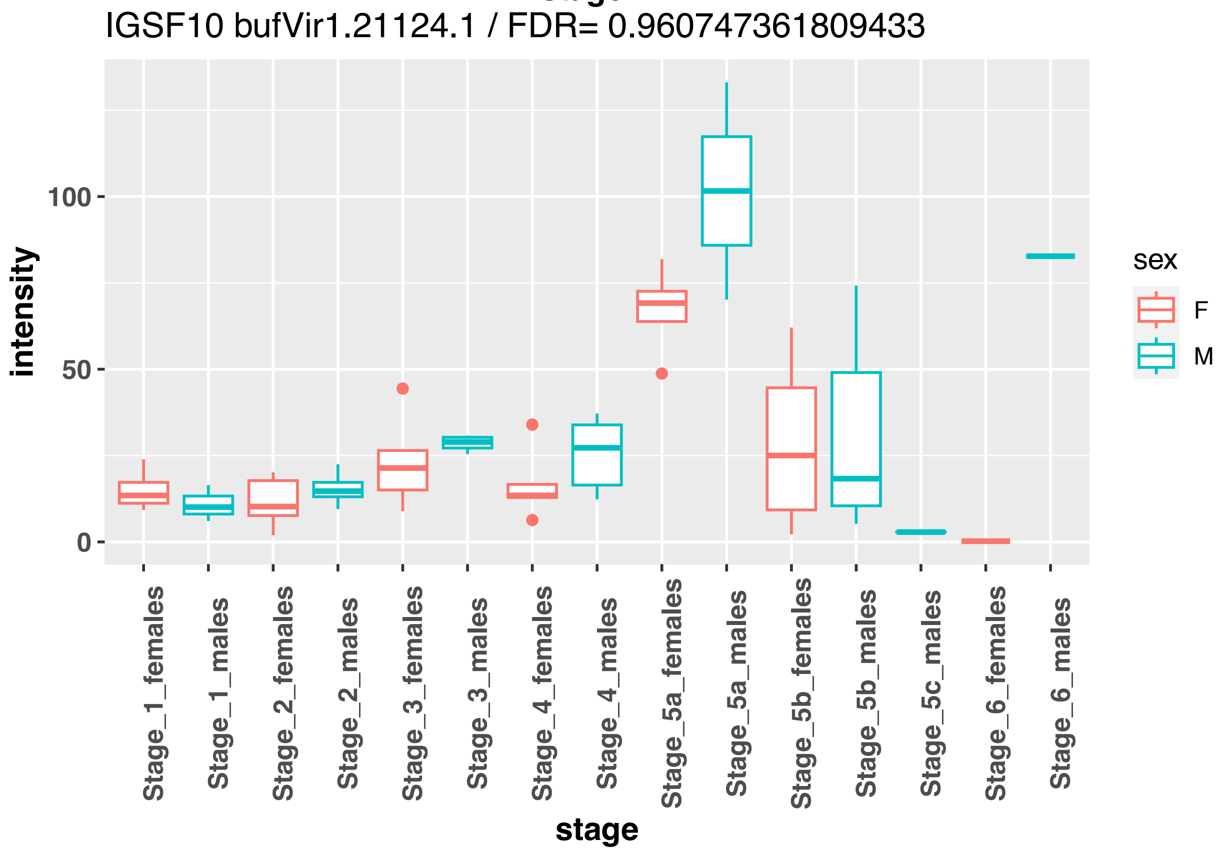
IGSF10 bufVir1.21124.1 / FDR= 0.960747361809433
intensity
0
50
100
Stage_1_females
Stage_1_males
Stage_2_females
Stage_2_males
Stage_3_females
Stage_3_males
Stage_4_females
Stage_4_males
Stage_5a_females
Stage_5a_males
Stage_5b_females
Stage_5b_males
Stage_5c_males
Stage_6_females
Stage_6_males
stage
sex
F
M

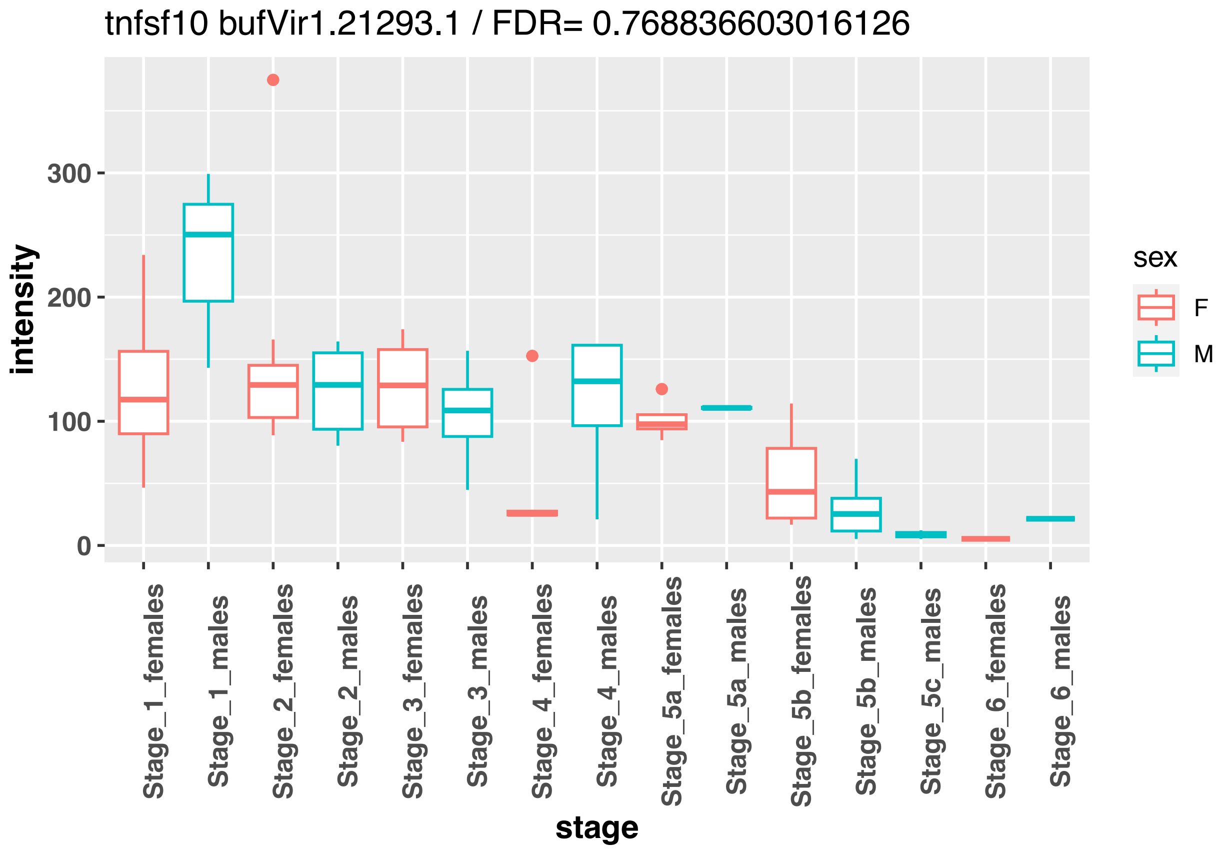
tnfsf10 bufVir1.21293.1 / FDR= 0.768836603016126
intensity
300
200
100
0
Stage_1_females
Stage_1_males
Stage_2_females
Stage_2_males
Stage_3_females
Stage_3_males
Stage_4_females
Stage_4_males
Stage_5a_females
Stage_5a_males
Stage_5b_females
Stage_5b_males
Stage_5c_males
Stage_6_females
Stage_6_males
stage
sex
F
M

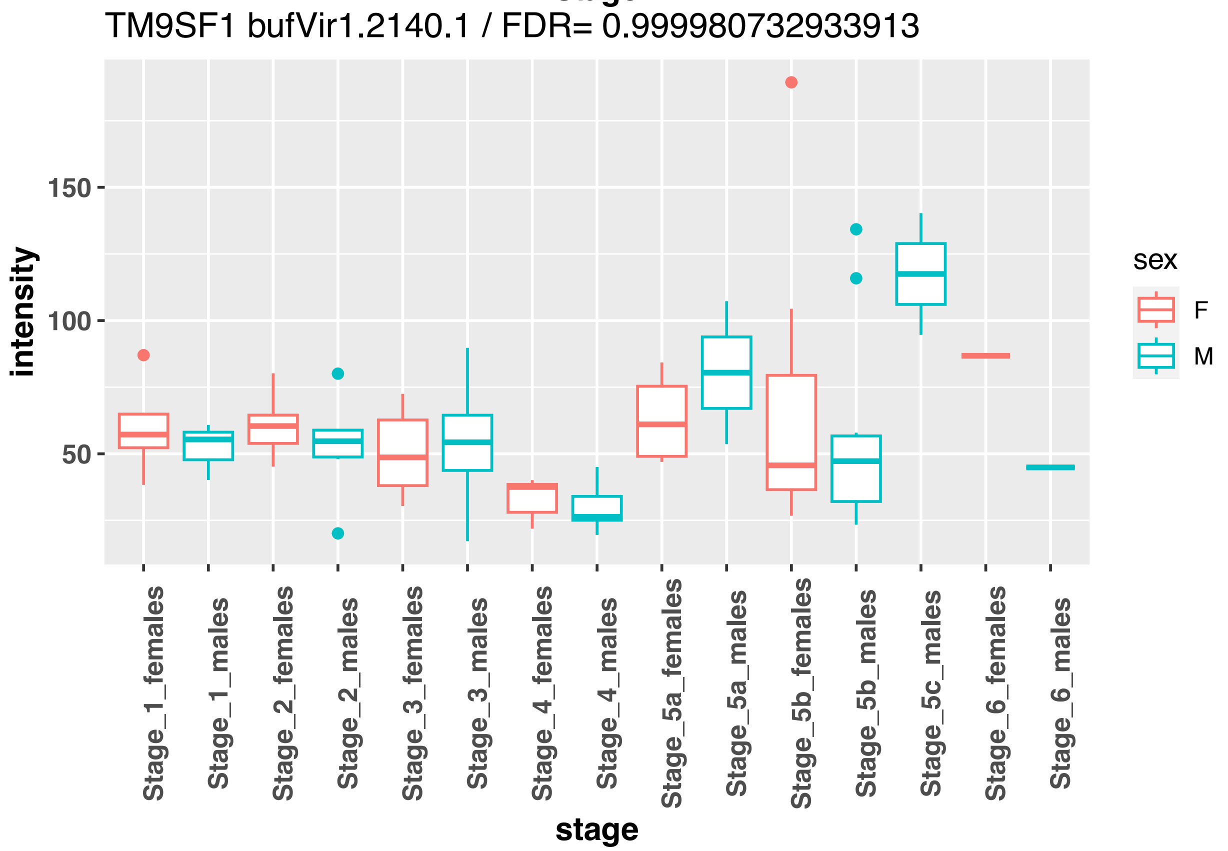
TM9SF1 bufVir1.2140.1 / FDR= 0.999980732933913
intensity
150
100
50
Stage_1_females
Stage_1_males
Stage_2_females
Stage_2_males
Stage_3_females
Stage_3_males
Stage_4_females
Stage_4_males
Stage_5a_females
Stage_5a_males
Stage_5b_females
Stage_5b_males
Stage_5c_males
Stage_6_females
Stage_6_males
stage
sex
F
M

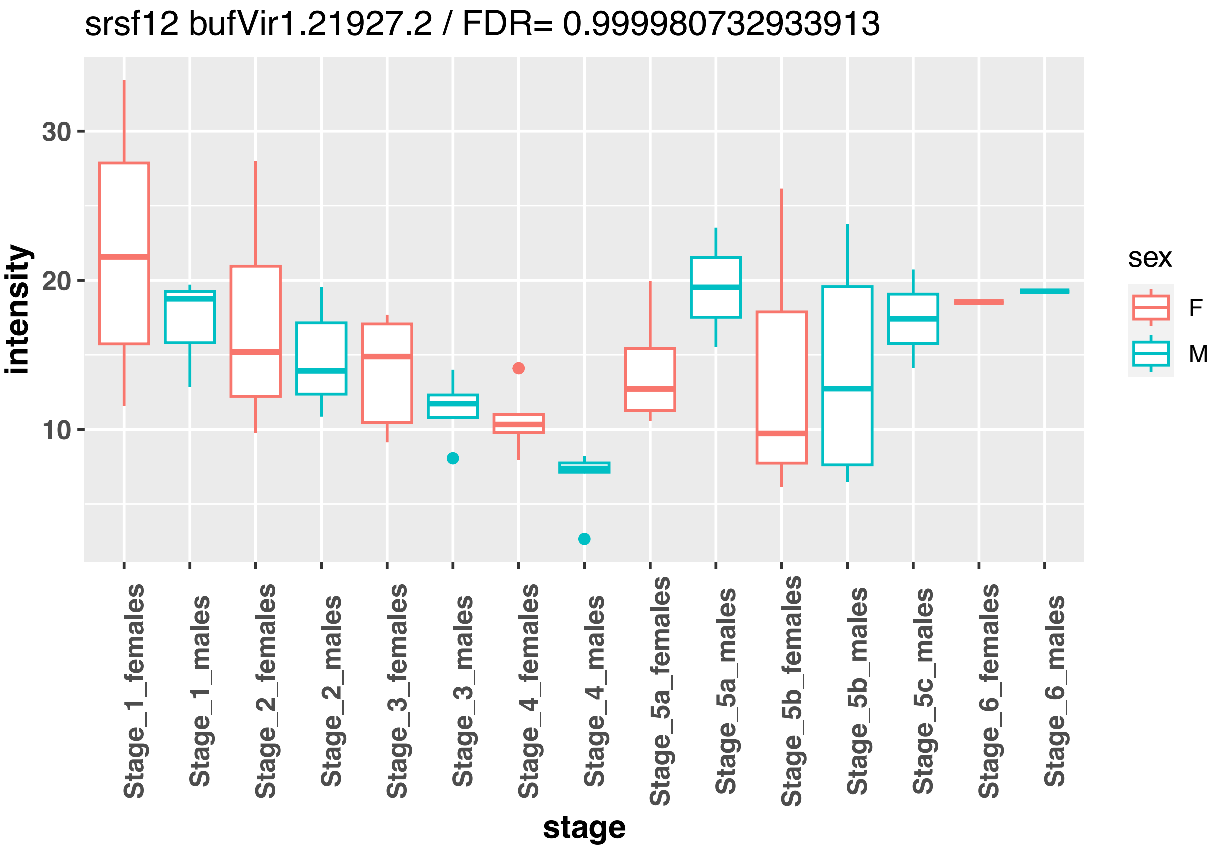
srsf12 bufVir1.21927.2 / FDR= 0.999980732933913
intensity
30
20
10
sex
F
M
Stage_1_females
Stage_1_males
Stage_2_females
Stage_2_males
Stage_3_females
Stage_3_males
Stage_4_females
Stage_4_males
Stage_5a_females
Stage_5a_males
Stage_5b_females
Stage_5b_males
Stage_5c_males
Stage_6_females
Stage_6_males
stage

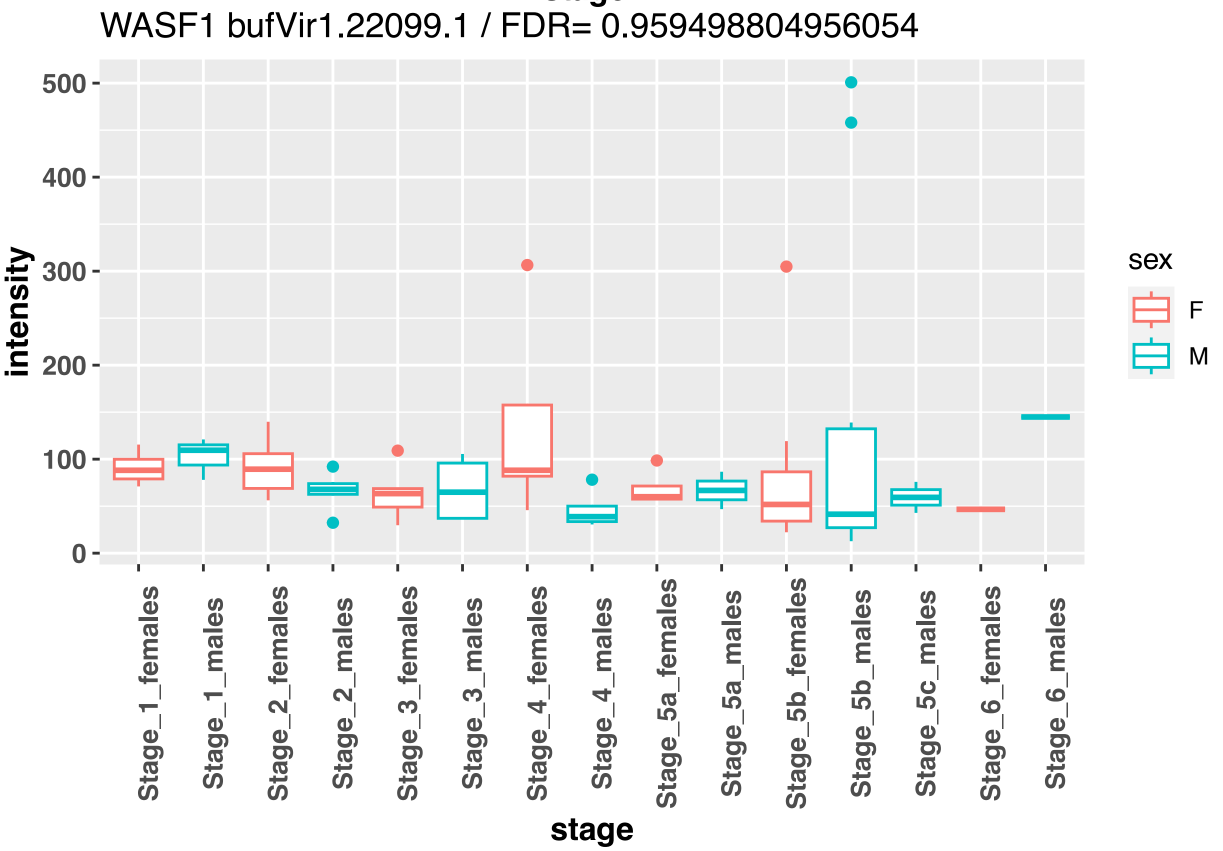
WASF1 bufVir1.22099.1 / FDR= 0.959498804956054
intensity
500
400
300
200
100
0
sex
F
M
Stage_1_females
Stage_1_males
Stage_2_females
Stage_2_males
Stage_3_females
Stage_3_males
Stage_4_females
Stage_4_males
Stage_5a_females
Stage_5a_males
Stage_5b_females
Stage_5b_males
Stage_5c_males
Stage_6_females
Stage_6_males
stage

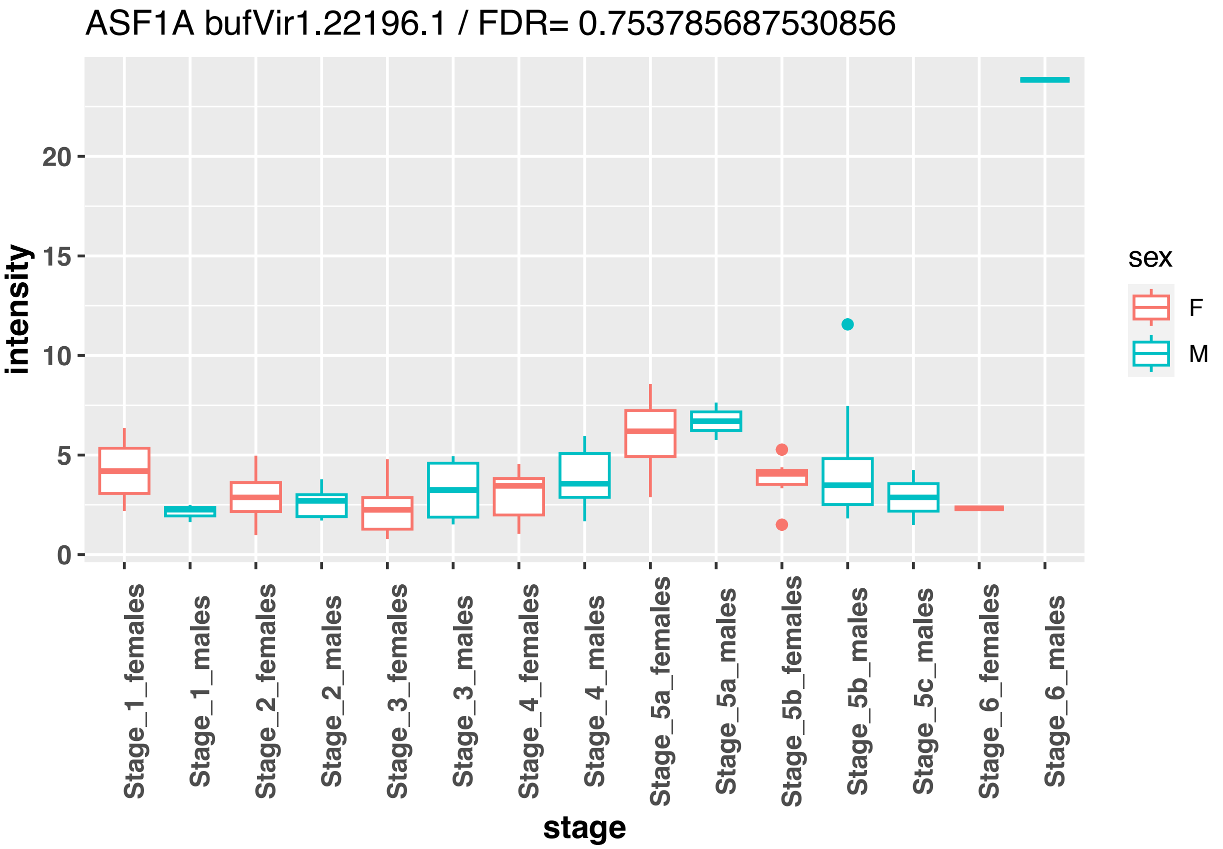
ASF1A bufVir1.22196.1 / FDR= 0.753785687530856
intensity
stage
sex
F
M
Stage_1_females
Stage_1_males
Stage_2_females
Stage_2_males
Stage_3_females
Stage_3_males
Stage_4_females
Stage_4_males
Stage_5a_females
Stage_5a_males
Stage_5b_females
Stage_5b_males
Stage_5c_males
Stage_6_females
Stage_6_males

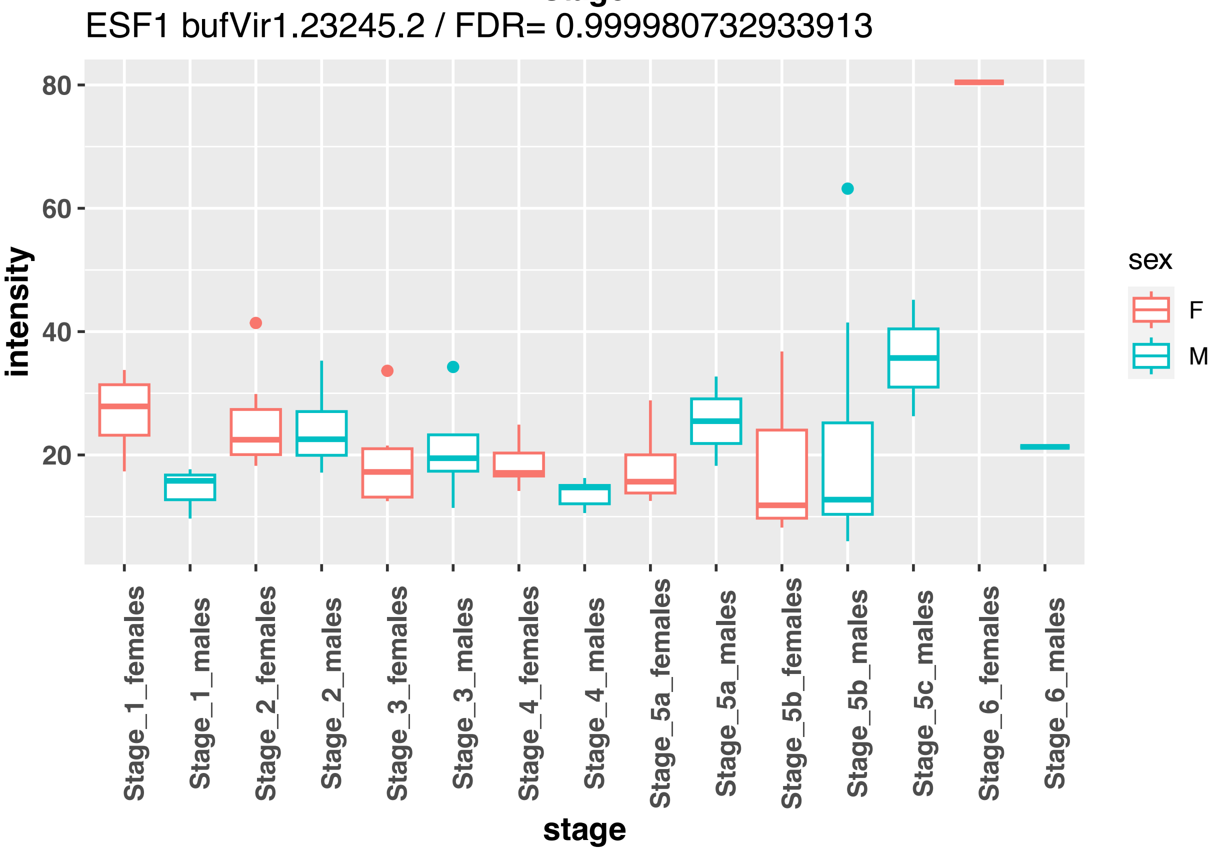
ESF1 bufVir1.23245.2 / FDR= 0.999980732933913
intensity
stage
sex
F
M
Stage_1_females
Stage_1_males
Stage_2_females
Stage_2_males
Stage_3_females
Stage_3_males
Stage_4_females
Stage_4_males
Stage_5a_females
Stage_5a_males
Stage_5b_females
Stage_5b_males
Stage_5c_males
Stage_6_females
Stage_6_males

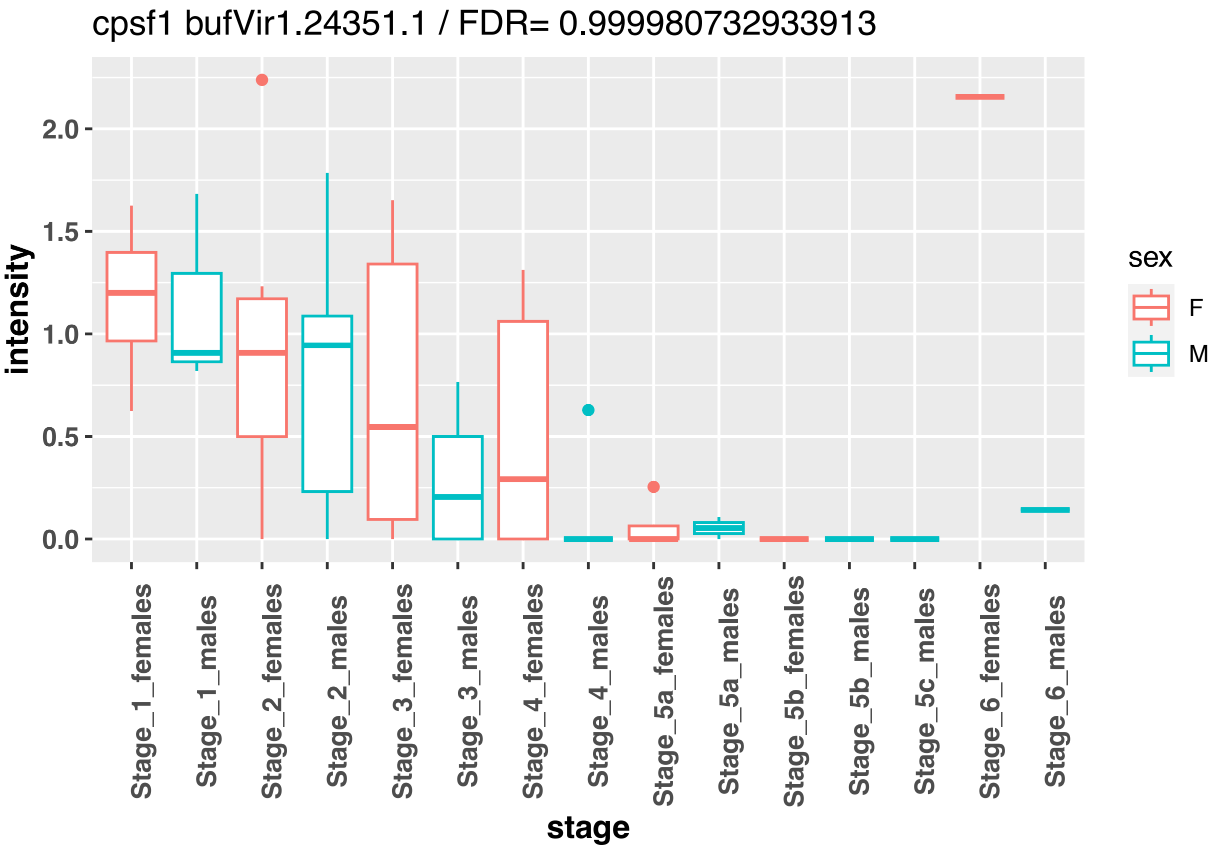
cpsf1 bufVir1.24351.1 / FDR= 0.999980732933913
intensity
0.0
0.5
1.0
1.5
2.0
stage
Stage_1_females
Stage_1_males
Stage_2_females
Stage_2_males
Stage_3_females
Stage_3_males
Stage_4_females
Stage_4_males
Stage_5a_females
Stage_5a_males
Stage_5b_females
Stage_5b_males
Stage_5c_males
Stage_6_females
Stage_6_males
sex
F
M

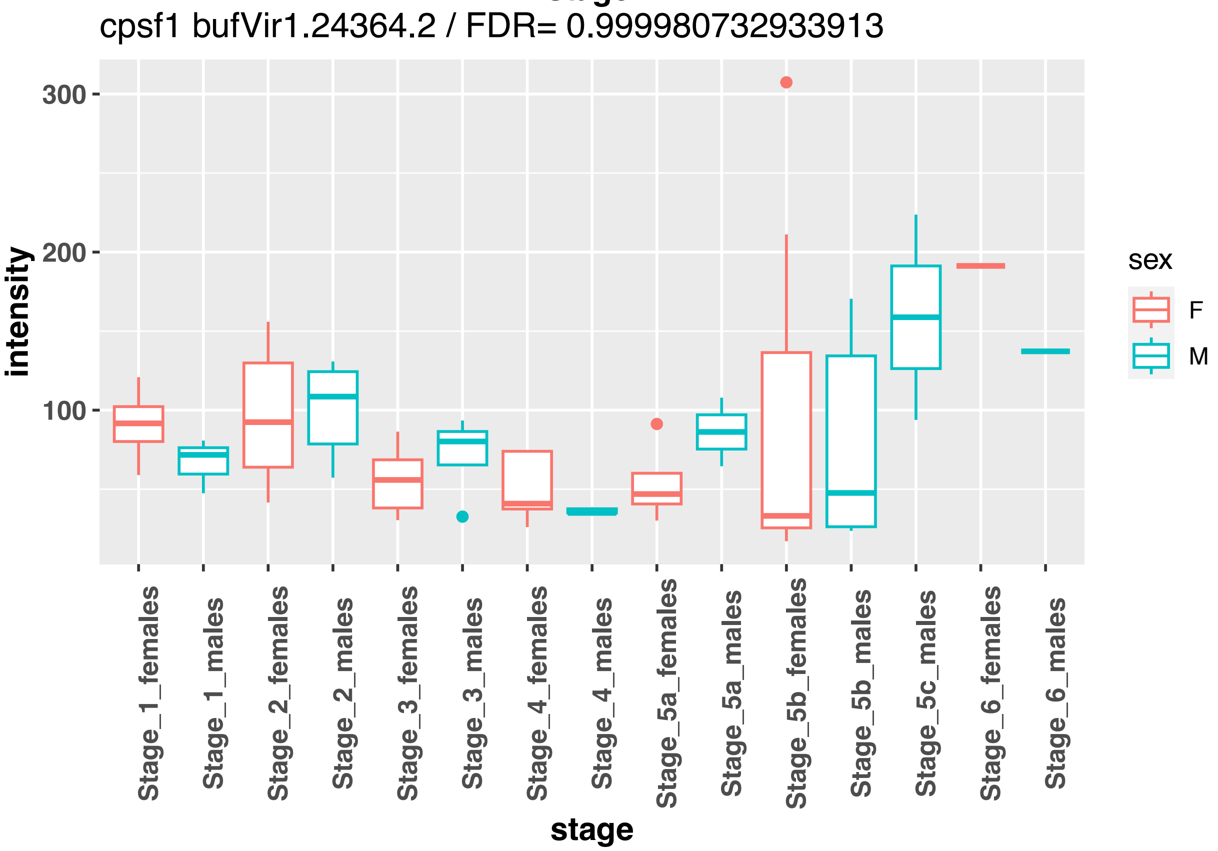
cpsf1 bufVir1.24364.2 / FDR= 0.999980732933913
intensity
100
200
300
stage
Stage_1_females
Stage_1_males
Stage_2_females
Stage_2_males
Stage_3_females
Stage_3_males
Stage_4_females
Stage_4_males
Stage_5a_females
Stage_5a_males
Stage_5b_females
Stage_5b_males
Stage_5c_males
Stage_6_females
Stage_6_males
sex
F
M

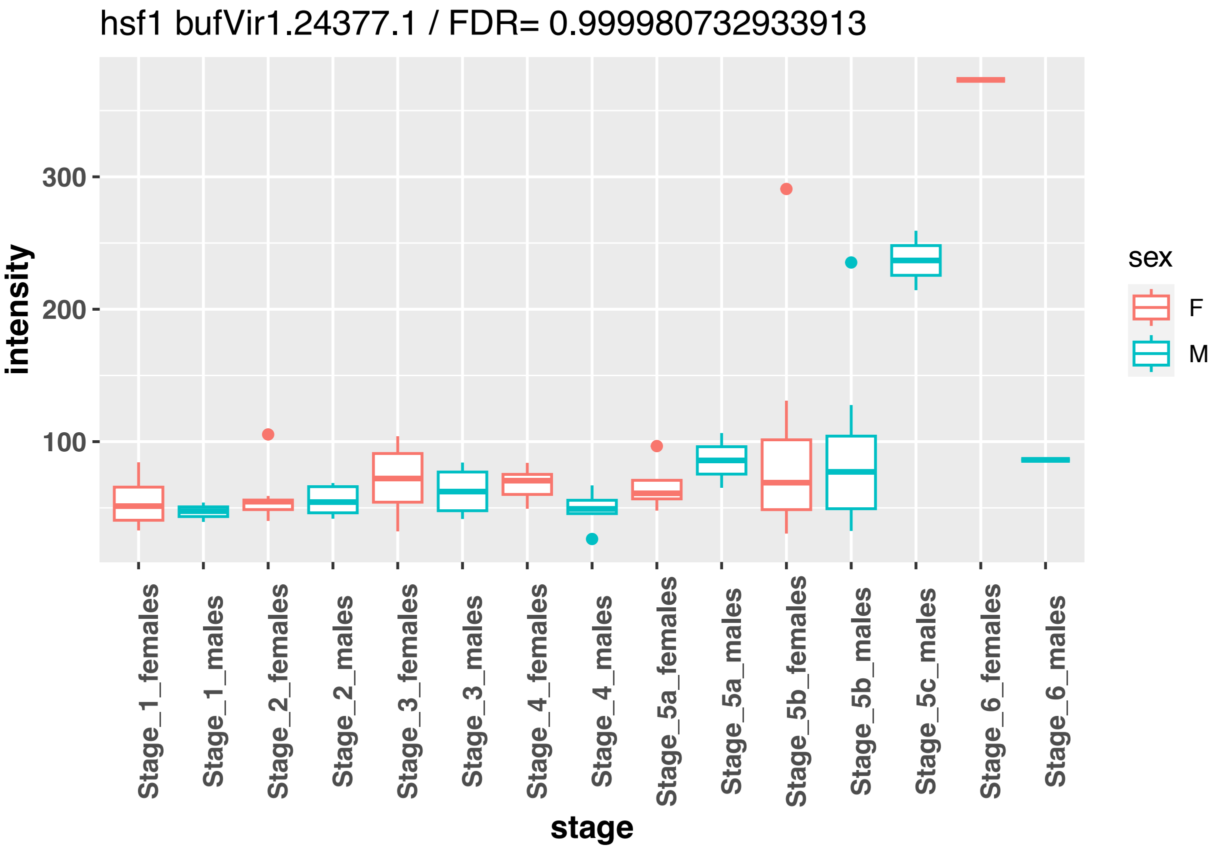
hsf1 bufVir1.24377.1 / FDR= 0.999980732933913
intensity
300
200
100
stage
Stage_1_females
Stage_1_males
Stage_2_females
Stage_2_males
Stage_3_females
Stage_3_males
Stage_4_females
Stage_4_males
Stage_5a_females
Stage_5a_males
Stage_5b_females
Stage_5b_males
Stage_5c_males
Stage_6_females
Stage_6_males
sex
F
M

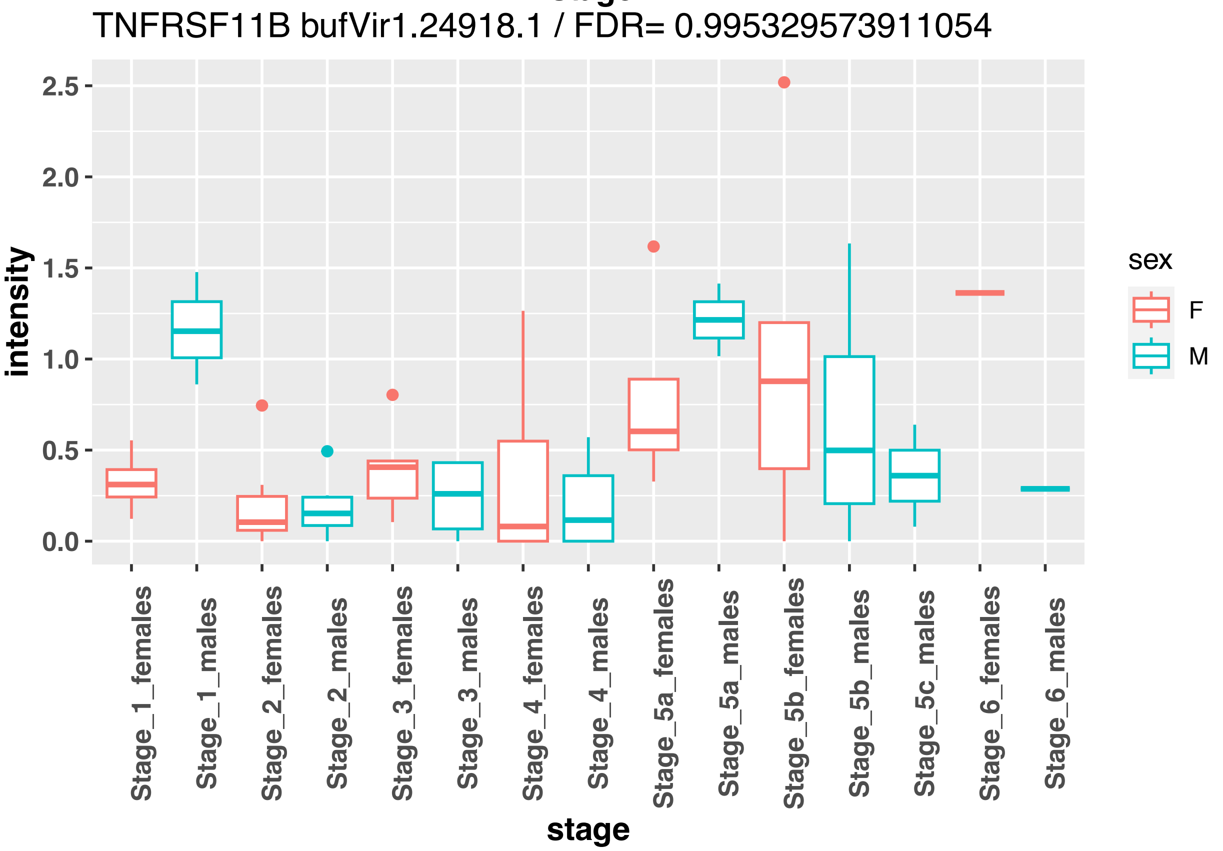
TNFRSF11B bufVir1.24918.1 / FDR= 0.995329573911054
intensity
2.5
2.0
1.5
1.0
0.5
0.0
stage
Stage_1_females
Stage_1_males
Stage_2_females
Stage_2_males
Stage_3_females
Stage_3_males
Stage_4_females
Stage_4_males
Stage_5a_females
Stage_5a_males
Stage_5b_females
Stage_5b_males
Stage_5c_males
Stage_6_females
Stage_6_males
sex
F
M

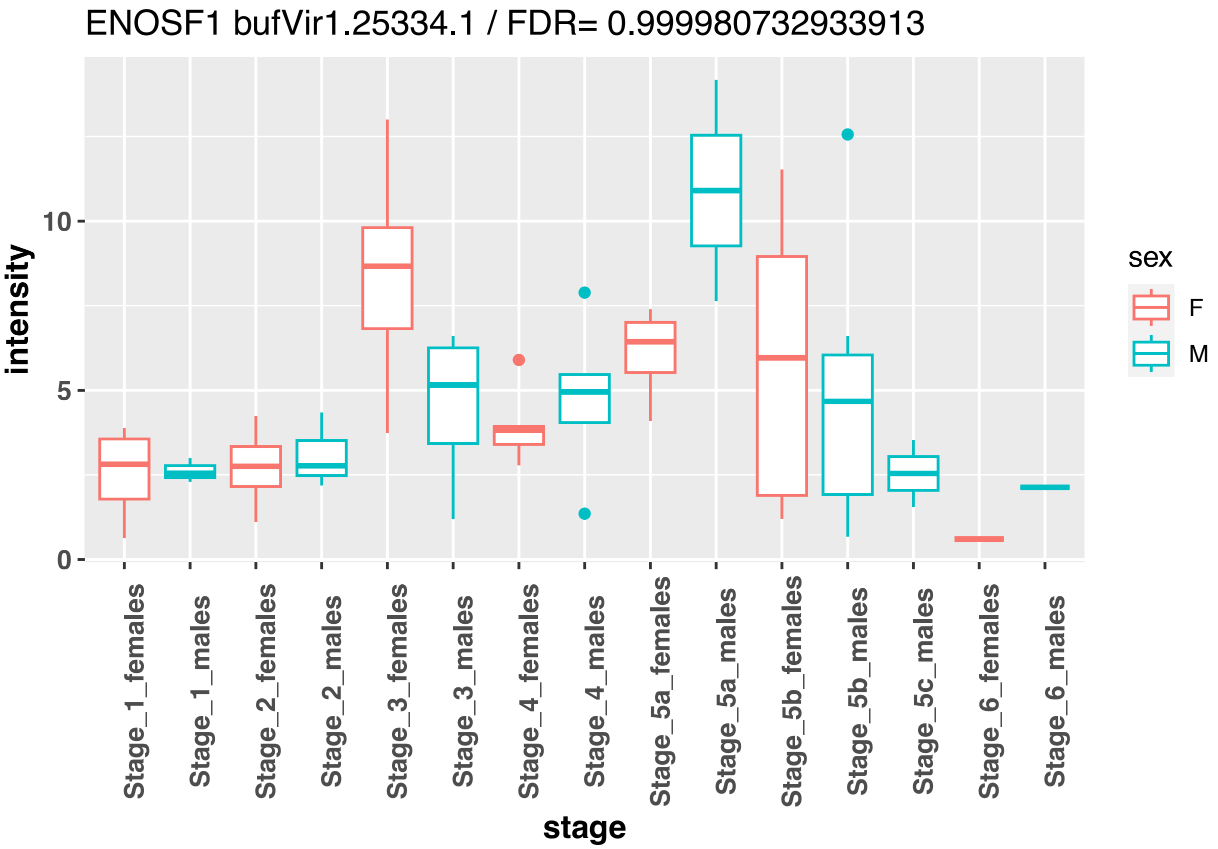
ENOSF1 bufVir1.25334.1 / FDR= 0.999980732933913
intensity
10
5
0
Stage_1_females
Stage_1_males
Stage_2_females
Stage_2_males
Stage_3_females
Stage_3_males
Stage_4_females
Stage_4_males
Stage_5a_females
Stage_5a_males
Stage_5b_females
Stage_5b_males
Stage_5c_males
Stage_6_females
Stage_6_males
stage
sex
F
M

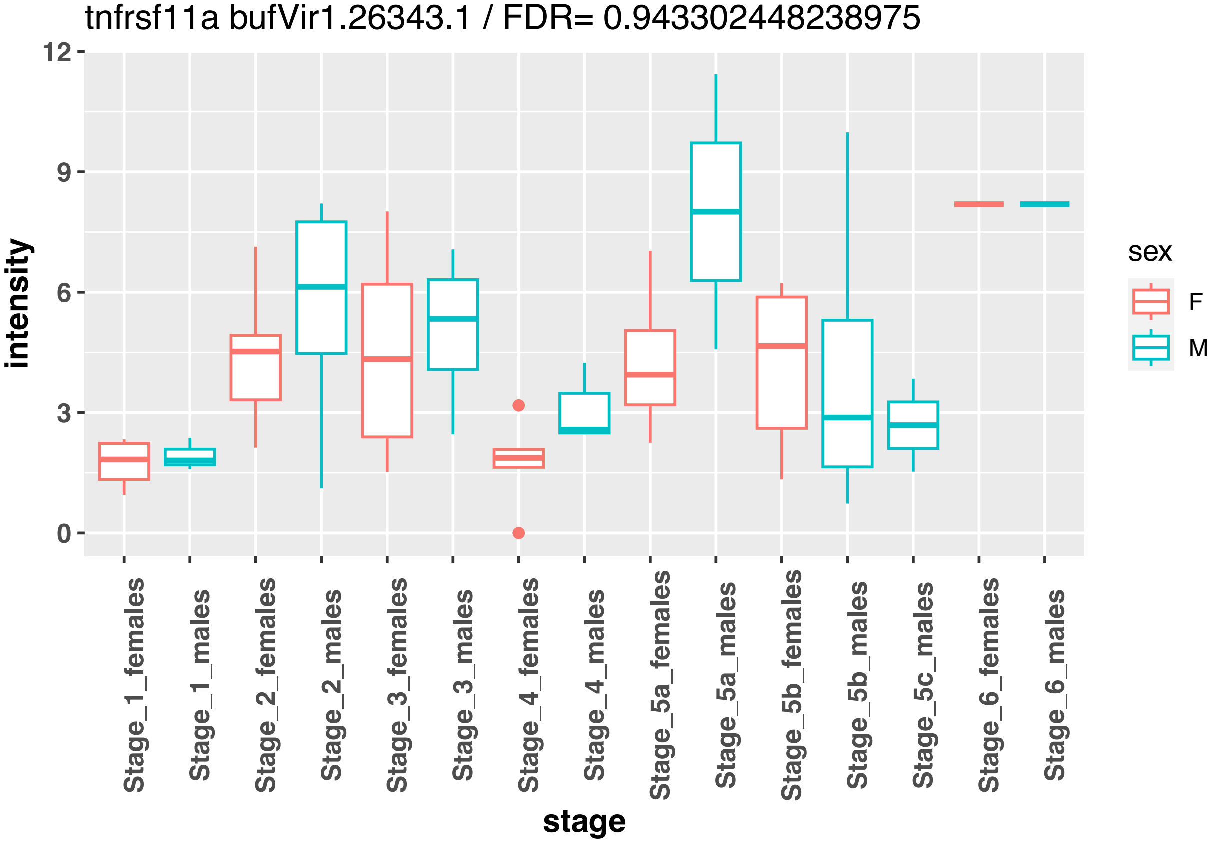
tnfrsf11a bufVir1.26343.1 / FDR= 0.943302448238975
intensity
12
9
6
3
0
Stage_1_females
Stage_1_males
Stage_2_females
Stage_2_males
Stage_3_females
Stage_3_males
Stage_4_females
Stage_4_males
Stage_5a_females
Stage_5a_males
Stage_5b_females
Stage_5b_males
Stage_5c_males
Stage_6_females
Stage_6_males
stage
sex
F
M

cpsf1 bufVir1.27782.1 / FDR= 0.999980732933913

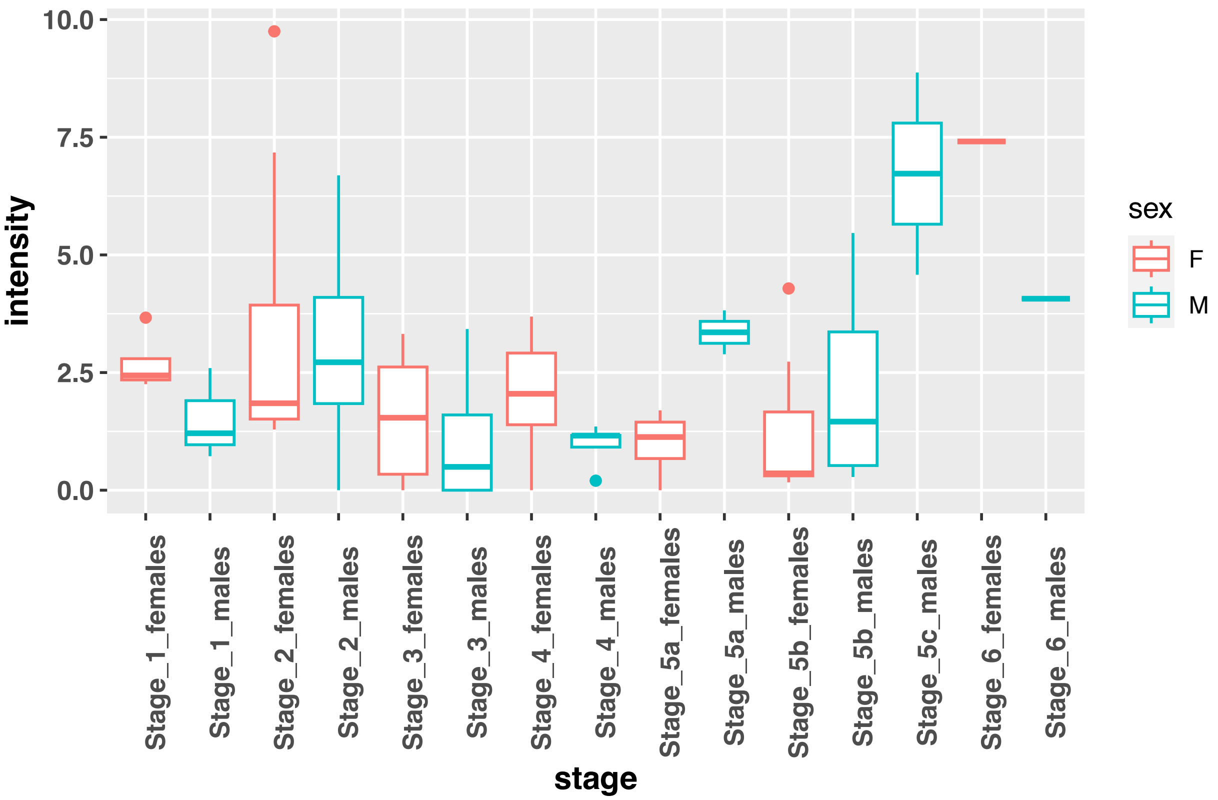


cpsf1 bufVir1.27783.1 / FDR= 0.920820971685537

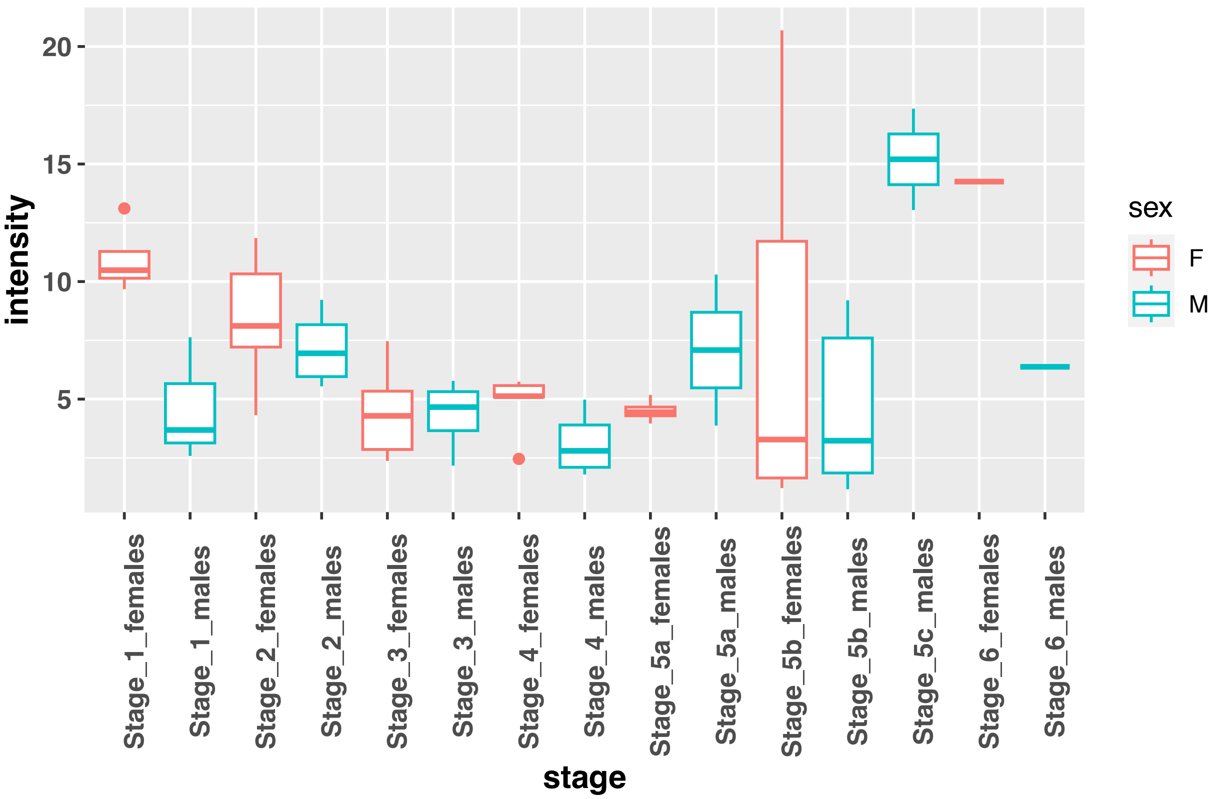

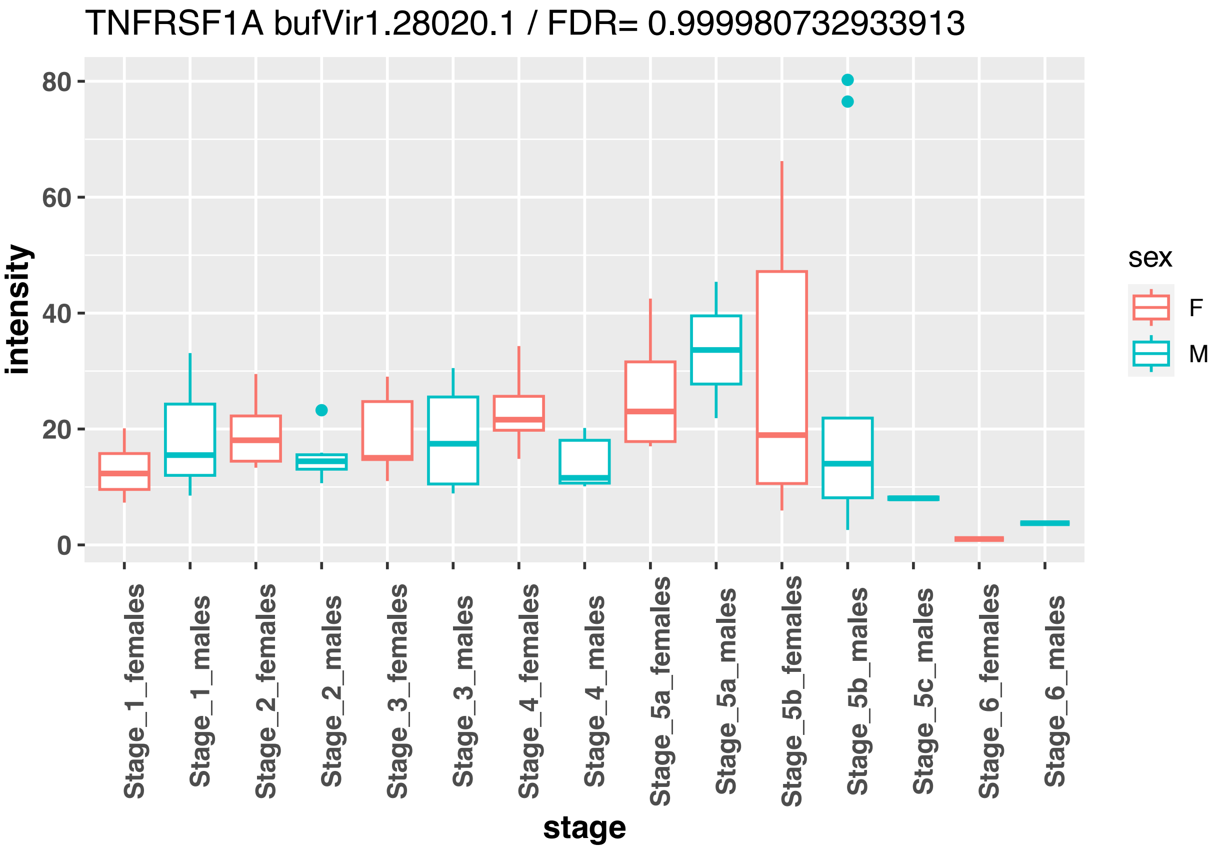
TNFRSF1A bufVir1.28020.1 / FDR= 0.999980732933913
intensity
80
60
40
20
0
stage
Stage_1_females
Stage_1_males
Stage_2_females
Stage_2_males
Stage_3_females
Stage_3_males
Stage_4_females
Stage_4_males
Stage_5a_females
Stage_5a_males
Stage_5b_females
Stage_5b_males
Stage_5c_males
Stage_6_females
Stage_6_males
sex
F
M

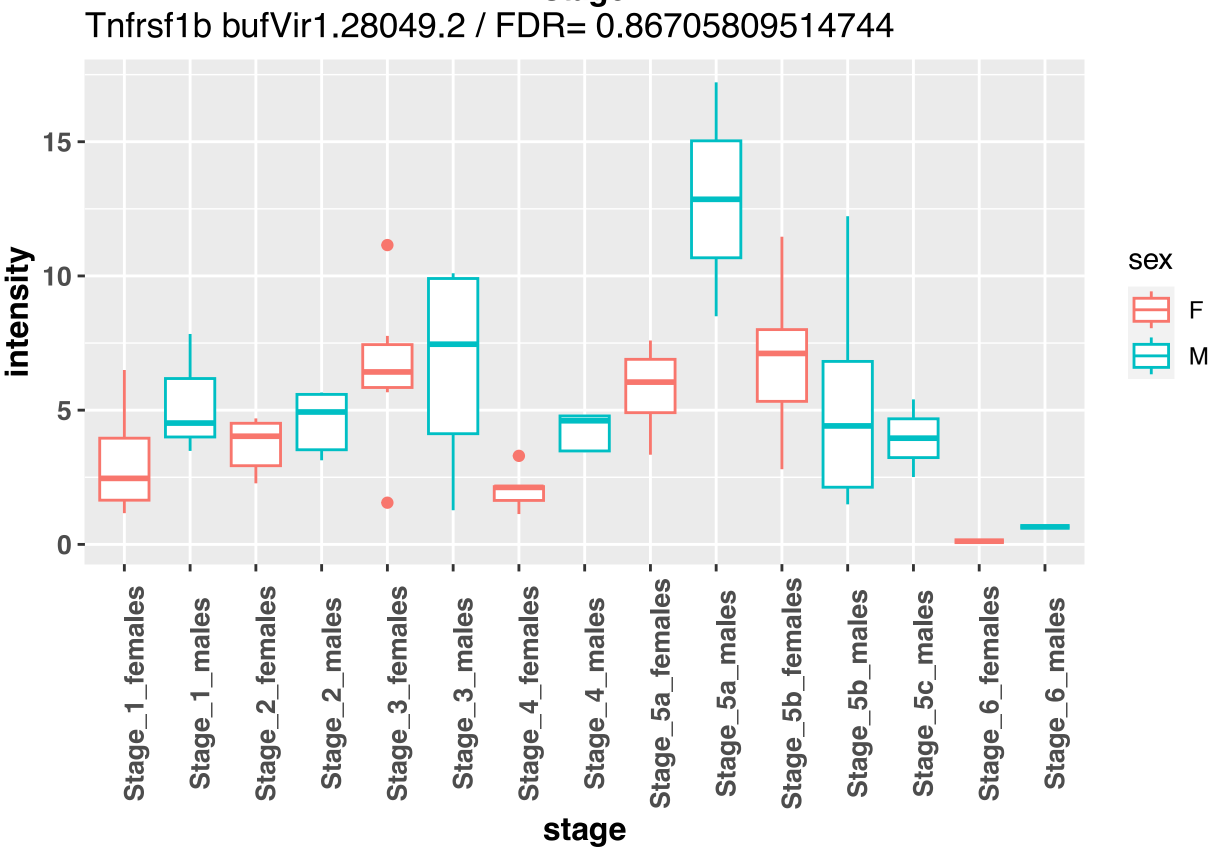
Tnfrsf1b bufVir1.28049.2 / FDR= 0.86705809514744
intensity
15
10
5
0
stage
Stage_1_females
Stage_1_males
Stage_2_females
Stage_2_males
Stage_3_females
Stage_3_males
Stage_4_females
Stage_4_males
Stage_5a_females
Stage_5a_males
Stage_5b_females
Stage_5b_males
Stage_5c_males
Stage_6_females
Stage_6_males
sex
F
M

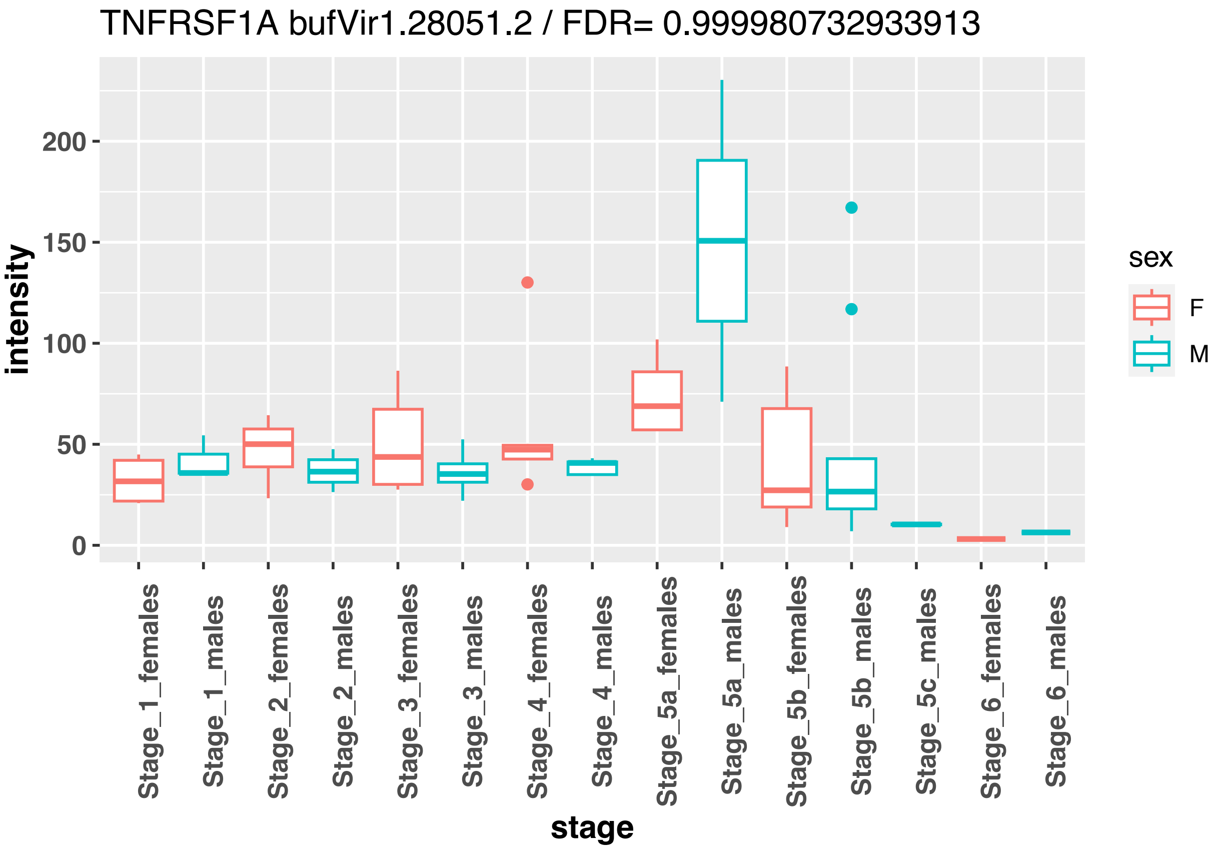
TNFRSF1A bufVir1.28051.2 / FDR= 0.999980732933913
intensity
0
50
100
150
200
stage
Stage_1_females
Stage_1_males
Stage_2_females
Stage_2_males
Stage_3_females
Stage_3_males
Stage_4_females
Stage_4_males
Stage_5a_females
Stage_5a_males
Stage_5b_females
Stage_5b_males
Stage_5c_males
Stage_6_females
Stage_6_males
sex
F
M

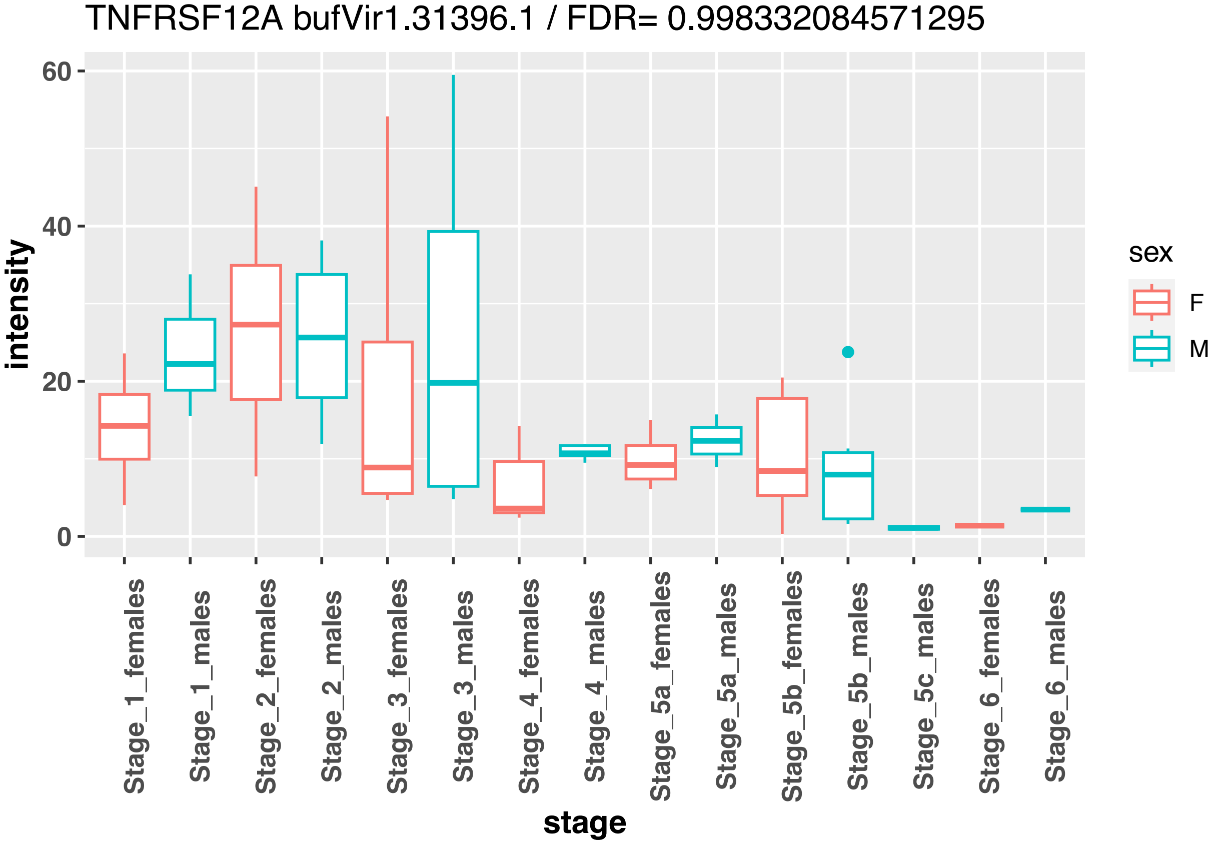
TNFRSF12A bufVir1.31396.1 / FDR= 0.998332084571295
intensity
0
20
40
60
stage
Stage_1_females
Stage_1_males
Stage_2_females
Stage_2_males
Stage_3_females
Stage_3_males
Stage_4_females
Stage_4_males
Stage_5a_females
Stage_5a_males
Stage_5b_females
Stage_5b_males
Stage_5c_males
Stage_6_females
Stage_6_males
sex
F
M

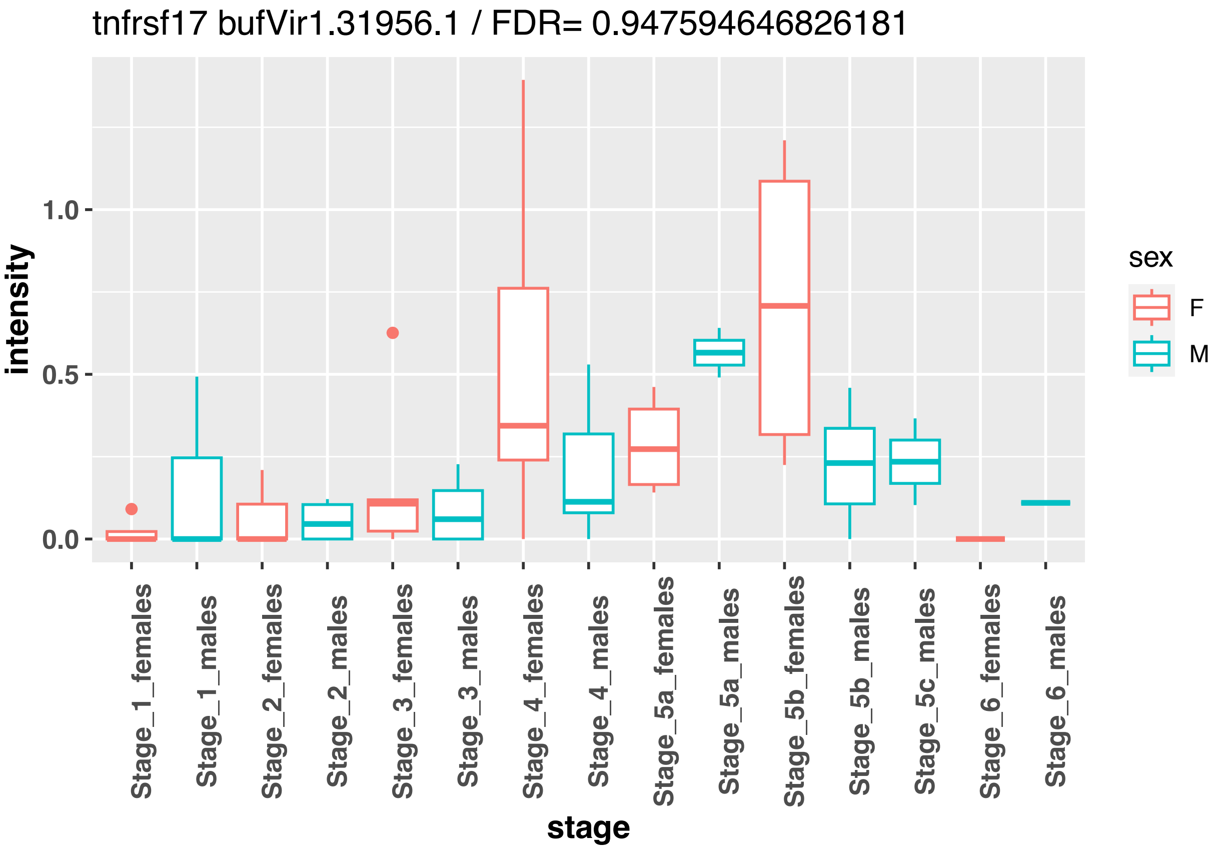
tnfrsf17 bufVir1.31956.1 / FDR= 0.947594646826181
intensity
1.0
0.5
0.0
Stage_1_females
Stage_1_males
Stage_2_females
Stage_2_males
Stage_3_females
Stage_3_males
Stage_4_females
Stage_4_males
Stage_5a_females
Stage_5a_males
Stage_5b_females
Stage_5b_males
Stage_5c_males
Stage_6_females
Stage_6_males
stage
sex
F
M

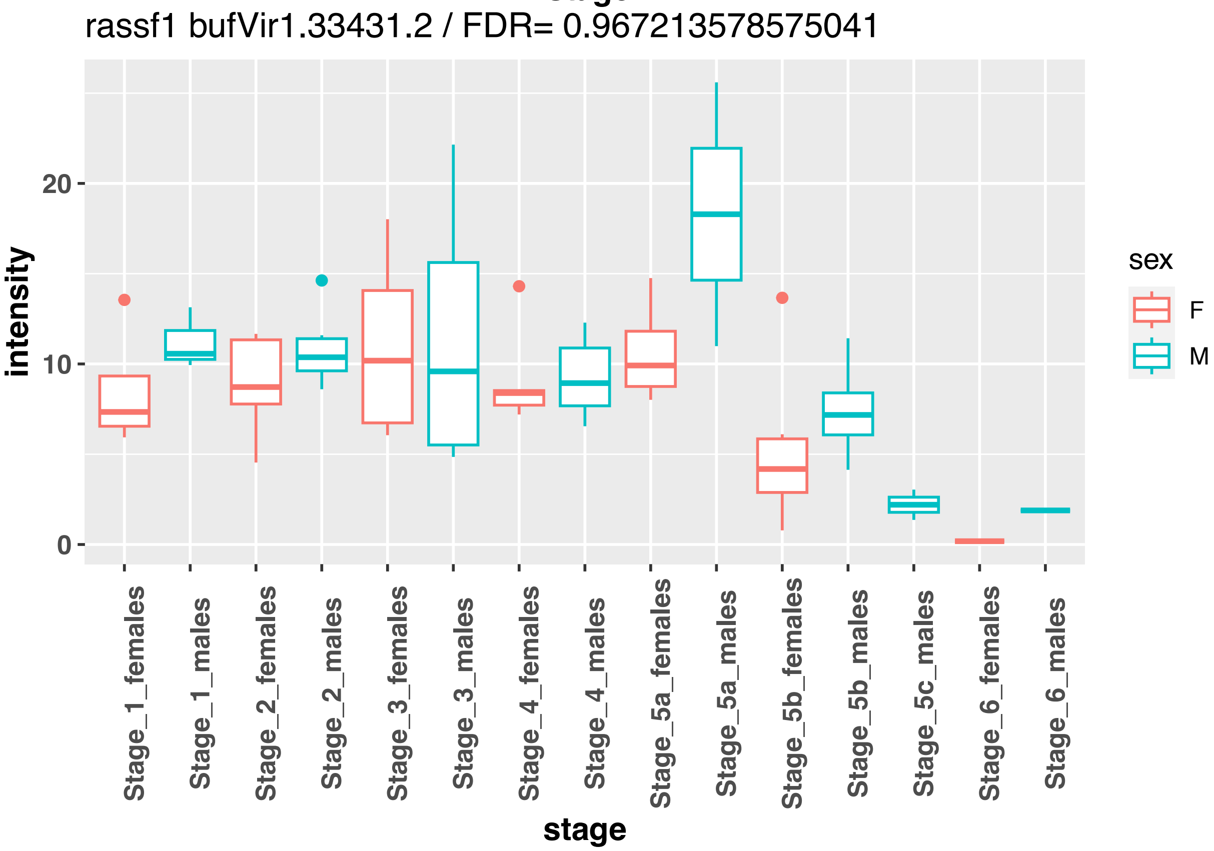
rassf1 bufVir1.33431.2 / FDR= 0.967213578575041
intensity
20
10
0
Stage_1_females
Stage_1_males
Stage_2_females
Stage_2_males
Stage_3_females
Stage_3_males
Stage_4_females
Stage_4_males
Stage_5a_females
Stage_5a_males
Stage_5b_females
Stage_5b_males
Stage_5c_males
Stage_6_females
Stage_6_males
stage
sex
F
M

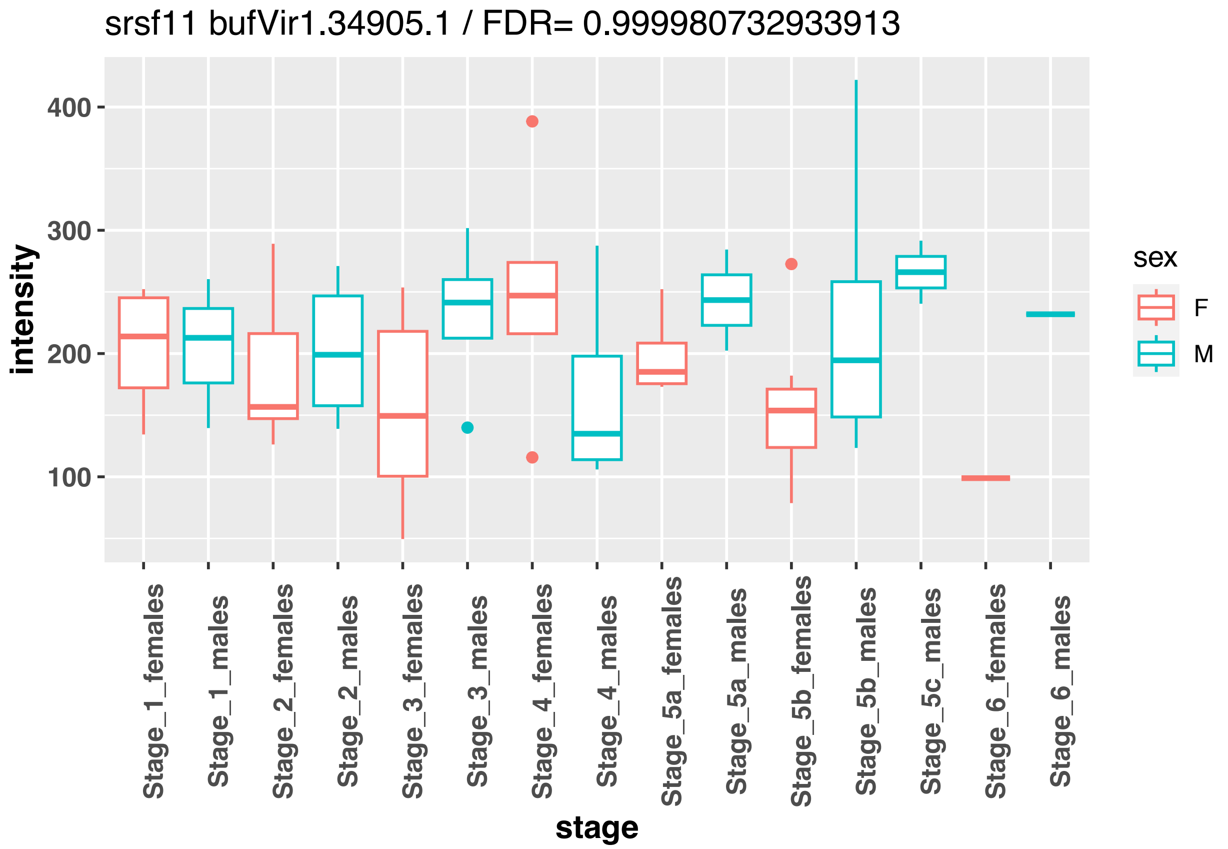
srsf11 bufVir1.34905.1 / FDR= 0.999980732933913
intensity
400
300
200
100
stage
Stage_1_females
Stage_1_males
Stage_2_females
Stage_2_males
Stage_3_females
Stage_3_males
Stage_4_females
Stage_4_males
Stage_5a_females
Stage_5a_males
Stage_5b_females
Stage_5b_males
Stage_5c_males
Stage_6_females
Stage_6_males
sex
F
M

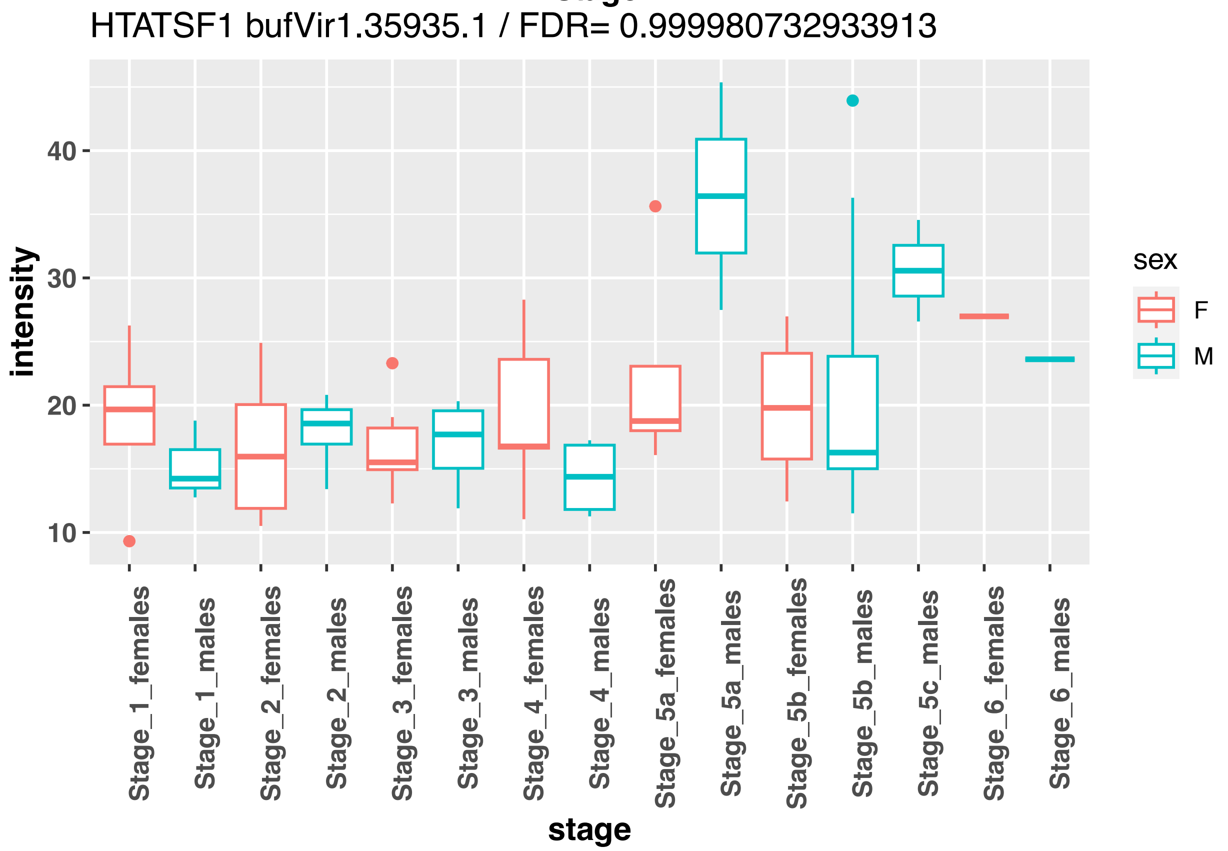
HTATSF1 bufVir1.35935.1 / FDR= 0.999980732933913
intensity
40
30
20
10
stage
Stage_1_females
Stage_1_males
Stage_2_females
Stage_2_males
Stage_3_females
Stage_3_males
Stage_4_females
Stage_4_males
Stage_5a_females
Stage_5a_males
Stage_5b_females
Stage_5b_males
Stage_5c_males
Stage_6_females
Stage_6_males
sex
F
M

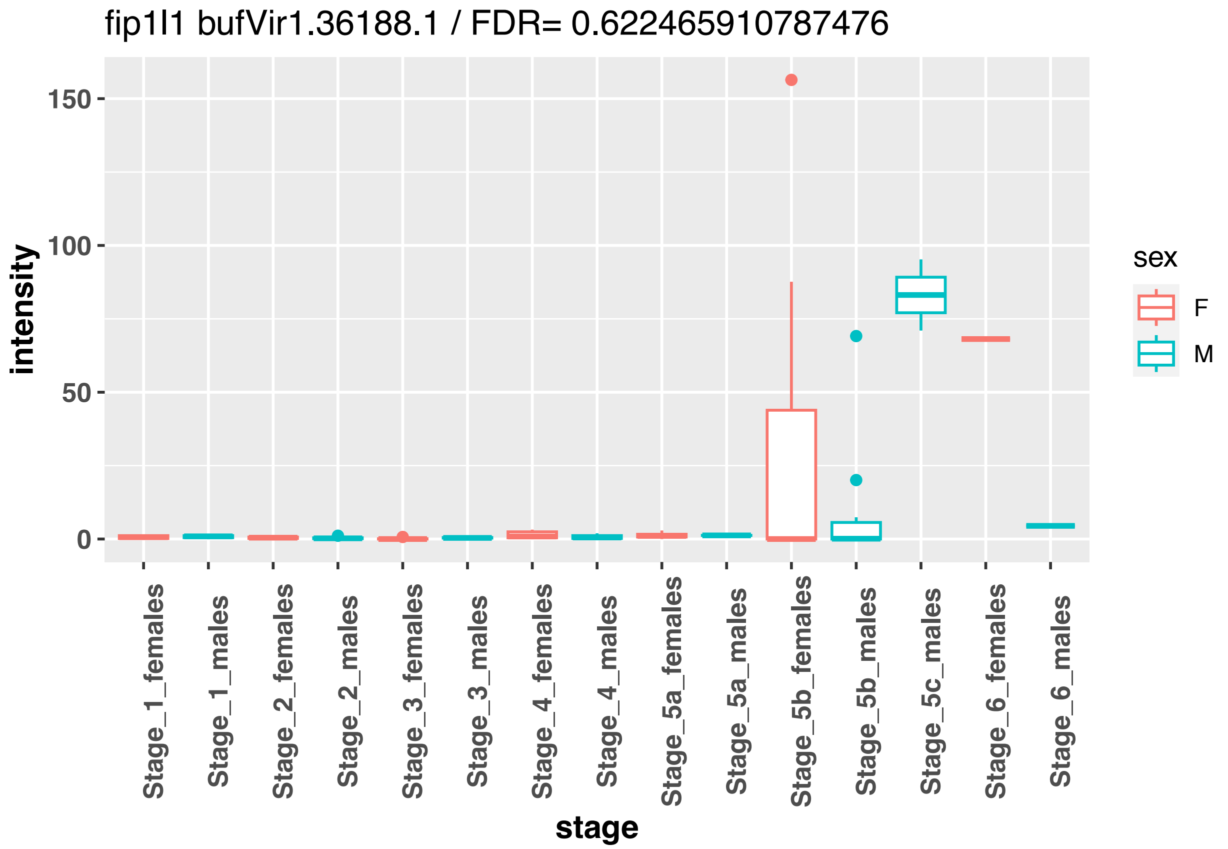
fip1l1 bufVir1.36188.1 / FDR= 0.622465910787476
intensity
150
100
50
0
Stage_1_females
Stage_1_males
Stage_2_females
Stage_2_males
Stage_3_females
Stage_3_males
Stage_4_females
Stage_4_males
Stage_5a_females
Stage_5a_males
Stage_5b_females
Stage_5b_males
Stage_5c_males
Stage_6_females
Stage_6_males
stage
sex
F
M

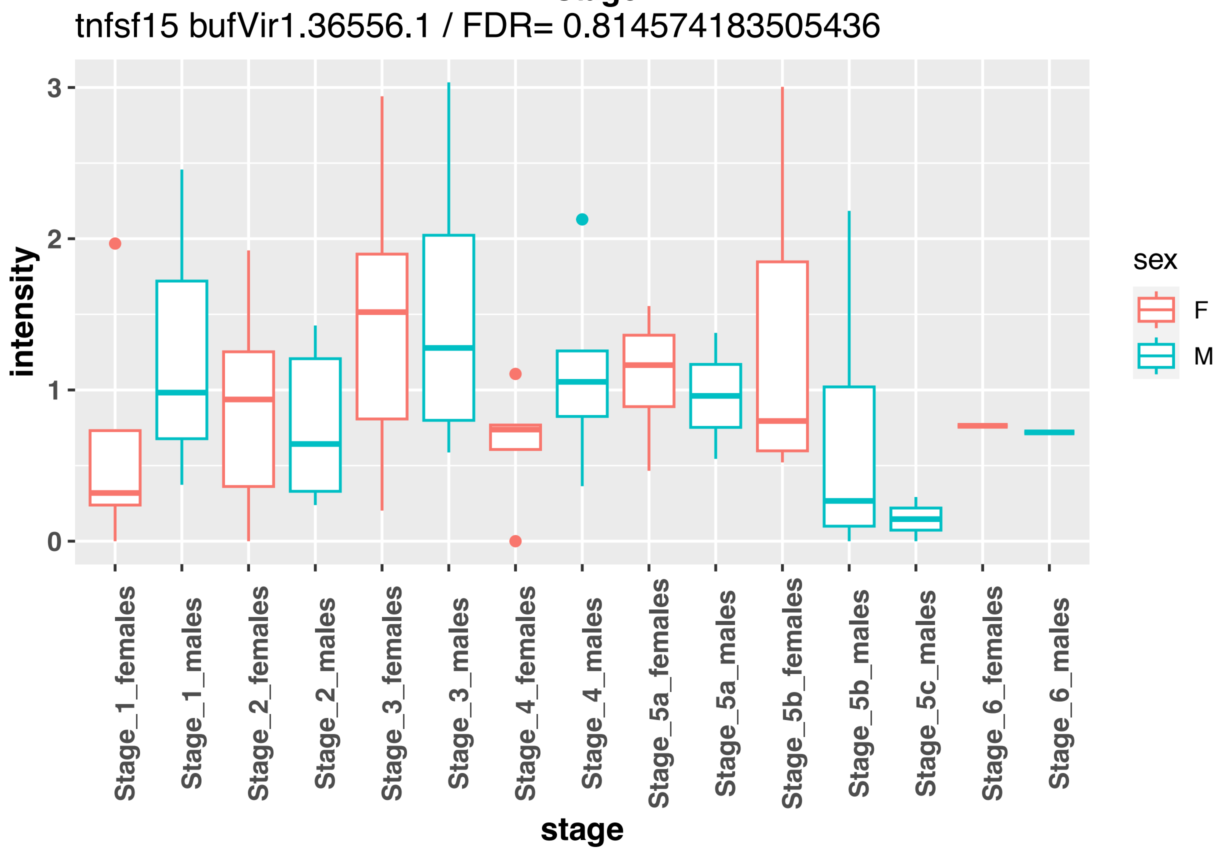
tnfsf15 bufVir1.36556.1 / FDR= 0.814574183505436
intensity
3
2
1
0
Stage_1_females
Stage_1_males
Stage_2_females
Stage_2_males
Stage_3_females
Stage_3_males
Stage_4_females
Stage_4_males
Stage_5a_females
Stage_5a_males
Stage_5b_females
Stage_5b_males
Stage_5c_males
Stage_6_females
Stage_6_males
stage
sex
F
M

grsf1 bufVir1.3712.1 / FDR= 0.999980732933913

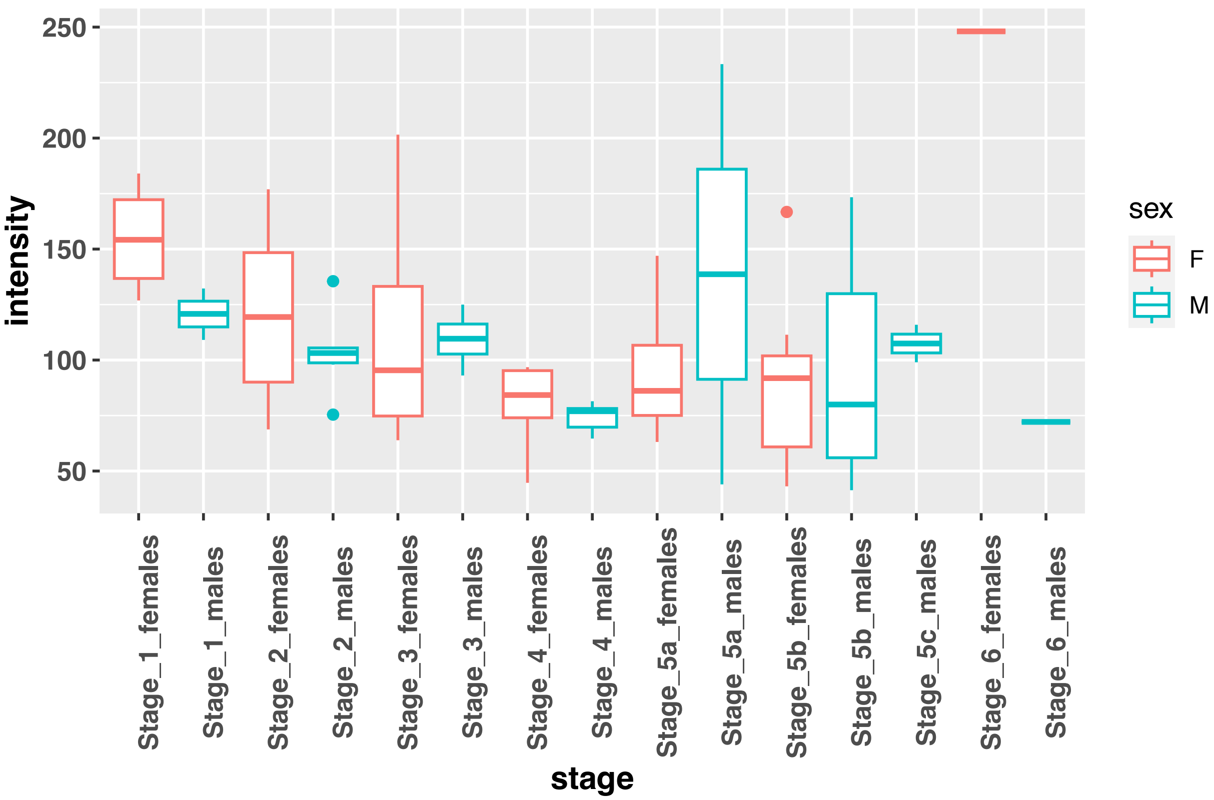


FIP1L1 bufVir1.4660.2 / FDR= 0.956707485324111

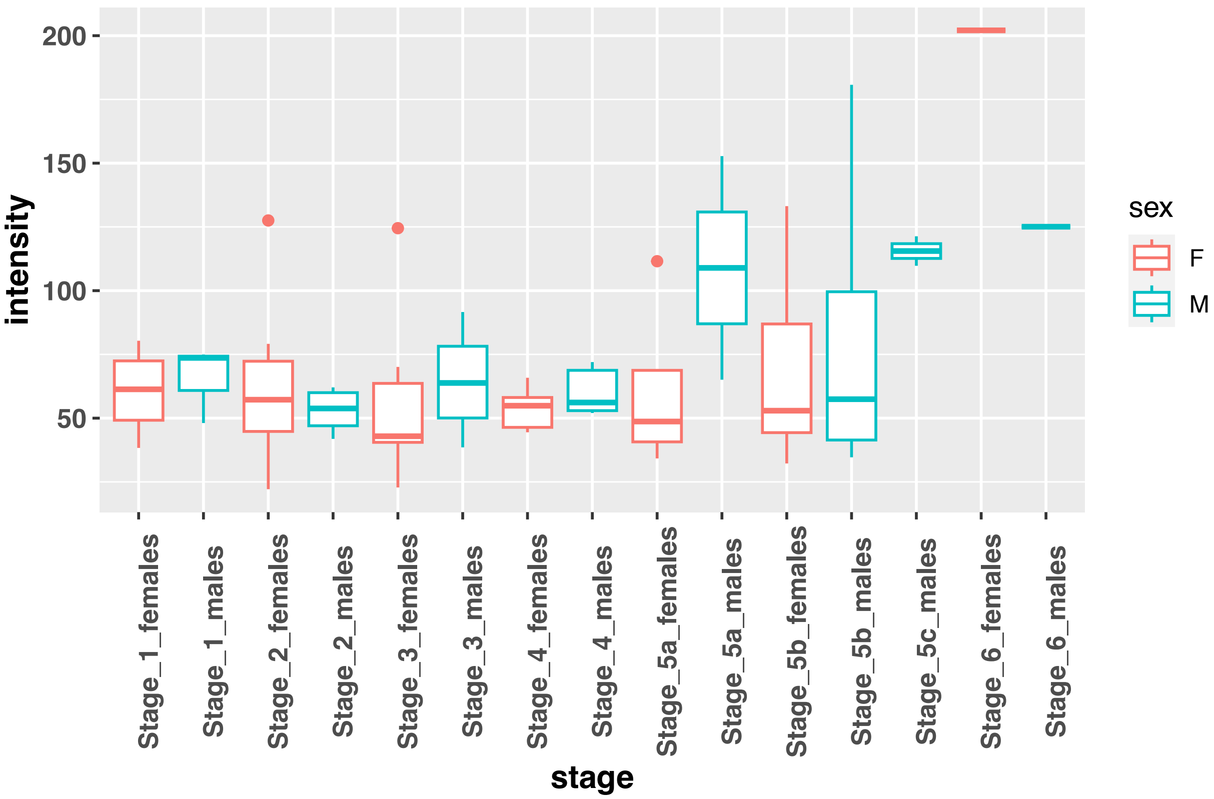

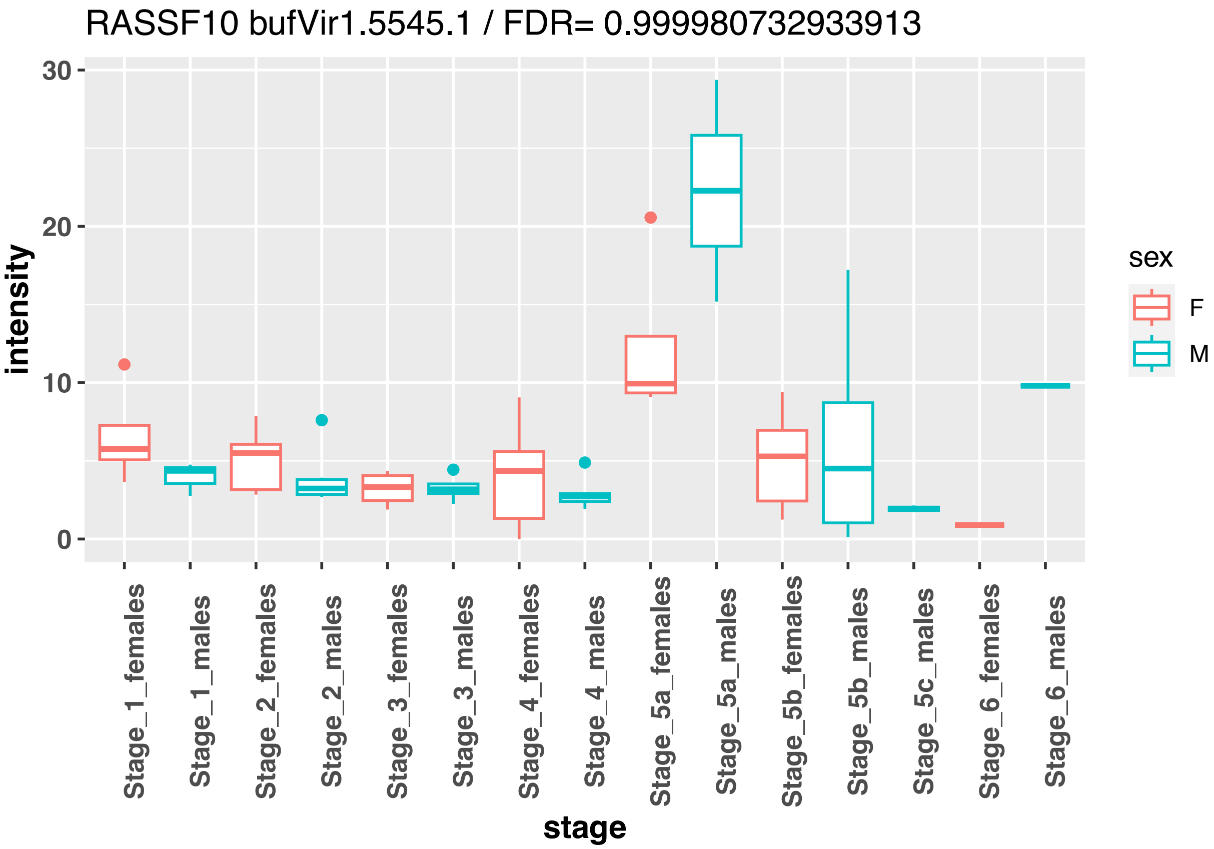
RASSF10 bufVir1.5545.1 / FDR= 0.999980732933913
intensity
30
20
10
0
stage
Stage_1_females
Stage_1_males
Stage_2_females
Stage_2_males
Stage_3_females
Stage_3_males
Stage_4_females
Stage_4_males
Stage_5a_females
Stage_5a_males
Stage_5b_females
Stage_5b_males
Stage_5c_males
Stage_6_females
Stage_6_males
sex
F
M

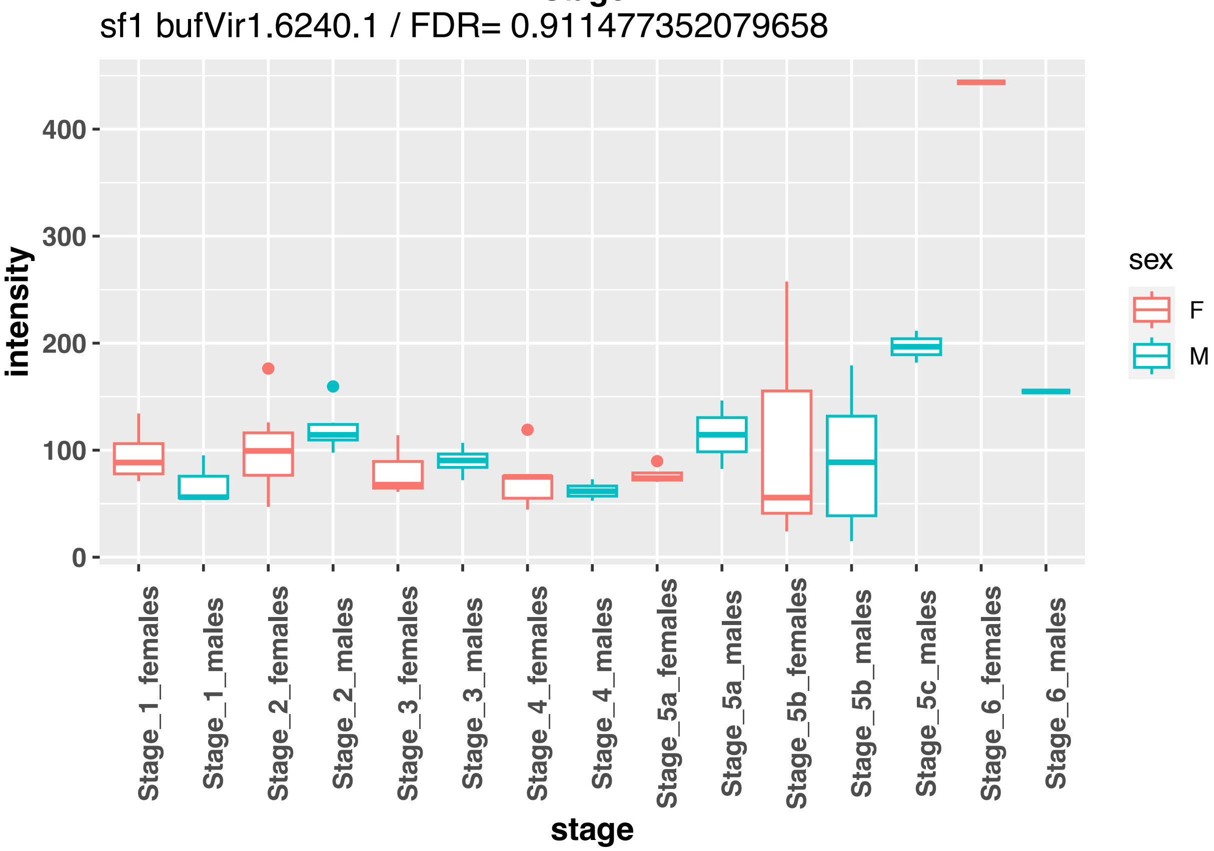
sf1 bufVir1.6240.1 / FDR= 0.911477352079658
intensity
400
300
200
100
0
stage
Stage_1_females
Stage_1_males
Stage_2_females
Stage_2_males
Stage_3_females
Stage_3_males
Stage_4_females
Stage_4_males
Stage_5a_females
Stage_5a_males
Stage_5b_females
Stage_5b_males
Stage_5c_males
Stage_6_females
Stage_6_males
sex
F
M

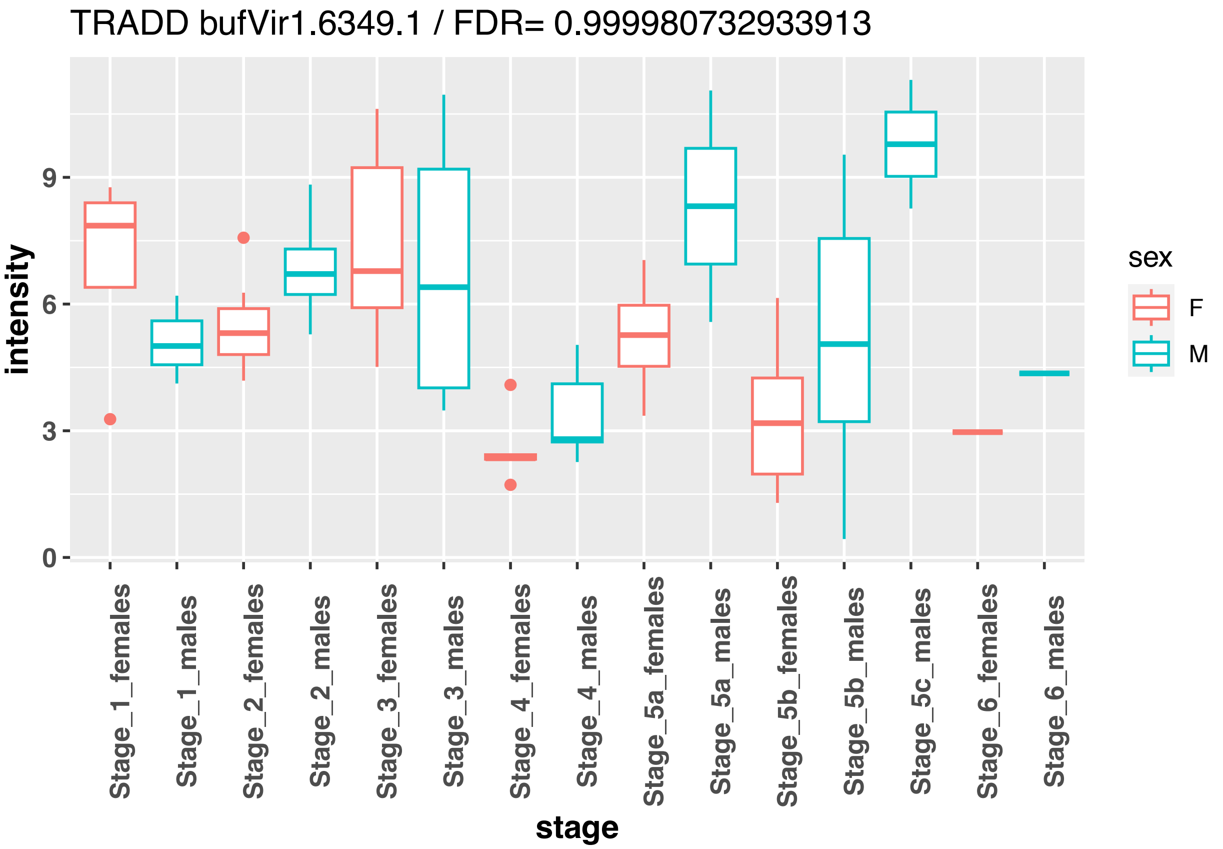

TRADD bufVir1.6349.1 / FDR= 0.999980732933913
intensity
0
3
6
9
stage
Stage_1_females
Stage_1_males
Stage_2_females
Stage_2_males
Stage_3_females
Stage_3_males
Stage_4_females
Stage_4_males
Stage_5a_females
Stage_5a_males
Stage_5b_females
Stage_5b_males
Stage_5c_males
Stage_6_females
Stage_6_males
sex
F
M


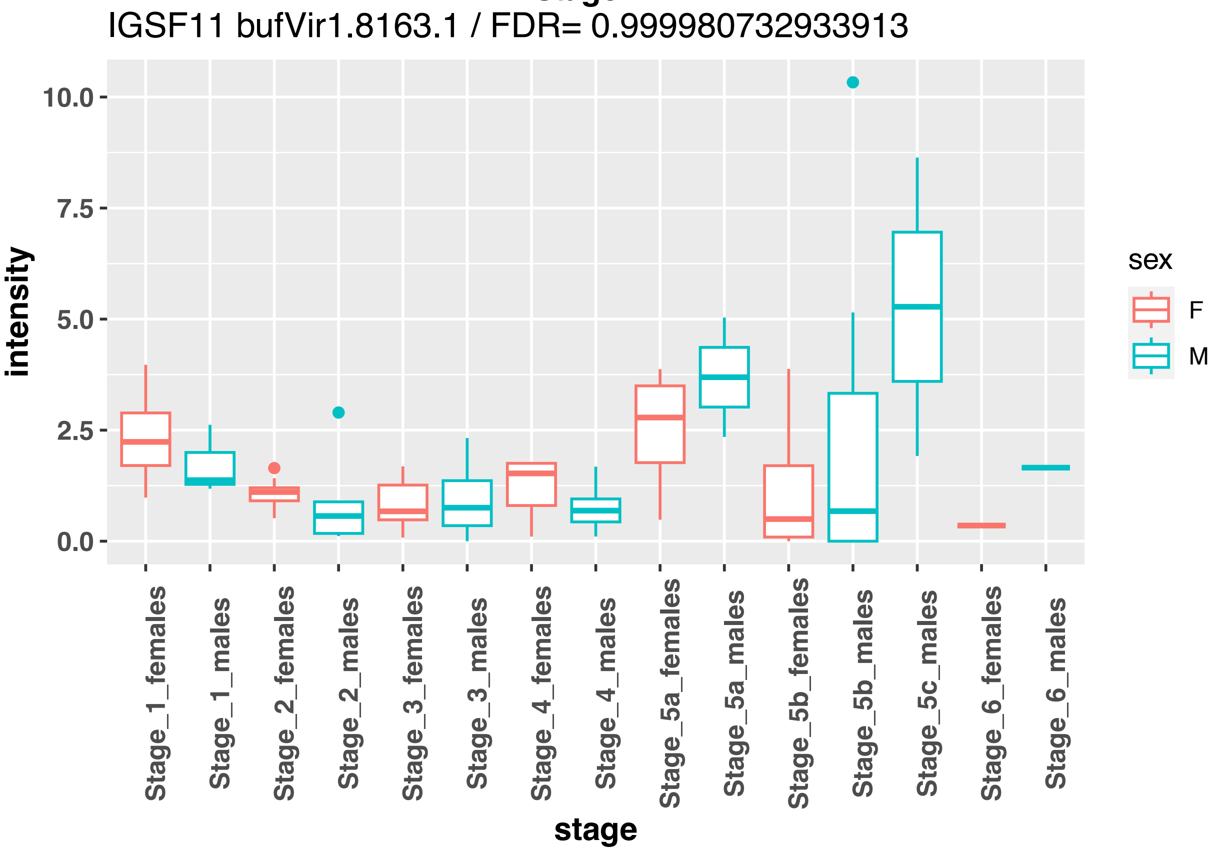

IGSF11 bufVir1.8163.1 / FDR= 0.999980732933913
intensity
0.0
2.5
5.0
7.5
10.0
stage
Stage_1_females
Stage_1_males
Stage_2_females
Stage_2_males
Stage_3_females
Stage_3_males
Stage_4_females
Stage_4_males
Stage_5a_females
Stage_5a_males
Stage_5b_females
Stage_5b_males
Stage_5c_males
Stage_6_females
Stage_6_males
sex
F
M

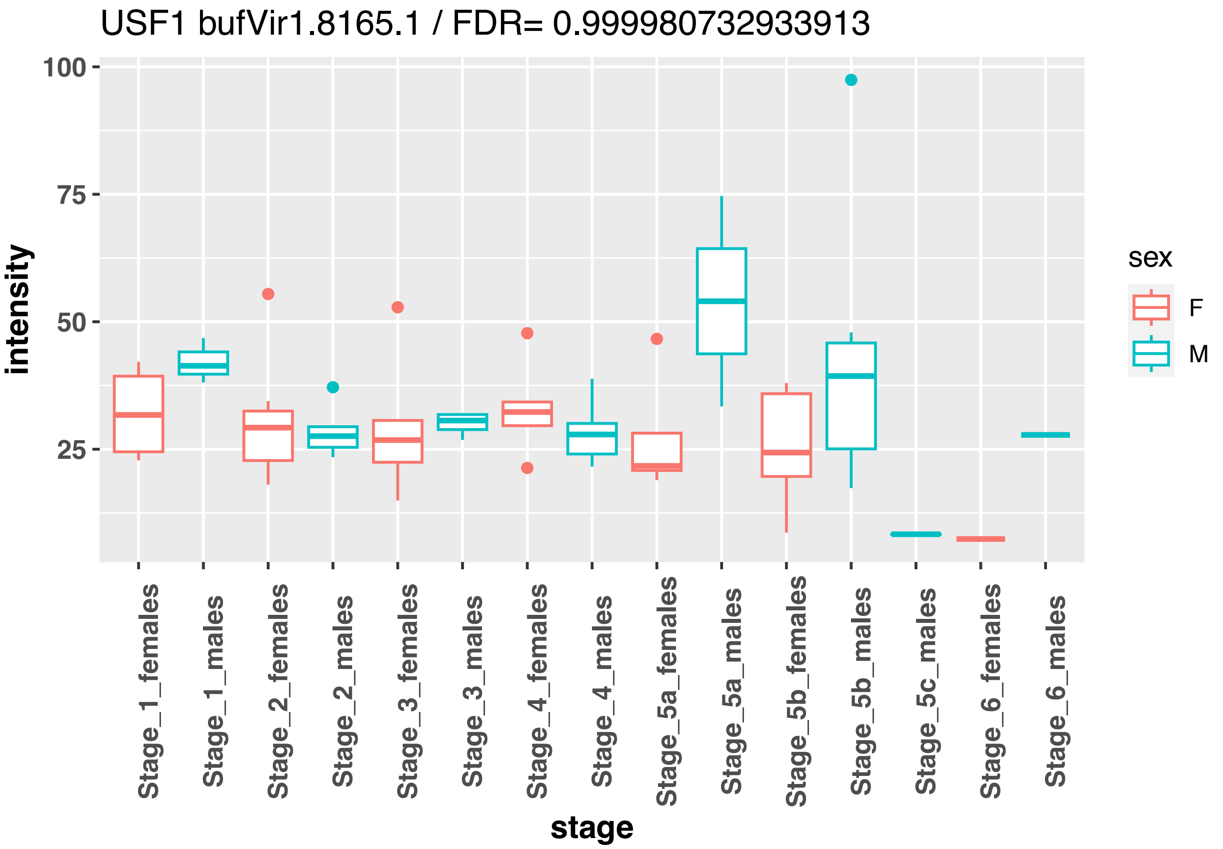
USF1 bufVir1.8165.1 / FDR= 0.999980732933913
100
75
50
25
intensity
Stage_1_females
Stage_1_males
Stage_2_females
Stage_2_males
Stage_3_females
Stage_3_males
Stage_4_females
Stage_4_males
Stage_5a_females
Stage_5a_males
Stage_5b_females
Stage_5b_males
Stage_5c_males
Stage_6_females
Stage_6_males
stage
sex
F
M

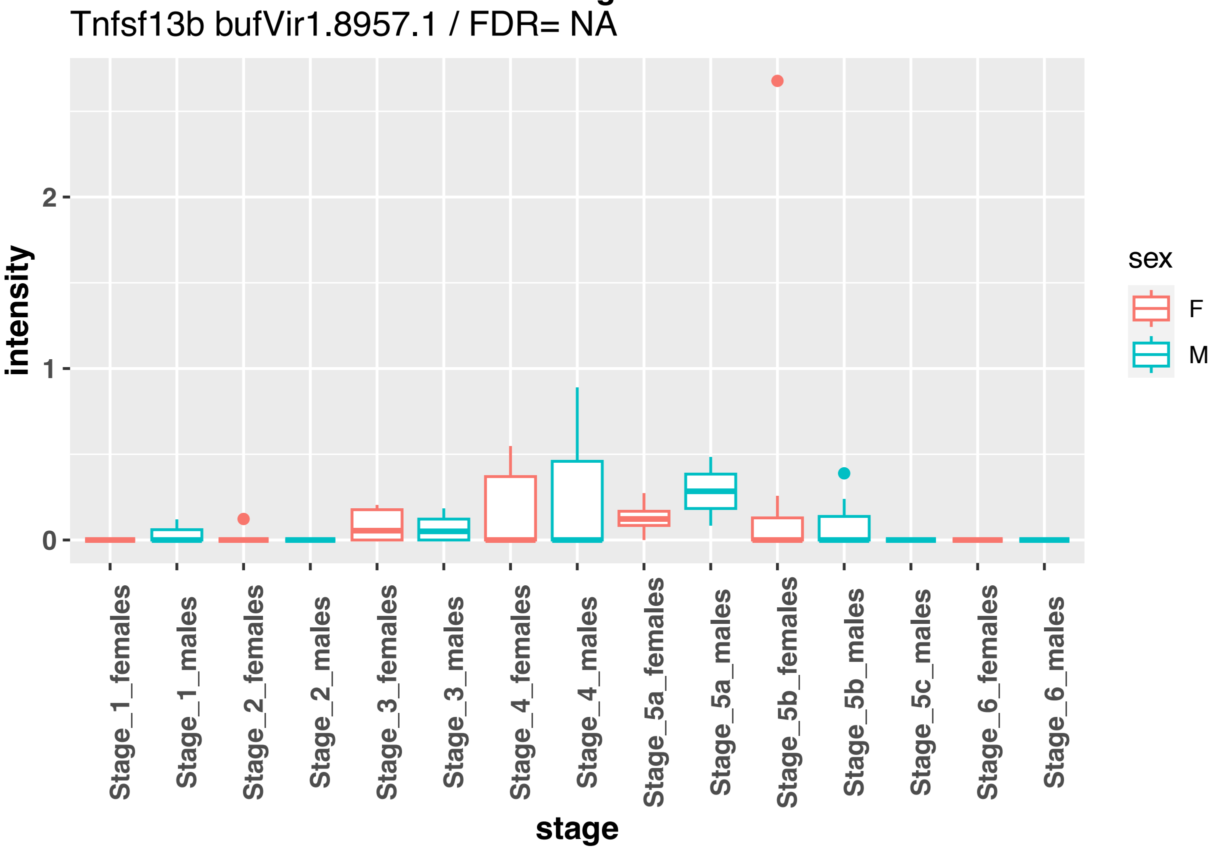
Tnfsf13b bufVir1.8957.1 / FDR= NA
2
1
0
intensity
Stage_1_females
Stage_1_males
Stage_2_females
Stage_2_males
Stage_3_females
Stage_3_males
Stage_4_females
Stage_4_males
Stage_5a_females
Stage_5a_males
Stage_5b_females
Stage_5b_males
Stage_5c_males
Stage_6_females
Stage_6_males
stage
sex
F
M

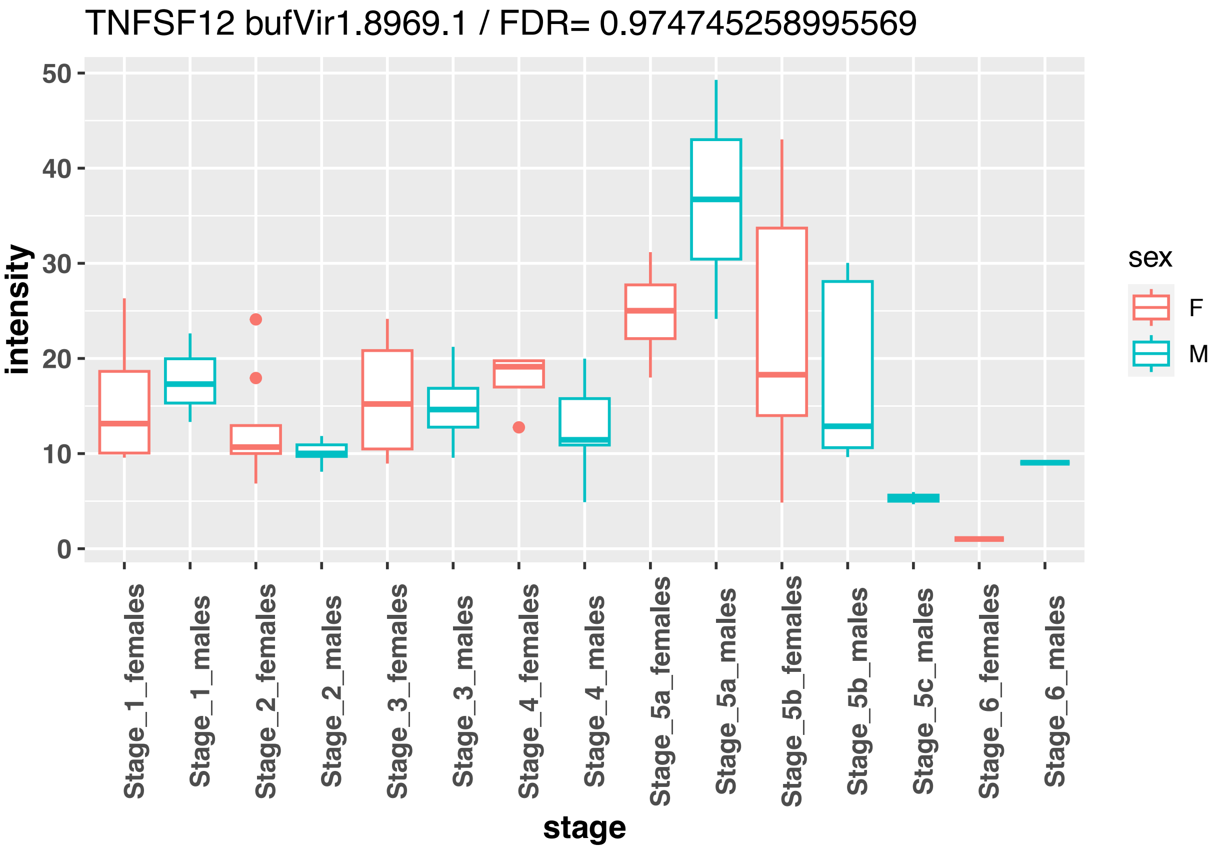
TNFSF12 bufVir1.8969.1 / FDR= 0.974745258995569
intensity
50
40
30
20
10
0
Stage_1_females
Stage_1_males
Stage_2_females
Stage_2_males
Stage_3_females
Stage_3_males
Stage_4_females
Stage_4_males
Stage_5a_females
Stage_5a_males
Stage_5b_females
Stage_5b_males
Stage_5c_males
Stage_6_females
Stage_6_males
stage
sex
F
M

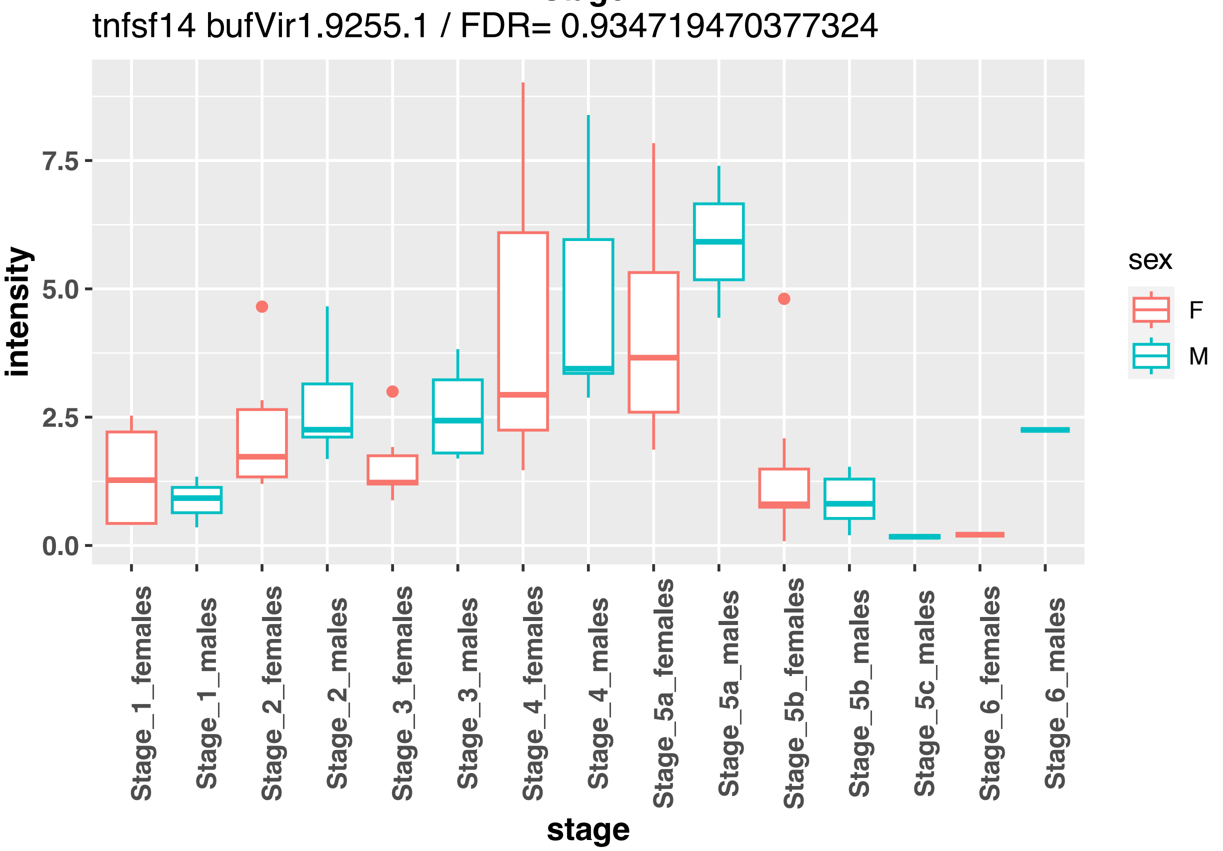
tnfsf14 bufVir1.9255.1 / FDR= 0.934719470377324
intensity
7.5
5.0
2.5
0.0
Stage_1_females
Stage_1_males
Stage_2_females
Stage_2_males
Stage_3_females
Stage_3_males
Stage_4_females
Stage_4_males
Stage_5a_females
Stage_5a_males
Stage_5b_females
Stage_5b_males
Stage_5c_males
Stage_6_females
Stage_6_males
stage
sex
F
M

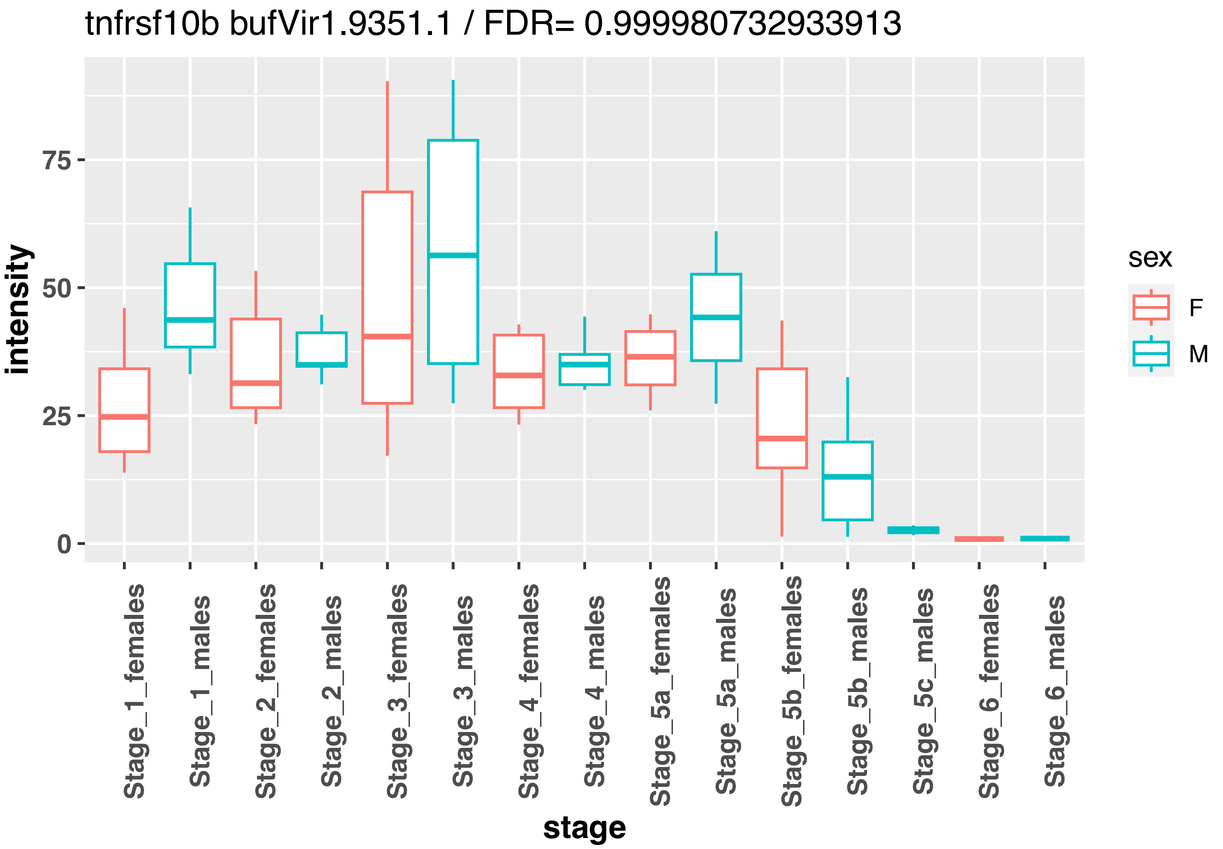
tnfrsf10b bufVir1.9351.1 / FDR= 0.999980732933913
intensity
0
25
50
75
sex
F
M
Stage_1_females
Stage_1_males
Stage_2_females
Stage_2_males
Stage_3_females
Stage_3_males
Stage_4_females
Stage_4_males
Stage_5a_females
Stage_5a_males
Stage_5b_females
Stage_5b_males
Stage_5c_males
Stage_6_females
Stage_6_males
stage

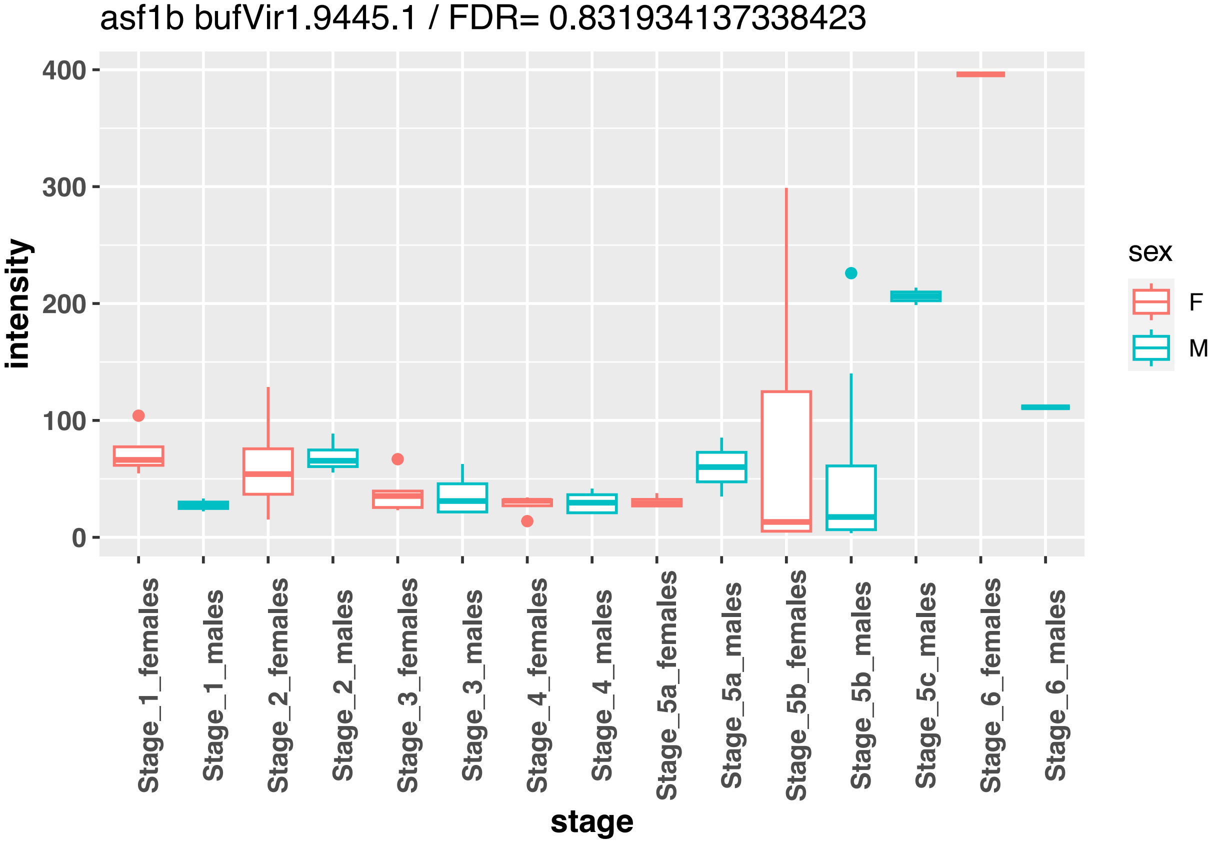
asf1b bufVir1.9445.1 / FDR= 0.831934137338423
intensity
0
100
200
300
400
sex
F
M
Stage_1_females
Stage_1_males
Stage_2_females
Stage_2_males
Stage_3_females
Stage_3_males
Stage_4_females
Stage_4_males
Stage_5a_females
Stage_5a_males
Stage_5b_females
Stage_5b_males
Stage_5c_males
Stage_6_females
Stage_6_males
stage

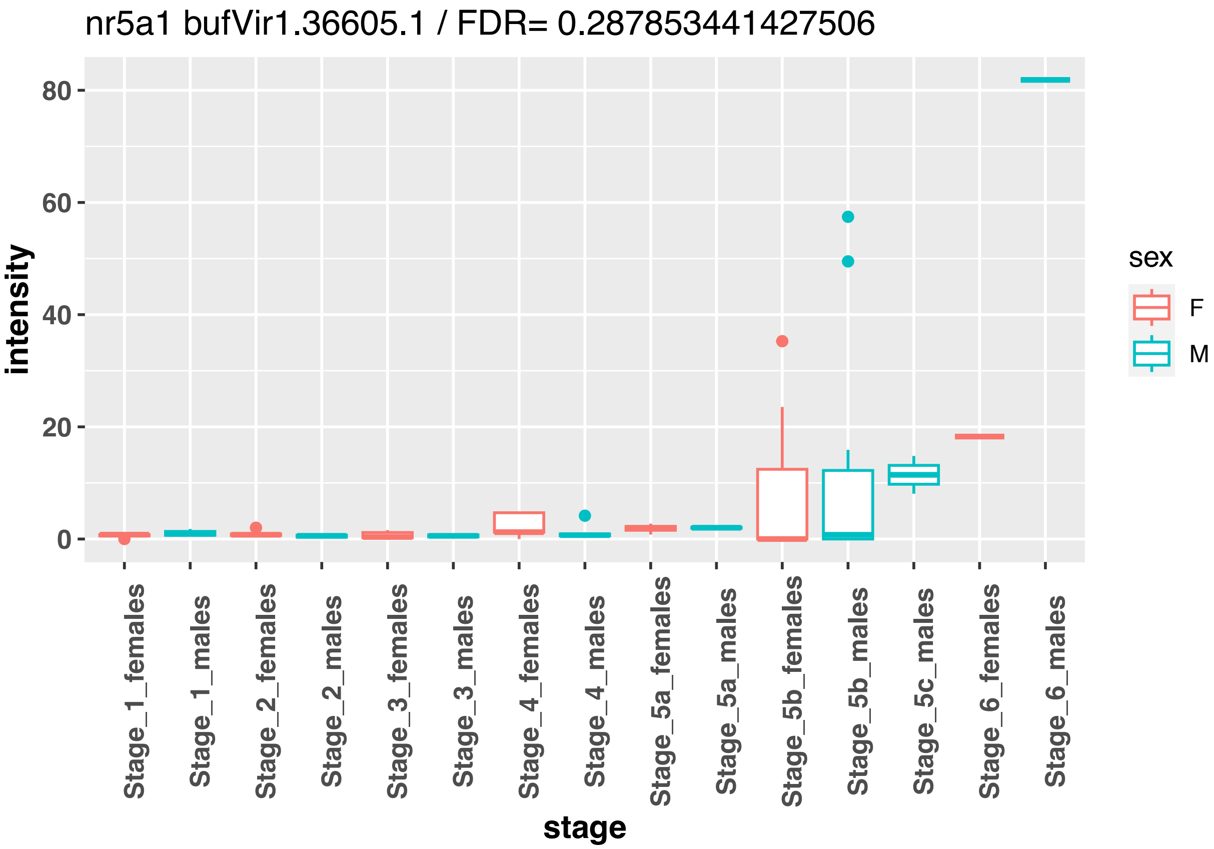
nr5a1 bufVir1.36605.1 / FDR= 0.287853441427506
intensity
0
20
40
60
80
Stage_1_females
Stage_1_males
Stage_2_females
Stage_2_males
Stage_3_females
Stage_3_males
Stage_4_females
Stage_4_males
Stage_5a_females
Stage_5a_males
Stage_5b_females
Stage_5b_males
Stage_5c_males
Stage_6_females
Stage_6_males
stage
sex
F
M

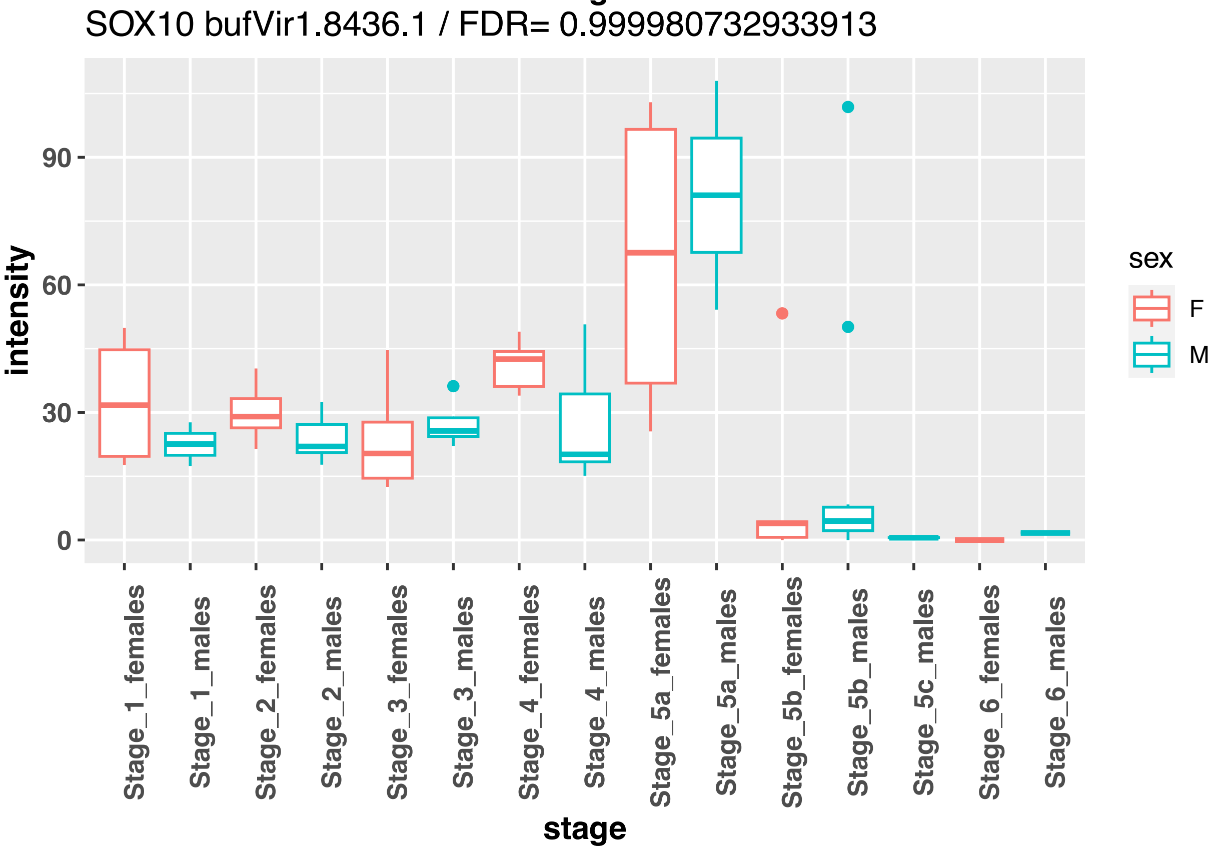
SOX10 bufVir1.8436.1 / FDR= 0.999980732933913
intensity
0
30
60
90
Stage_1_females
Stage_1_males
Stage_2_females
Stage_2_males
Stage_3_females
Stage_3_males
Stage_4_females
Stage_4_males
Stage_5a_females
Stage_5a_males
Stage_5b_females
Stage_5b_males
Stage_5c_males
Stage_6_females
Stage_6_males
stage
sex
F
M

sox17a bufVir1.25154.1 / FDR= 0.907967094460556

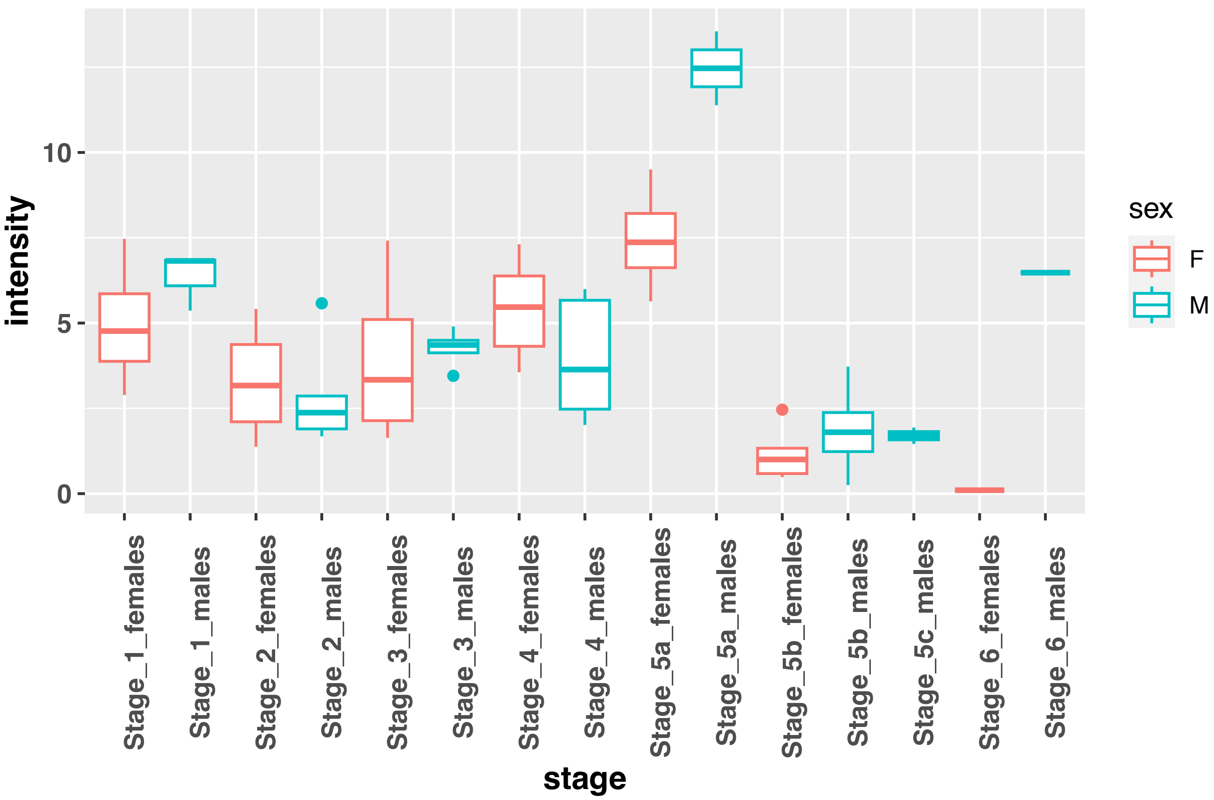


sox17a bufVir1.25155.1 / FDR= 0.530787821700476

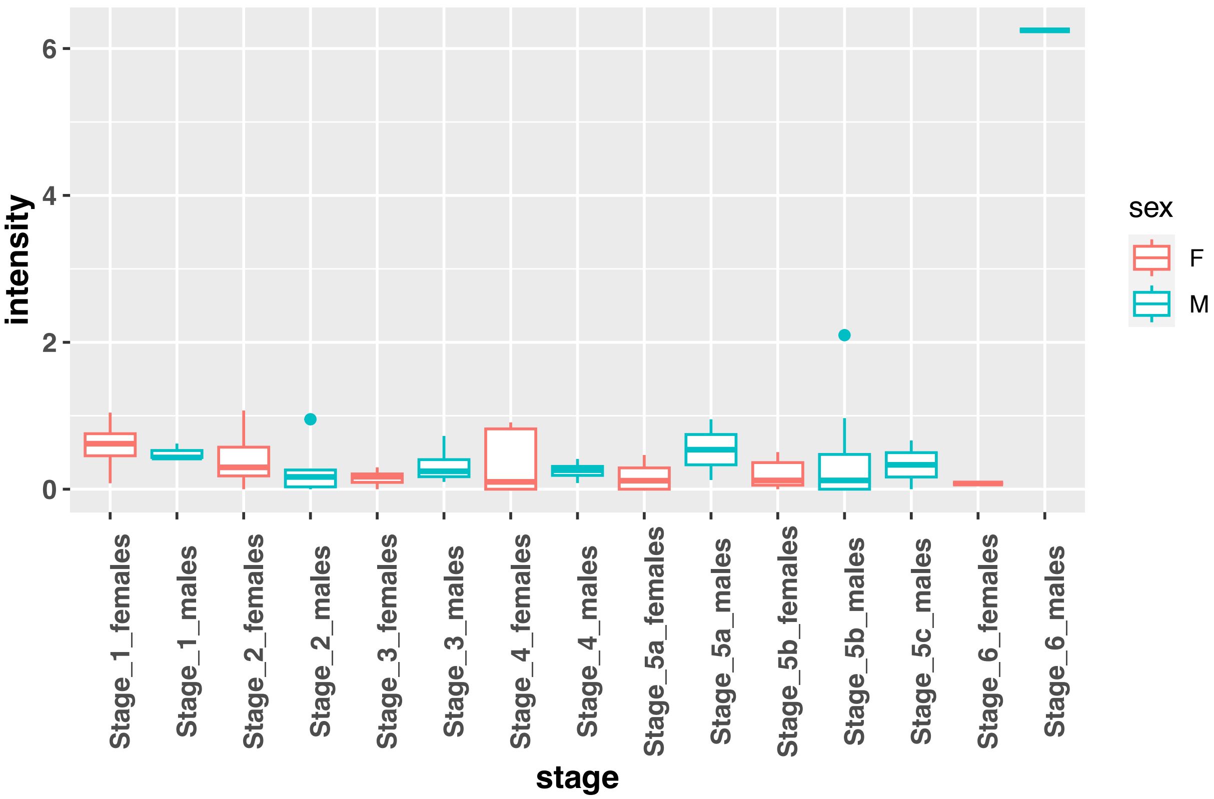

Sox30 bufVir1.12521.1 / FDR= 0.0251154392826941

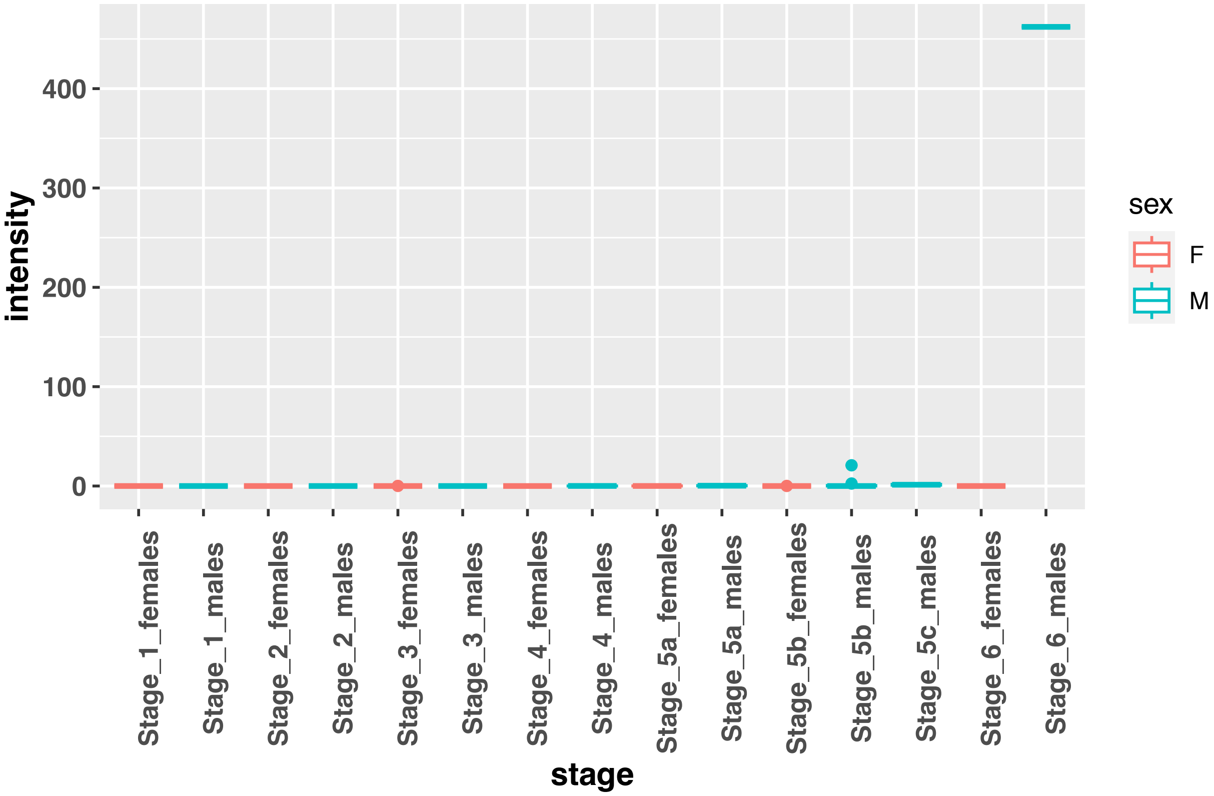


SOX30 bufVir1.3569.1 / FDR= 0.754801944156059

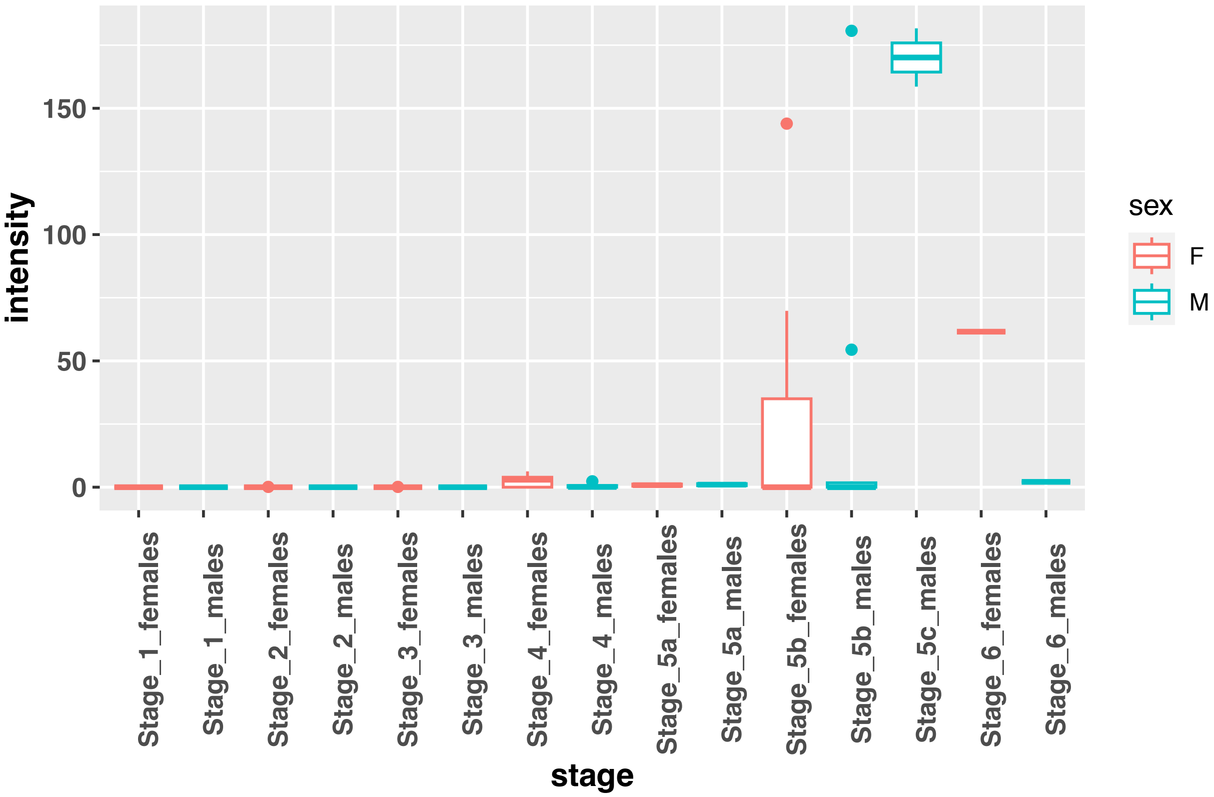

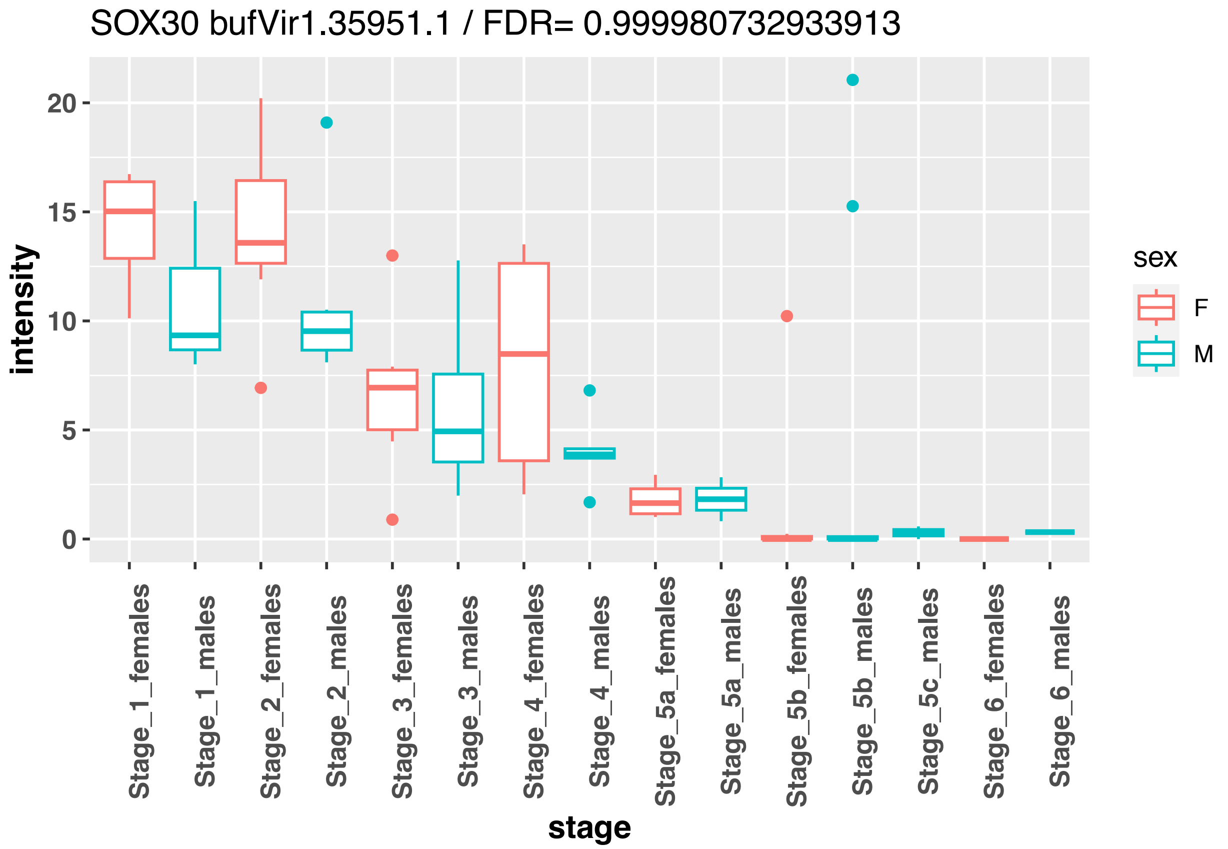
SOX30 bufVir1.35951.1 / FDR= 0.999980732933913
intensity
stage
sex
F
M
Stage_1_females
Stage_1_males
Stage_2_females
Stage_2_males
Stage_3_females
Stage_3_males
Stage_4_females
Stage_4_males
Stage_5a_females
Stage_5a_males
Stage_5b_females
Stage_5b_males
Stage_5c_males
Stage_6_females
Stage_6_males

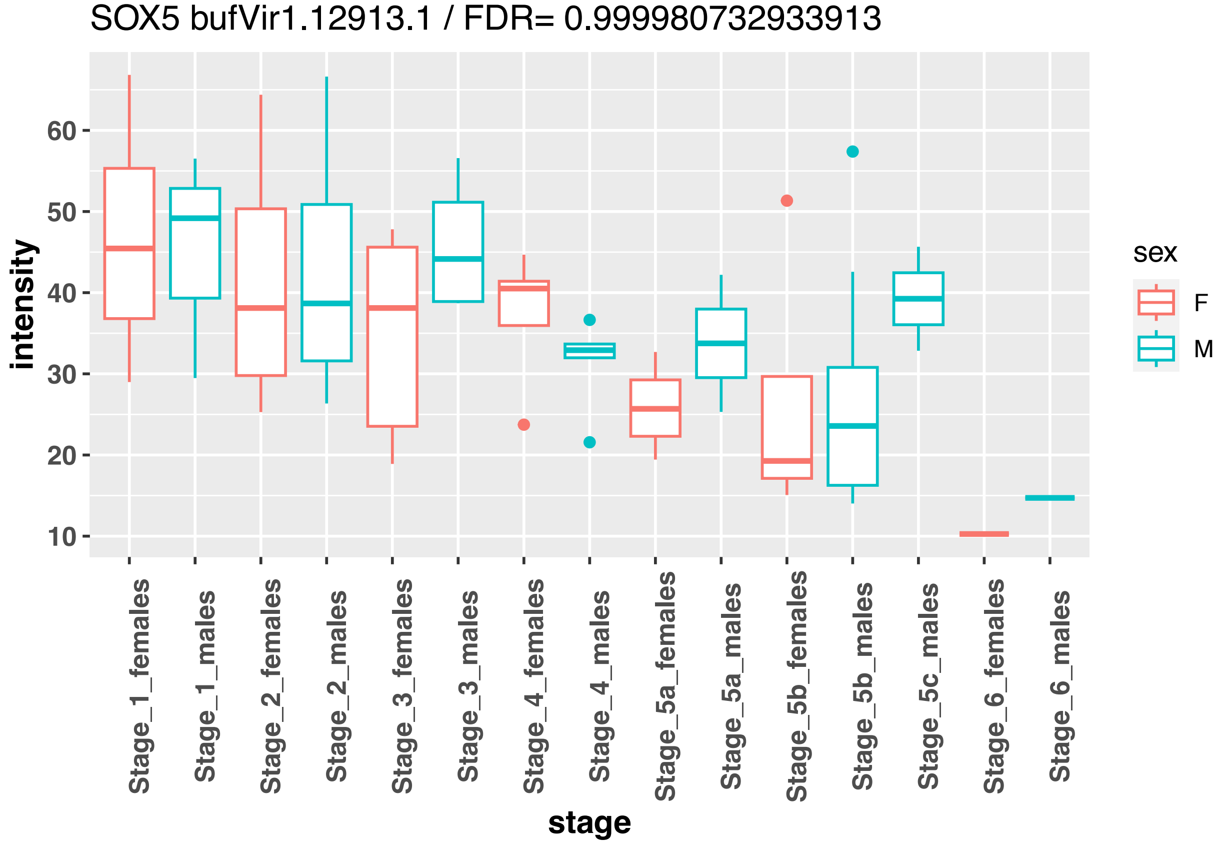
SOX5 bufVir1.12913.1 / FDR= 0.999980732933913
intensity
stage
sex
F
M
Stage_1_females
Stage_1_males
Stage_2_females
Stage_2_males
Stage_3_females
Stage_3_males
Stage_4_females
Stage_4_males
Stage_5a_females
Stage_5a_males
Stage_5b_females
Stage_5b_males
Stage_5c_males
Stage_6_females
Stage_6_males

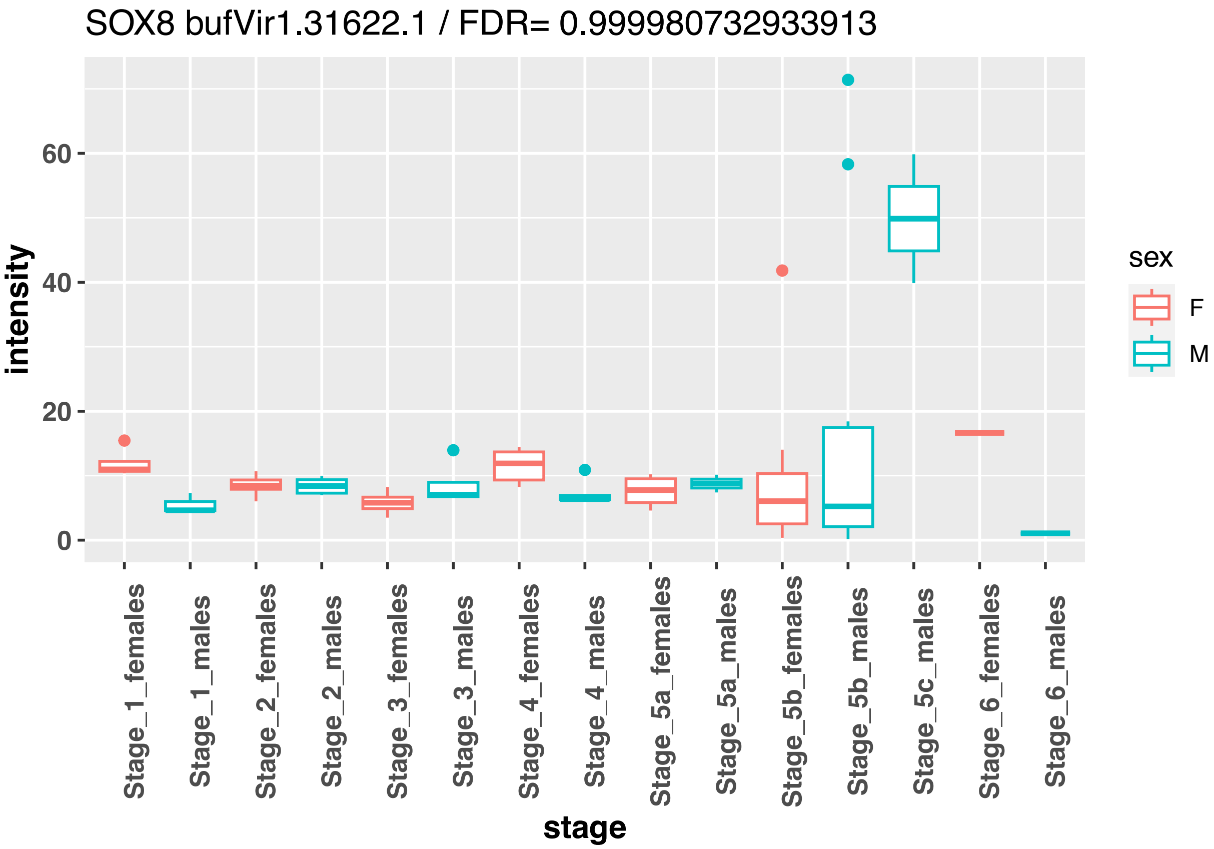
SOX8 bufVir1.31622.1 / FDR= 0.999980732933913
intensity
stage
sex
F
M
Stage_1_females
Stage_1_males
Stage_2_females
Stage_2_males
Stage_3_females
Stage_3_males
Stage_4_females
Stage_4_males
Stage_5a_females
Stage_5a_males
Stage_5b_females
Stage_5b_males
Stage_5c_males
Stage_6_females
Stage_6_males

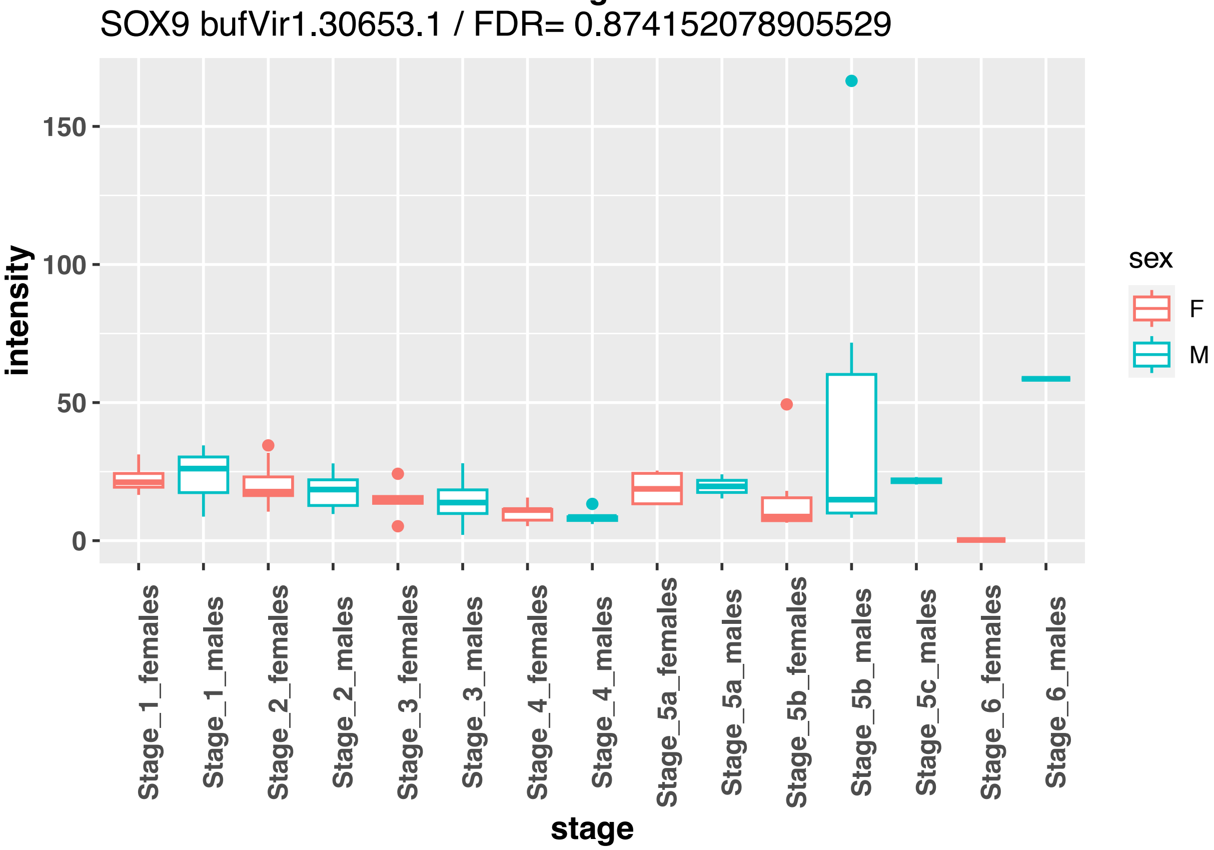
SOX9 bufVir1.30653.1 / FDR= 0.874152078905529
intensity
stage
sex
F
M
Stage_1_females
Stage_1_males
Stage_2_females
Stage_2_males
Stage_3_females
Stage_3_males
Stage_4_females
Stage_4_males
Stage_5a_females
Stage_5a_males
Stage_5b_females
Stage_5b_males
Stage_5c_males
Stage_6_females
Stage_6_males

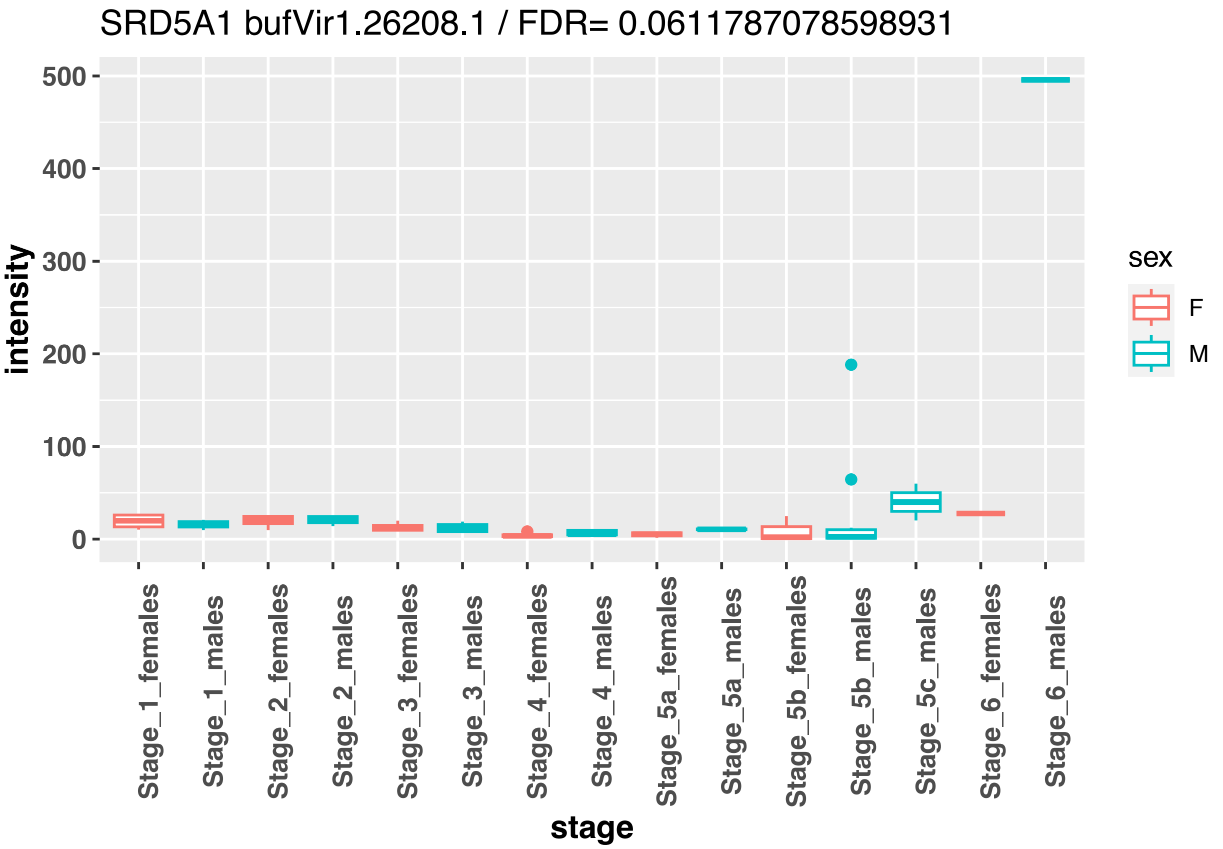
SRD5A1 bufVir1.26208.1 / FDR= 0.0611787078598931
intensity
stage
sex
F
M
0
100
200
300
400
500
Stage_1_females
Stage_1_males
Stage_2_females
Stage_2_males
Stage_3_females
Stage_3_males
Stage_4_females
Stage_4_males
Stage_5a_females
Stage_5a_males
Stage_5b_females
Stage_5b_males
Stage_5c_males
Stage_6_females
Stage_6_males

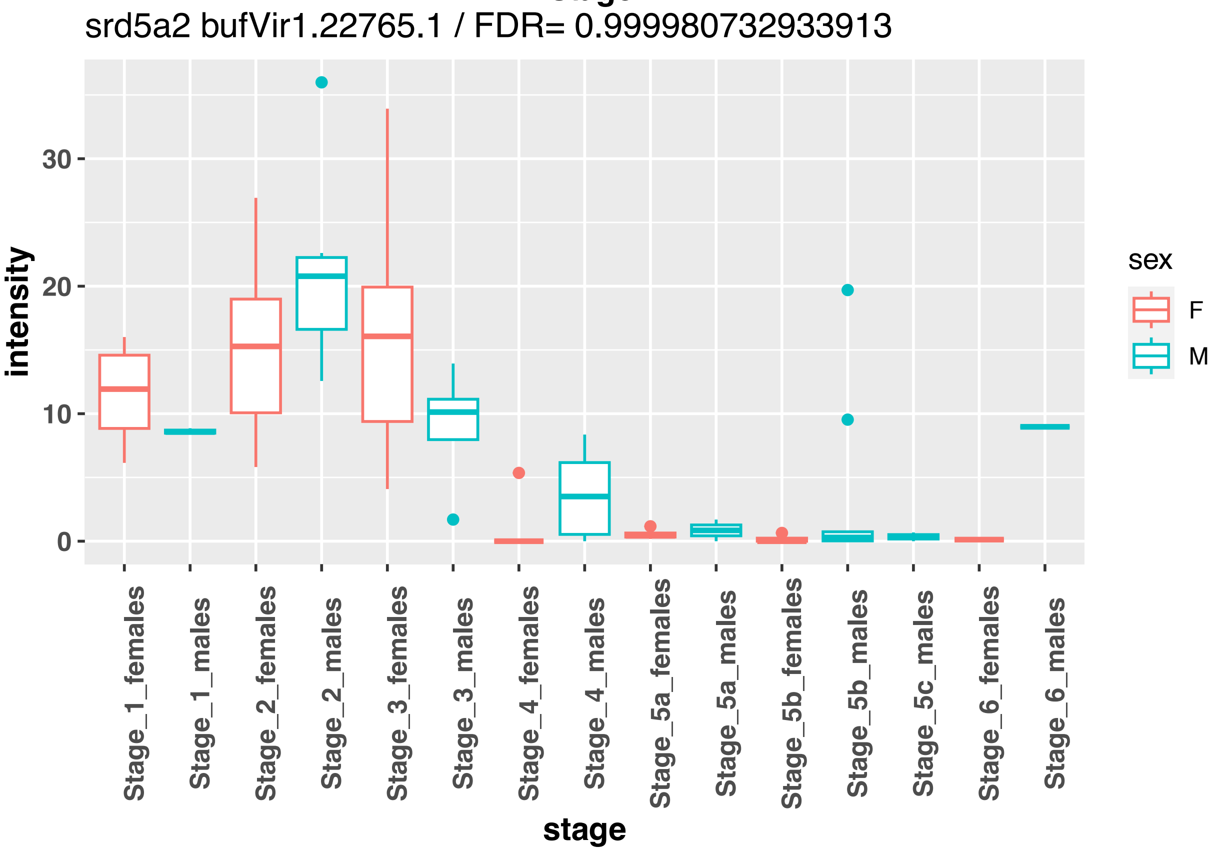
srd5a2 bufVir1.22765.1 / FDR= 0.999980732933913
intensity
stage
sex
F
M
0
10
20
30
Stage_1_females
Stage_1_males
Stage_2_females
Stage_2_males
Stage_3_females
Stage_3_males
Stage_4_females
Stage_4_males
Stage_5a_females
Stage_5a_males
Stage_5b_females
Stage_5b_males
Stage_5c_males
Stage_6_females
Stage_6_males

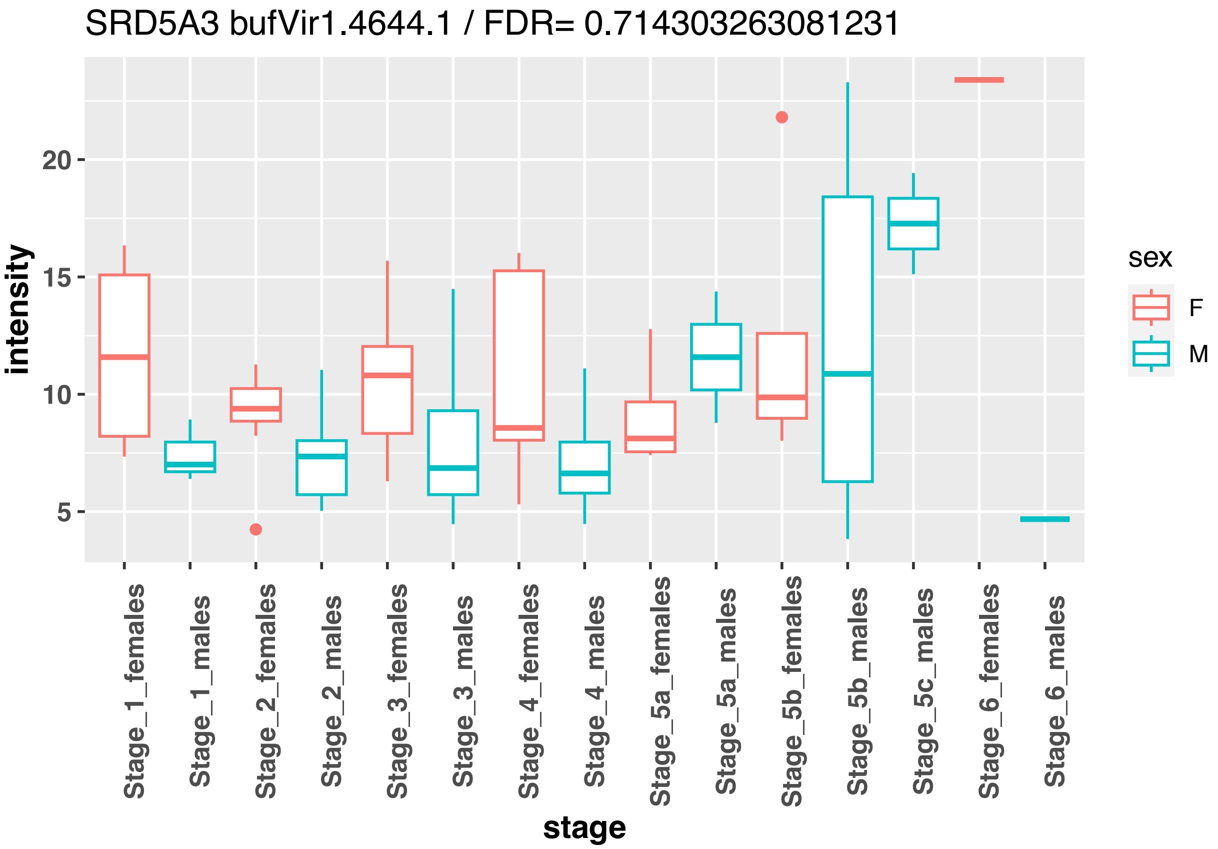
SRD5A3 bufVir1.4644.1 / FDR= 0.714303263081231
intensity
stage
sex
F
M
Stage_1_females
Stage_1_males
Stage_2_females
Stage_2_males
Stage_3_females
Stage_3_males
Stage_4_females
Stage_4_males
Stage_5a_females
Stage_5a_males
Stage_5b_females
Stage_5b_males
Stage_5c_males
Stage_6_females
Stage_6_males

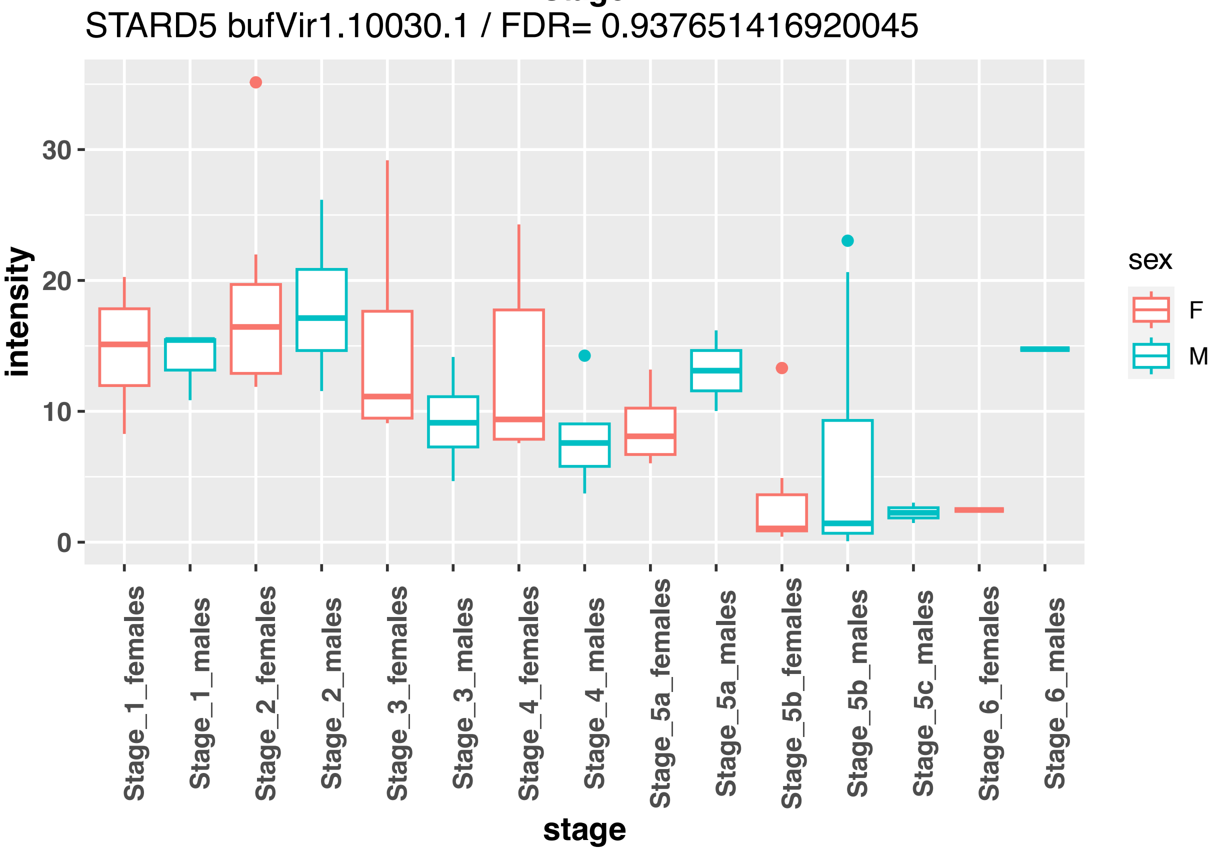
STARD5 bufVir1.10030.1 / FDR= 0.937651416920045
intensity
stage
sex
F
M
Stage_1_females
Stage_1_males
Stage_2_females
Stage_2_males
Stage_3_females
Stage_3_males
Stage_4_females
Stage_4_males
Stage_5a_females
Stage_5a_males
Stage_5b_females
Stage_5b_males
Stage_5c_males
Stage_6_females
Stage_6_males

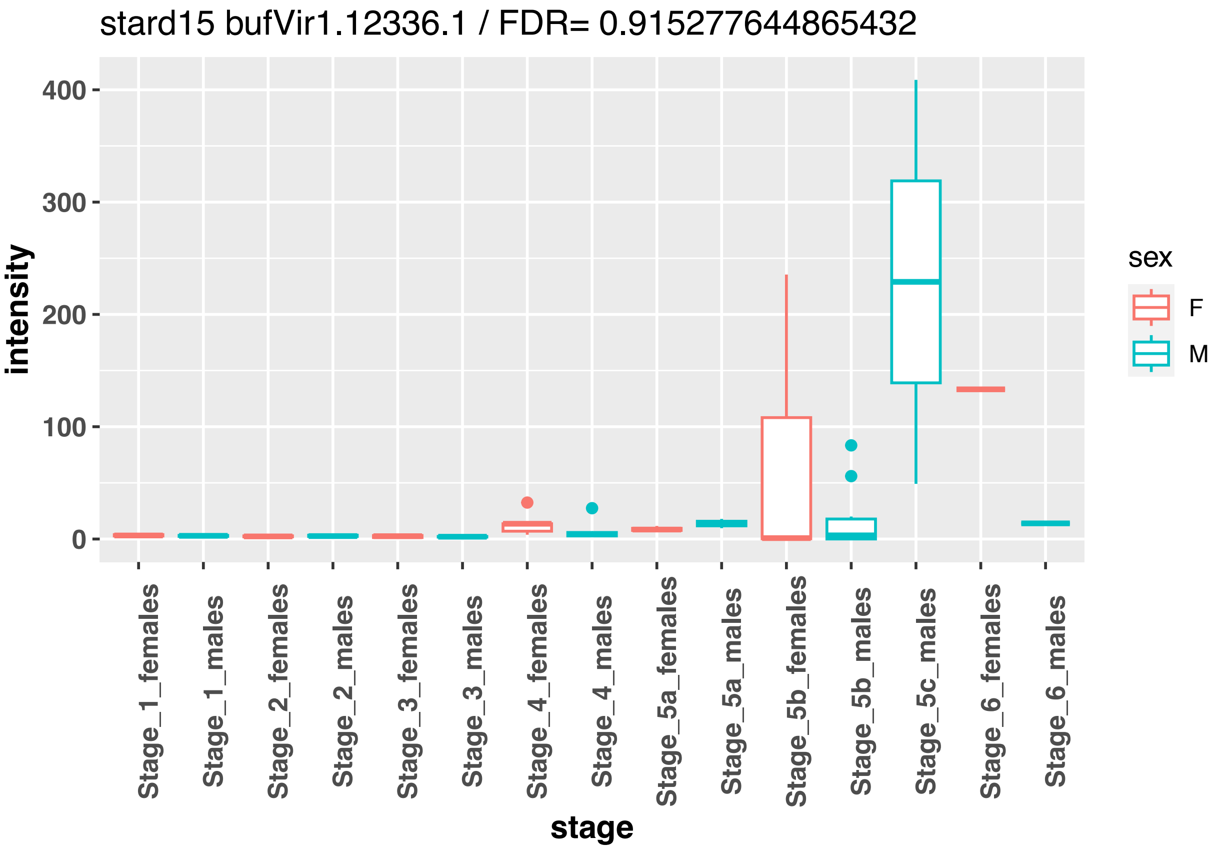
stard15 bufVir1.12336.1 / FDR= 0.915277644865432
intensity
400
300
200
100
0
stage
Stage_1_females
Stage_1_males
Stage_2_females
Stage_2_males
Stage_3_females
Stage_3_males
Stage_4_females
Stage_4_males
Stage_5a_females
Stage_5a_males
Stage_5b_females
Stage_5b_males
Stage_5c_males
Stage_6_females
Stage_6_males
sex
F
M

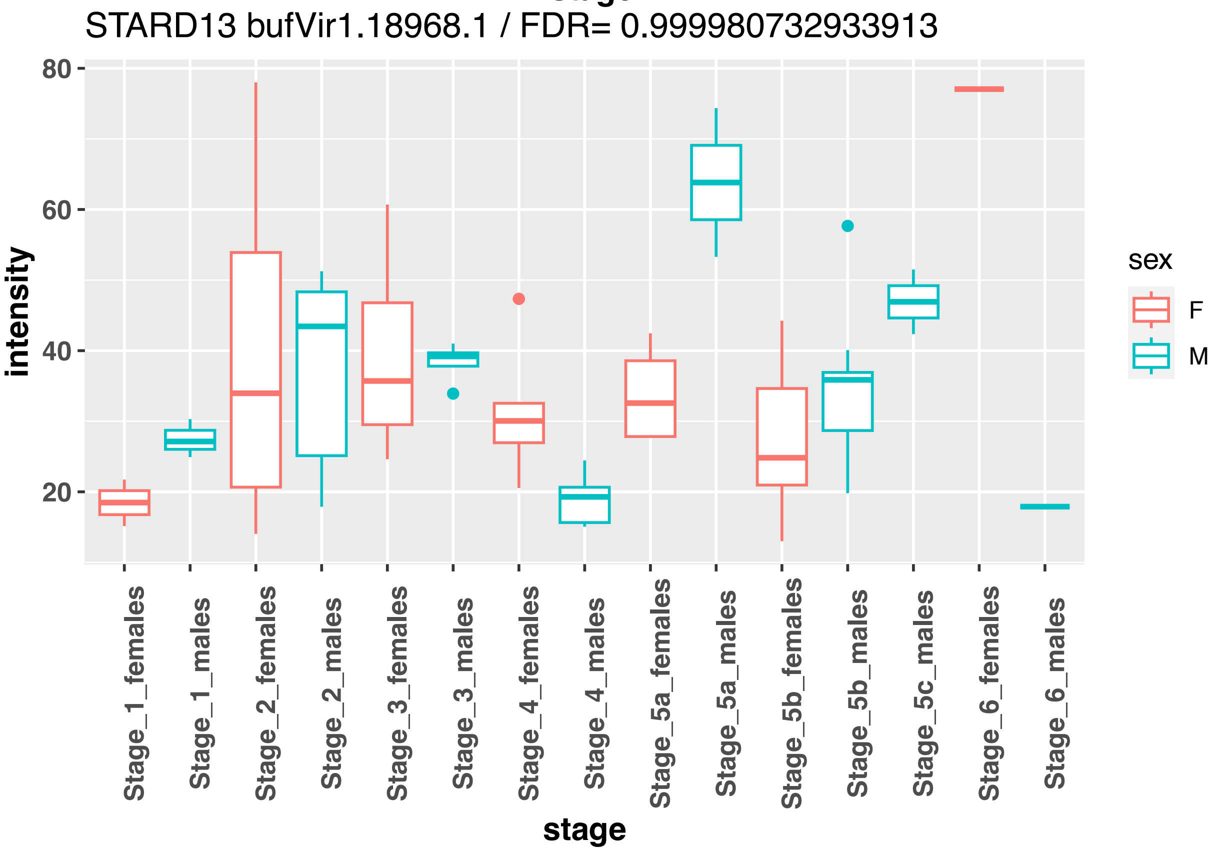
STARD13 bufVir1.18968.1 / FDR= 0.999980732933913
intensity
80
60
40
20
stage
Stage_1_females
Stage_1_males
Stage_2_females
Stage_2_males
Stage_3_females
Stage_3_males
Stage_4_females
Stage_4_males
Stage_5a_females
Stage_5a_males
Stage_5b_females
Stage_5b_males
Stage_5c_males
Stage_6_females
Stage_6_males
sex
F
M

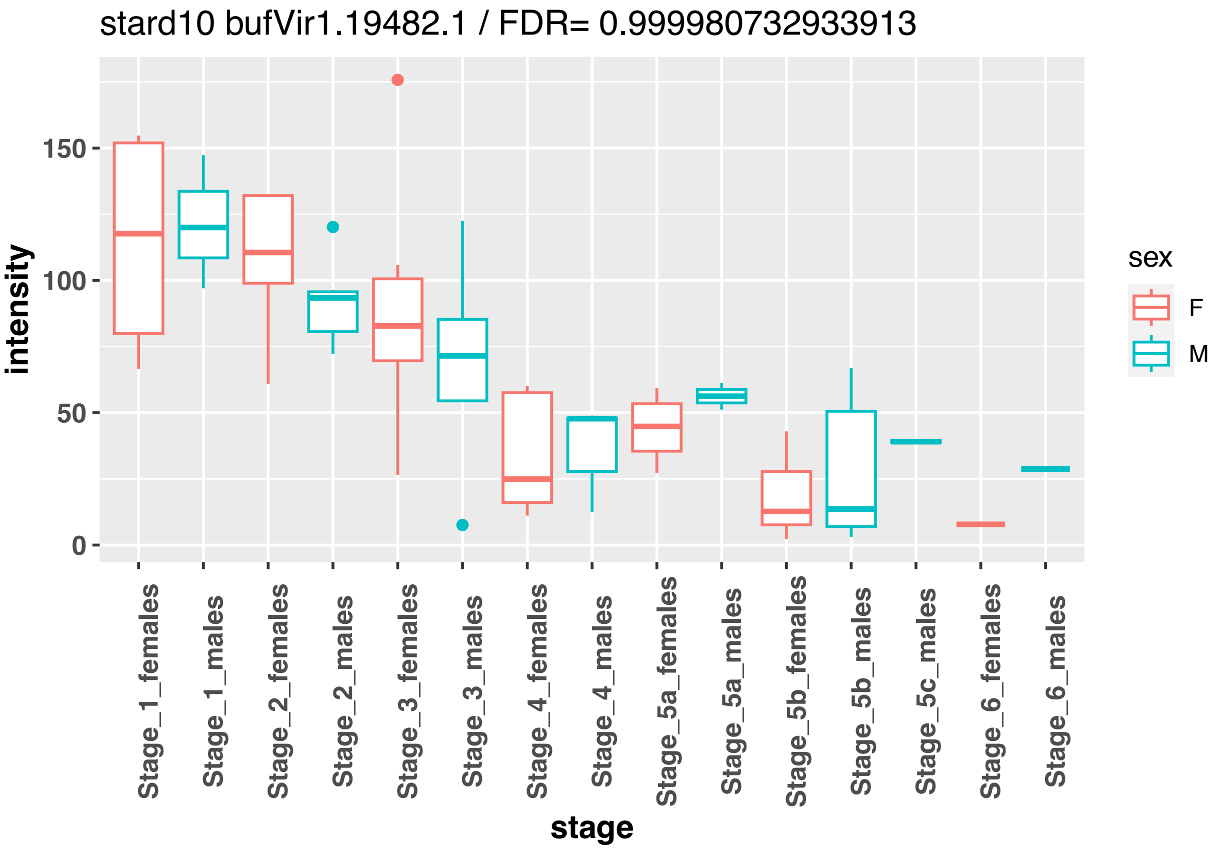
stard10 bufVir1.19482.1 / FDR= 0.999980732933913
intensity
150
100
50
0
Stage_1_females
Stage_1_males
Stage_2_females
Stage_2_males
Stage_3_females
Stage_3_males
Stage_4_females
Stage_4_males
Stage_5a_females
Stage_5a_males
Stage_5b_females
Stage_5b_males
Stage_5c_males
Stage_6_females
Stage_6_males
stage
sex
F
M

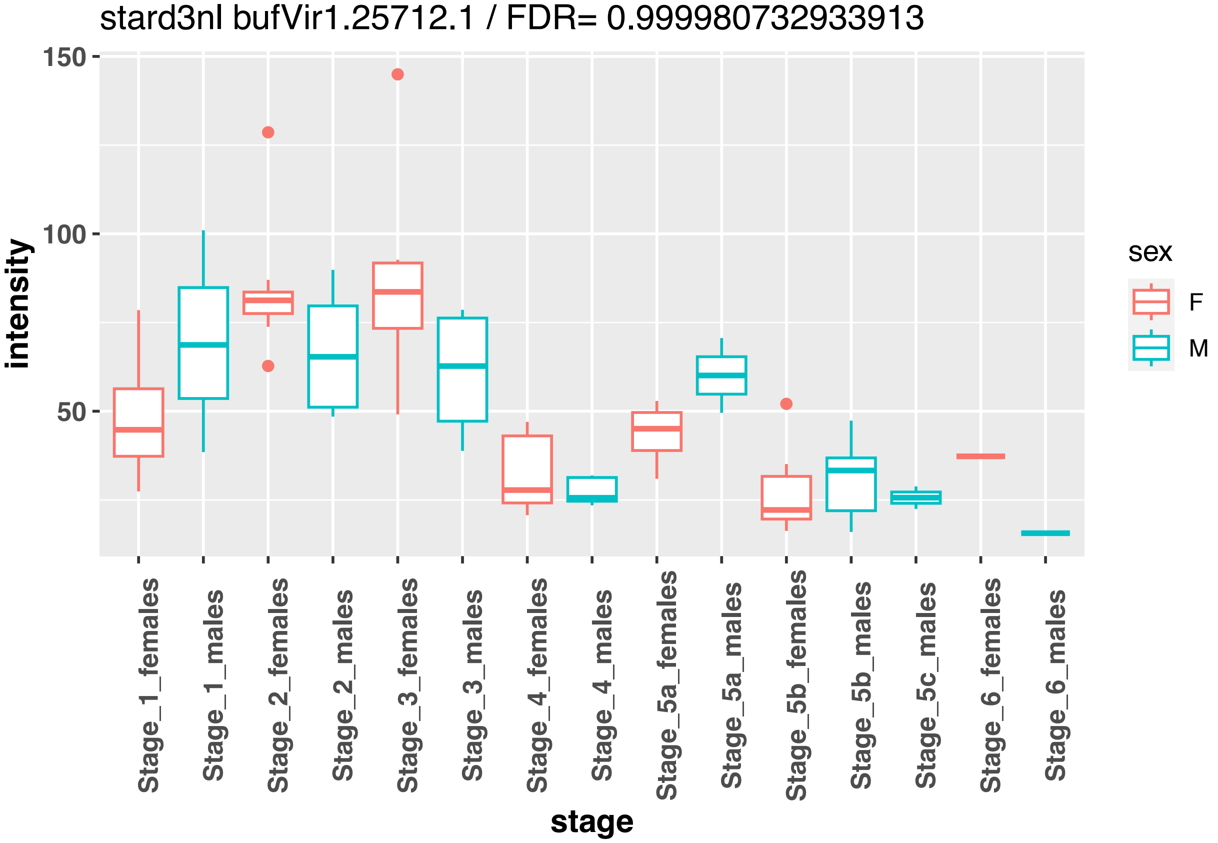
stard3nl bufVir1.25712.1 / FDR= 0.999980732933913
intensity
150
100
50
Stage_1_females
Stage_1_males
Stage_2_females
Stage_2_males
Stage_3_females
Stage_3_males
Stage_4_females
Stage_4_males
Stage_5a_females
Stage_5a_males
Stage_5b_females
Stage_5b_males
Stage_5c_males
Stage_6_females
Stage_6_males
stage
sex
F
M

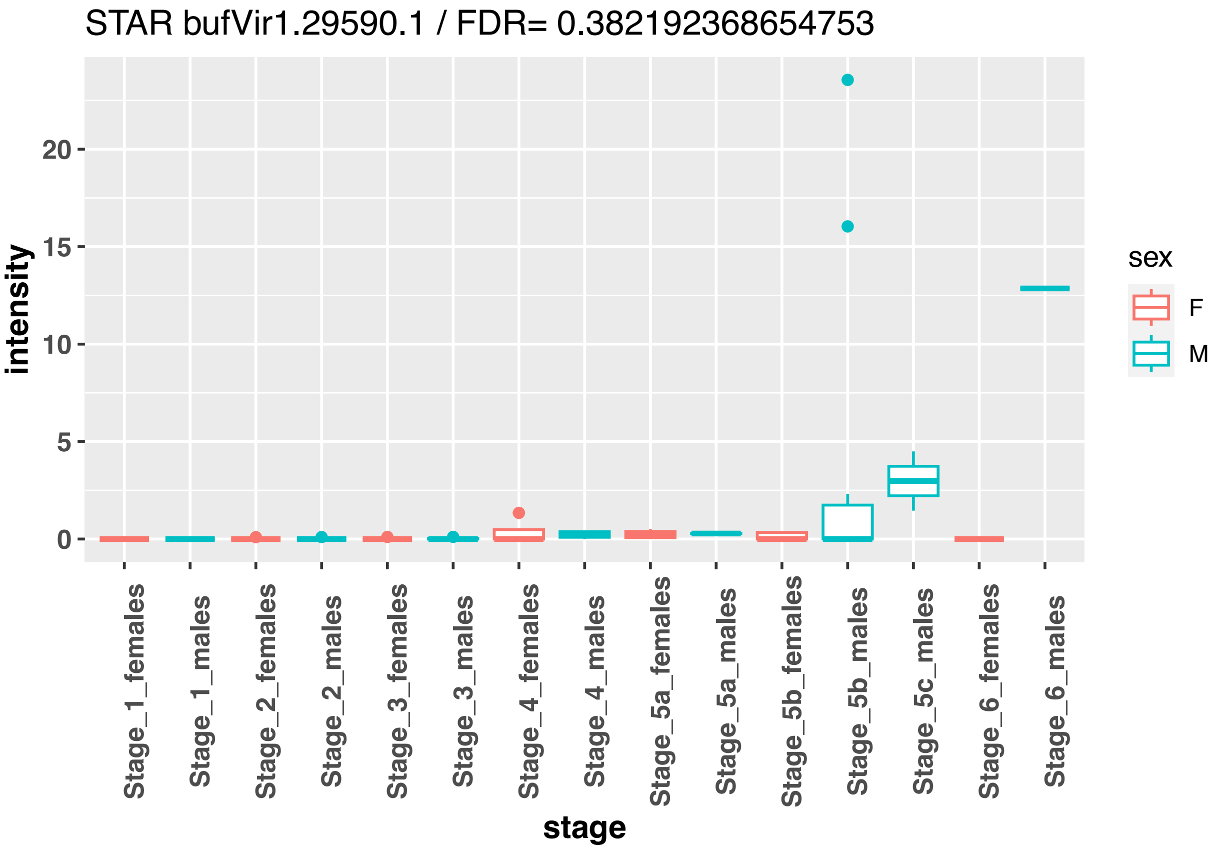
STAR bufVir1.29590.1 / FDR= 0.382192368654753
intensity
stage
sex
F
M
Stage_1_females
Stage_1_males
Stage_2_females
Stage_2_males
Stage_3_females
Stage_3_males
Stage_4_females
Stage_4_males
Stage_5a_females
Stage_5a_males
Stage_5b_females
Stage_5b_males
Stage_5c_males
Stage_6_females
Stage_6_males

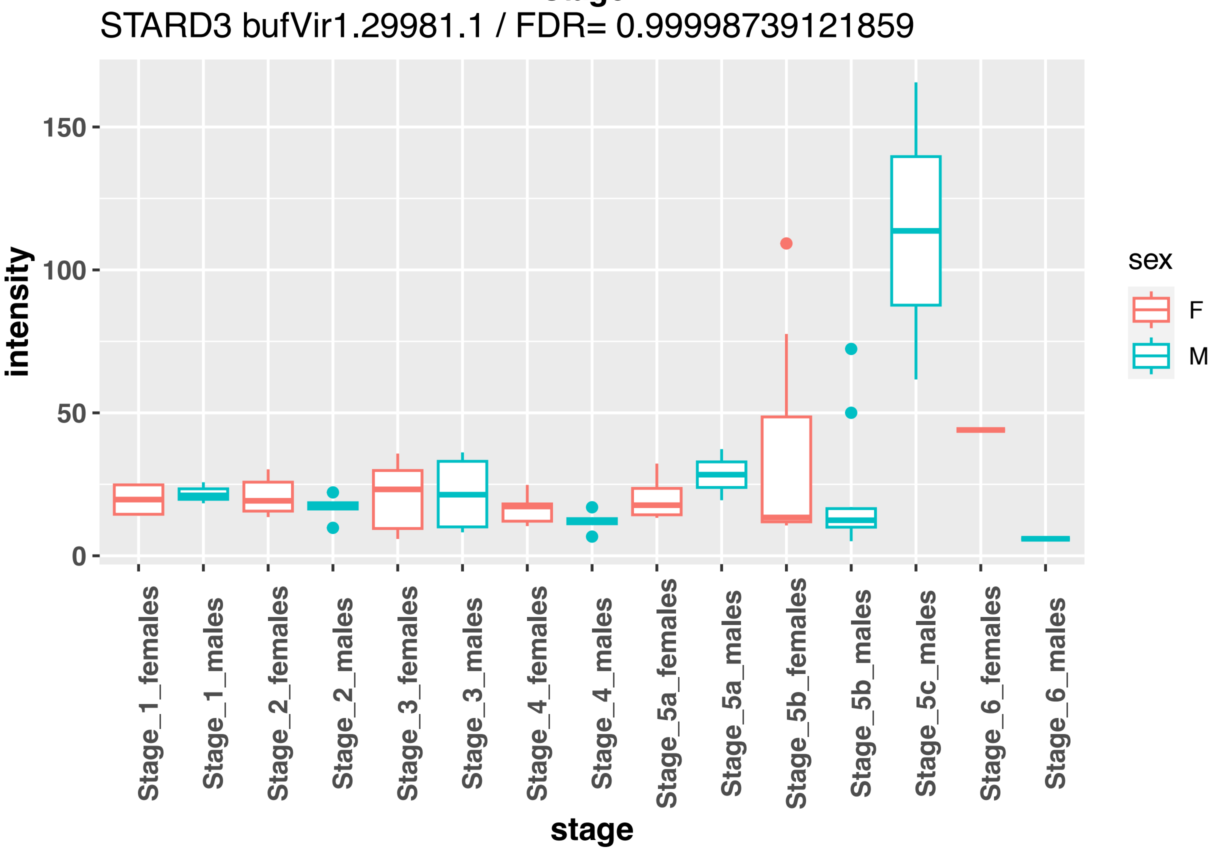
STARD3 bufVir1.29981.1 / FDR= 0.99998739121859
intensity
stage
sex
F
M
Stage_1_females
Stage_1_males
Stage_2_females
Stage_2_males
Stage_3_females
Stage_3_males
Stage_4_females
Stage_4_males
Stage_5a_females
Stage_5a_males
Stage_5b_females
Stage_5b_males
Stage_5c_males
Stage_6_females
Stage_6_males

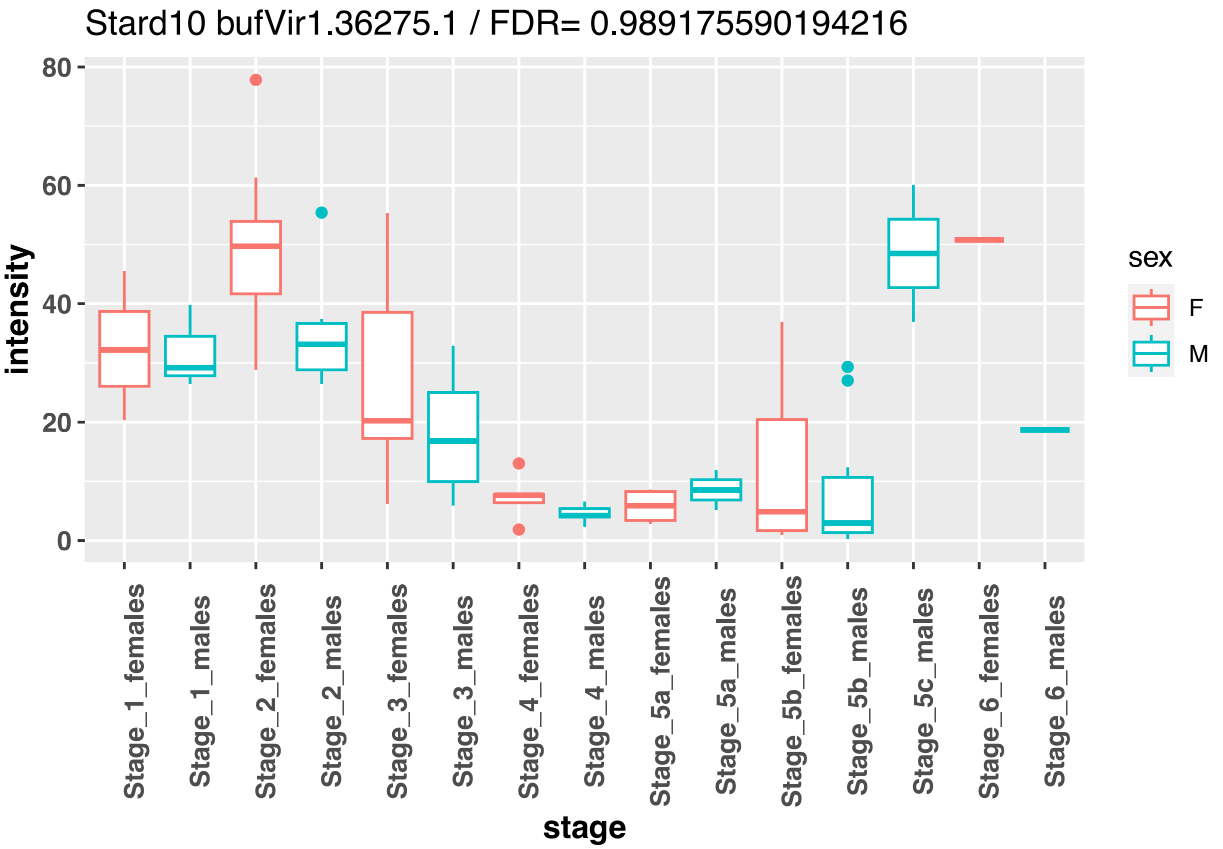
Stard10 bufVir1.36275.1 / FDR= 0.989175590194216
intensity
stage
sex
F
M
Stage_1_females
Stage_1_males
Stage_2_females
Stage_2_males
Stage_3_females
Stage_3_males
Stage_4_females
Stage_4_males
Stage_5a_females
Stage_5a_males
Stage_5b_females
Stage_5b_males
Stage_5c_males
Stage_6_females
Stage_6_males

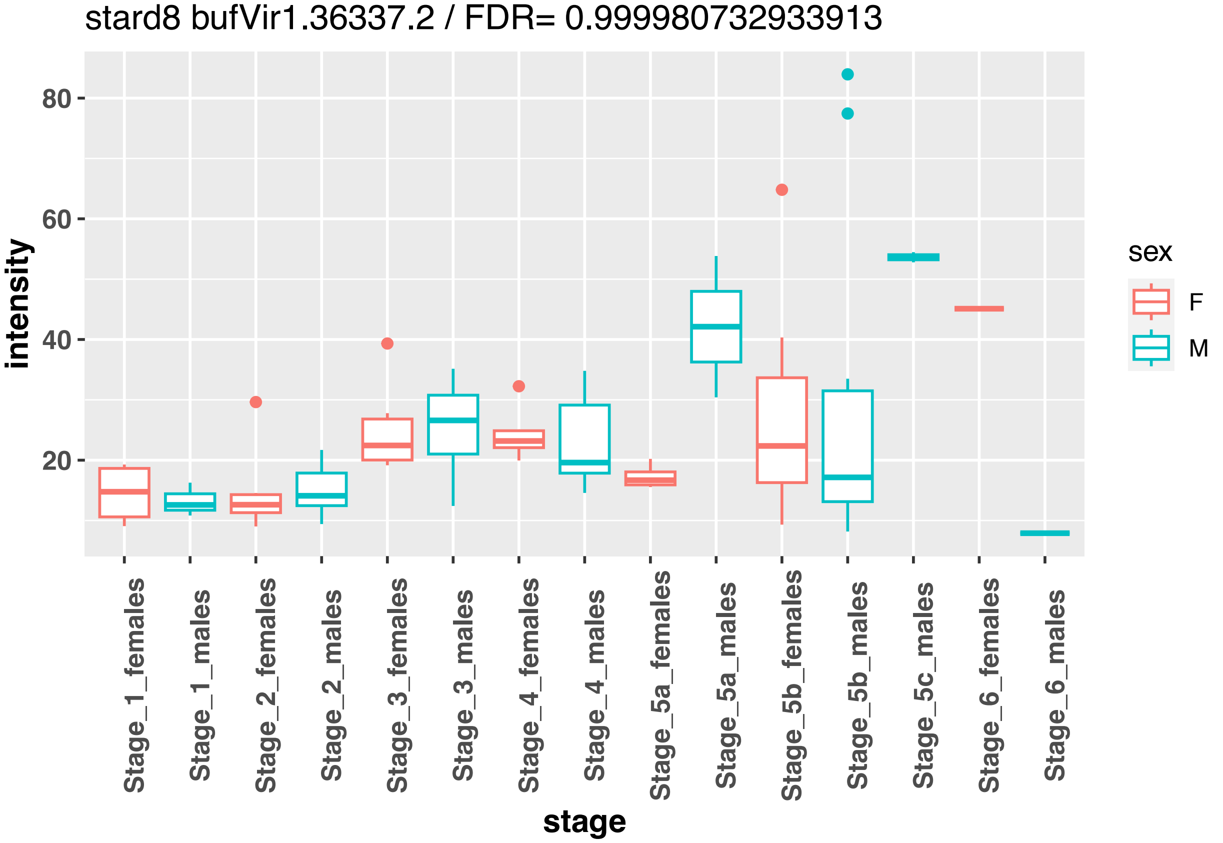
stard8 bufVir1.36337.2 / FDR= 0.999980732933913
intensity
stage
sex
F
M
Stage_1_females
Stage_1_males
Stage_2_females
Stage_2_males
Stage_3_females
Stage_3_males
Stage_4_females
Stage_4_males
Stage_5a_females
Stage_5a_males
Stage_5b_females
Stage_5b_males
Stage_5c_males
Stage_6_females
Stage_6_males

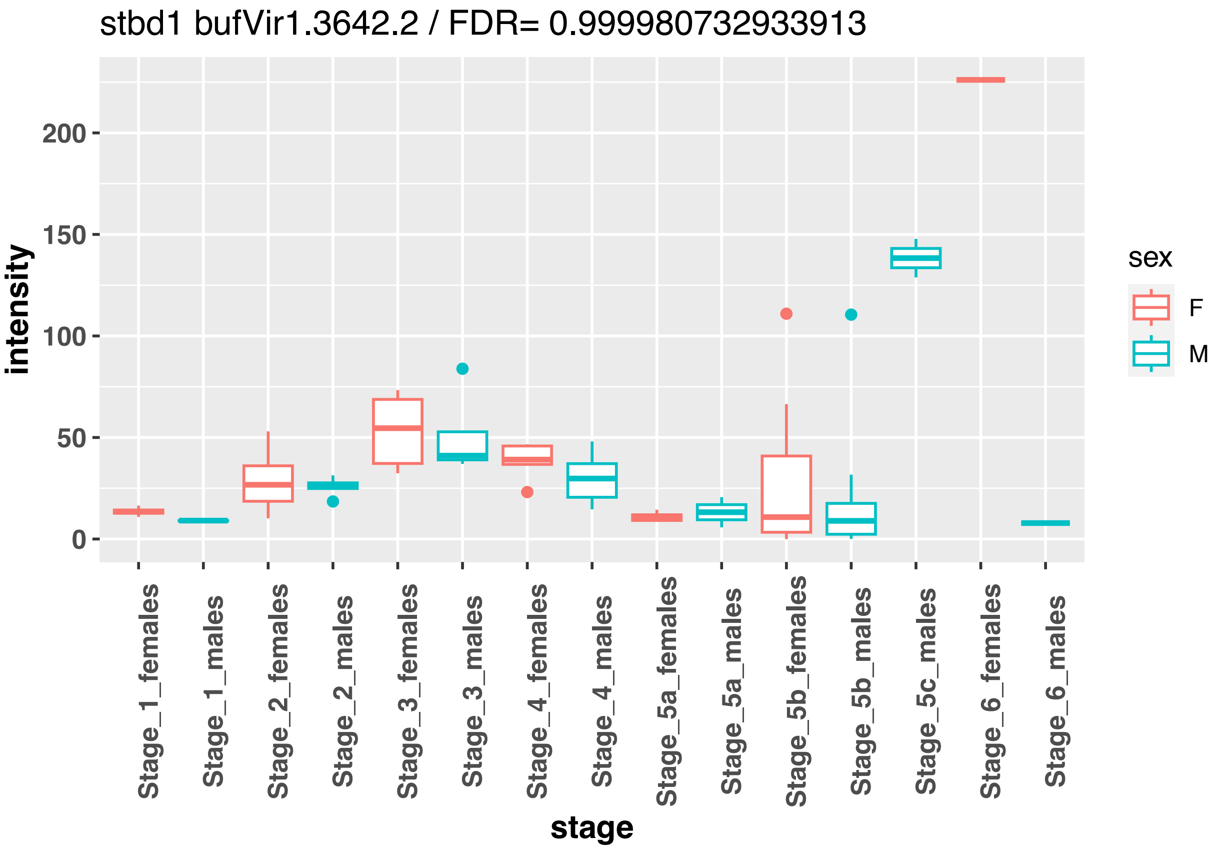
stbd1 bufVir1.3642.2 / FDR= 0.999980732933913
intensity
200
150
100
50
0
Stage_1_females
Stage_1_males
Stage_2_females
Stage_2_males
Stage_3_females
Stage_3_males
Stage_4_females
Stage_4_males
Stage_5a_females
Stage_5a_males
Stage_5b_females
Stage_5b_males
Stage_5c_males
Stage_6_females
Stage_6_males
stage
sex
F
M

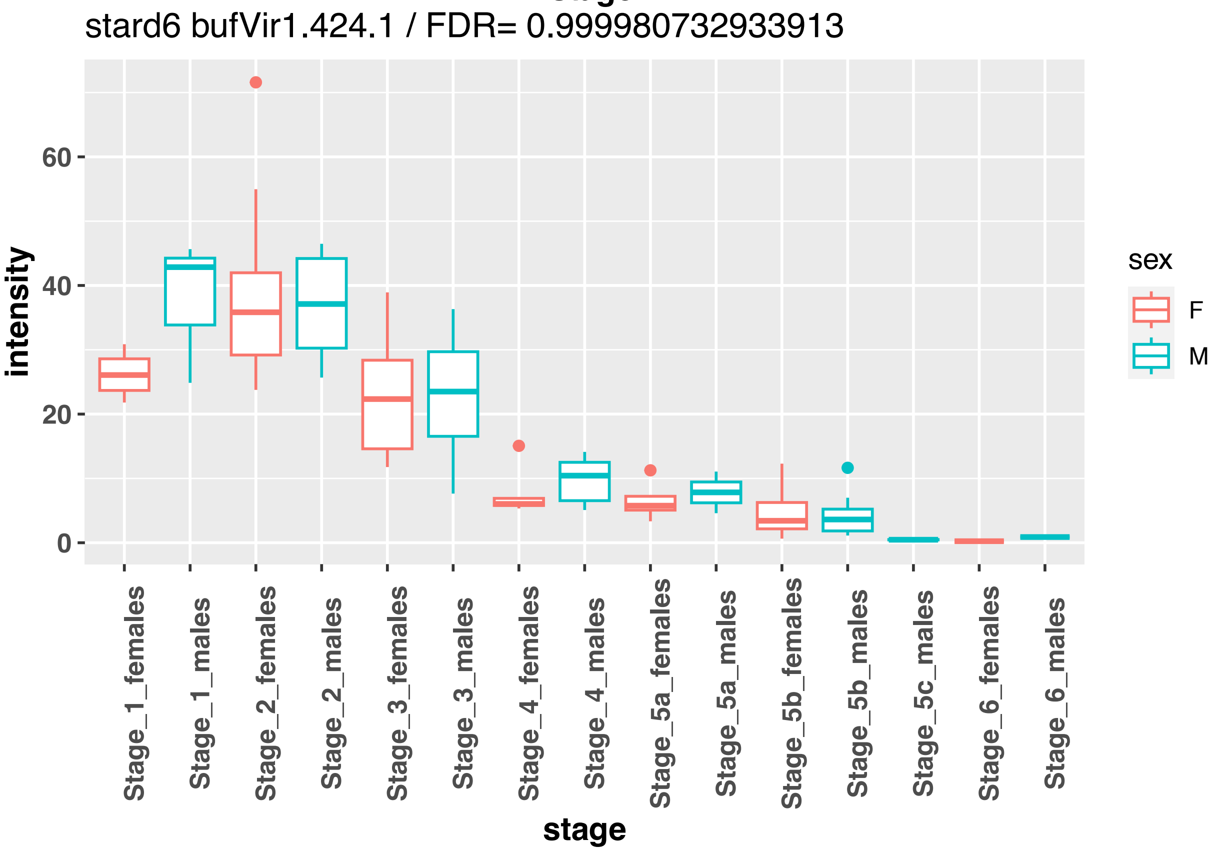
stard6 bufVir1.424.1 / FDR= 0.999980732933913
intensity
60
40
20
0
Stage_1_females
Stage_1_males
Stage_2_females
Stage_2_males
Stage_3_females
Stage_3_males
Stage_4_females
Stage_4_males
Stage_5a_females
Stage_5a_males
Stage_5b_females
Stage_5b_males
Stage_5c_males
Stage_6_females
Stage_6_males
stage
sex
F
M

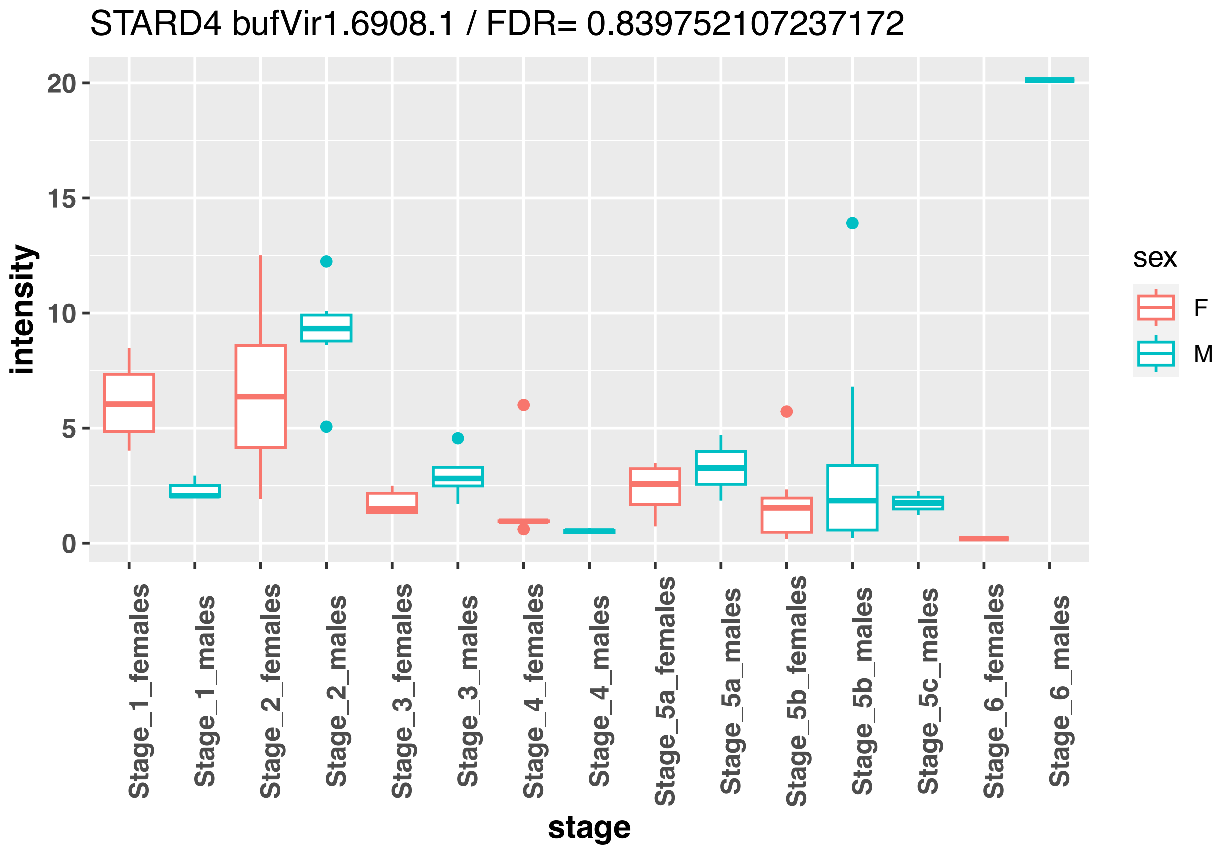
STARD4 bufVir1.6908.1 / FDR= 0.839752107237172
intensity
20
15
10
5
0
Stage_1_females
Stage_1_males
Stage_2_females
Stage_2_males
Stage_3_females
Stage_3_males
Stage_4_females
Stage_4_males
Stage_5a_females
Stage_5a_males
Stage_5b_females
Stage_5b_males
Stage_5c_males
Stage_6_females
Stage_6_males
stage
sex
F
M

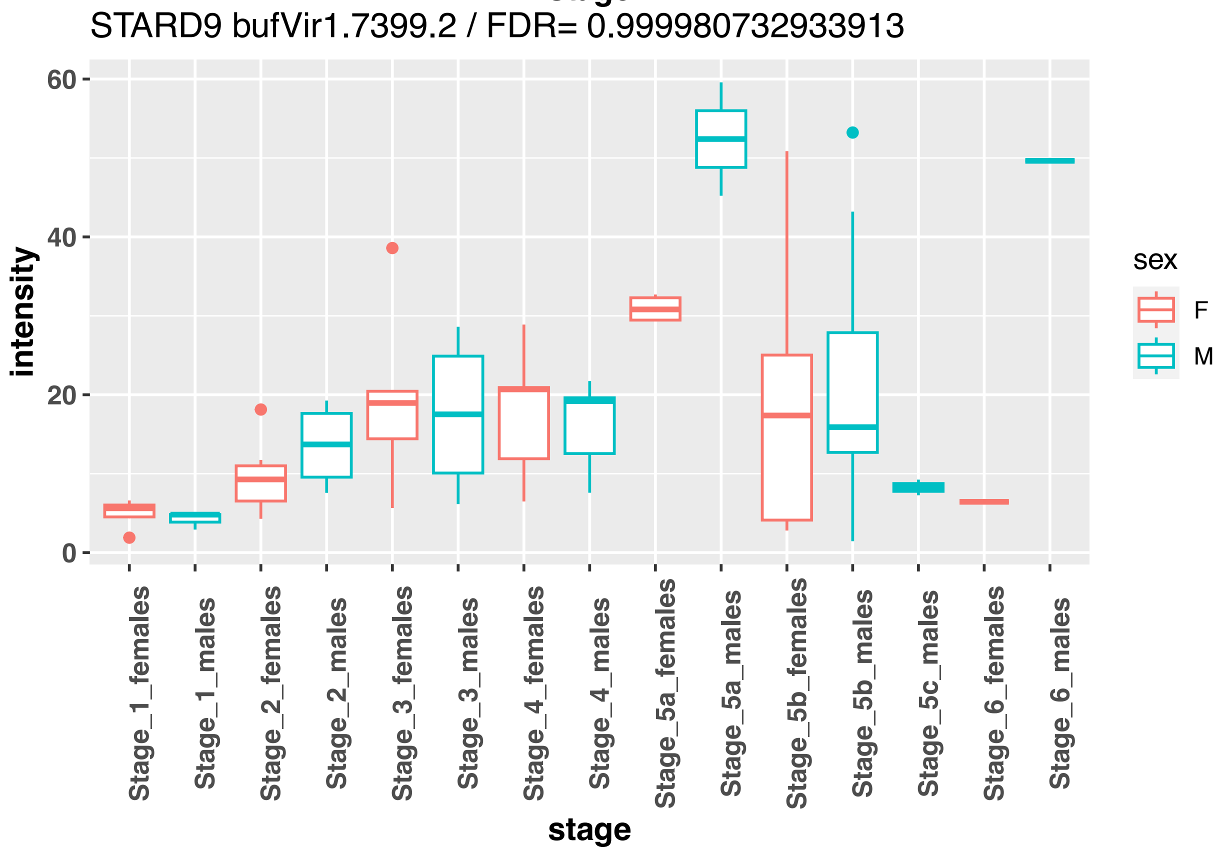
STARD9 bufVir1.7399.2 / FDR= 0.999980732933913
intensity
60
40
20
0
Stage_1_females
Stage_1_males
Stage_2_females
Stage_2_males
Stage_3_females
Stage_3_males
Stage_4_females
Stage_4_males
Stage_5a_females
Stage_5a_males
Stage_5b_females
Stage_5b_males
Stage_5c_males
Stage_6_females
Stage_6_males
stage
sex
F
M

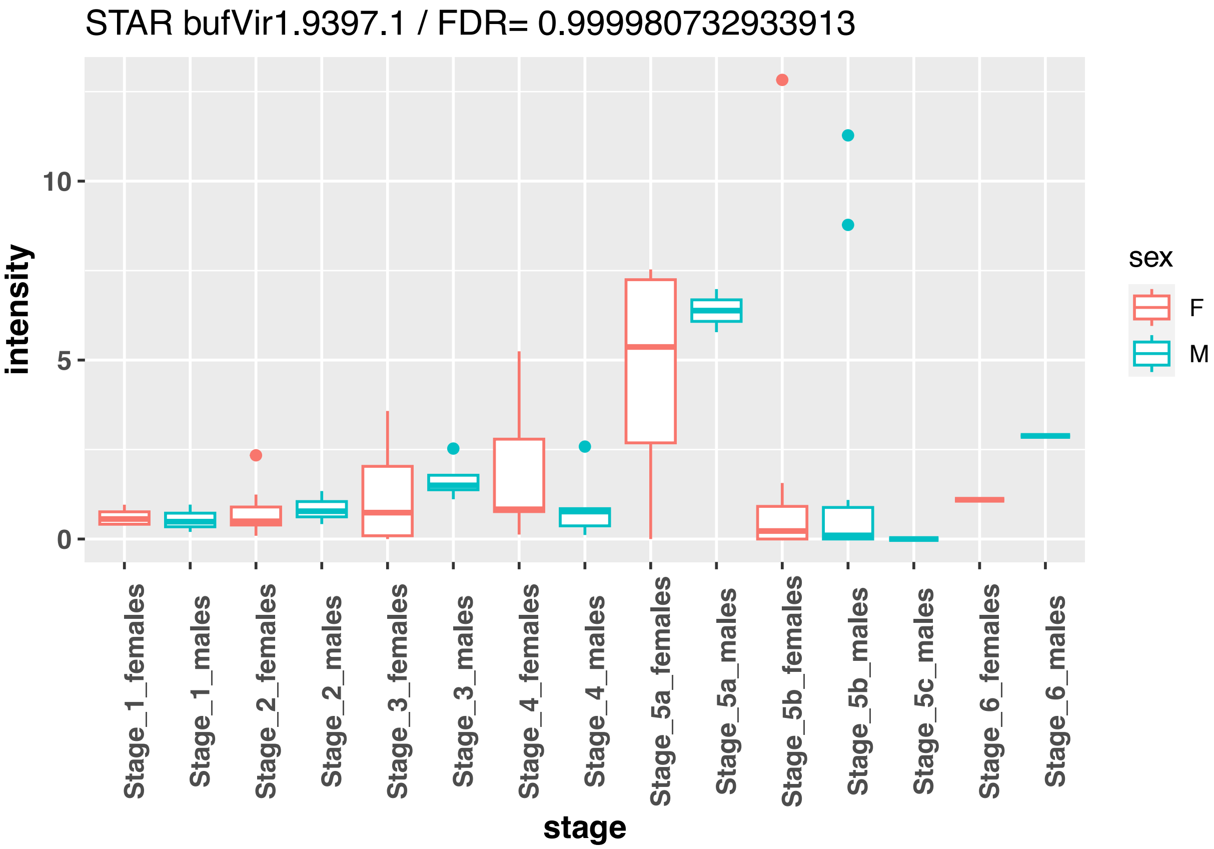
STAR bufVir1.9397.1 / FDR= 0.999980732933913
intensity
10
5
0
Stage_1_females
Stage_1_males
Stage_2_females
Stage_2_males
Stage_3_females
Stage_3_males
Stage_4_females
Stage_4_males
Stage_5a_females
Stage_5a_males
Stage_5b_females
Stage_5b_males
Stage_5c_males
Stage_6_females
Stage_6_males
stage
sex
F
M

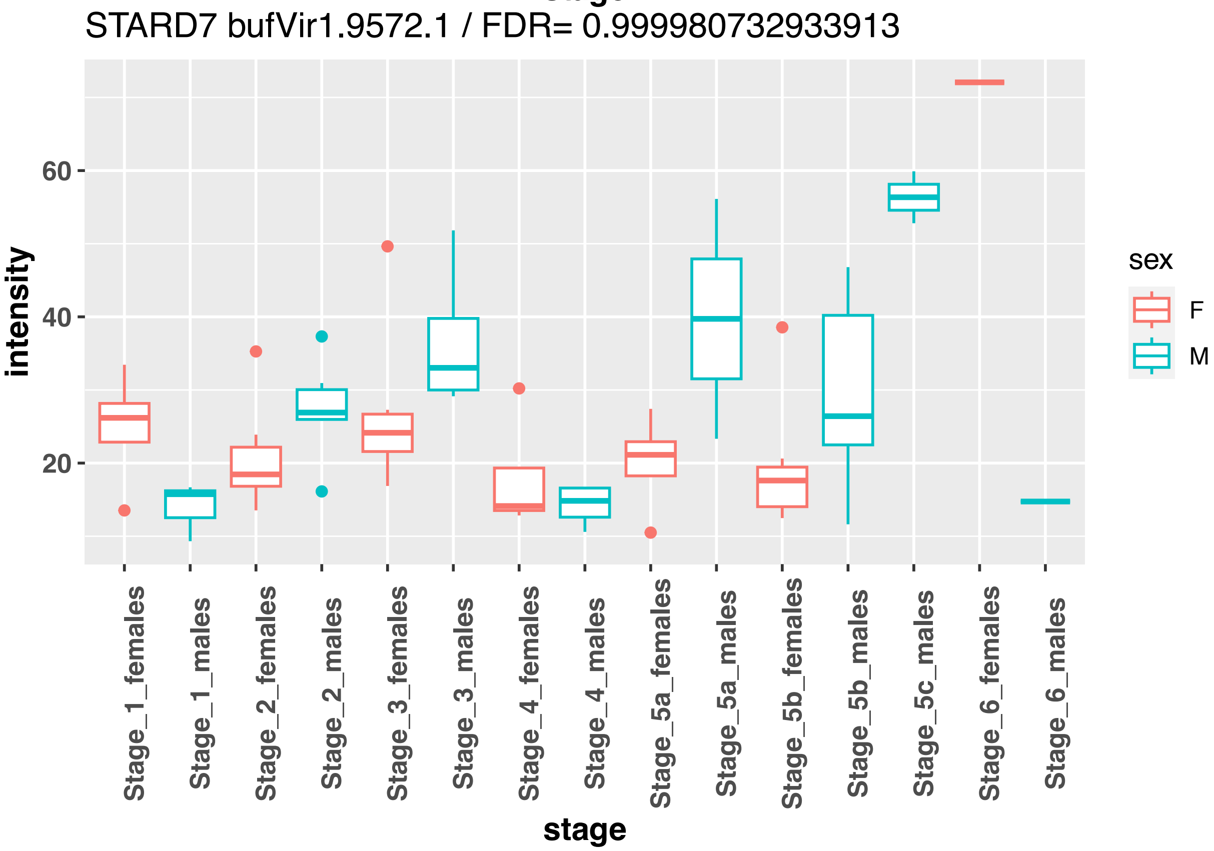
STARD7 bufVir1.9572.1 / FDR= 0.999980732933913
intensity
60
40
20
Stage_1_females
Stage_1_males
Stage_2_females
Stage_2_males
Stage_3_females
Stage_3_males
Stage_4_females
Stage_4_males
Stage_5a_females
Stage_5a_males
Stage_5b_females
Stage_5b_males
Stage_5c_males
Stage_6_females
Stage_6_males
stage
sex
F
M

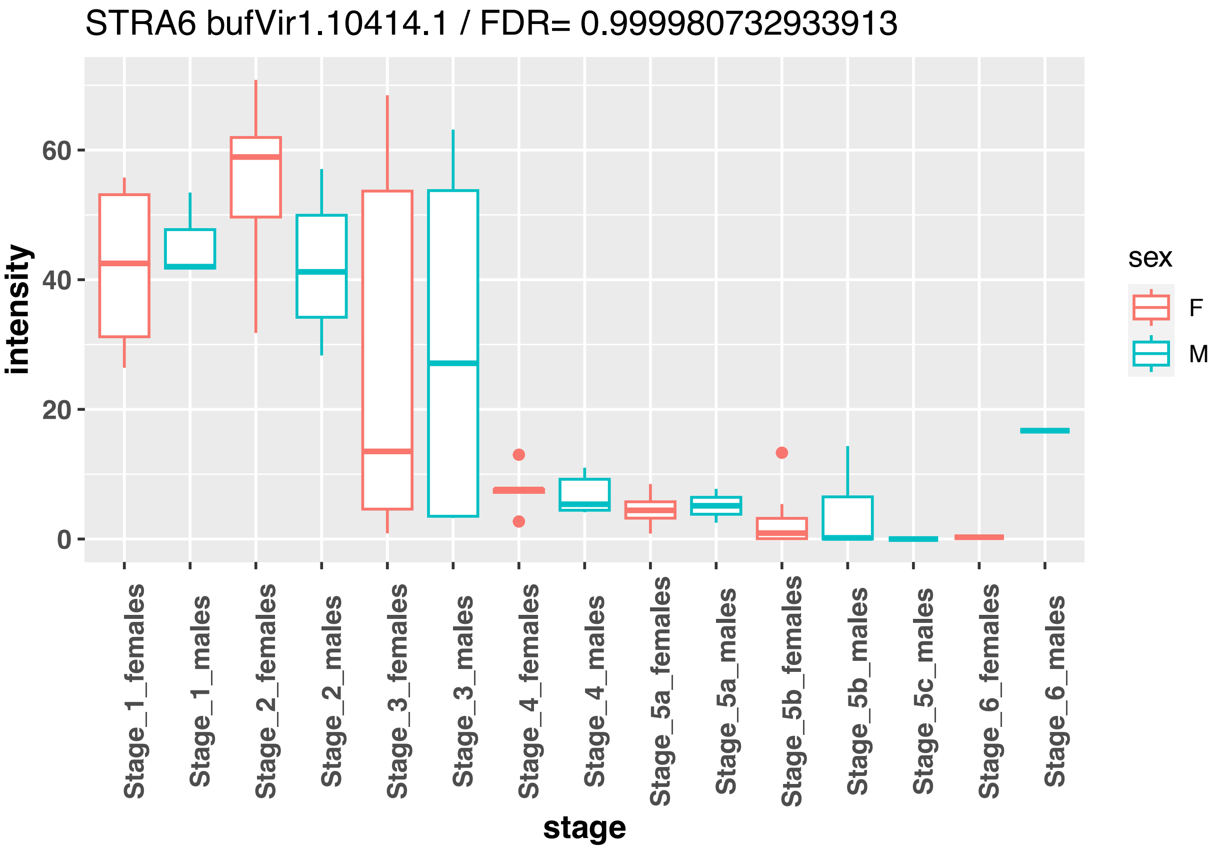
STRA6 bufVir1.10414.1 / FDR= 0.999980732933913
intensity
60
40
20
0
Stage_1_females
Stage_1_males
Stage_2_females
Stage_2_males
Stage_3_females
Stage_3_males
Stage_4_females
Stage_4_males
Stage_5a_females
Stage_5a_males
Stage_5b_females
Stage_5b_males
Stage_5c_males
Stage_6_females
Stage_6_males
stage
sex
F
M

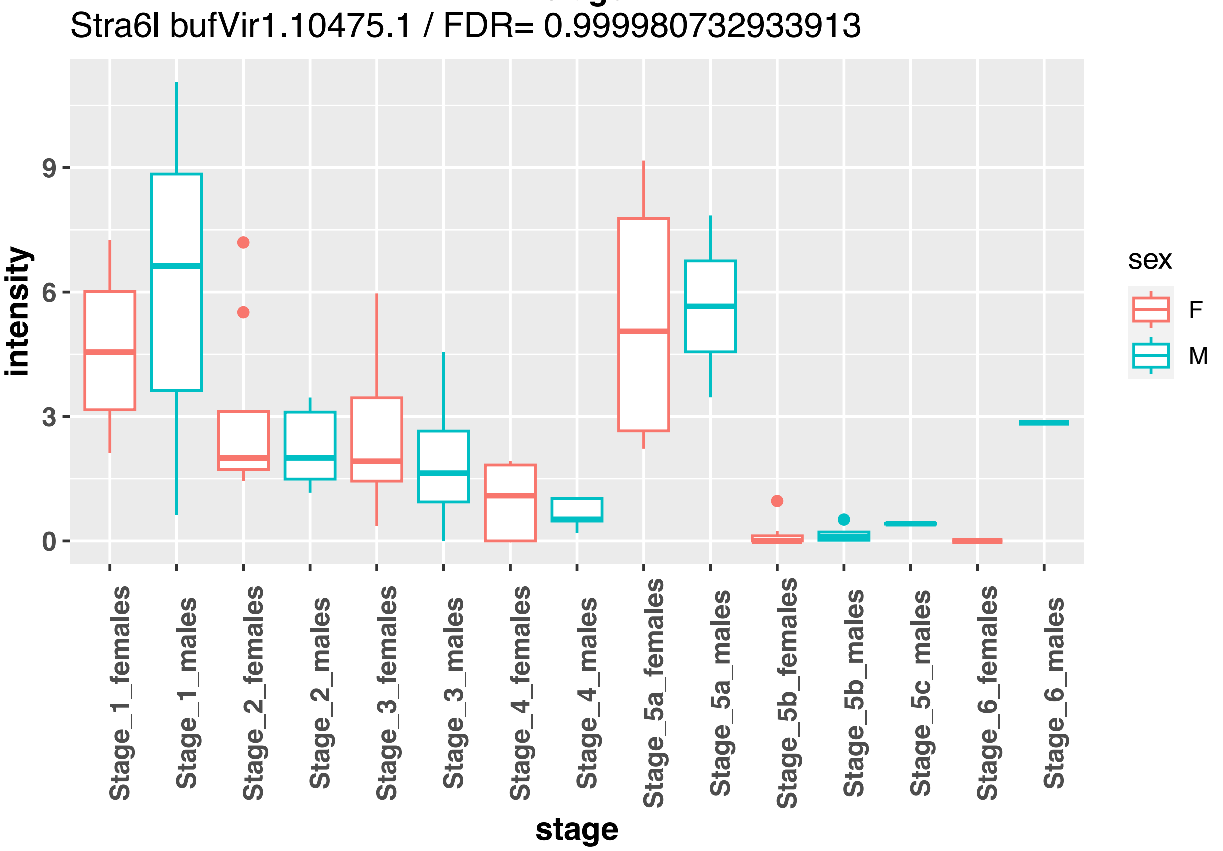
Stra6l bufVir1.10475.1 / FDR= 0.999980732933913
intensity
9
6
3
0
Stage_1_females
Stage_1_males
Stage_2_females
Stage_2_males
Stage_3_females
Stage_3_males
Stage_4_females
Stage_4_males
Stage_5a_females
Stage_5a_males
Stage_5b_females
Stage_5b_males
Stage_5c_males
Stage_6_females
Stage_6_males
stage
sex
F
M

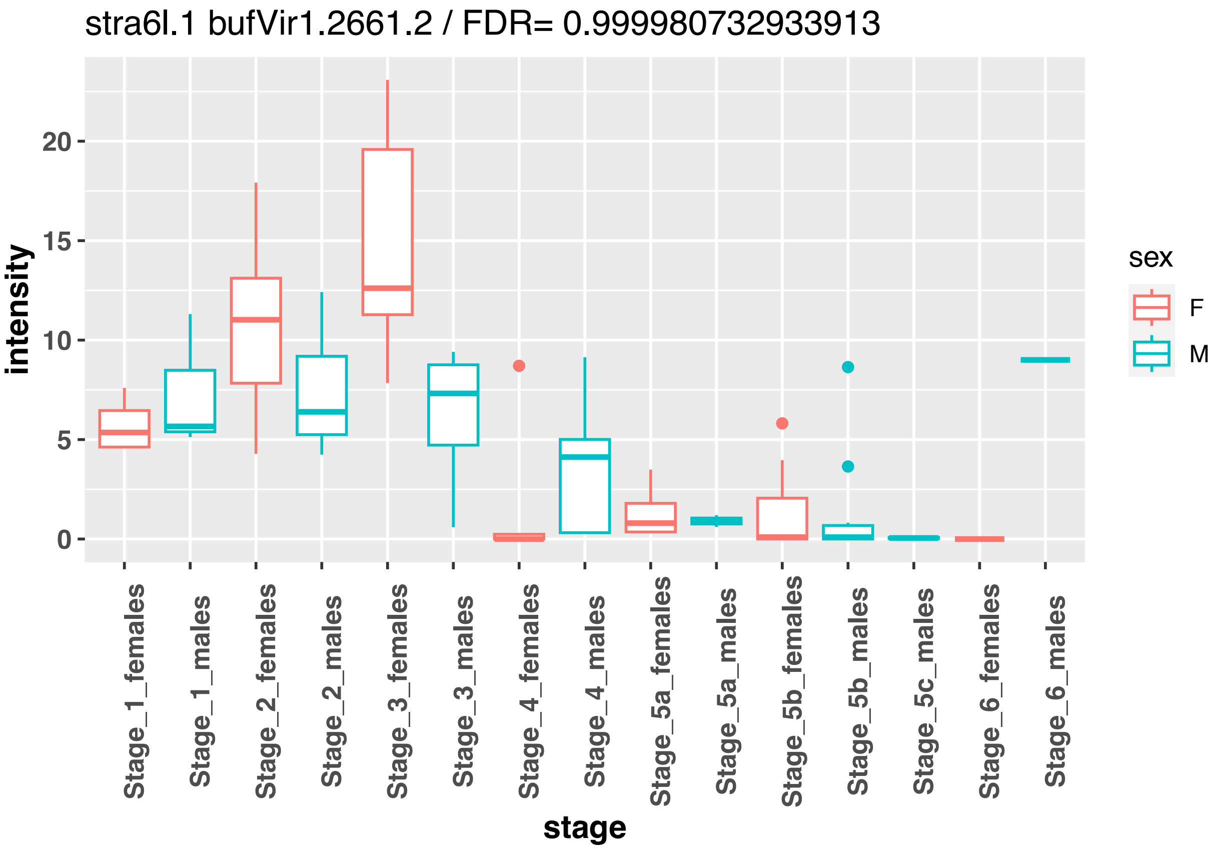
stra6l.1 bufVir1.2661.2 / FDR= 0.999980732933913
intensity
20
15
10
5
0
stage
Stage_1_females
Stage_1_males
Stage_2_females
Stage_2_males
Stage_3_females
Stage_3_males
Stage_4_females
Stage_4_males
Stage_5a_females
Stage_5a_males
Stage_5b_females
Stage_5b_males
Stage_5c_males
Stage_6_females
Stage_6_males
sex
F
M

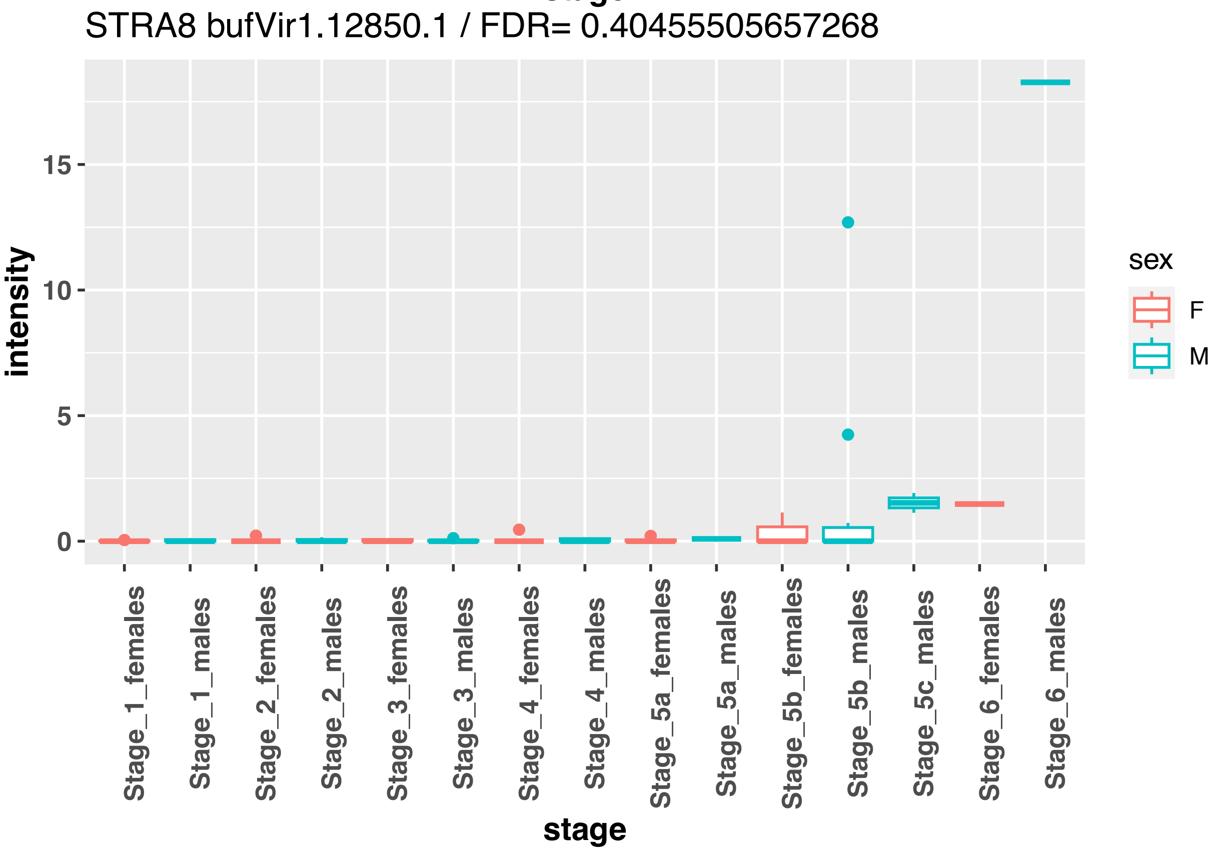
STRA8 bufVir1.12850.1 / FDR= 0.40455505657268
intensity
15
10
5
0
stage
Stage_1_females
Stage_1_males
Stage_2_females
Stage_2_males
Stage_3_females
Stage_3_males
Stage_4_females
Stage_4_males
Stage_5a_females
Stage_5a_males
Stage_5b_females
Stage_5b_males
Stage_5c_males
Stage_6_females
Stage_6_males
sex
F
M

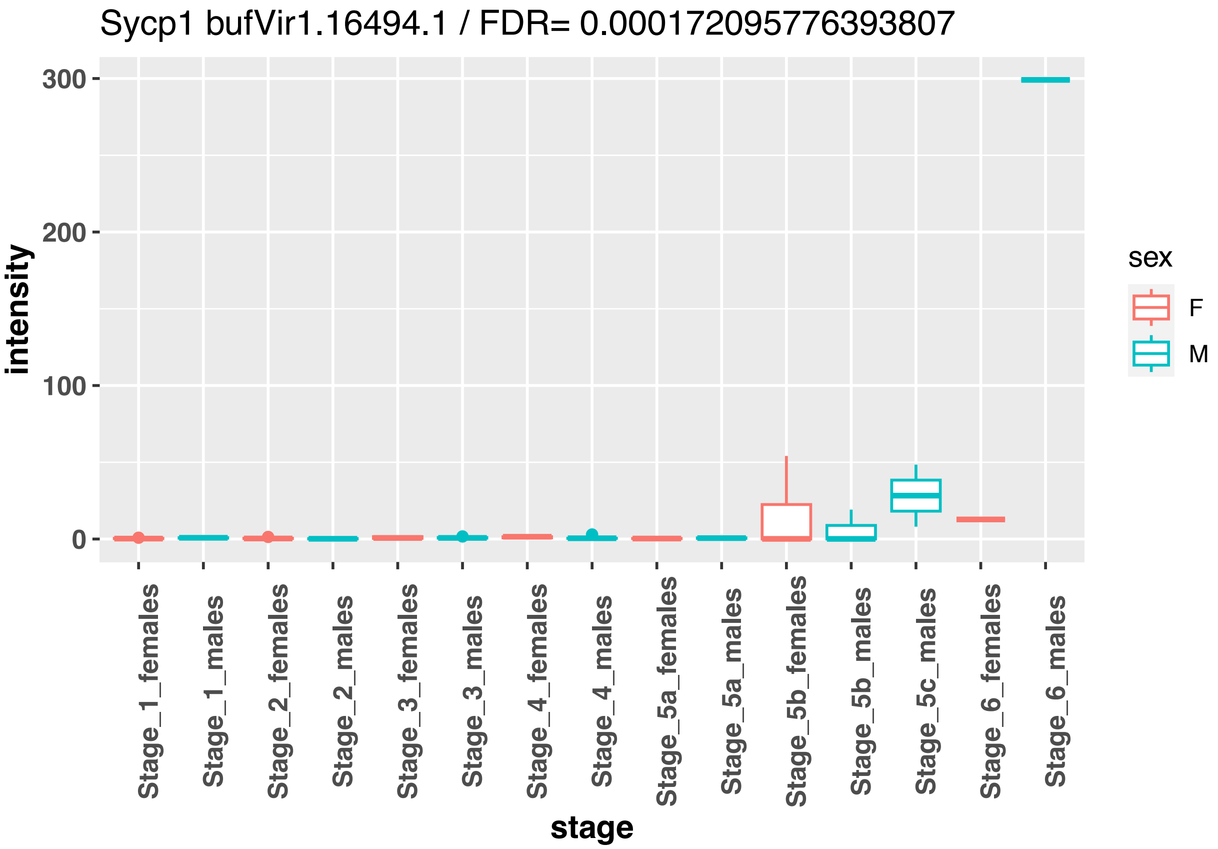
Sycp1 bufVir1.16494.1 / FDR= 0.000172095776393807
intensity
300
200
100
0
stage
Stage_1_females
Stage_1_males
Stage_2_females
Stage_2_males
Stage_3_females
Stage_3_males
Stage_4_females
Stage_4_males
Stage_5a_females
Stage_5a_males
Stage_5b_females
Stage_5b_males
Stage_5c_males
Stage_6_females
Stage_6_males
sex
F
M

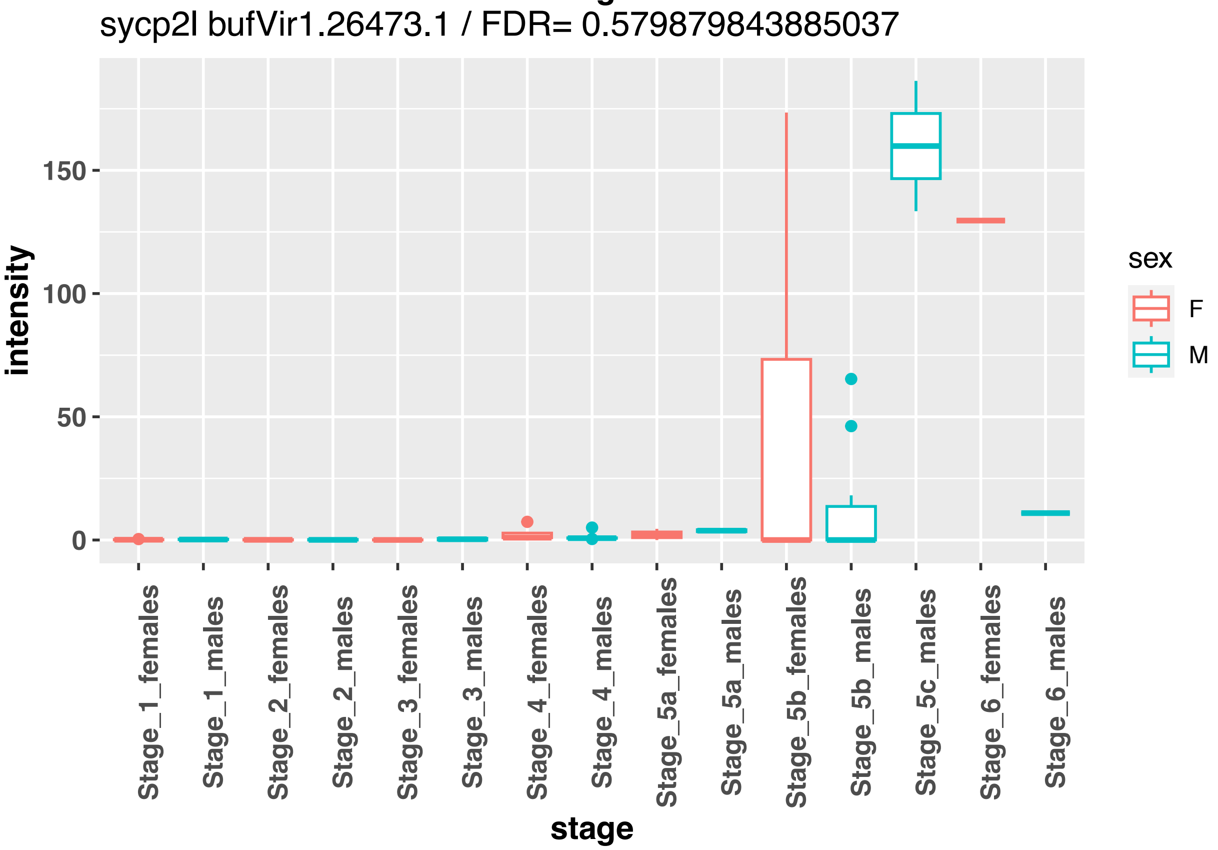
sycp2l bufVir1.26473.1 / FDR= 0.579879843885037
intensity
150
100
50
0
stage
Stage_1_females
Stage_1_males
Stage_2_females
Stage_2_males
Stage_3_females
Stage_3_males
Stage_4_females
Stage_4_males
Stage_5a_females
Stage_5a_males
Stage_5b_females
Stage_5b_males
Stage_5c_males
Stage_6_females
Stage_6_males
sex
F
M

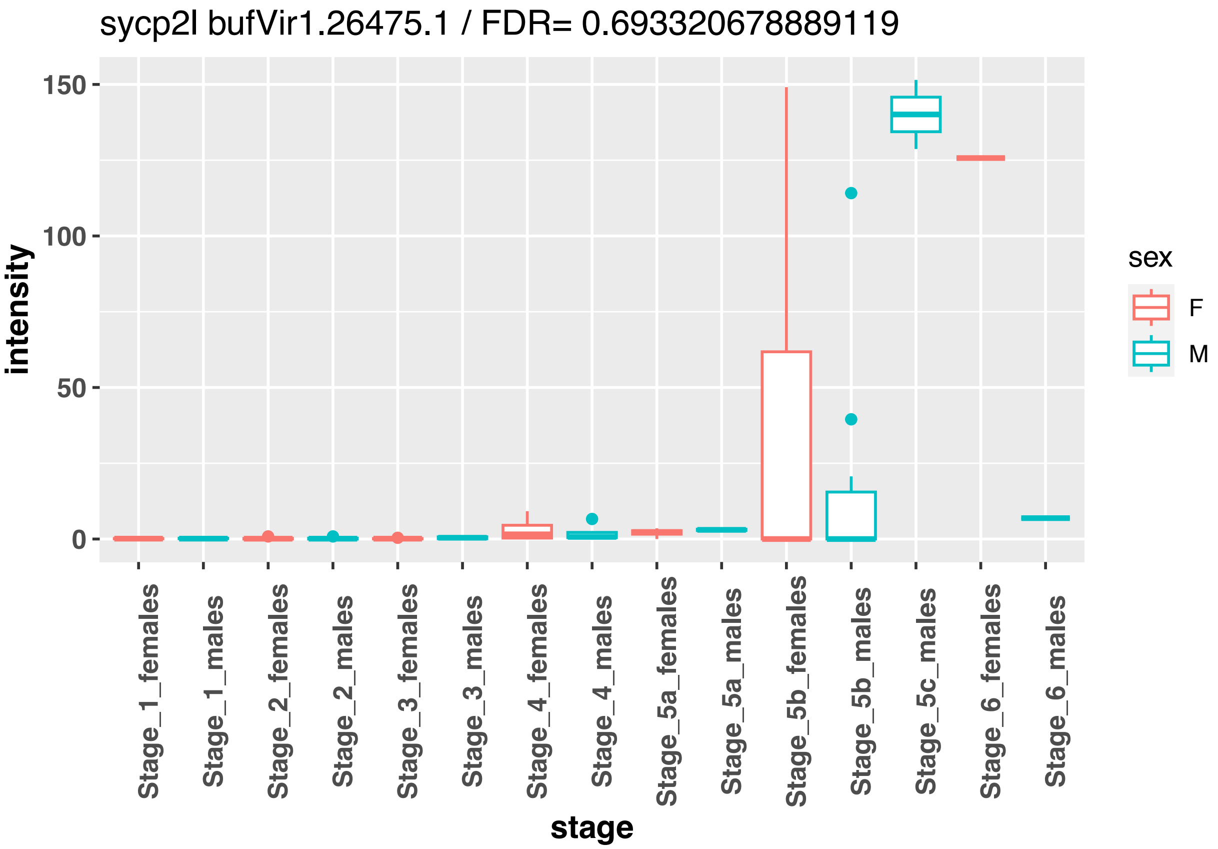
sycp2l bufVir1.26475.1 / FDR= 0.693320678889119
150
100
50
0
intensity
Stage_1_females
Stage_1_males
Stage_2_females
Stage_2_males
Stage_3_females
Stage_3_males
Stage_4_females
Stage_4_males
Stage_5a_females
Stage_5a_males
Stage_5b_females
Stage_5b_males
Stage_5c_males
Stage_6_females
Stage_6_males
stage
sex
F
M

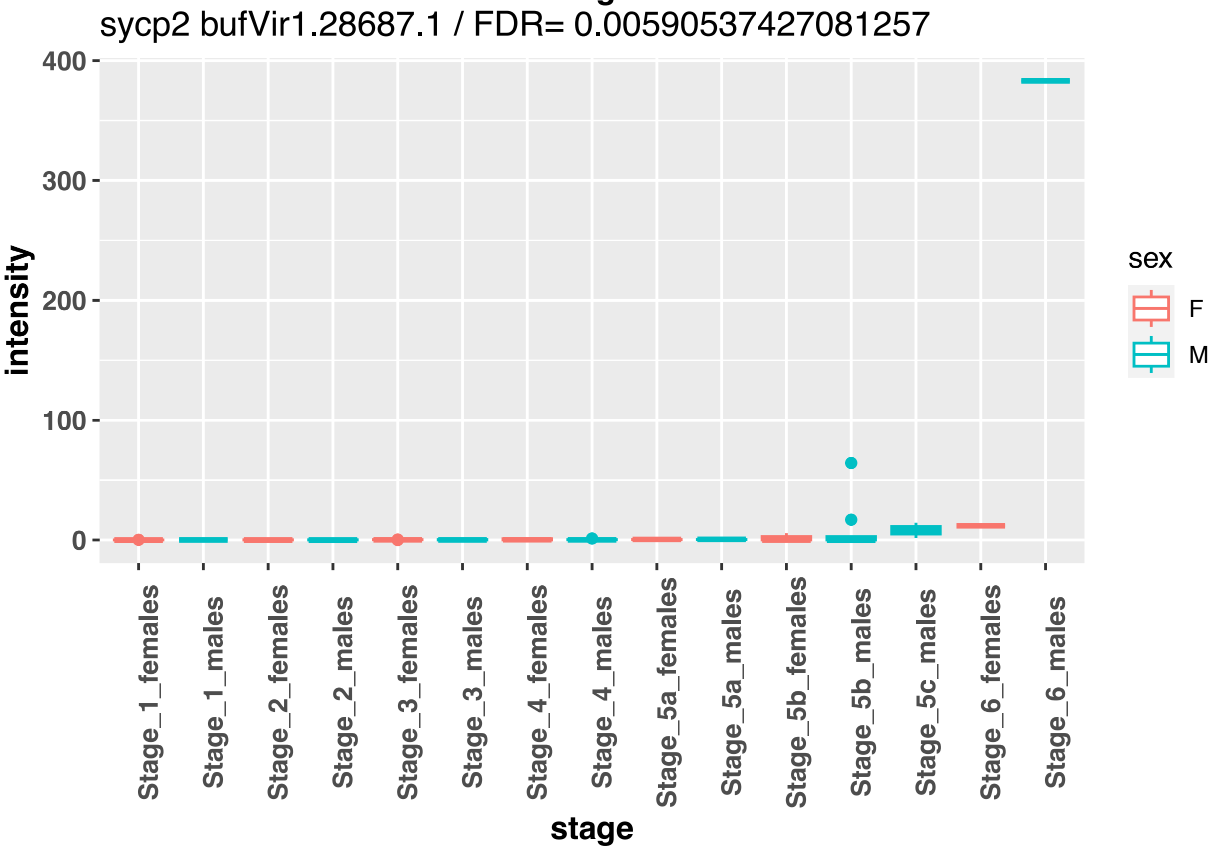
sycp2 bufVir1.28687.1 / FDR= 0.00590537427081257
400
300
200
100
0
intensity
Stage_1_females
Stage_1_males
Stage_2_females
Stage_2_males
Stage_3_females
Stage_3_males
Stage_4_females
Stage_4_males
Stage_5a_females
Stage_5a_males
Stage_5b_females
Stage_5b_males
Stage_5c_males
Stage_6_females
Stage_6_males
stage
sex
F
M

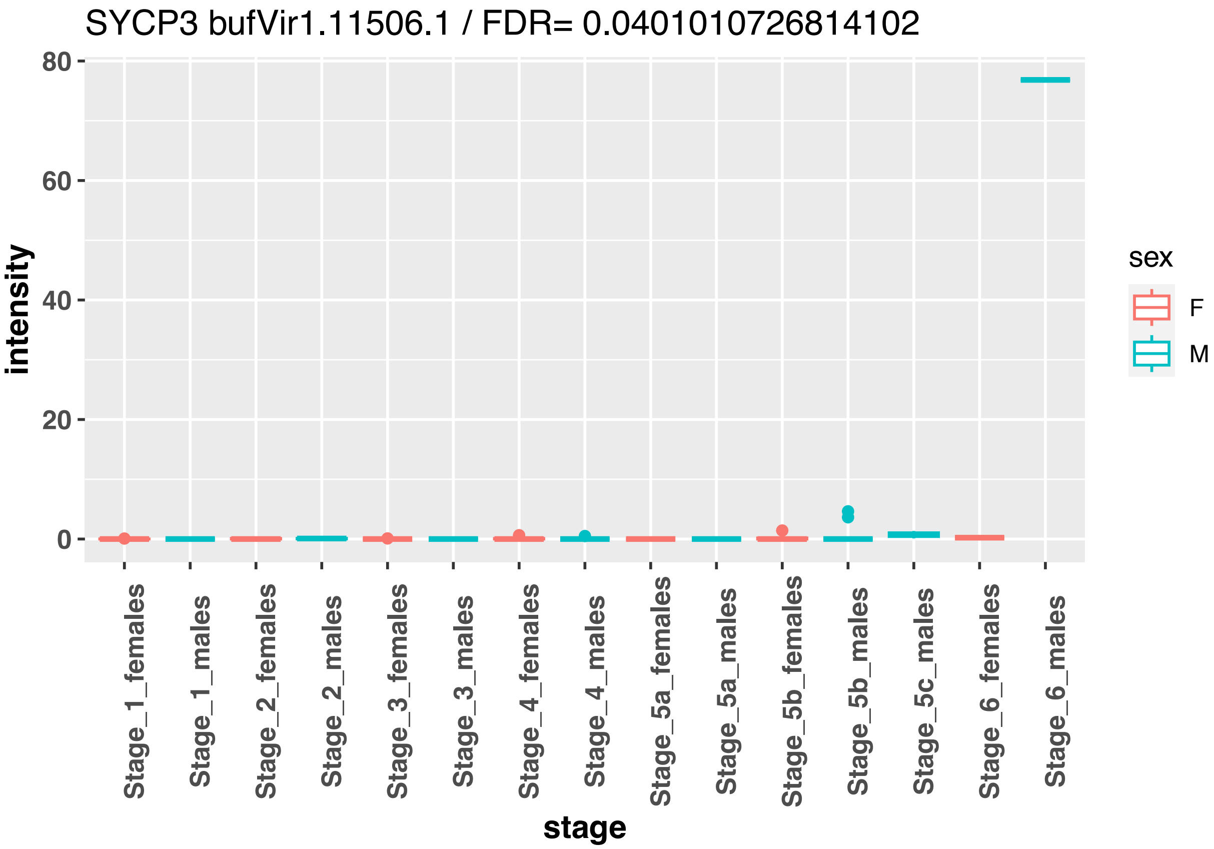


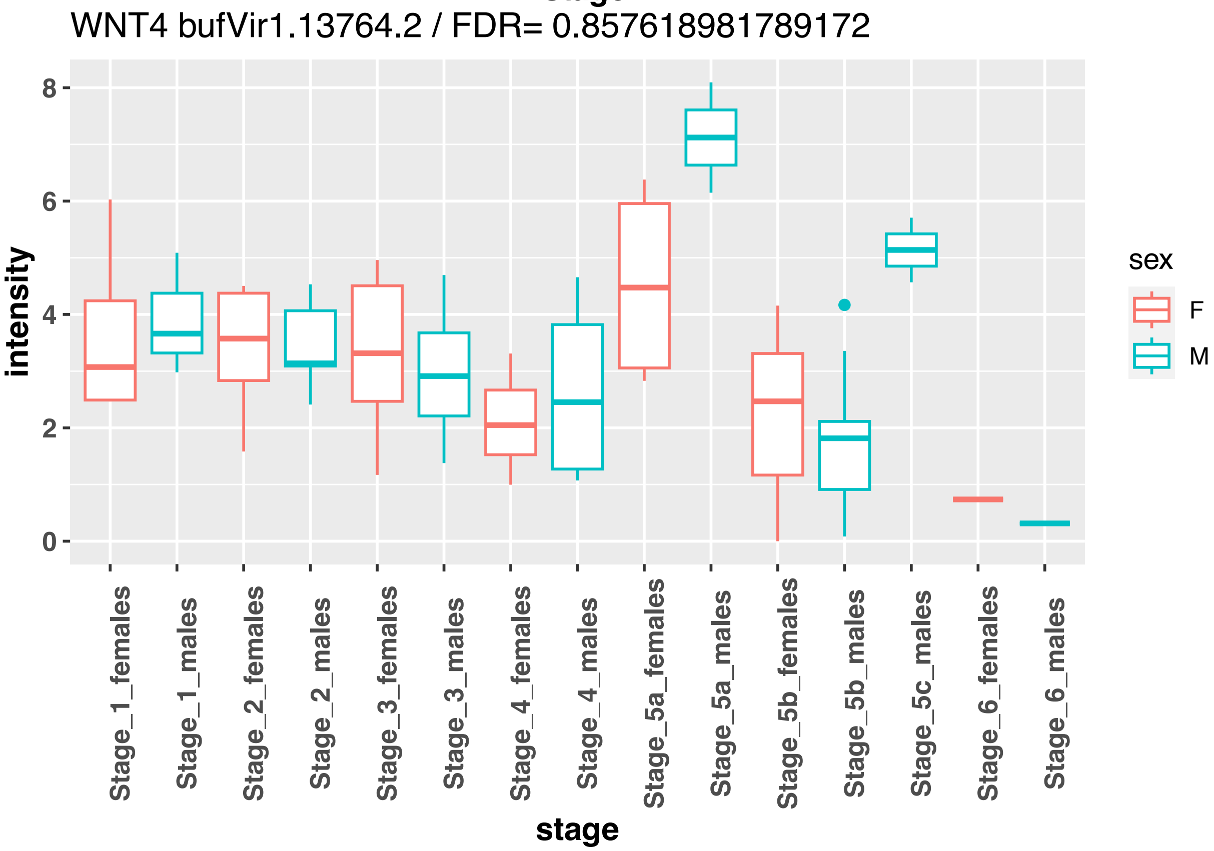


```
## R version 4.3.0 (2023-04-21)
```

```
## Platform: x86_64-pc-linux-gnu (64-bit)
## Running under: Ubuntu 22.04.3 LTS
##
## Matrix products: default
## BLAS:   /usr/lib/x86_64-linux-gnu/blas/libblas.so.3.10.0
## LAPACK: /usr/lib/x86_64-linux-gnu/lapack/liblapack.so.3.10.0
##
## locale:
##  [1] LC_CTYPE=fr_FR.UTF-8       LC_NUMERIC=C
##  [3] LC_TIME=fr_FR.UTF-8        LC_COLLATE=fr_FR.UTF-8
##  [5] LC_MONETARY=fr_FR.UTF-8    LC_MESSAGES=fr_FR.UTF-8
##  [7] LC_PAPER=fr_FR.UTF-8       LC_NAME=C
##  [9] LC_ADDRESS=C               LC_TELEPHONE=C
## [11] LC_MEASUREMENT=fr_FR.UTF-8 LC_IDENTIFICATION=C
##
## time zone: Europe/Paris
## tzcode source: system (glibc)
##
## attached base packages:
## [1] stats     graphics  grDevices utils     datasets  methods   base
##
## other attached packages:
## [1] knitr_1.43      plyr_1.8.8      edgeR_3.42.4    limma_3.56.2
## [5] pheatmap_1.0.12 ggplot2_3.4.2   reshape2_1.4.4
##
## loaded via a namespace (and not attached):
##  [1] gtable_0.3.3       dplyr_1.1.2        compiler_4.3.0     tidyselect_1.2.0
##  [5] Rcpp_1.0.10        stringr_1.5.0      splines_4.3.0      scales_1.2.1
##  [9] yaml_2.3.7         fastmap_1.1.1      lattice_0.21-8     R6_2.5.1
## [13] labeling_0.4.2     generics_0.1.3     tibble_3.2.1       munsell_0.5.0
## [17] pillar_1.9.0       RColorBrewer_1.1-3 rlang_1.1.1        utf8_1.2.3
## [21] stringi_1.7.12     xfun_0.39          cli_3.6.1          withr_2.5.0
## [25] magrittr_2.0.3     digest_0.6.31      grid_4.3.0         locfit_1.5-9.8
## [29] rstudioapi_0.14    lifecycle_1.0.3    vctrs_0.6.3        evaluate_0.21
## [33] glue_1.6.2         farver_2.1.1       fansi_1.0.4        colorspace_2.1-0
## [37] rmarkdown_2.22     tools_4.3.0        pkgconfig_2.0.3    htmltools_0.5.5
```
