## Supplementary Figures 20212102 for "A candidate sex determination locus in amphibians, evolved by structural variation between X- and Y-chromosomes"

**Supplementary Figure 11: *Vasa*, *bod1l* expression and H3K4me3-methylation in *B. viridis* gonads, overview images and additional representative examples.**

**a-m, v***asa* and *bod1L* expression shown by RNA *in-situ* hybridization (ISH) on 20 µm transverse cryosections of female (**a, b, g, h**) and male (**c, d, f, i-m**) gonads, Gosner 36-37. DAPI was used to stain nuclei. Sense controls for RNA ISH are shown in **a, c, e**, **g**, right panels. Red boxes in **a, c, g, i** (left panel) indicate zoomed images shown in **b, d, h, k,** respectively. Yellow boxes in **b, d, h, k** (right panel) indicate zoomed images shown in **Fig. 4a**-b**,** respectively. Red boxes in l**, m** (left panel) indicate zoomed images shown in right panels of l**, m**. *Bod1L* staining on three different cryosections of male gonads are shown in k**, l, m**. Magenta asterisks indicate nuclei of germ cells; magenta arrows: cytoplasm of germ cells; yellow arrows: somatic cells not stained by RNA ISH; green arrows: somatic cells stained by RNA ISH. **n, o,** H3K4me3 and DAPI staining on 20 µm transverse cryosections of female (**l**) and male (**m**) *B. viridis* Bidder's organ and gonads, Gosner stage 43-44. Yellow boxes in **l, m** indicate zoomed images shown in **Fig. 4g**, **h,** respectively. White asterisks indicate H3K4me3-positive germ cell nuclei; white arrowheads, H3K4me3-positive somatic cell nuclei; yellow arrowheads, H3K4me3-negative somatic cell nuclei. Scale bars: (a, c, g, i) 50 µm, (b, d-f, h, k-o) 20 µm.

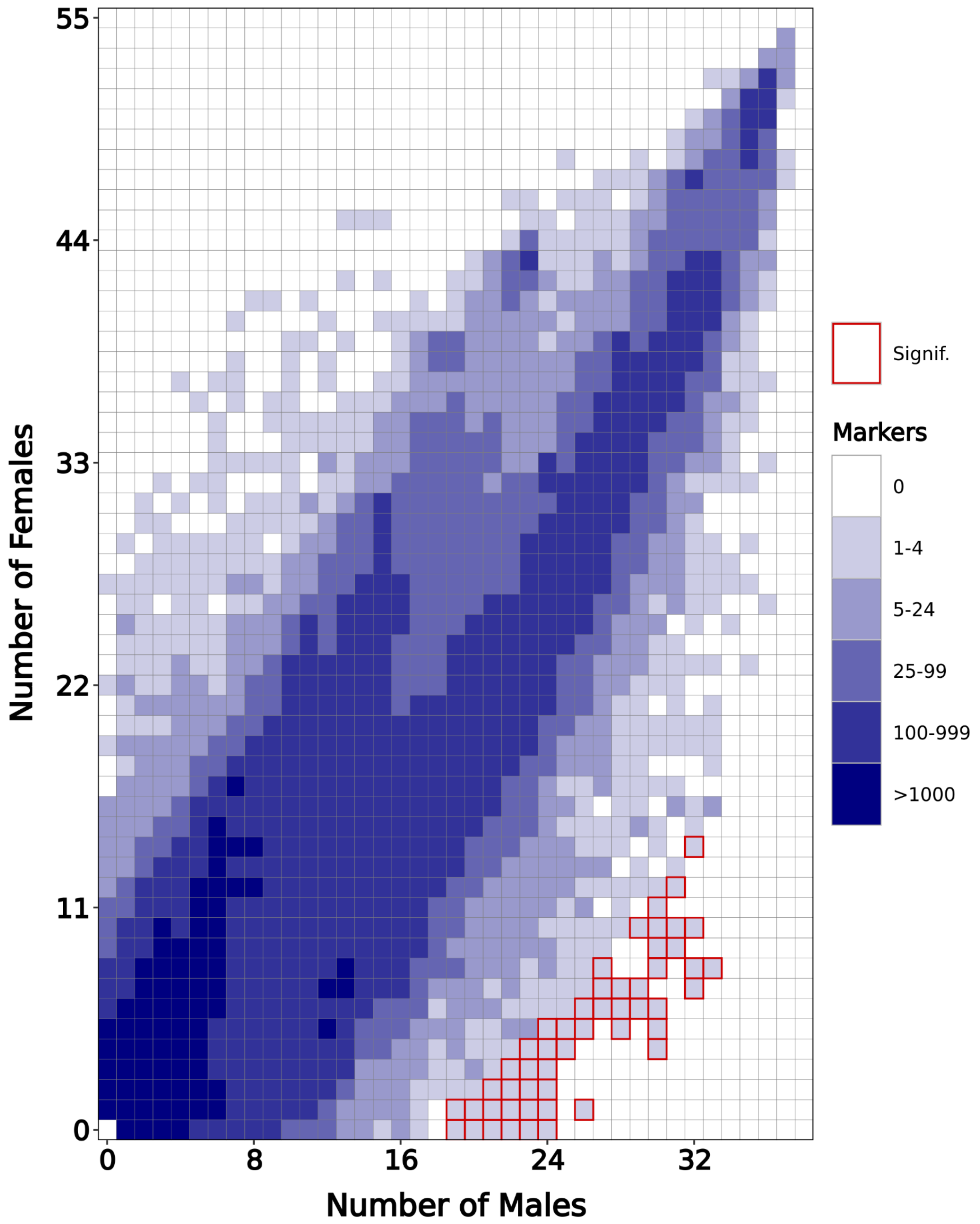


**Supplementary Figure 1**

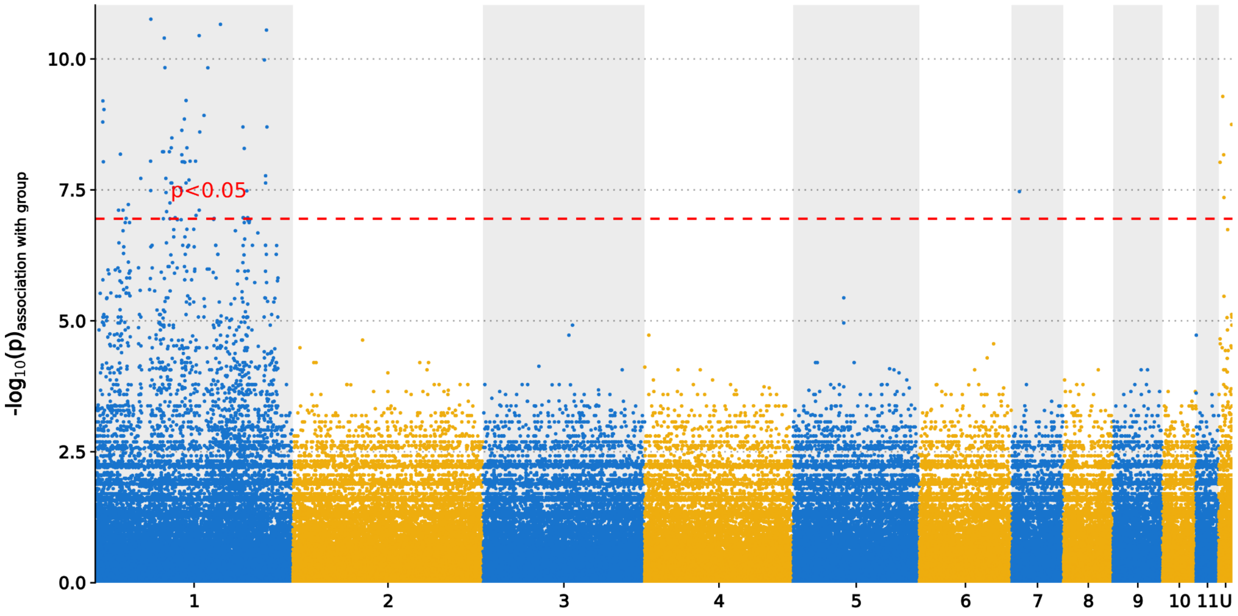


**Supplementry Figure 2**

Supplementary Figure 3

a

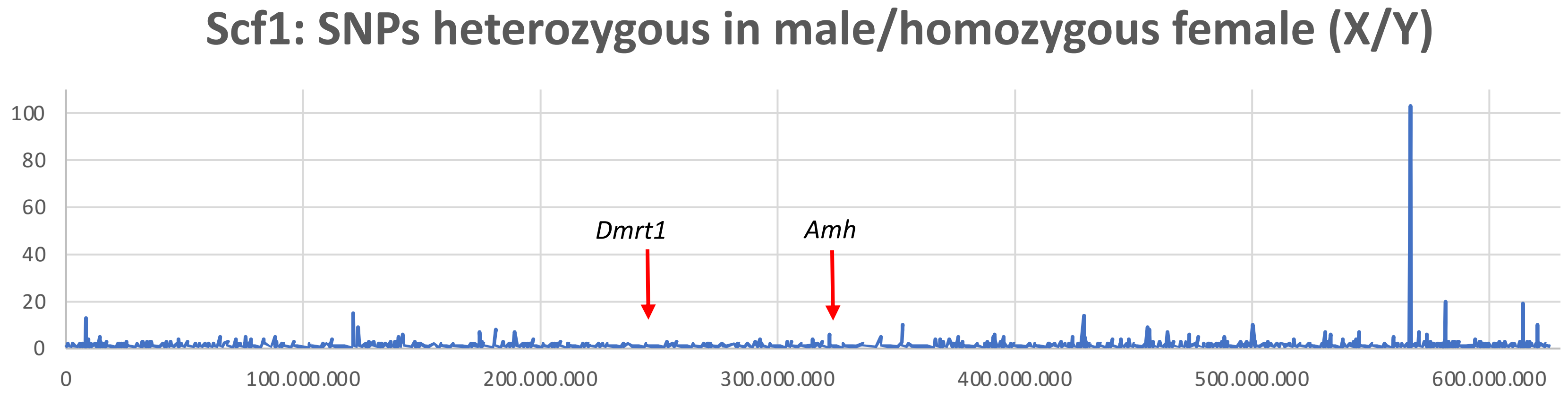


b

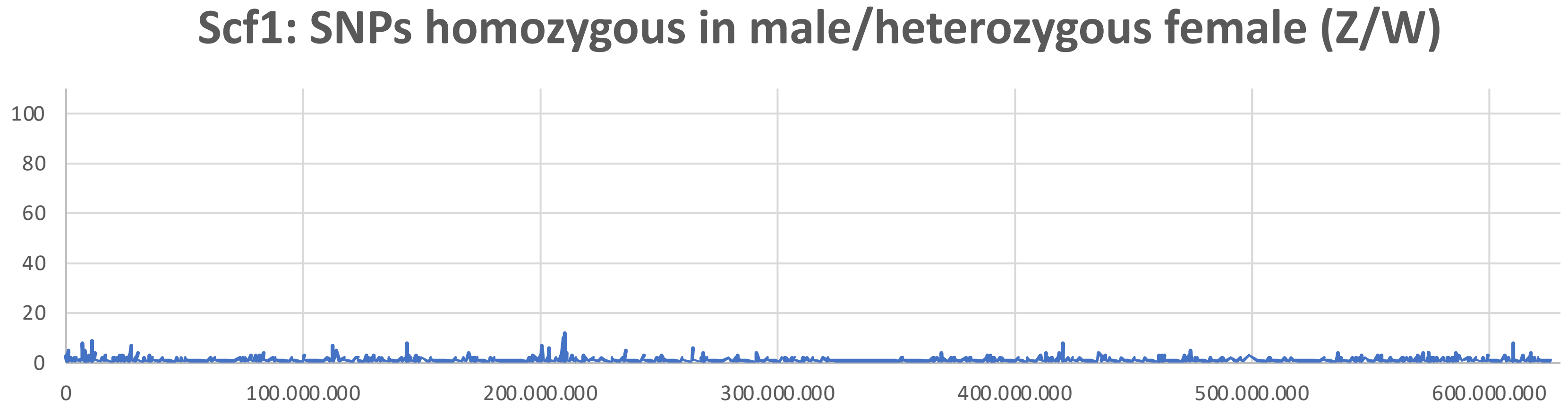

### Supplementary Figure 4

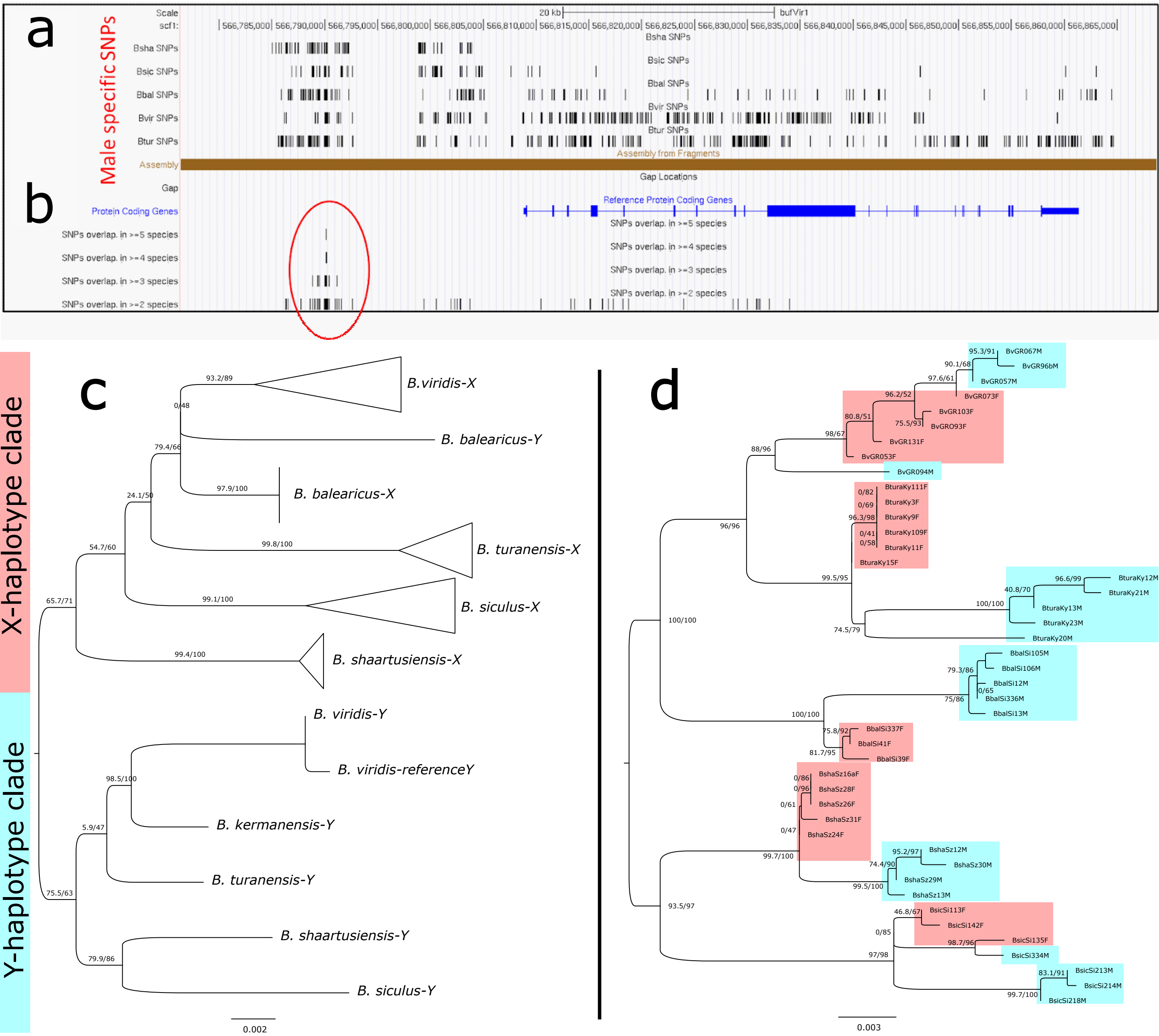

Supplementary Figure 5: Coverage distributions of X/Y haplotypes depending on female/male poolseq in *B. viridis*:

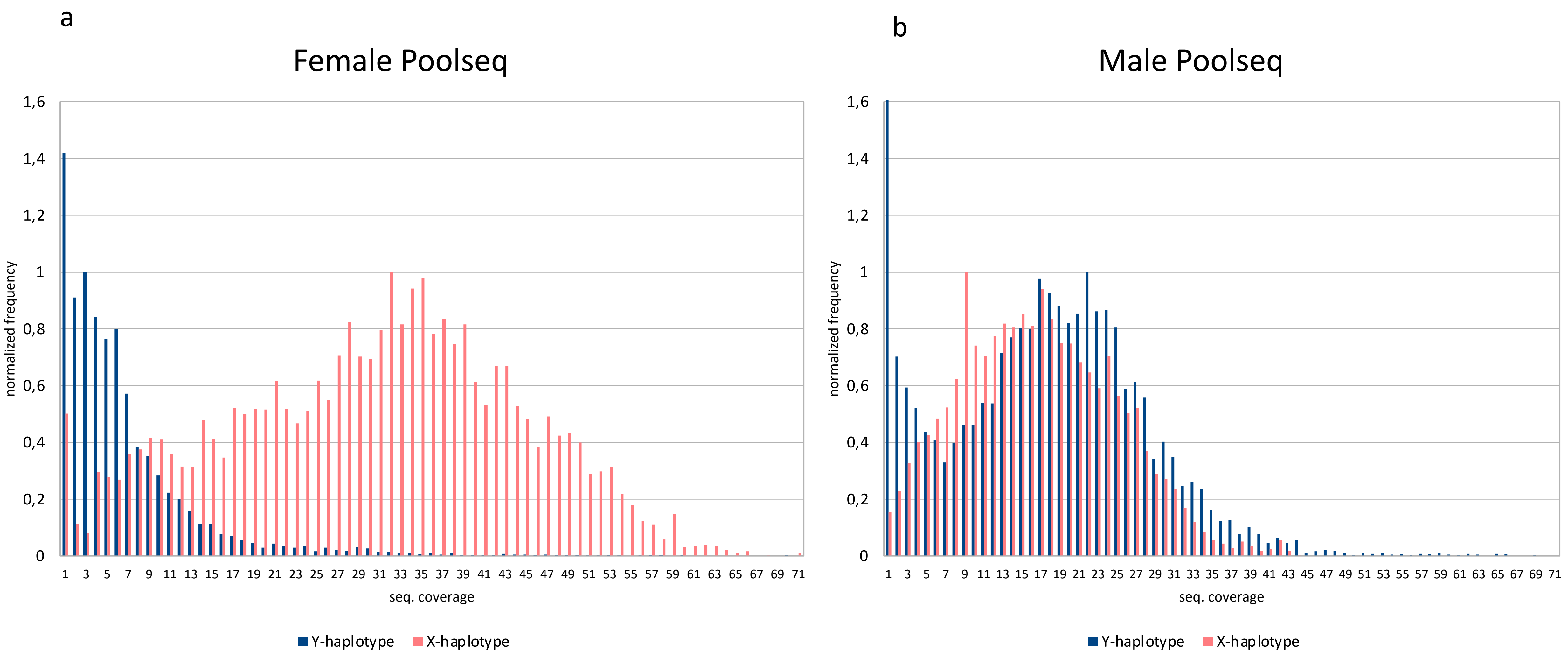

Supplementary Figure 6: Female/male poolseq coverage over haplotypes in *B. viridis*

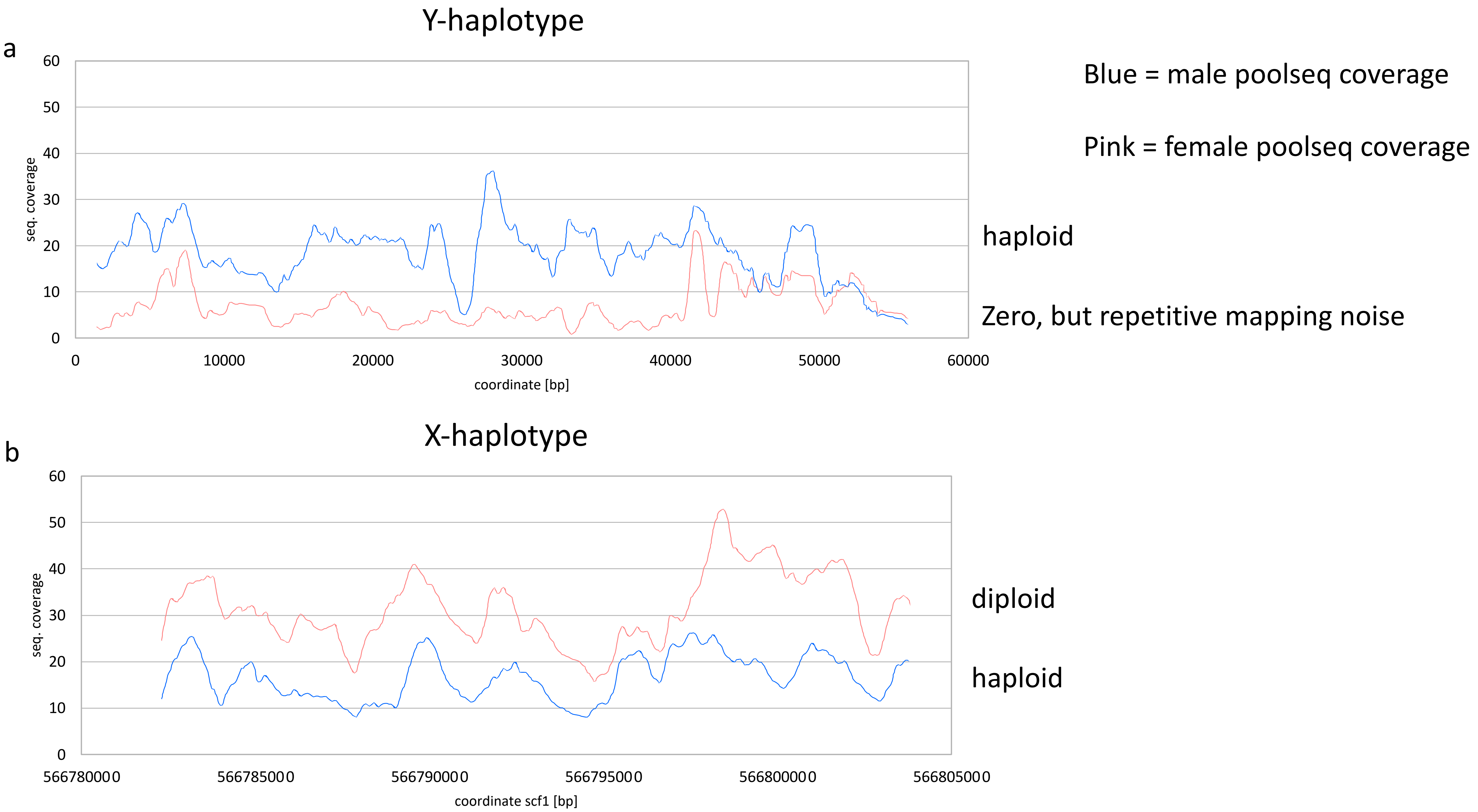

Supplementary Figure 7: X- and Y-haplotype comparisons by dot plots in the *Bod1L*-region

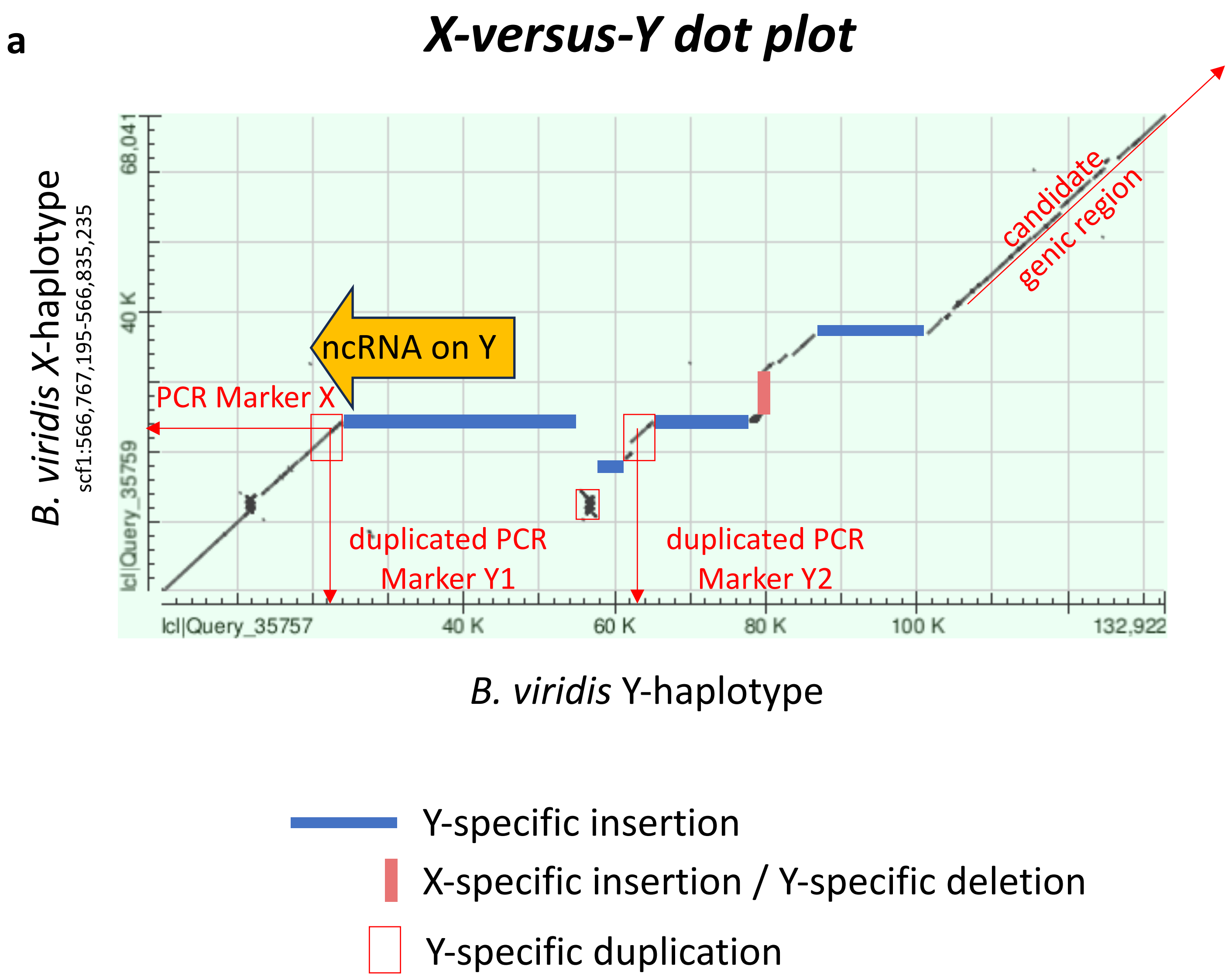

**b**

#### *Y-versus-Y dot plot*

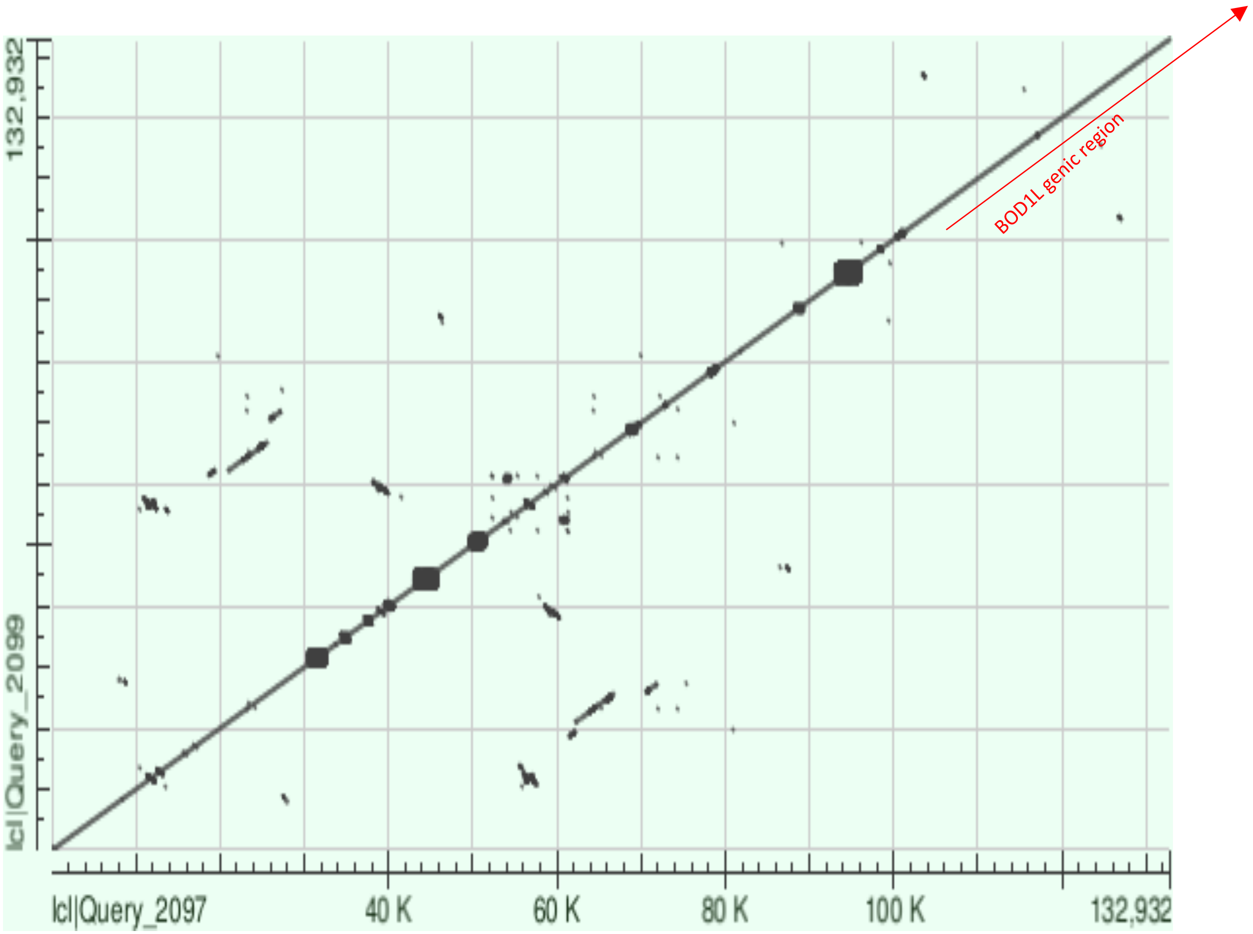


*B. viridis* Y-haplotype

*B. viridis* Y-haplotype

c

#### *X-versus-X dot plot*

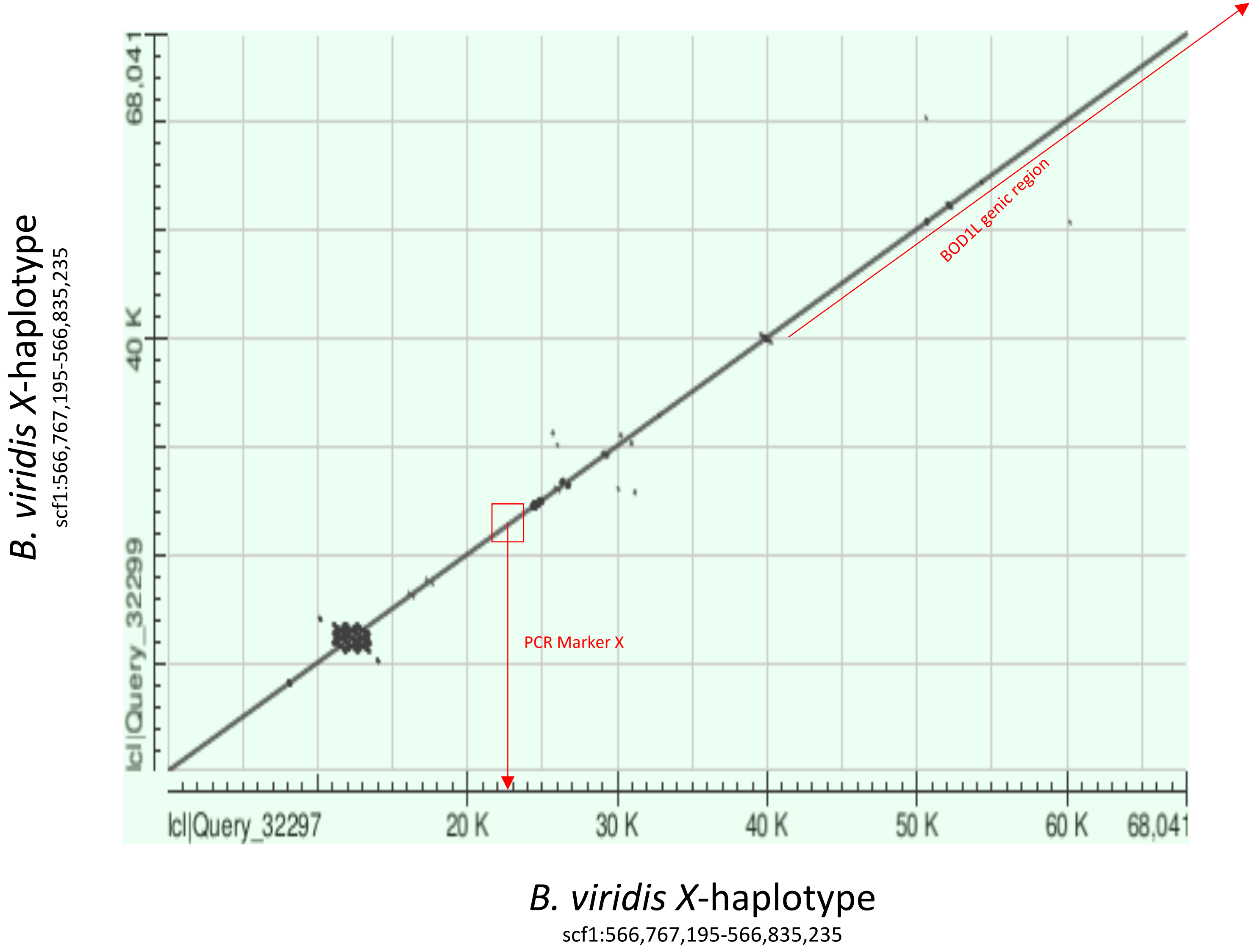

Supplementary Fig. 8

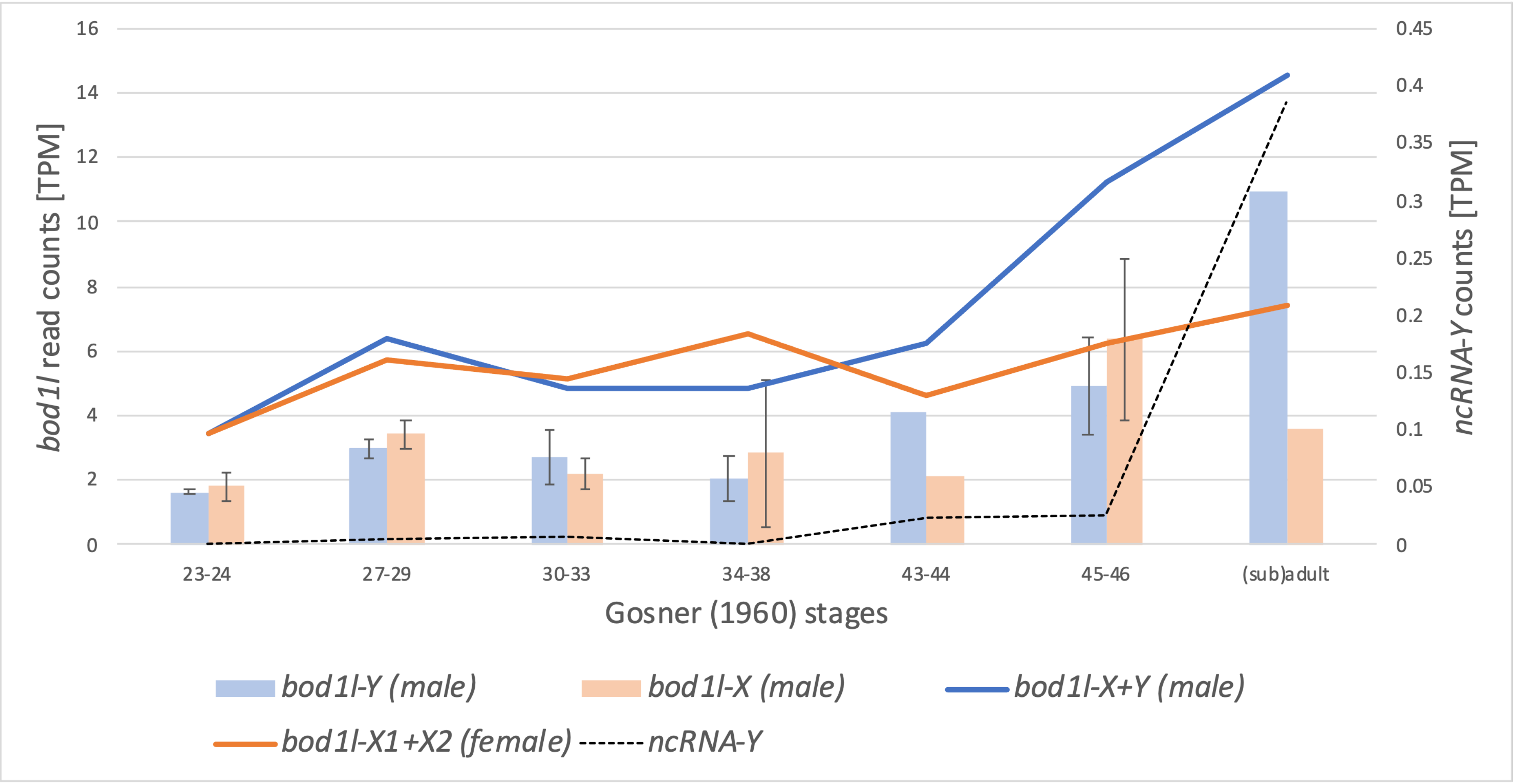

#### Supplementary Figure 9

a

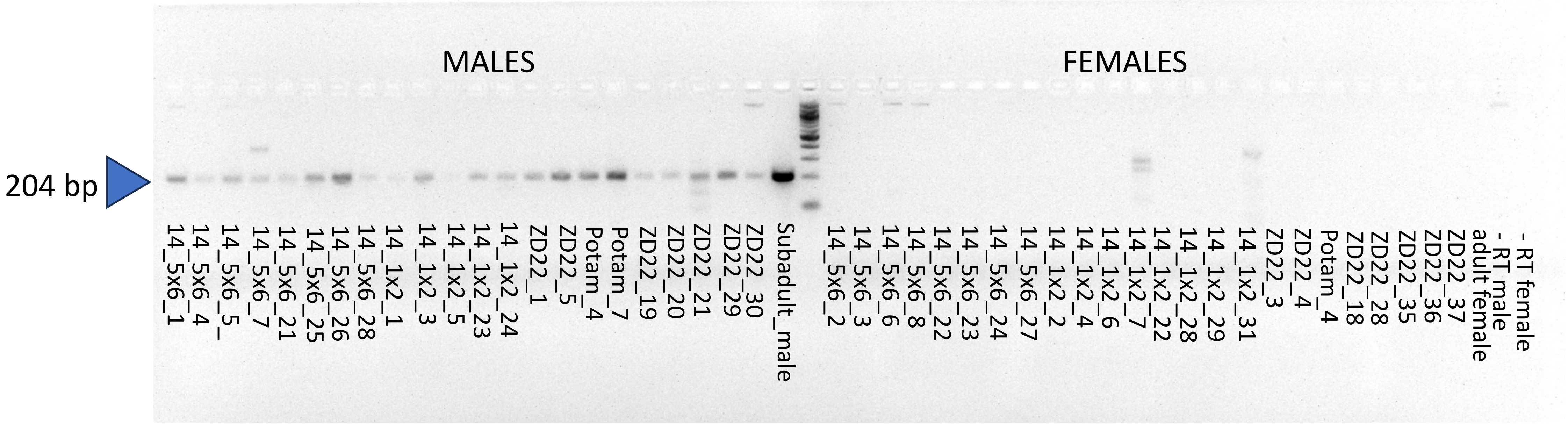


Primers: NcRNAY_Ex1to2_F + NcRNAY_Ex3to2_R

b

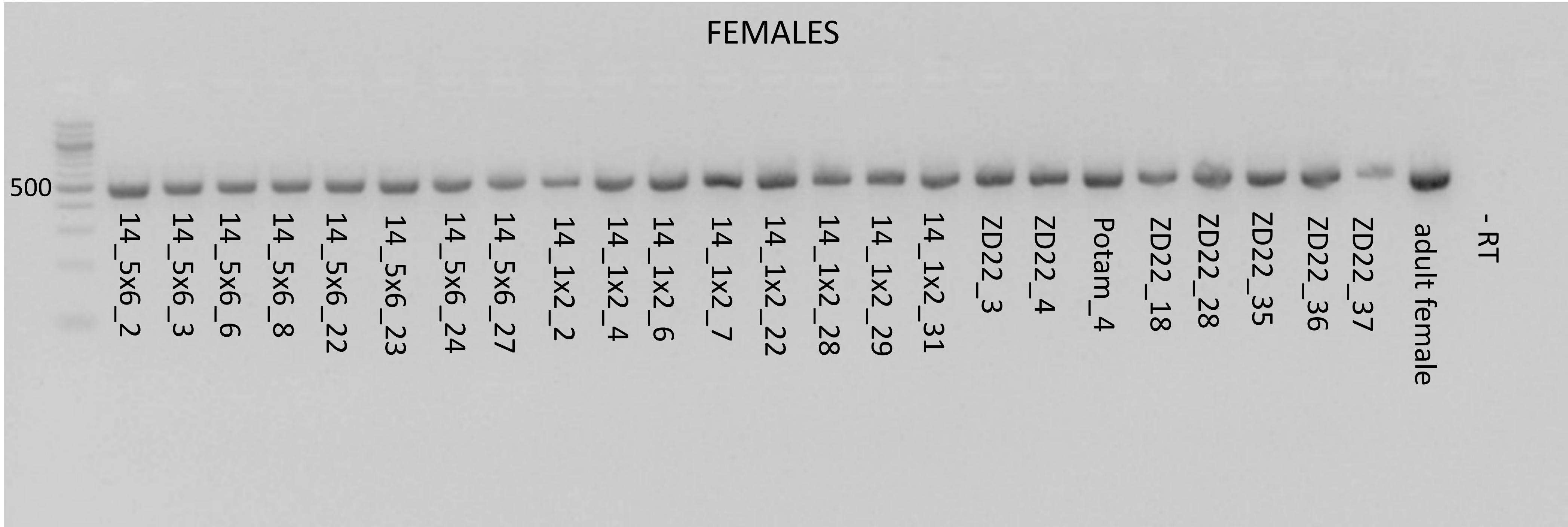


Primers: vasa_1F (5’_AACCAAAAAGAAGGCGGATT_3') and vasa_1R (5’_ATGCCAGGATTGCTGTAACC_3')

### Supplementary Figure 10

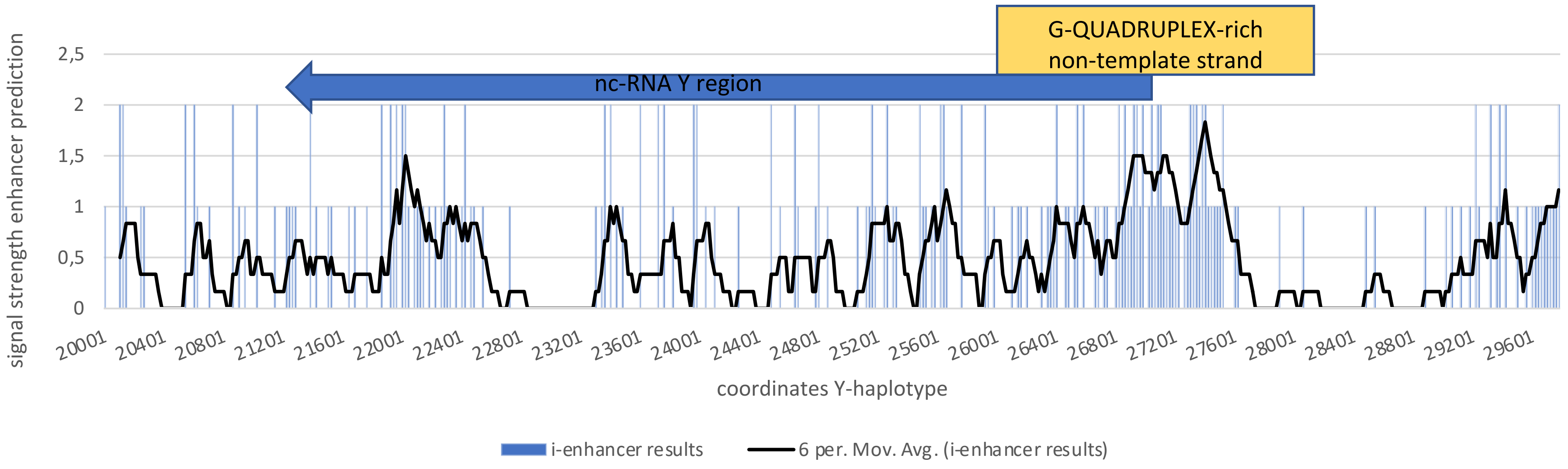

### Supplementary Figure 11

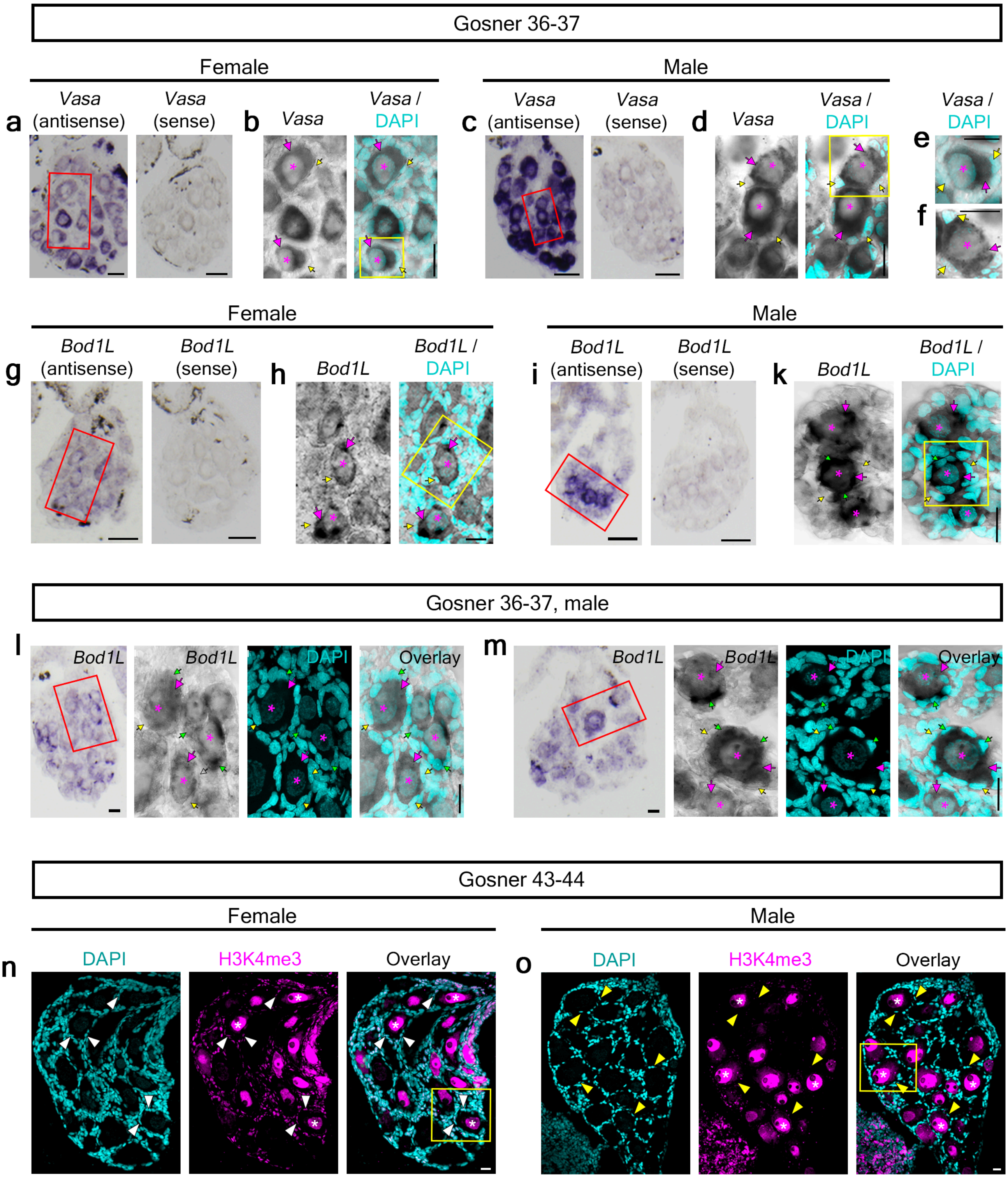
