## Supplementary Text for "A candidate sex determination locus in amphibians, evolved by structural variation between X- and Y-chromosomes"

**Suppelementary Text 1**

**Structural comparison of the gene products from the X- and Y-copy of *Bod1L***

A closer look at the SNPs in the transcriptomes (Supplementary File 5) revealed that two coding SNPs led to non-synonymous changes between the X and the Y copy, specifically scf1:566,831,998 G/A (AA609: Arginine to Lysine) and scf1:566,832,081 T/G (AA637: Serine to Alanine). To test if these amino-acid changes may cause structural and thus functional differences to explain sex-specific action of the X and Y copies of the manually curated *Bod1L* gene model, we evaluated them using the platforms Alphafold and RaptorX. Indeed, folding differences occur, but their predictions remained highly insecure due to the low quality of the alphafold model for these extremely large proteins.

While our results in *B. viridis* could neither confirm nor fully reject that structural and thus functional differences of the *Bod1L*-proteins might play sex determining roles, together with the results of the AmpliSeq-approach from multiple related green toad species, that were not showing any conserved Y-specific coding mutations between the species, we currently consider the X- and Y-specific differences in the coding region not to be of primary importance for sex determination.
